# Computational Analysis of Silent Mutation Effects on SARS-CoV-2 RNA–Host RNA-Binding Protein Interactome

**DOI:** 10.1101/2025.09.22.677528

**Authors:** Adebayo J. Bello, Onyeka S. Chukwudozie, Olajumoke B. Oladapo, Ayomide O. Omotuyi, Nnamdi A. Nzoniwu, Oluwafemi D. Amusa, Bodunrin O. Ottu, Ugochukwu Okeke, Abeebat O. Adewole, Ololade O. Akinnusi, Ayomikun E. Kade, Joseph B. Minari, Joy Okpuzor, Onikepe Folarin, Mujeeb O. Shittu

## Abstract

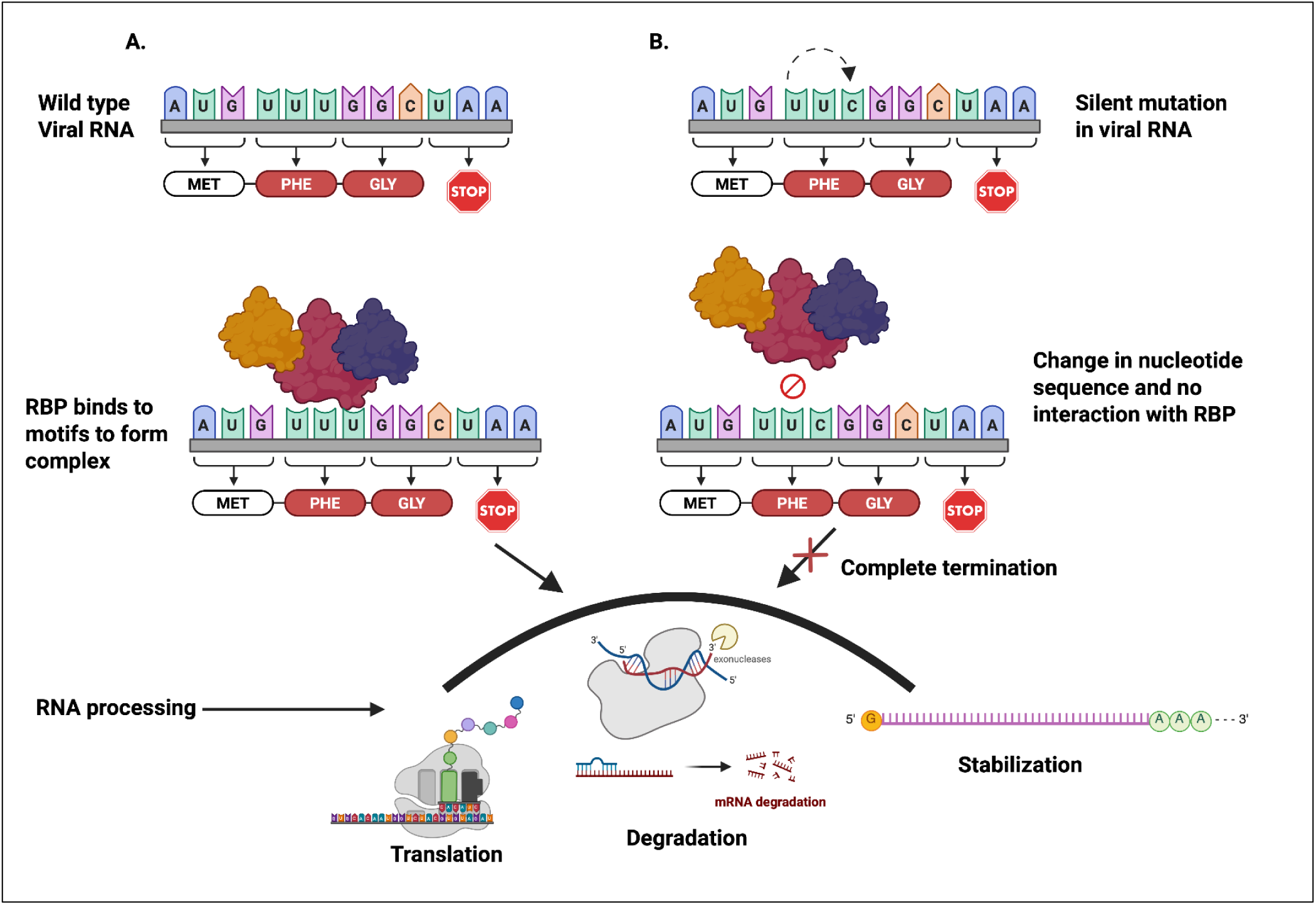

**Abstract:** RNA-Binding proteins (RBPs) play critical roles in host-virus interaction. They facilitate the regulation of viral RNA (vRNA) turnover by recognizing and forming complexes with the vRNA structure via specific RNA motifs-RNA binding domain interaction. However, due to consistent evolving nature of viruses, silent mutations in the viral genome can impact RBP-vRNA binding thereby altering the RNA processing. While efforts have been made in characterizing other forms of mutations leading to changes in amino acids sequence in SARS-CoV-2 variants, details on how silent mutations impact RBP-vRNA interaction remain limited. Here, we use extensive *in silico* mutagenesis to introduce silent mutations in the SARS-CoV-2 genome to generate four different synthetic variants and map the interaction of the variants and the wild-type with a catalogue of human RBPs. Our result shows variation in accumulation and reduction of the RBPs binding motifs in the variants compared to the virus reference sequence on a global scale and at the UTRs. The majority of the RBPs with AU-rich binding motifs are reduced in the variants, while RBPs with mostly GC-rich motifs accumulate more binding positions, suggesting that a single change from U/A to G/C and vice versa can impact RBP- viral interactions. Furthermore, we use structural analysis to show the interaction of the vRNA with PUF60 and KHDRBS3 proteins, two RBPs that have not been previously implicated in SARS-CoV- 2 interactome. Our findings show that loss to the conserved poly(U) in PUF60 binding motifs in some of the variants affects its interaction with the protein at the 5′ end, which may disrupt the function of the protein as an anti-viral RNA regulator. We also predicted the key residues in KHDRBS3 interacting with its binding motif in the wild-type at the 3′ end, while noting that the vRNA structural changes in the variants may contribute to the loss of this interaction. Overall, our predictions contribute to the insights into virus evolution and pathogenicity of potential new variants due to the impact of synonymous changes in the nucleotide sequences on protein-RNA interaction.

## Introduction

Severe Acute Respiratory Syndrome Coronavirus 2 (SARS-CoV-2) is an enveloped, positive-sense, single-stranded RNA virus of approximately 30 kb (Pal et al., 2020). The SARS-CoV-2 genome contains 3′ poly(A) and 5′ cap structures as mRNA in the translation activity of the replicase enzyme (Romano et al., 2020). The SARS-CoV-2 RNA genome is encoded by structural proteins (membrane protein (M), nucleocapsid protein (N), envelope protein (E), and spike protein (S)) for viral assembly, non-structural proteins, and accessory proteins (Brant et al., 2021; Malone et al., 2022). The genome consists of 14 functional open reading frames (ORFs) and two nonstructural proteins (NSPs), which are expressed at the 3′ end of the genomic RNA (Kandwal and Fayne, 2023). The virus is responsible for the COVID-19 disease, which has caused over 7 million deaths globally (WHO, 2025). Despite vaccine development and efforts to curtail the spread of the virus, more studies are needed to understand the virus’s evolution and host-pathogen interaction.

Upon infection of a host cell, the virus deploys a ‘translation-ready’ RNA molecule, which uses the protein synthesis machinery of the host to express a set of proteins crucial for replication (Schmidt et al., 2021). During this process, replication of the full-length viral genome and transcription of subgenomic RNAs both involve the synthesis of negative-strand RNA intermediates (Miller and Koev, 2000; Grellet et al., 2022). In common with other RNA viruses, SARS-CoV-2 depends on effectively engaging the host cell factors as regulators of this process, as well as regulating its RNA stability, localization, translation, and production of progeny (Brant et al., 2021). However, the virus does not have the independent mechanism to synthesize all the proteins required for the processes but hijacks and uses the host cellular RNA-binding proteins (RBPs) to regulate the processes. The host cell, as a response mechanism, uses these regulators to detect the pathogen and initiate appropriate innate immune response pathways against the virus.

RNA-binding proteins (RBPs) bind to RNA to form ribonucleoproteins (RNPs). Over 2,000 classes of RBPs that form an interactome with RNA in its diversity (Gerstberger et al., 2014; Hentze et al., 2018). The RBPs potently and ubiquitously regulate transcripts throughout their life cycle. They do so by binding to sequence and/or structural motifs in RNA via modular combinations of a limited set of structurally well-defined RNA-binding domains (RBDs), such as the RNA recognition motif (RRM) (Cassola et al., 2010), the hnRNP K homology domain (KH)4 (Lunde et al., 2007), or the DEAD box helicase domain (Ryan and Schröder, 2022), and the interaction of the RBDs with RNA motifs facilitates the regulation of genes encoded in a genome. The identification of RNA-binding proteins (RBPs) that bind to the SARS-CoV-2 RNA interactome is crucial for discovering and uncovering the reprogramming of viral gene regulation and the activation of the antiviral immune system in host cells (Mathur et al., 2024). RBP sensing of SARS-CoV-2 RNA can also trigger the cellular antiviral state, which suppresses viral gene expression through the inhibition of protein synthesis and the production of interferons (Kamel et al., 2021). Therefore, cellular RBPs are key regulators of the virus life cycle, either promoting or restricting infection. Several studies have reported the interaction of SARS-CoV-2 RNA with human RBPs. For example, Srivastava et al. (2020) used motif enrichment analysis to show the interaction of SRSFs, PCBPs, ELAVs, and HNRNPs RBPs with the SARS-CoV-2 genome. Lee et al. (2021) used transcriptome analyses and knockdown experiments to uncover 17 and 8 RBPs acting as SARS-CoV-2 antiviral and proviral RBPs, respectively. Multiple studies have also reported that SARS-CoV-2 RNA can interact with both the antiviral and proviral RBPs (Lisy et al., 2021, Koliński et al., 2022). Other experimental assays such as mass spectrometry (ChIRP-MS) (Flynn et al., 2021; Labeau et al., 2022) and RNA antisense purification and quantitative mass spectrometry (RAP-MS) (Schmidt et al., 2021) have identified numerous RBPs interacting with SARS-CoV-2 RNA. DEAD-box helicase 1 (DDX1), an RBP known for RNA processing and repair of DNA damage, can influence replication of the virus genomic RNA by initiating template read-through (Ariumi et al., 2007), while heterogeneous nuclear ribonucleoprotein A1 (hnRNPA1), which acts as pre-mRNA processing, can act as a regulator of the viral RNA synthesis (Kaur and Lal, 2020). Putting all these together, there is strong evidence that the virus can re-engineer the RNA-binding proteins (RBPs) in the hosts’ cell and evade the anti-viral RBPs (Lee et al., 2021).

Due to the consistent evolving nature of coronaviruses, silent mutations (changes to the virus nucleotide sequence without changes to the amino acid sequence) in the virus elements can alter the RNA motifs, thereby affecting the binding of the RBPs. When this happens, the change can alter the viral RNA processing (Horlacher et al., 2023). One of the analyses that has succeeded in the implementation of the pathogenesis and lineage tracking is the silent synonymous mutations that have been detected in the SARS-CoV-2 protein variants (Hossain et al., 2021). In their report, 37.2% of the total 3334545 mutations identified from 259044 SARS-CoV-2 isolates were silent mutations, while others were missense mutations and extragenic single nucleotide polymorphisms. Notably, they found that F106F in the NSP3 region may affect the mRNA processing and subsequently the viral protein. Pachetti et al. (2020) identified, out of a random collection of 220 SARS-CoV-2 genomes on the GISAID database, that 3 out of the 12 most frequent mutations at positions 3036, 8782, and 18060 were silent mutations. However, reports on how the silent mutations affect the binding motifs for RBPs are lacking.

Genetic recombination has been associated with the emergence of novel SARS-CoV-2 strains, which can alter the RBP binding motifs or introduce new motifs into the viral genome, potentially affecting the properties of the viral RNA-protein interaction and subsequently RNA processing (Haddad et al., 2022, Syed et al., 2024). This recombination often leads to changes in the amino acid sequence, leading to new variants. However, despite the vast analysis of the SARS-CoV-2- RBP interactome (Ferrarini et al., 2021, Kamel et al., 2021, Enguita et al., 2022, Labeau et al., 2022, Lee et al., 2021, Wu et al., 2024), there are limited reports on the impact of silent mutation on the virus-host RBP interaction. While Ziesel and Jabbari, (2025) predicted the impact of these mutations on the RNA secondary structure and stability of six Variants of Concern, Horlacher et al. (2023) used deep learning models, pysster and DeepRiPe that were trained on CLIP-seq data to identify some loss and gain of RBPs binding positions at nucleotide resolution across seven human coronaviruses, and identified MBNL1, FTO, and FXR2 RBPs as probable clinical biomarkers (Horlacher et al., 2023). However, significant gaps persist in elucidating how silent mutations can create new SARS-CoV-2 RNA variants and contribute to their pathogenicity through interactions with host RBPs.

Here, we performed extensive *in silico* mutagenesis across the whole SARS-CoV-2 genome without changing the amino acid composition. Then, we used RBPmap (Paz et al., 2014) to map 132 known human RBPs on 4 synthetic variants of the virus. By predicting and mapping the RNA-RBP interaction, we identified hundreds of loss or gain of binding positions for different RBPs. Using computational and structural analysis, we provide, to the best of our knowledge, how changes in RBP binding motifs affect the interaction of PUF60 and KHDRBS3 at the 5’ and 3’ of the SARS-CoV- 2 genome, respectively. Our study on the impact of silent mutations within the RNA-RNA binding protein interactome of SARS-CoV-2 is important in the predictive mapping of RNA-binding protein (RBP) binding sites, offering insights into the intricate interactions within cellular systems and unveiling pivotal insights into viral evolution and pathogenicity.

## Results

To perform *in silico* probing of RBPs binding to the genome, we first introduced silent mutations in the genome of SAR-CoV-2 reference sequence (29,903 bp) without altering the amino acid sequence of the global transcripts. To ensure this process, we generated four different variants of the SARS-CoV-2 genome using four publicly available codon optimization tools: JCat (VT1), Vector Builder (VT2), IDT (VT3), and GeneArt (VT4) and renamed them accordingly (**Suppl file**). To avoid errors during the optimization process, we partitioned entire genome into 10 sets (each containing at most 3000 bp). The sequences were codon optimized for human/Homo sapiens as indicated by each tool, since SAR-CoV-2 infects human cells. Then we performed multiple sequence alignment and generated a phylogenetic tree for the wild-type genome (WT) and 5′ and 3′ regions to show the relationship between the virus reference genome and the new variants (**Suppl Figure 1**). 5′ and 3′ have been shown to play key roles in pro- or anti-viral defense, and evolutionary changes in this region may contribute to the diversity of a virus (Farkas et al., 2021). Here, our results showed that VT3 is closely related to WT across the whole genome, 5′ and 3′ regions. Variant 1 shows a closer relation to VT2 across the entire genome and at the 5’ (**Suppl Fig 1A and B)** but differs at the 3’ region with a close relation to VT4 (**Suppl Figure 1C**). We also determined the effect of the mutations on changes in the percentage of nucleotides present in the WT and the variants. There are similarities in the percentage of the nucleotides present in the whole genome and the 3′ UTR. In both, the pattern shows that adenine and uracil have a higher percentage compared to other variants, while guanine and cytosine show the highest in VT1 and VT4, respectively (Figures **1A and C**). Notably in the 5′ UTR (**Figure 1B**), there is a sharp increase in cytosine in WT, which suggests the nucleotide is more abundant in the region. Whereas in VT1, the mutation causes an increase in uracil compared to its abundance in the whole genome and at the 3′ region. Generally, the decrease or increase in the percentage of nucleotides across the variants suggests these silent mutations could cause rearrangement of the protein-binding motifs on the SARS-CoV-2 RNA, thereby leading to loss or gain of protein-RNA interaction.

**Figure 1.**
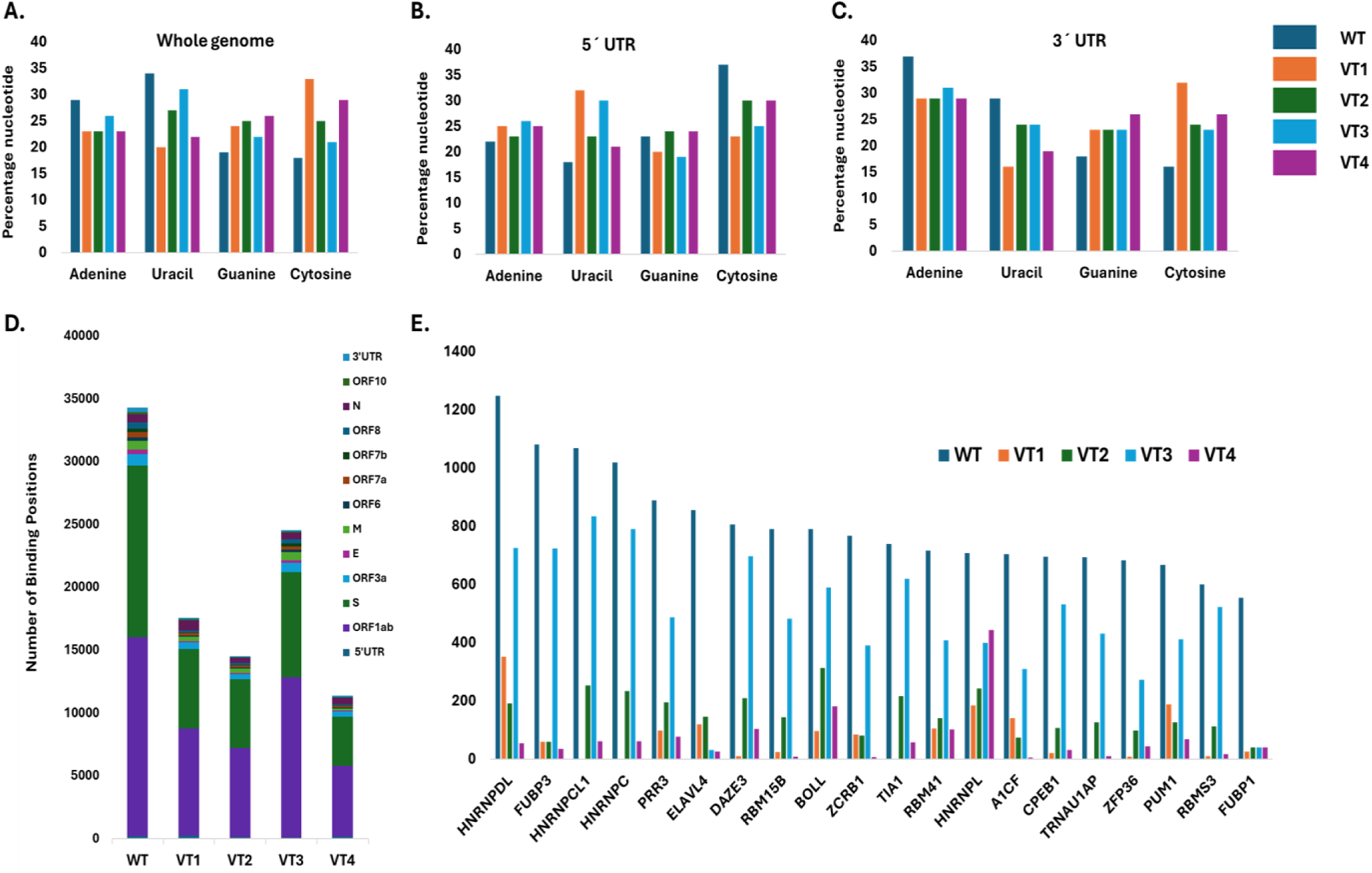
Impact of induced silent mutations on the SARS-CoV-2 genomic regions. Percentage of nucleotides for the **A.** Whole genome, **B.** 5’ UTR, and **C.** 3’ UTR sequences. **D**. Accumulation of loss or gain of binding positions in each genomic element of the wild type and the synthetic variants **E.** Accumulation of changes in binding positions in the variants compared to the wild type for the top 10 RBPs.

Next, we assessed the changes in the binding positions that accumulated in different regions of the viral genome due to the silent mutations. There is a clear difference in the total number of binding positions in the WT compared to the variants. Most mutations are noticeable in ORF1ab and the spike region (**Figure 1D**). Interestingly, VT3 harbours considerable changes in binding positions but is similar to the WT, suggesting a close similarity between the two genomic sequences. **Figure 1E** represents the top 20 RBPs with the most accumulated changes to binding positions across the entire genome. Like **Figure 1D**, the number of changes in VT3 presents a close similarity to the WT compared to other variants, although with reduced binding positions. However, the number of binding positions in different variants, especially VT4, reduced significantly, suggesting a drastic shift in RBP-RNA interactions could lead to altered RNA processing.

### Catalogue of predicted human RBPs interacting with the SARS-CoV-2 genome

New variants of the SARS-CoV-2 virus are emerging with new features due to the newly acquired sequence or sequence arrangement, making them important variants to alter the RNA process inside the host. To investigate the extent of silent mutation on the interaction of RBPs with the RNA in the entire genome and the UTRs, we used the full 131 human RBP catalogue on RBPmap to construct a comprehensive RBP interaction with the SARS-CoV-2 reference genome and the induced variants by in-silico mutagenesis (supplementary file). With some degree of variation, our prediction shows clustering of RBPs with similar function and motif recognition mechanisms. For example, SFSF5, SRSF7, MBNL1, RMB24, RMB6, and RMB45, which are involved in RNA splicing, show an increase in the number of RNA motif recognitions in VT1, VT2, and VT4 compared to the WT and are clustered together in the whole genome (**Figure 2A**). Likewise, YXB1, XYB2, SRSF2, SRSF4, and MBNL1 with similar RNA splicing functions are also clustered together in VT1, VT2, and VT4 at the 5′ UTR. The RBPs show a reduction in their recognition motifs in VT3 compared to the WT, apart from MBNL1, with an increase in recognition motifs across all the variants (**Figure 2B**). In the 3′ UTR, PAPBC1 and PAPBC4, known for mRNA stability regulation and translation, cluster with splicing factor SART1 and show a significant increase in binding positions in VT2 and VT3 compared to the WT (**Figure 2C**), suggesting that nucleotide changes in the region could lead to a pro- or anti-viral response depending on the favourable RBP binding motif involved.

**Figure 2.**
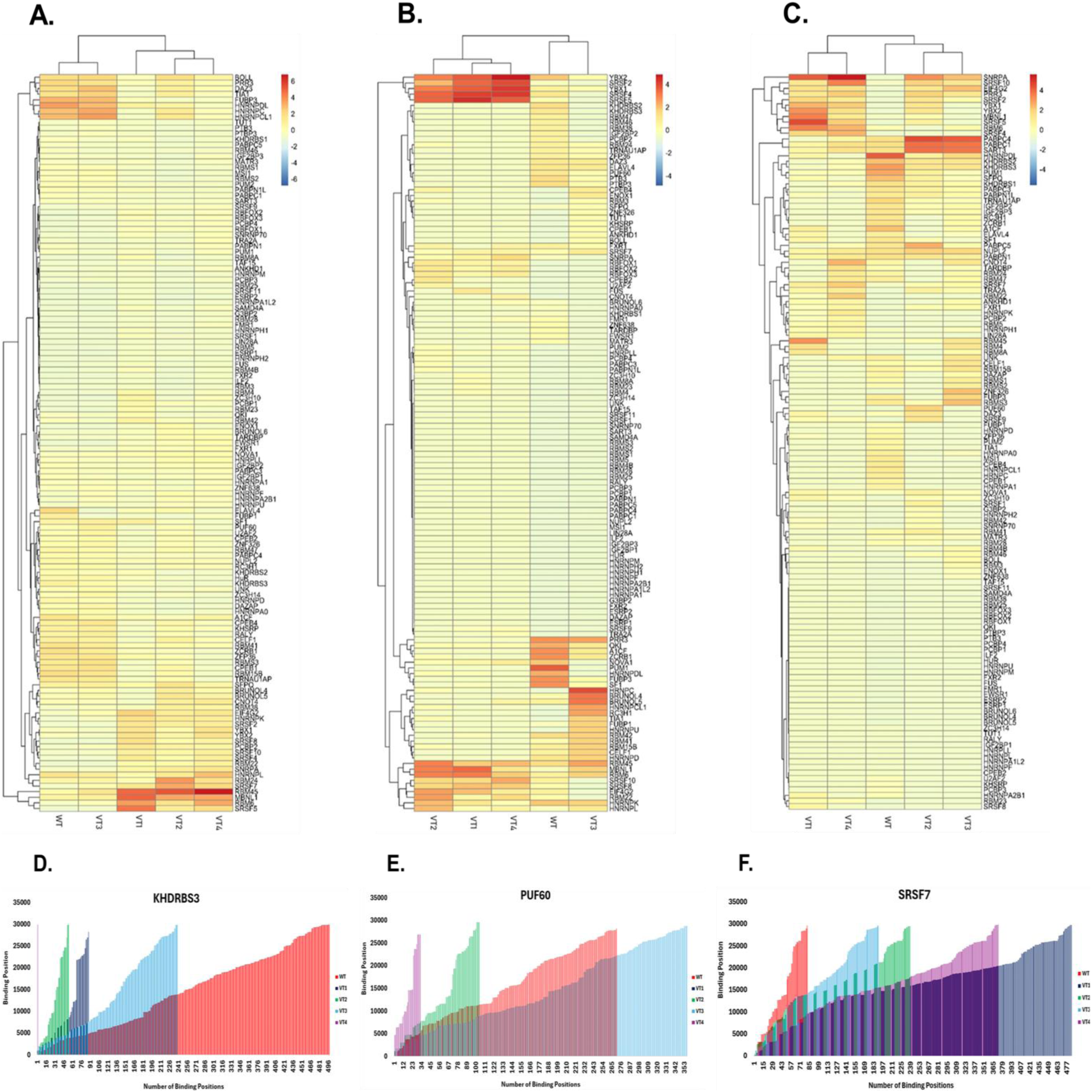
SARS-CoV-2 RNA distribution and interactome showing clustering of the RBPs based on the binding positions. **A.** whole genome, **B.** 5’ UTR, and **C.** 3’ UTR. Clustering analysis was applied to evaluate the reference genome and variants in terms of binding positions. RBPs with higher numbers of binding positions are represented in red color, while the color intensity decreases with a decrease in the binding positions. Binding positions of the **D.** KHDRBS1, **E.** PUF60, and **F.** SRSF7 RBPs relative to the total number of binding positions in each variant are represented.

Next, our assessment of the total number of binding positions of each RBP motif shows variation in the accumulation and reduction of the binding positions in the variants compared to the wild type (**Suppl file**). For example, KHDRBS3 was predicted to have 498 binding positions in the WT whole genome, compared to 88, 54, and 239 in VT1, VT2, and VT3, respectively (**Figure 2D**). In VT4, the RBP has only one binding position at A29878. Although the binding position is present in the WT at the poly(A) region of the 3′ UTR, losing 497 binding positions in the global genome could play a key role in altering the KHDRBS3-RNA interaction and processing. Similarly, the total number of binding positions for PUF60 decreases from 271 in WT to 104 and 32 in VT2 and VT4, respectively, while it increases to 356 in VT3 (**Figure 2E**). The binding positions were entirely lost in VT1, whereas the U47, U58, U63, and U83 at the 5’ UTR were lost in all the variants. The number of binding positions increased from 82 in the WT. to 484, 238, 190, and 372 in VT1, VT2, VT3, and VT4, respectively (**Figure 2F**), suggesting an increase in the interaction of SRSF7 with a silent mutated genome of the SARS-CoV-2.

### RBPs with loss and gain of binding positions

With previous reports showing that UTRs of viral genome may not be under pressure to translate into proteins, they could become more prone to mutations that would exert changes in the regulatory pattern of the virus by host binding proteins (Jaafar and Kieft, 2019). We then put together the top 10 RBPs with loss or gain of binding positions at the UTRs. On the top of our list in **Table 1** is PUM1, a regulator of mRNA in different cellular processes. Overexpression of PUM1 was reported to suppress RNA replication in cells infected with Newcastle disease virus (Narita et al., 2014). From our prediction, we identified ten different binding positions of PUM1 in WT, and surprisingly, all the positions were lost in all the variants. Our findings suggest that in the event of silent mutation in the 5′ UTR of the SARS-CoV-2 virus leading to new variants similar to our predicted variants, PUM1 recognition motifs may be lost, which could cause proliferation of the virus in infected cells. Similarly, the recognition motifs for FUBP3 and A1CF are lost in all the variants except VT3, which has a close relation to WT. Nevertheless, the number of binding positions reduce from 8 to 1 and 2 for FUBP3 and A1CF, respectively.

**Table 1:**
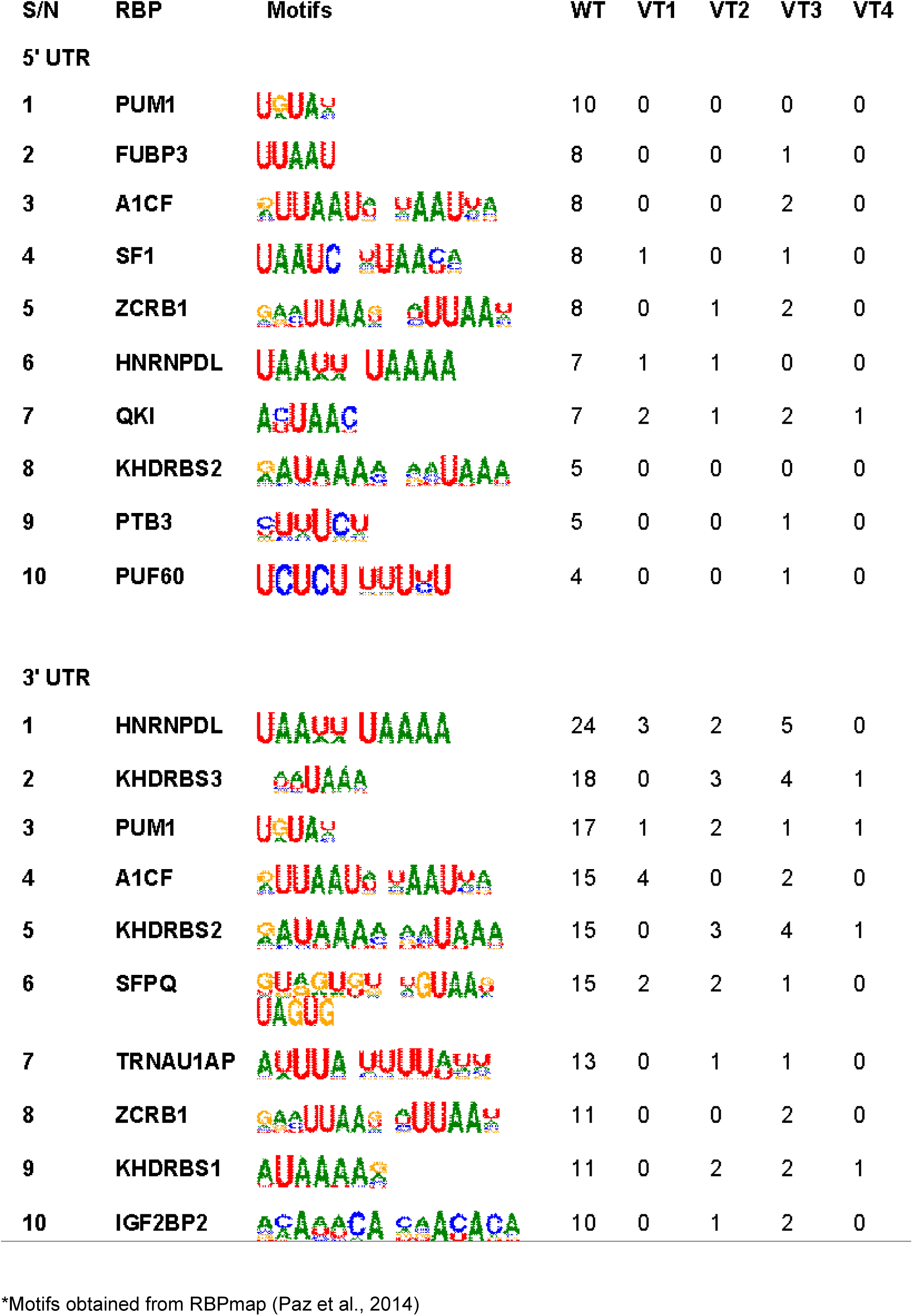
Top 10 RNA-binding proteins with loss of binding motif* positions at 5’ and 3’ UTRs.

Poly(U) binding splicing factor 60, also known as PUF60, is a regulator of RNA turnover when overexpressed in cells infected with hepatitis virus (Sun et al., 2017). In our study, the binding position of the protein recognition motif was reduced from 4 in WT to 1 in VT3 but lost in other variants (**Table 1**). At the 3′ UTR, we noticed that PUM1 and A1CF also have high binding positions in WT but were reduced significantly in the variants. KHDRBS3, another splicing factor that was identified to play a key role in the successful completion of the sarcoma-associated herpesvirus (KSHV) life cycle (Lee et al. 2025), shows a reduction in binding positions in the variants. The protein functions by reactivating the KSHV lytic machinery and replication by influencing the viral gene expression, making it an attractive antiviral target (Lee et al., 2025). Notably, VT1 has no binding site for KHDRBS3 compared to 18 in the WT 3′ UTR (**Table 1**), suggesting less influence of the RBP as a proviral factor on the variants.

In **Table 2**, we show the top 10 RBPs with significant increases in their binding positions at the UTRs. In the 5′ UTR, the binding positions for MBNL1 increase from 1 (WT) to 16 (VT1), 7 (VT2), 3 (VT3), and 5 (VT4). Surprisingly, in VT3, SRSF2, SRSF4, SRSF5, SRSF8, SRSF10, and EIF4G2 lost their binding positions but increase in all other variants. YXB1 and YXB2 also decrease to 1 from 3 and 6, respectively, in VT3 but increase across other variants. Nevertheless, VT1 and VT4 present the most significant increase in the binding positions for the RBPs compared to VT2 and VT3. On the other hand, SNRPA, SRSF2, SRSF5, YXB1, YXB2, EIF4G2, and PRR2 have no binding positions in the WT in the 3′ UTR. However, an increase in the binding position was observed in other variants for the RBPs apart from SRSF5, YXB1, and YXB2 with no binding positions in VT3. Similarly, MBNL2 has no binding position in VT3 but increase from 1 (WT and VT2) to 12 (VT1) and 3 (VT4). Likewise, the RBM45 binding position retains only 1 binding position in VT2 but increase to 11 (VT1) and 4 (VT3) and is lost in VT4.

**Table 2:**
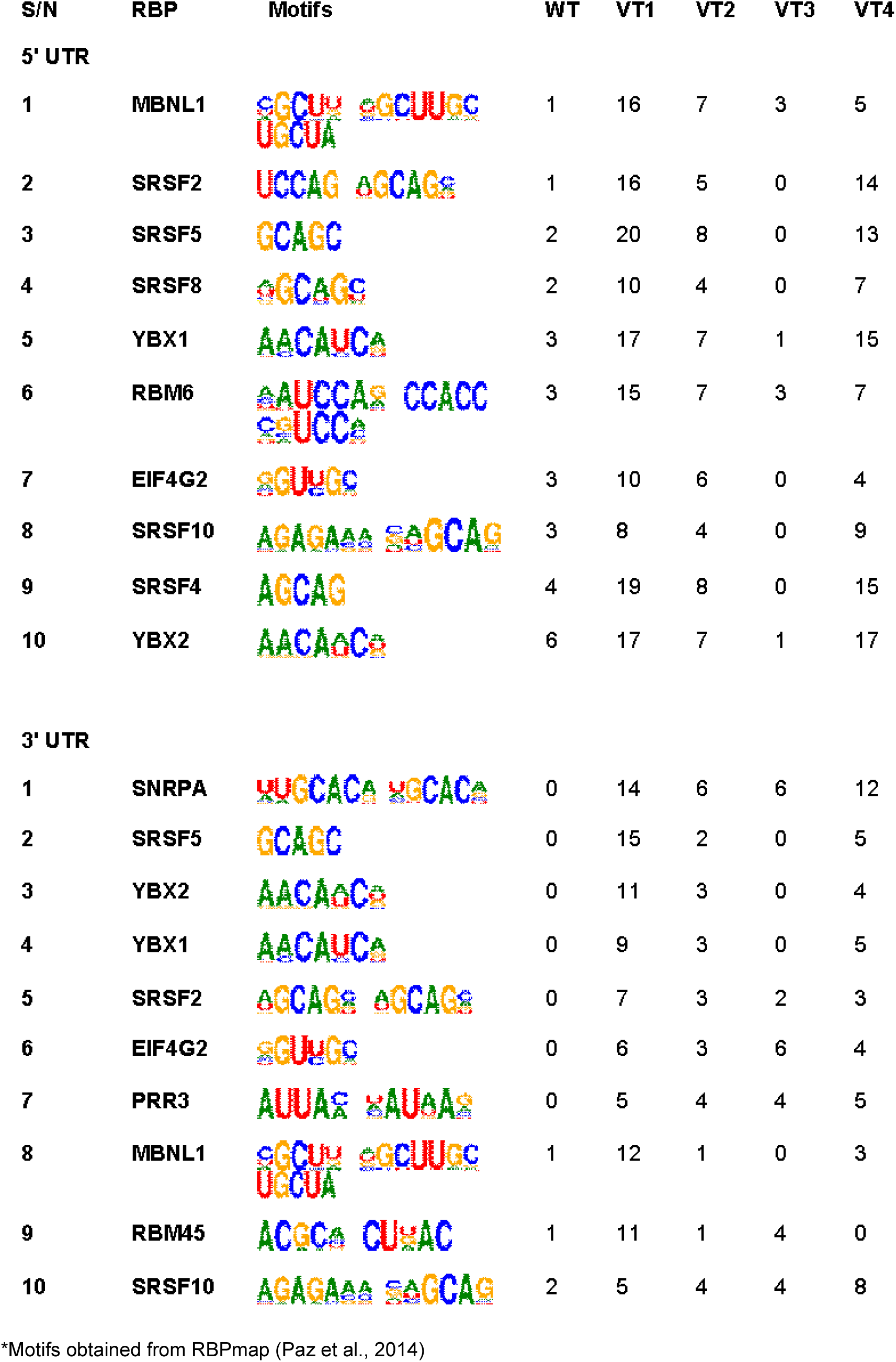
Top 10 RNA-binding proteins with gain of binding motif* positions at 5’ and 3’ UTRs.

| S/N | RBP | Motifs | WT | VT1 | VT2 | VT3 | VT4 |
| --- | --- | --- | --- | --- | --- | --- | --- |
| **5' UTR** |  |  |  |  |  |  |  |
| 1 | MBNL1 |  | 1 | 16 | 7 | 3 | 5 |
| 2 | SRSF2 |  | 1 | 16 | 5 | 0 | 14 |
| 3 | SRSF5 |  | 2 | 20 | 8 | 0 | 13 |
| 4 | SRSF8 |  | 2 | 10 | 4 | 0 | 7 |
| 5 | YBX1 |  | 3 | 17 | 7 | 1 | 15 |
| 6 | RBM6 |  | 3 | 15 | 7 | 3 | 7 |
| 7 | EIF4G2 |  | 3 | 10 | 6 | 0 | 4 |
| 8 | SRSF10 |  | 3 | 8 | 4 | 0 | 9 |
| 9 | SRSF4 |  | 4 | 19 | 8 | 0 | 15 |
| 10 | YBX2 |  | 6 | 17 | 7 | 1 | 17 |
| **3' UTR** |  |  |  |  |  |  |  |
| 1 | SNRPA |  | 0 | 14 | 6 | 6 | 12 |
| 2 | SRSF5 |  | 0 | 15 | 2 | 0 | 5 |
| 3 | YBX2 |  | 0 | 11 | 3 | 0 | 4 |
| 4 | YBX1 |  | 0 | 9 | 3 | 0 | 5 |
| 5 | SRSF2 |  | 0 | 7 | 3 | 2 | 3 |
| 6 | EIF4G2 |  | 0 | 6 | 3 | 6 | 4 |
| 7 | PRR3 |  | 0 | 5 | 4 | 4 | 5 |
| 8 | MBNL1 |  | 1 | 12 | 1 | 0 | 3 |
| 9 | RBM45 |  | 1 | 11 | 1 | 4 | 0 |
| 10 | SRSF10 |  | 2 | 5 | 4 | 4 | 8 |

\*Motifs obtained from RBPmap (Paz et al., 2014)

The EIF4G2 is a key player in translation initiation in eukaryotes. It works as a scaffold protein by recruiting ribosomal subunits and bypassing upstream open reading frames to allow scanning and translation continuation of downstream genes by the ribosome (Smirnova et al. 2022, Meril et al. 2024). Meanwhile, viruses that depend on internal ribosome entry site (IRES)-mediated translation can hijack and recruit EIF4G2 to promote their mRNA translation (Liu et al. 2023), while some viruses such as poliovirus secrete proteases to destroy and inhibit the function of EIF4G2 in synthesizing host protein, thereby promoting their protein synthesis (Gradi et al. 1998). However, as noticed in our findings, silent mutations leading to an increase in the number of binding positions at both UTRs for EIF4G2 may significantly suppress host protein translation and increase the stability and synthesis of the viral proteins.

### Loss of binding motifs on 5’ UTR

Next, we used a structural approach to investigate the loss of RNA-binding motifs across all the variants compared to the wild-type 5’ UTR. Given SARS-CoV-2’s dependence on RBPs for replication and transcriptional regulation, we focused on the interactions between the PUF60 splicing factor and RNA nucleotide bases to elucidate how these RNA variants lack the essential pyrimidine motifs that PUF60 recognizes, thereby impeding viral RNA processing. The wild-type 5’ UTR and PUF60 protein’s structural conformation revealed that PUF60 occupies the central region of the RNA, providing a conformational balance for both ends of the RNA. Based on structural visualization, the central area, where PUF60 resides, exhibits an enriched pyrimidine signature, comprising uridine-rich motifs that play a crucial role in safeguarding against viral RNA manipulation. The wild-type RNA displayed a high content of pyrimidines, predominantly uracil, establishing most contacts with PUF60 out of the 23 hydrogen bonds formed (Figures **3A**, **4A, 3F, Table 3**). Different atomic subdomains of uracil formed hydrogen bonds with residues from PUF 60. U89, U137, U168, U184, and U212 mediate hydrogen bond contacts. More hydrogen bond contacts on uracil were found around U137, with arginine (R551) atomic subdomains interacting with the U137 (O4’) and U212 (O2) atoms on the side chain and U137 (OP1) on the backbone of the base.

**Figure 3:**
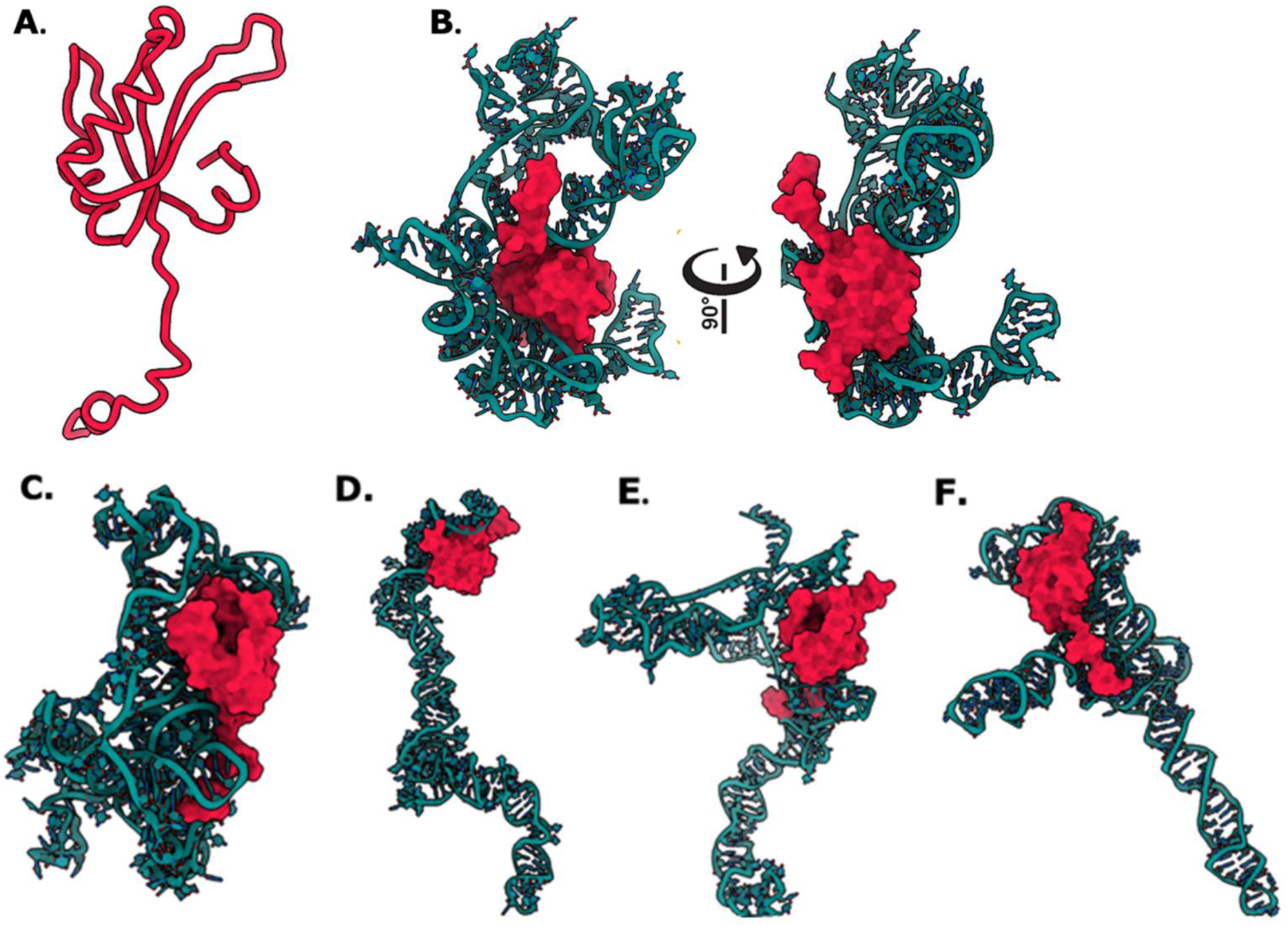
Different computationally induced variants of the SARS-CoV-2 RNA in complex with the PUF 60 protein. **The A**. structure of PUF 60 protein (PDB: 3US5) is fully represented in licorice B. The wild type of 5’ UTR and PUF60 protein complex: PUF60 resides at the center of the RNA structure, containing the needed motif for its recognition. The RNA is represented in teal color and the PUF60 in crimson. **C**. VT1 variant: PUF 60 binds to the end of the RNA structure. **D**. VT2 variant: PUF 60 binds at the terminal end of the RNA loop before the stem at the terminal. **E**. VT3 variant **F**. VT4 variant.

**Table 3:** Interactions between PUF60 and 5’ UTR of the wild-type and the induced variants.

|  | PUF60 | Dist. [Å] | RNA |
| --- | --- | --- | --- |
| **Wild-type 5'UTR** |  |  |  |
| 1 | ASN 468[ ND2] | 3.74 | A 142[ OP1] |
| 2 | ASN 468[ ND2] | 3.12 | G 124[ O2'] |
| 3 | SER 523[ OG ] | 3.50 | C 193[ O2'] |
| 4 | HIS 526[ ND1] | 3.10 | U 168[ O3'] |
| 5 | LYS 527[ NZ ] | 2.94 | U 212[ O2 ] |
| 6 | GLN 530[ NE2] | 3.31 | G 213[ O3'] |
| 7 | GLN 530[ NE2] | 2.30 | C 214[ OP1] |
| 8 | ASN 533[ ND2] | 3.33 | A 139[ O2'] |
| 9 | GLY 534[ N ] | 2.45 | C 140[ O2'] |
| 10 | ARG 535[ NH1] | 3.83 | G 185[ O2'] |
| 11 | ARG 535[ NH1] | 2.34 | G 185[ O3'] |
| 12 | ARG 535[ NH2] | 2.75 | C 186[ OP1] |
| 13 | GLY 539[ N ] | 2.91 | U 89[ O4'] |
| 14 | ARG 540[ NH1] | 2.16 | A 158[ OP1] |
| 15 | ARG 540[ NH1] | 3.80 | C 123[O2] |
| 16 | ARG 540[ NH1] | 3.21 | C 123[ O2'] |
| 17 | ARG 540[ NH2] | 3.52 | C 123[ O2'] |
| 18 | ARG 540[ NH2] | 3.90 | A 143[ OP1] |
| 19 | LYS 541[NZ] | 3.68 | U 184[ O2'] |
| 20 | ARG 551[ NH1] | 2.05 | U 137[ O4'] |
| 21 | SER 558[OG] | 3.66 | U 137[ OP1] |
| 22 | GLY 539[O] | 3.85 | G 124[N2] |
| 23 | GLN 530[OE1] | 3.16 | C 140[N4] |
| 24 | SER 523[O] | 3.90 | U 168[ N3] |
| **VT1** |  |  |  |
| 1 | GLN 459[N] | 3.57 | C 165[O3'] |
| 2 | SER 461[N] | 2.90 | G 199[OP1] |
| 3 | SER 461[N] | 2.19 | G 199[O5'] |
| 4 | ARG 467[NH2] | 3.88 | C 64[O2'] |
| 5 | ILE 475[N] | 2.47 | C 33[N3] |
| 6 | GLN 500[N] | 2.74 | A 38[OP1] |
| 7 | ASN 533[ND2] | 2.27 | G 66[O2'] |
| 8 | GLN 549[N] | 2.13 | G 199[O2'] |
| 9 | GLU 550[N] | 3.81 | G 199[O2'] |
| 10 | ARG 551[NH1] | 2.79 | U 65[OP2] |
| 11 | ARG 551[NH1] | 3.60 | C 198[OP2] |
| 12 | ARG 551[NH1] | 3.24 | C 64[O3'] |
| 13 | B:PRO 472[O] | 2.76 | C 33[N4] |
| 14 | GLU 480[OE1] | 3.01 | A 35[N6] |
| 15 | GLU 507[OE1] | 3.88 | G 39[N2] |
| 16 | GLN 459[O] | 3.82 | A 155[N6] |
| 17 | GLU 545[OE2] | 3.53 | A 205[N1] |
| 18 | GLU 545[OE1] | 3.74 | G 206[N2] |
| 19 | GLN 453[OE1] | 2.82 | G 247[N2] |
| **VT2** |  |  |  |
| 1 | HIS 526[ND1] | 2.68 | A 14[OP2] |
| 2 | ASN 533[ND2] | 2.63 | G 12[O3'] |
| 3 | ARG 540[NH1] | 2.59 | U 224[OP1] |
| 4 | ARG 540[NH1] | 3.86 | A 223[O3'] |
| 5 | ARG 540[NH2] | 3.23 | U 224[O3'] |
| 6 | LYS 541[NZ] | 3.68 | G 260[O4'] |
| 7 | SER 558[OG] | 3.28 | C 234[O2] |
| 8 | SER 558[OG] | 2.83 | C 249[O2] |
| 9 | SER 558[OG] | 3.65 | G 233[N2] |
| 10 | ASP 556[OD2] | 2.10 | G 248[N2] |
| 11 | SER 558[OG] | 3.62 | G 248[N2] |
| **VT3** |  |  |  |
| 1 | GLY 0[N] | 3.02 | G 101[ OP1] |
| 2 | SER 445[OG] | 3.83 | C 151[ OP1] |
| 3 | SER 445[OG] | 3.56 | C 150[ O3'] |
| 4 | SER 445[OG] | 3.50 | C 150[ O2'] |
| 5 | ARG 448[NE] | 2.96 | C 151[OP2] |
| 6 | LYS 454[N] | 3.70 | A 215[O2'] |
| 7 | LYS 454[NZ] | 2.10 | C 218[OP2] |
| 8 | LEU 455[N] | 3.74 | A 215[O3'] |
| 9 | LEU 455[N] | 3.39 | A 216[O5'] |
| 10 | ARG 457[N] | 3.25 | U 137[O4] |
| 11 | ARG 457[N] | 3.60 | U 85[O4] |
| 12 | ARG 457[NH1] | 2.53 | C 217[O5'] |
| 13 | ARG 457[NH2] | 3.05 | U 137[O2] |
| 14 | SER 461[N] | 3.10 | A 80[OP1] |
| 15 | THR 462[N] | 3.38 | U 225[O4] |
| 16 | TRP 536[NE1] | 2.04 | A 70[OP1] |
| 17 | LYS 541[NZ] | 3.86 | A 69[O2'] |
| 18 | LYS 541[NZ] | 3.71 | A 70[OP1] |
| 19 | ASP 548[N] | 2.78 | A 79[OP1] |
| 20 | GLU 545[OE2] | 3.90 | C 75[N4] |
| 21 | LYS 454[O] | 3.33 | C 234[N4] |
| 22 | ARG 457[O] | 2.36 | A 235[N6] |
| 23 | GLN 530[OE1] | 3.74 | G 242[N2] |
| **VT4** |  |  |  |
| 1 | ARG 448[N] | 3.32 | G 56[O2'] |
| 2 | GLN 453[NE2] | 3.82 | C 98[O4'] |
| 3 | GLN 453[NE2] | 3.45 | A 97[O2'] |
| 4 | ARG 467[NH2] | 3.85 | C 230[N3] |
| 5 | LYS 473[N] | 3.90 | A 243[OP2] |
| 6 | ASP 474[N] | 3.25 | C 244[OP1] |
| 7 | LYS 489[NZ] | 3.24 | U 76[OP2] |
| 8 | GLN 530[NE2] | 2.54 | G 63[O6] |
| 9 | ASN 533[ND2] | 2.79 | A 226[OP2] |
| 10 | ARG 535[NH1] | 2.47 | C 64[O2] |
| 11 | ARG 535[NH1] | 2.46 | G 66[ O4'] |
| 12 | ARG 535[NH2] | 2.66 | G 63[O6] |
| 13 | TRP 536[N] | 2.76 | G 66[ OP2] |
| 14 | TRP 536[NE1] | 2.18 | U 65[O2'] |
| 15 | VAL 546[N] | 3.15 | G 227[OP1] |
| 16 | ARG 551[NH1] | 2.31 | A 229[OP1] |
| 17 | TRP 536[O] | 3.84 | A 74[N6] |
| 18 | ALA 538[O] | 3.53 | A 4[N1] |
| 19 | ALA 559[OXT] | 2.67 | C 230[N4] |

Additionally, some of these contacts, mediated by the backbone of the RNA, include the OP4 and OP5 atoms of uracil and other backbones of a few cytosine nucleotides. Notably, several hidden atoms within the RNA backbone also formed electrostatic contacts that contributed to the stability of the RNA-protein complex. The number of contacts mediated by the nucleotide residues is also represented in **Suppl Figure 2.**

Regarding the interactions between wild-type 5’ UTR and PUF60, our investigation aimed to comprehend how the uridine-rich motifs are lost in different variants of the 5’ UTR. The VT2 showed a distinct conformation in which PUF60 is housed at the end of the RNA architecture (**Figure 3A**). This structural conformation implies that it allows for fewer hydrogen bonds with PUF 60 (**Table 3**). Most of these interactions were centered on the purines, with only two hydrogen bonds formed by uracil and cytosine (**Figure 4B**). Guanine dominated the interactions, signifying a loss of the uridine motifs. Conversely, the VT3 variant had 23 hydrogen bonds with PUF 60 (**Table 3**), with adenine dominating with nine hydrogen bonds, also indicating the loss of the pyrimidine motifs despite the number of contacts.

**Figure 4:**
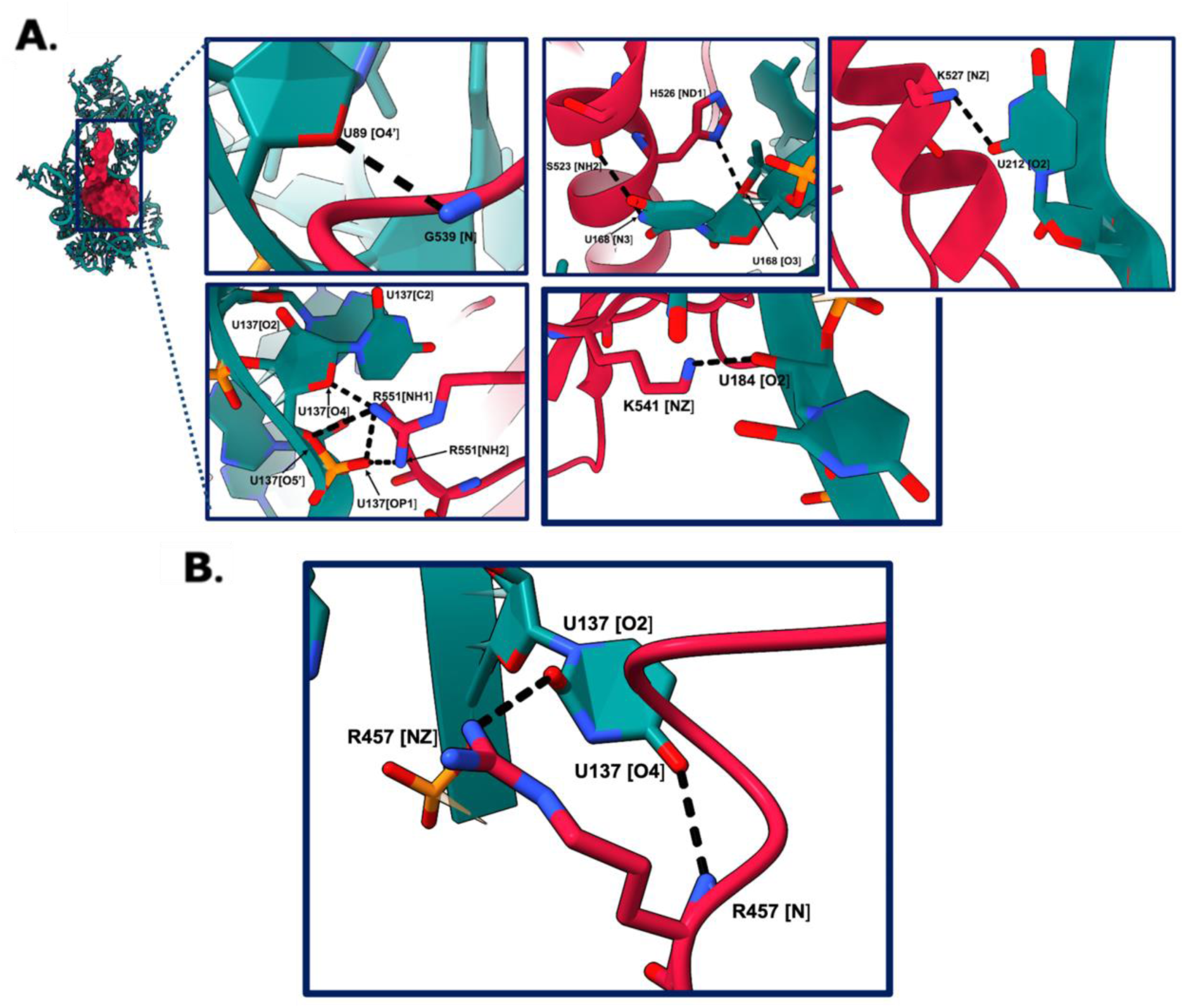
The wild type of 5’ UTR and PUF60 protein complex. PUF 60 is represented in crimson, and the 5’ UTR is in teal color. Atomic representatives of the residues that participate in hydrogen interactions are shown in the ribbon. **A**. R551 and U137 mediate the formation of four hydrogen bonds between the side chain of G539 and U89. Two hydrogen bonds are also mediated by U168’s side chains. The NZ atom of K541 and the O2 atom of U184 create a single hydrogen bond, whereas K527 and U212 form a second hydrogen link. **B**. The U137 conserved motif in VT3. Arginine interacts with the U137 using both the side chain and backbone atomic subdomains to make contact with the O atoms of U137.

We observed a conserved interaction for VT3. Despite the induced global mutation, we found that a conserved motif, U137, also present in the wild-type 5’ UTR, remained in this variant. Notably, we observed that the O2 and O4 atoms of U137 formed interactions with the arginine (R457 NH2) residue from PUF 60 (**Figure 4**). This finding suggests that the mutation of U137 could significantly alter the viral RNA, thereby affecting PUF 60’s ability to recognize and process it. Moreover, the conserved recognition of U137 by arginine in both the wild type and VT3 raises the possibility that a mutation in the arginine residue might affect PUF 60’s capacity to recognize crucial motifs on the viral RNA. In addition, based on the observed results, we suggest that the stable contacts between the SARS-CoV-2 viral RNA and PUF60 require both the side chain and backbone for a stable conformation, and a mutation across either of these could disrupt these interactions.

### 3’ UTR-derived variants and loss of binding motifs

We also studied the 3’ UTR and its induced derivative variants to understand how the loss of functions contributes to the protein’s inability to regulate the RNA’s gene expression. Using the same structural approach as we did for the 5’ UTR, we could juxtapose the structural differences induced by introducing the global mutation in the RNA. We used the KHDRBS3 protein, also known as T-STAR, as a reference 3’ UTR RBP to study the loss of binding motifs. The T-STAR is a protein that regulates the alternative splicing of mRNAs (Ehrmann et al., 2013). Unlike other splicing factors, they contain just a single RNA-binding domain (Feracci et al., 2016), which makes them suitable as a reference for the study. Additionally, like other RBPs, KHDRBS3 plays a crucial role in regulating viral replication. Their interaction with viral RNA affects viral stability, translation, and replication. We retrieved the T-STAR protein from the PDB repository (PDB: 5EL3) (**Figure 5A**).

**Figure 5:**
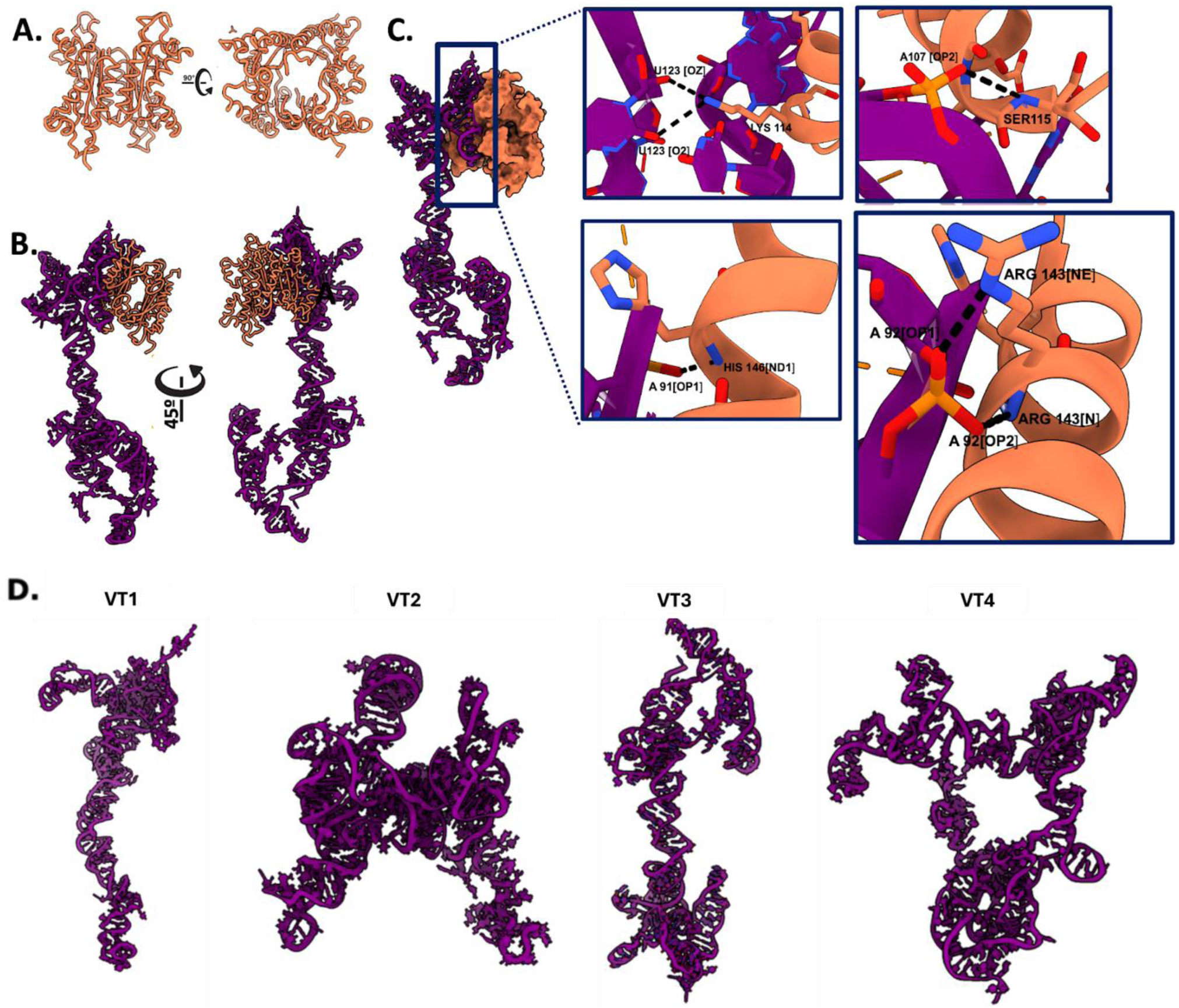
Interaction between the 3’UTR wildtype and the T-STAR protein. **A**. Structure of KHDRBS3 protein, also known as T-STAR (PDB: 5EL3). **B**. Binding of T-STAR protein to the 3’UTR wild type **C**. Zoomed overall interactions of the 3’UTR wildtype and the T-STAR. **D**. Different structural conformations of the 3’ UTR-induced variants.

The wild-type 3’ UTR had an RNA duplex geometry and an almost 90-degree kink at the top end, giving way to a right-angled 5-way loop at the top serving as a scaffold where T-STAR binds (**Figure 5B**). T-STAR binds to the edge of this 5-way loop, making contact with the protein’s helical core containing the relevant motifs. We also observed that the signatory motifs are in the helical core of the KHDRBS3 protein (**Figure 5C**). Some of these motifs pierced through the sugar-phosphate backbone of the RNA. They made contact with the nitrogenous bases, as shown in HIS 146, forming a hydrogen bond with the phosphate atom of adenine at position 91 using its backbone. The KHDRBS3 mediated some interactions through the amino backbones. Another significant interaction is through the phosphorus atom surrounded by oxygen atoms. A notable example of this backbone interaction is ARG 143 (**Figure 5C**), which makes a hydrogen bond contact with its side chain and almost hidden carbonyl backbone. The sidechain of ARG 143 also makes close contact with the phosphate backbone of adenine at position 92 in the wild-type but is not seen in VT3. The structural changes to the RNA elements at the 3’ UTR in all the variants (Figure 5D) may contribute to the loss of interactions between ARG 143 and adenine at position 92. As shown in **Figure 5C**, the presence of the phosphate backbone OP4 and OP5 contributes to the overall negative charge of the RNA molecule. We have summarized the rest of the interactions in **Table 4**, which also contains the interactions between the induced variants and the T-STAR.

**Table 4:** Interactions between PUF60 and 3’ UTR of the wild-type and the variant 3.

|  | T-STAR | Dist. [Å] | RNA |
| --- | --- | --- | --- |
| **Wild type 3'UTR** |  |  |  |
| 1 | B:ARG 113[ NH1] | 2.03 | A:G 149[ OP2] |
| 2 | B:LYS 114[ NZ] | 3.62 | A:A 147[ OP1] |
| 3 | B:LYS 114[ NZ] | 3.23 | A:G 145 |
| 4 | C:LYS 114[ NZ] | 3.50 | A:U 123[O2'] |
| 5 | C:LYS 114[ NZ] | 3.49 | A:U 123[O2] |
| 6 | C:SER 115[ N] | 3.84 | A:A 107[OP2] |
| 7 | C:GLU 111[ OE2] | 2.23 | A:C 113[N4] |
| 8 | D:ASN 51[ ND2] | 3.15 | A:A 134[ OP1] |
| 9 | D:ARG 143[ N] | 2.32 | A:A 92[OP2] |
| 10 | D:HIS 146[ ND1] | 2.60 | A:A 91[O4'] |
| 11 | D:ASN 160[ ND2] | 3.18 | A:U 43[O3'] |
| 12 | E:ASN 53[ ND2] | 2.63 | A:A 150[OP2] |
| 13 | E:LYS 153[ NZ] | 3.17 | A:G 137[OP2] |
| 14 | E:PHE 134[ O] | 3.57 | A:A 141[N6] |
| **VT3** |  |  |  |
| 1 | E:ASN 71[ ND2] | 2.77 | A:A 318[OP1] |
| 2 | E:ASN 71[ ND2] | 3.82 | A:A 328[O4'] |
| 3 | E:ARG 80[ N] | 3.83 | A:A 327[ OP2] |
| 4 | E:ARG 80[ NH1] | 2.06 | A:A 20[ OP1] |
| 5 | E:ARG 80[ NH2] | 2.94 | A:U 90[ OP1] |
| 6 | E:ASN 82[ ND2] | 3.46 | A:A 311[O2'] |
| 7 | E:THR 91[ N] | 3.16 | A:G 27[ O2'] |
| 8 | E:LYS 94[ NZ] | 3.39 | A:A 78[ OP1] |
| 9 | E:LYS 153[ NZ] | 3.69 | A:C 26[O3'] |
| 10 | E:LYS 153[ NZ] | 2.33 | A:G 27[OP1] |
| 11 | E:LYS 69[ O] | 3.85 | A:A 332[ N1] |
| 12 | E:ARG 80[ O] | 2.26 | A:A 313[ N6] |
| 13 | E:GLU 89[ O] | 2.32 | A:A 30[ N6] |
| 14 | E:GLU 89[ O] | 2.70 | A:C 29[ N4] |
| 15 | E:THR 91[ O] | 2.01 | A:A 28[ N6] |
| 16 | B:ARG 104[ NE] | 3.88 | A:U 94[ O2] |
| 17 | B:LYS 108[ NZ] | 2.43 | A:C 81[ OP1] |
| 18 | B:LEU 112[ N] | 3.45 | A:G 79[ O6] |
| 19 | B:GLU 111[ O] | 3.80 | A:C 80[ N4] |
| 20 | B:ARG 86[ O] | 2.98 | A:A 253[ N6] |

### Folding patterns of the 5’ and 3’ UTRs of the wild type and variants

To uncover what could be responsible for the loss of essential, rich uracil motifs, we studied the folding patterns that could alter how these motifs could be presented to PUF60 to maintain the stability of the RNA. We noticed a folding pattern with meaningful biological relevance for the wild type. The presence of about eight stem-loops (hairpins) with a double-stranded stem with stacked base pairs and a single-stranded loop at the structure’s tip. This site is known to be the hotspot for protein binding, as PUF60 majorly binds in these areas of the RNA. We also noticed the 3-, 4-, and 5-way junctions, where multiple helices converging on the single loop. This could also be responsible for the kinks and the 90° angle we stated earlier. This region was also crucial for the binding of the PUF60, as shown in **Figure 6A**. We noticed that the stem loops have the essential motifs (uracil and a few cytosines) necessary for PUF60 binding.

**Figure 6:**
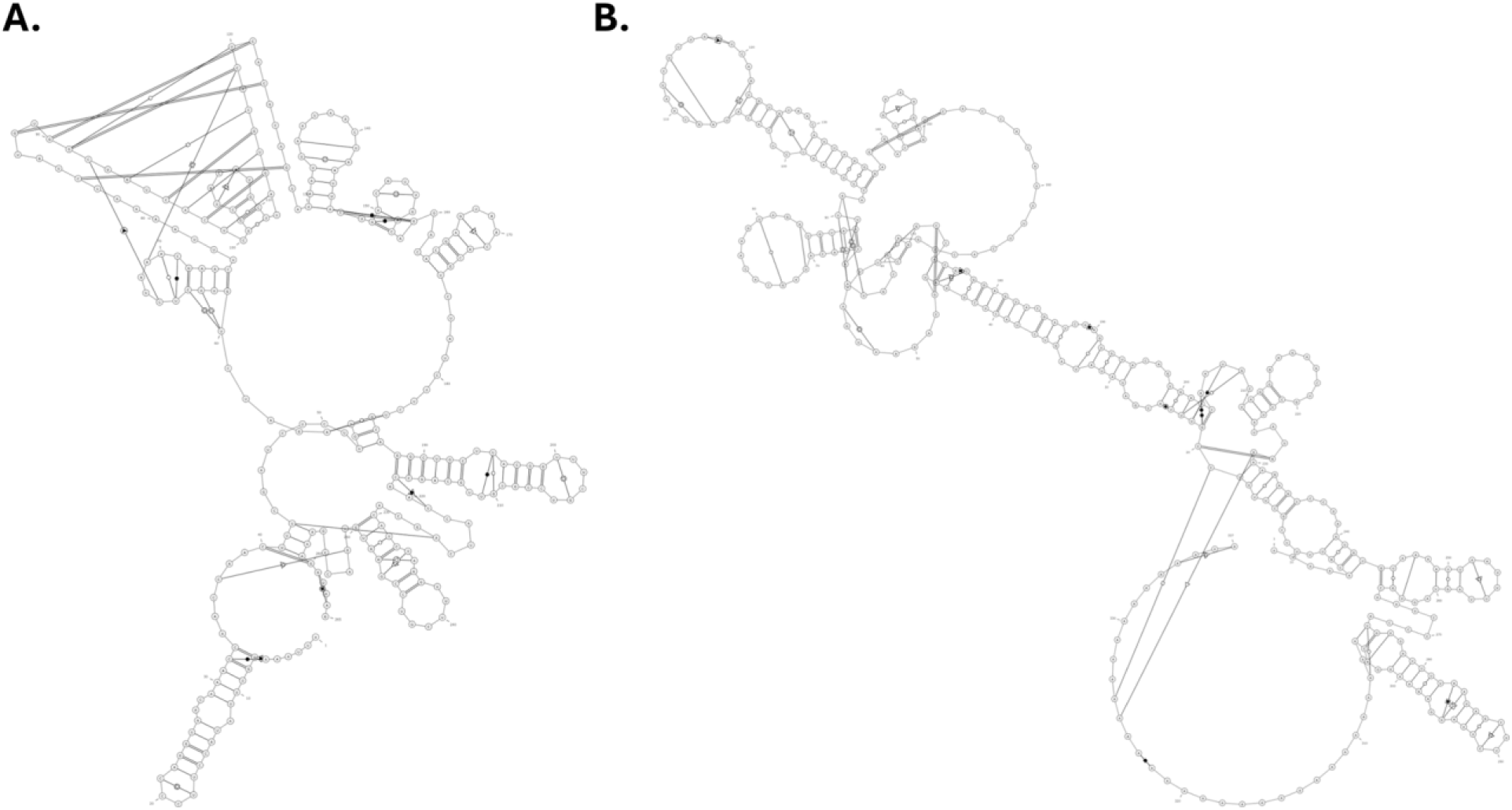
Folding patterns of **A.** 5’ UTR wild type and **B.** 3’ UTR wild type.

Additionally, we observed some extended regions without base pairing, which may be accessible for interactions with the translation machinery or other RNP complexes. Another observation was a large internal loop, which could also play a critical role. We noticed that PUF60 also binds to this region of the wild type, accessing the uracil embedded within it. For the VT3, we also saw similar patterns of stem loops, bulges, and extended internal loops that contain the necessary motifs for PUF60 recognition. This phenomenon was observed for the VT3, which contains up to five stem loops with the motifs required for PUF60 recognition, three internal central loops that include uracil, and a few mismatches. A striking observation was the 5JC variant. This variant is characterized by a large, massive terminal loop, which has a circular appearance. There is also a long-range duplex, which might indicate an extensive pairing between the 5′ and 3′ ends. We also observed a few internal junctions, a smaller multi-loop region, and several stem loops. This contributes to the loss of proper folding and the inability of PUF60 to access essential motifs. The structure also lacks structural flexibility, as it is locked in a compact formation, providing less accessibility to crucial motifs (**Suppl Figure 3**).

In the 3’ UTR, stem loops (hairpins) containing the rich adenine motifs were present for the wild type. The same structural similarities were observed for some of the variants. However, for VT3, we observed a large circular terminal loop, which contributed to the rigidity of the structure and reduced the presentation of stem loops and adenine-rich nucleotides required for KHDRBS3 recognition and stabilization of the RNA (**Figure 6B and Supplementary Figure 4**).

## Discussion

Compelling evidence suggests that RBPs play critical roles in SARS-CoV-2 RNA turnover when the virus infects human cells. Yet, there is a scarcity of reports on how silent mutations in the viral genome affect its RNA interacting with host RBPs. To address this, we first map the human-virus RBP-RNA interaction for the reference sequence (wildtype) of SARS-CoV-2 using RBPmap (Paz et al., 2014). Then, we performed in silico mutagenesis on the wildtype (WT) to create 4 synthetic variants without changing the amino sequences of the coding regions. We used different codon optimization tools (see methods) to allow changes in the entire genome without altering the amino acid sequences. The aligned amino acid sequences of the variants and the WT show 100% similarity but vary in their nucleotide sequences, which thus confirms that the variants are silent mutants of the wild type.

RBPmap catalogues 223 defined motifs for human RBPs. While some of the RBPs contain one or two motifs, we observed loss and gain of binding motifs for some RBPs in the variants compared to the wild type. Previously, Sun et al. (2021) used the PRISMNet deep learning model to map binding motifs for 42 RBPs at the 5’ and 3’ UTRs of SARS-CoV-2, and while the predicted motifs are large, Horlacher et al. (2023) recently used two deep learning methods (pysster and DeepRiPe) to map over 100 human RBPs interacting with SARS-CoV-2 RNA at nucleotide resolution. However, our study addresses the impact of silent mutations across the entire viral genome and its synthetic variants and uses structural analysis to define the contacts between the RNA nucleotides and the amino acid residues in selected RBPs.

Our investigation predicted a loss or gain of binding motifs for new RBPs that do not interact with SARS-CoV-2 and experimentally verified RBPs. Furthermore, our predictions offer significant insights into the interactions between the host and the virus, particularly regarding silent mutations within the viral genome. Our study can facilitate understanding that RBPs can serve as pro- and antivirals when silent mutations occur within the coding and non-coding regions of the viral genome.

Likewise, these predictions may facilitate new therapeutic targets, for example, by developing drugs that could interfere with binding sites of pro-viral RBPs to disrupt the binding to their motifs. Additionally, our synthetic variants generated via *in silico* mutagenesis may offer insight into surveillance of new SARS-CoV-2 variants or other coronaviruses that could be of concern in the future.

The silent mutation impacts some of the RBPs that have been experimentally validated to interact with the SARS-CoV-2 genome. For example, out of the top 10 RBPs with loss of binding position at the 5’ UTR, PUM1 (Phan et al. 2025) lost its conserved binding positions in all the variants, which suggests a loss of function of protein-RNA interaction. We also predicted the binding of PUF60 and KHDRBS3 to the viral RNA. Although PUF60 regulates hepatitis B virus pregenomic RNA expression by interacting with transcription factor TCF7L2 and is involved in the viral RNA degradation and suppressing the RNA splicing (Sun et al., 2017), and the restriction of KHDRBS3 has been reported to suppress lytic reactivation of oncogenic herpesvirus by host microRNA-31- 5p (Lee et al., 2025), PUF60 and KHDRBS3 have not been previously reported to interact with SARS-CoV-2. However, our findings show that the PUF60 binding motif with poly(U) is conserved in the wild-type viral genome. Our prediction indicates the loss of this motif in the variants, which may impact the function of the RBP as a viral RNA regulator. Another surprising finding from our prediction is that the majority of RBPs with AU-rich binding motifs are reduced in the variants, while RBPs with mostly GC-rich motifs gain binding positions. Because SARS-CoV-2, like most viruses, has a mechanism of evading host protein interactions to facilitate its RNA processing, we hypothesized that a single change from U/A to G/C could impact RBP-viral interactions. We show evidence of this in the structural analysis of PUF60 interacting with the wildtype and the variant, where loss of U137 in some of the variants disrupts contact with R457 in the RBP (**Figure 4**). Our findings might suggest a critical role of PUF60 in regulating the viral RNA, in which experimental validation is necessary to assess the key important nucleotides and residues in the viral RNA and in the protein, respectively.

Viral RNA structure has been implicated in several processes such as replication, immune evasion, packaging, and synthesis of viral protein (Boerneke et al., 2023). The functional RNA structures in the virus are also important therapeutic targets. Our study shows that silent mutations change the structure of the viral RNA at both the 5’ and the 3’ UTR in all the variants. The newly shaped structure in the variants may confer new biophysical interactions with relevant RBPs or disrupt already known interactions, for example, at the 3’ end, where poly(A) plays a major role in RNA stability (Bar et al., 2010; Tang et al., 2019). They could also form pseudoknots, which may preferentially resist degradation by host exoribonucleases (Gezelle et al., 2024).

Although our study is computationally assessed, our findings contribute to the understanding of the impact of silent mutations on the host RBPs-SARS-CoV-2 viral RNA interactome. However, our study is limited by experimental validation. Extensive biochemical and structural studies are necessary to support these findings. We also believe that there are some overlaps in the mapping of binding motifs for more than one RBP in each location in the viral genome using the RBPmap, which are not captured in our report. Thus, mapping motif-specific RBP-RNA without overlap will improve on our current approach.

## Methods

### Sequence Retrieval

The SARS-CoV-2 reference sequence was retrieved from the nucleotide database of the National Center for Biotechnology Information (NCBI) (https://www.ncbi.nlm.nih.gov/nuccore/NC_045512). The reference sequence was used in this study, as it provides the basis for all mutations in the SARS-CoV-2 viral genome, leading to other variants.

### *In silico* Mutagenesis

Synonymous codon substitutions were carried out on the SARS-CoV-2 whole genome using codon optimization programs from JCat (Grote et al., 2005), VectorBuilder, Integrated DNA Technologies (IDT), and GeneArt (ThermoFisher) to introduce silent mutations without altering the amino acid sequence. Human/Homo sapiens was selected as the expression host in all the optimization and other default settings in each programme were unchanged. The variants generated, JCat (VT1), Vector builder (VT2), IDT (VT3), and GeneArt (VT4), were further compared with the wild-type SARS-CoV-2 nucleotide sequence (WT) for the RBP probing.

### Multiple Sequence Alignment

The four variant sequences generated were aligned with the reference sequence (wild-type) using the Multiple Sequence Alignment (MSA) Tool Clustal Omega (EMBL-EBI). ClustalW with character counts was selected as the output format, and other default parameters were left unchanged. The MSA result was used to construct the phylogenetic tree. The tree was constructed using the neighbour-joining method with MEGA 11 software (Tamura et al., 2021) by applying its MUSCLE program. Finally, the sequences and the phylogenetic tree for the 5’ and 3’ untranslated regions (UTRs) of the SARS-CoV-2 genome were aligned and generated, respectively.

### RNA Binding Proteins – RNA Interaction

To predict interactions of the RBPs and the SARS-CoV-2 WT and VT sequences, RBPmap version 1.2 was used to globally map the binding motif positions of RBPs on the viral sequences. RBPmap is an online algorithm containing 131 human RBPs with 223 defined human/mouse RNA binding motifs. The RBPmap program allows input of nucleotide sequences of interest, either as DNA or RNA. If DNA sequences are inputted, the algorithm will automatically convert the DNA into an RNA sequence as an output result during binding interaction. Users have the option to either insert the name of the RBP of interest or choose the RBPs and motifs from the full list provided on the web server. In this study, all 223 motifs for the human/mouse RBPs in the catalogue were selected. The RBPmap uses a weighted-rank approach to rank the motifs (Paz *et al.,* 2021) and statistically generate a z-score and p-value for each predicted binding interaction.

### Secondary RNA Structure Prediction

The secondary structures of the 5’ and 3’ UTRs of the RNA sequence were predicted using the RNAFold program (Lorenz *et al.,* 2016). The minimum free energy (MFE) and partition function were selected for the fold algorithms and basic options. The “avoid isolated base pairs,” which predicts the optimal secondary structure and removes lonely pairs, respectively, was also selected. Meanwhile, the non-canonical GU pairing was deselected to avoid weak pairs in the RNA structure. The dot-plot annotation generated was used to predict the 3D RNA structures using 3dRNA/DNA with default parameters (Zhang *et al.,* 2024).

### Protein-RNA structural analysis

Our studies used the two RBPs, which were PUF60 (PDB ID: 3US5) and KHDRBS3 (PDB: 5EL3), also known as T-STAR, as our representative splicing factor in complex with the 5’ UTR and 3’ UTR and their induced derived variants. We conducted molecular docking using a simulation approach to explore the interaction between the RBPs and these RNA variants to obtain exact docked protein-RNA complex structures. For this purpose, we utilized NPDOCK (Tuszynska et al., 2015), a program that integrates the Global Range Molecular Matching (GRAMM) methodology to dock molecules based on their atomic coordinates. The docking process was further refined through statistical estimation of scores, followed by clustering of structures exhibiting the best scores based on their hierarchy. From this cluster, the program automatically selected the optimal representatives of the RNA-protein complexes (decoys), which then underwent iterative rounds of refinement. During the simulation, 20,000 decoys are generated for the RBPs and the RNA structures using GRAMM, subsequently focusing on the 100 decoys with the highest scores for clustering. We chose a Root Mean Square Deviation (RMSD) of 7 Å for the clustering process. The docking process involved 1000 iterative steps with an initial starting temperature of 15,000 K and cooling down to 295 K. Eventually, we selected the decoy with the best statistical score and a clash score of 0 as our final structure. We visualize the obtained protein-RNA complex structure using Chimera X (Meng et al., 2023).

### RNA Intramolecular Interactions

We studied the intramolecular interactions of the wild-type 3’ UTR and 5’ UTR compared to their various induced variants. These interactions include the internal hydrogen bond, hydrophobic interactions, base stacking, hydrophobic interactions, electrostatic interactions, and other relevant internal interactions that contribute to the structural characterizations of the wild-type and induced variant RNAs. We used the RNApdbee v2.0 web server to generate the RNA architecture of the wild types and variants (Antczak *et al*., 2018; Zok *et al*., 2018).

