## Supplementary Figures for "Computational Analysis of Silent Mutation Effects on SARS-CoV-2 RNA–Host RNA-Binding Protein Interactome"

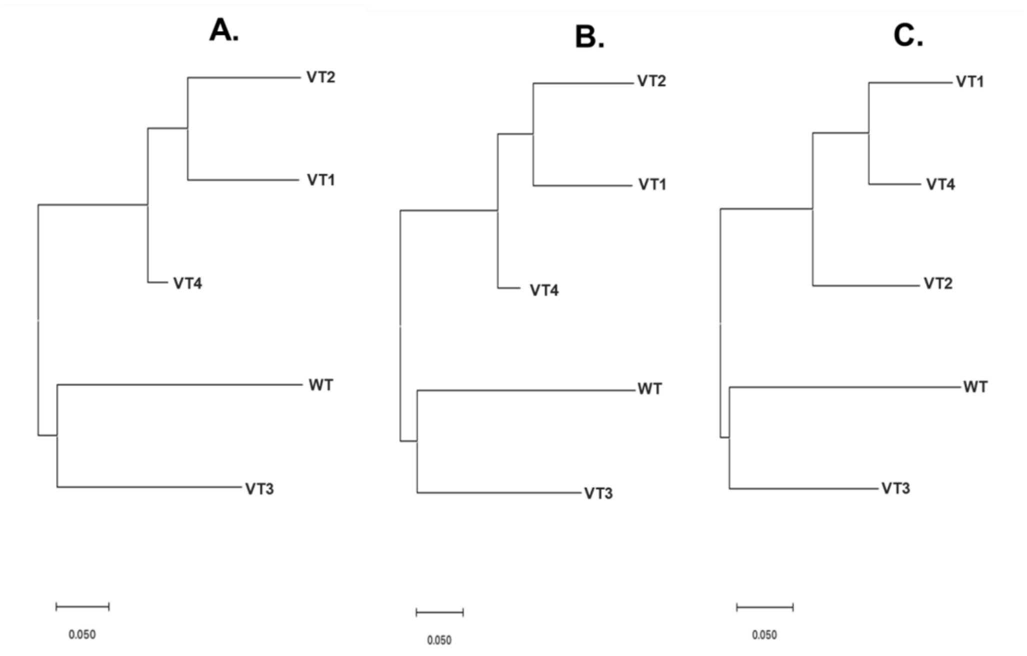


**Supplementary Figure 1:** Phylogenetic relationship among the wild type and the variants. **A.** whole genome. **B.** 5 UTR. **C.** 3 UTR

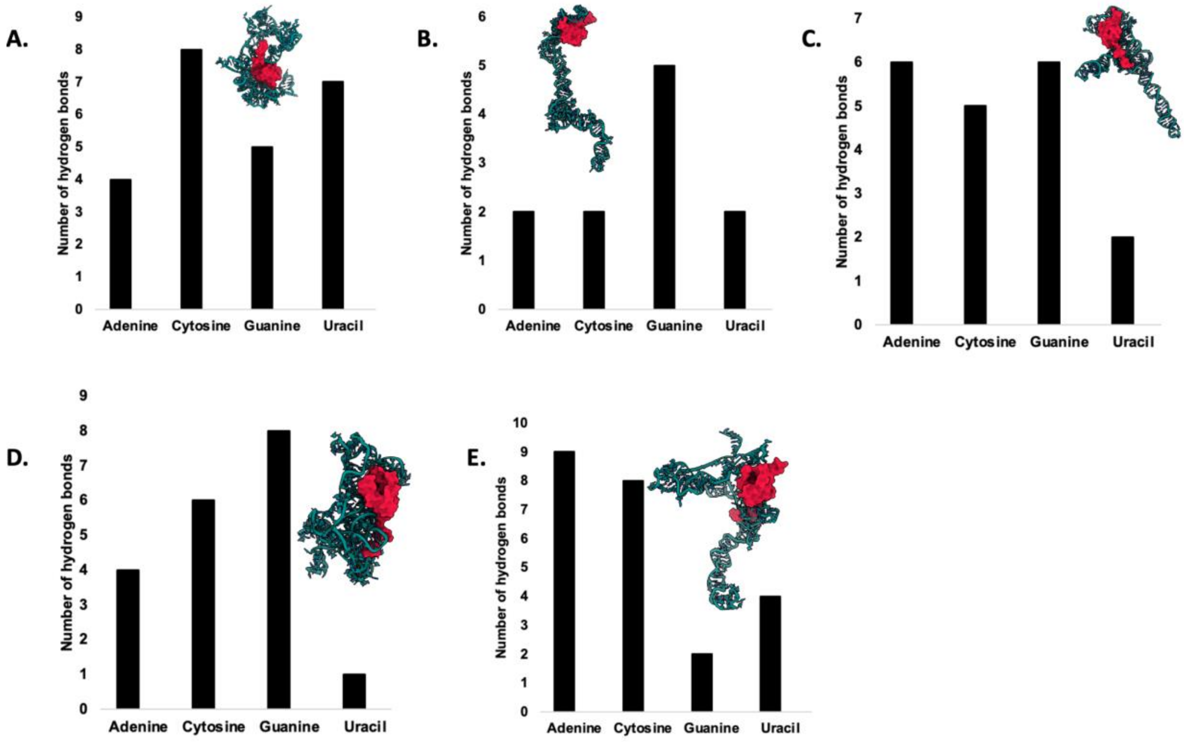


**Supplementary Figure 2**: Graphical representation of the number of contacts mediated by the Nucleotide residues. **A**. Wild type. **B**. VT1. **C**. VT2. **D**. VT3. **E**. VT4

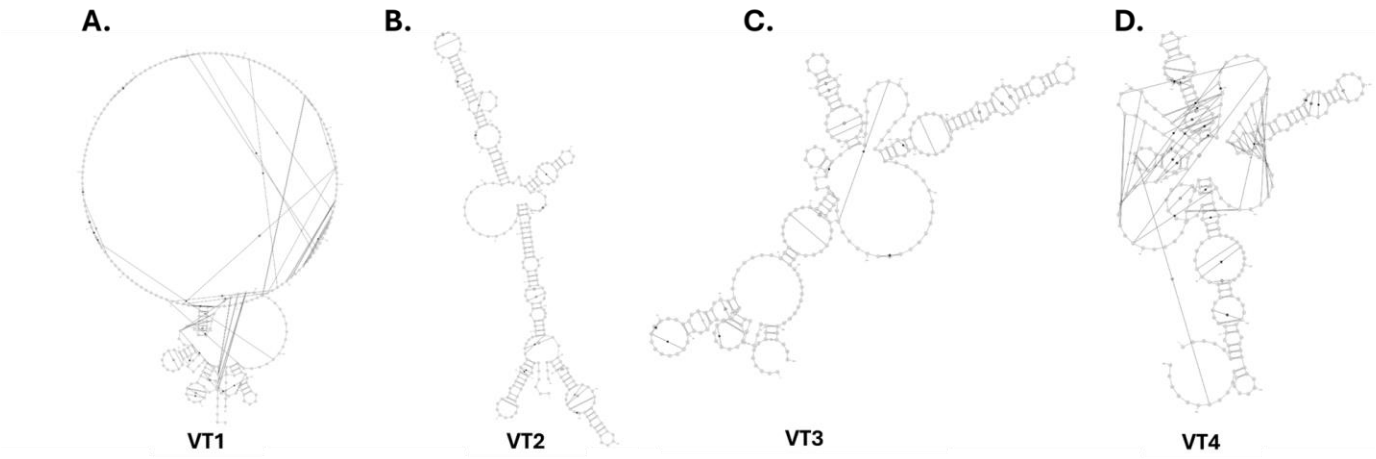


**Suppl Figure 3:** The folding pattern of 5' UTR variants.

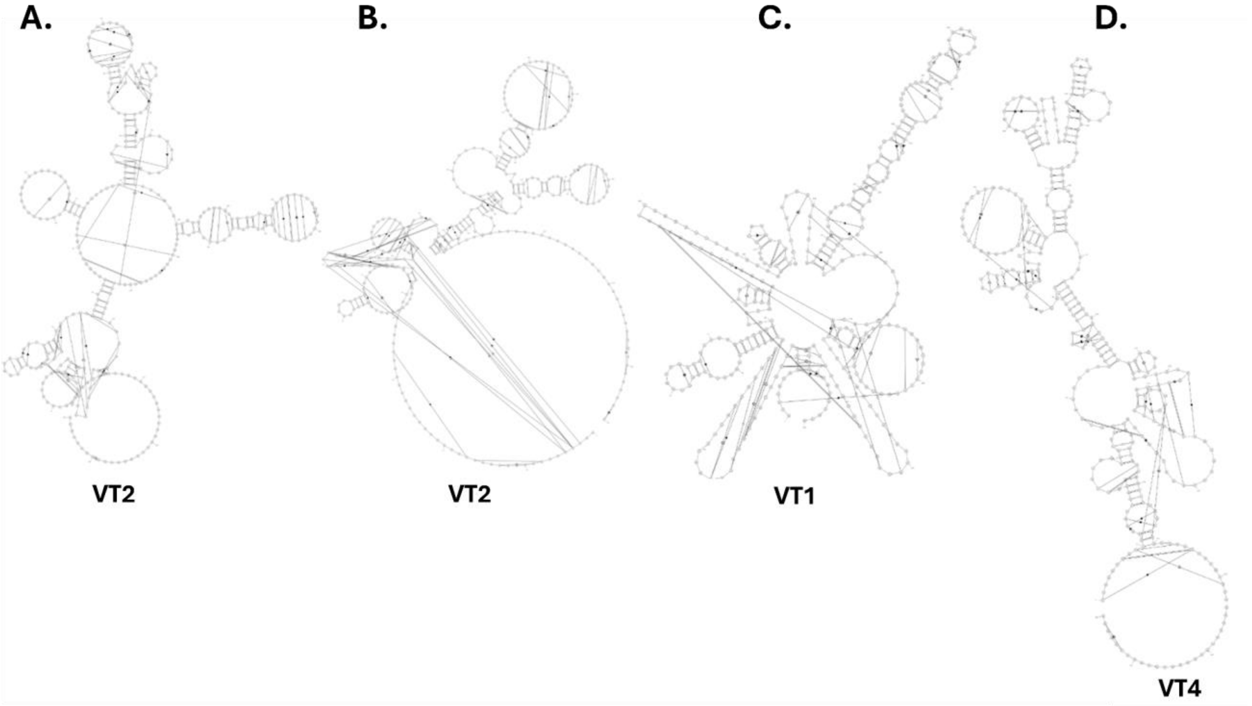


**Suppl Figure 4:** The folding pattern of 5' UTR variants.
