## Supplementary File for "Computational Analysis of Silent Mutation Effects on SARS-CoV-2 RNA–Host RNA-Binding Protein Interactome"

**>WT_NC_045512.2 Severe acute respiratory syndrome coronavirus 2 isolate Wuhan- Hu-1, complete genome**

ATTAAAGGTTTATACCTTCCCAGGTAACAAACCAACCAACTTTCGATCTCTTGTAGATCTGTTCTCTAAACGAACTT
TAAAATCTGTGTGGCTGTCACTCGGCTGCATGCTTAGTGCACTCACGCAGTATAATTAATAACTAATTACTGTCGTT
GACAGGACACGAGTAACTCGTCTATCTTCTGCAGGCTGCTTACGGTTTCGTCCGTGTTGCAGCCGATCATCAGCACA
TCTAGGTTTCGTCCGGGTGTGACCGAAAGGTAAGATGGAGAGCCTTGTCCCTGGTTTCAACGAGAAAACACACGTCC
AACTCAGTTTGCCTGTTTTACAGGTTCGCGACGTGCTCGTACGTGGCTTTGGAGACTCCGTGGAGGAGGTCTTATCA
GAGGCACGTCAACATCTTAAAGATGGCACTTGTGGCTTAGTAGAAGTTGAAAAAGGCGTTTTGCCTCAACTTGAACA
GCCCTATGTGTTCATCAAACGTTCGGATGCTCGAACTGCACCTCATGGTCATGTTATGGTTGAGCTGGTAGCAGAAC
TCGAAGGCATTCAGTACGGTCGTAGTGGTGAGACACTTGGTGTCCTTGTCCCTCATGTGGGCGAAATACCAGTGGCT
TACCGCAAGGTTCTTCTTCGTAAGAACGGTAATAAAGGAGCTGGTGGCCATAGTTACGGCGCCGATCTAAAGTCATT
TGACTTAGGCGACGAGCTTGGCACTGATCCTTATGAAGATTTTCAAGAAAACTGGAACACTAAACATAGCAGTGGTG
TTACCCGTGAACTCATGCGTGAGCTTAACGGAGGGGCATACACTCGCTATGTCGATAACAACTTCTGTGGCCCTGAT
GGCTACCCTCTTGAGTGCATTAAAGACCTTCTAGCACGTGCTGGTAAAGCTTCATGCACTTTGTCCGAACAACTGGA
CTTTATTGACACTAAGAGGGGTGTATACTGCTGCCGTGAACATGAGCATGAAATTGCTTGGTACACGGAACGTTCTG
AAAAGAGCTATGAATTGCAGACACCTTTTGAAATTAAATTGGCAAAGAAATTTGACACCTTCAATGGGGAATGTCCA
AATTTTGTATTTCCCTTAAATTCCATAATCAAGACTATTCAACCAAGGGTTGAAAAGAAAAAGCTTGATGGCTTTAT
GGGTAGAATTCGATCTGTCTATCCAGTTGCGTCACCAAATGAATGCAACCAAATGTGCCTTTCAACTCTCATGAAGT
GTGATCATTGTGGTGAAACTTCATGGCAGACGGGCGATTTTGTTAAAGCCACTTGCGAATTTTGTGGCACTGAGAAT
TTGACTAAAGAAGGTGCCACTACTTGTGGTTACTTACCCCAAAATGCTGTTGTTAAAATTTATTGTCCAGCATGTCA
CAATTCAGAAGTAGGACCTGAGCATAGTCTTGCCGAATACCATAATGAATCTGGCTTGAAAACCATTCTTCGTAAGG
GTGGTCGCACTATTGCCTTTGGAGGCTGTGTGTTCTCTTATGTTGGTTGCCATAACAAGTGTGCCTATTGGGTTCCA
CGTGCTAGCGCTAACATAGGTTGTAACCATACAGGTGTTGTTGGAGAAGGTTCCGAAGGTCTTAATGACAACCTTCT
TGAAATACTCCAAAAAGAGAAAGTCAACATCAATATTGTTGGTGACTTTAAACTTAATGAAGAGATCGCCATTATTT
TGGCATCTTTTTCTGCTTCCACAAGTGCTTTTGTGGAAACTGTGAAAGGTTTGGATTATAAAGCATTCAAACAAATT
GTTGAATCCTGTGGTAATTTTAAAGTTACAAAAGGAAAAGCTAAAAAAGGTGCCTGGAATATTGGTGAACAGAAATC
AATACTGAGTCCTCTTTATGCATTTGCATCAGAGGCTGCTCGTGTTGTACGATCAATTTTCTCCCGCACTCTTGAAA
CTGCTCAAAATTCTGTGCGTGTTTTACAGAAGGCCGCTATAACAATACTAGATGGAATTTCACAGTATTCACTGAGA
CTCATTGATGCTATGATGTTCACATCTGATTTGGCTACTAACAATCTAGTTGTAATGGCCTACATTACAGGTGGTGT
TGTTCAGTTGACTTCGCAGTGGCTAACTAACATCTTTGGCACTGTTTATGAAAAACTCAAACCCGTCCTTGATTGGC
TTGAAGAGAAGTTTAAGGAAGGTGTAGAGTTTCTTAGAGACGGTTGGGAAATTGTTAAATTTATCTCAACCTGTGCT
TGTGAAATTGTCGGTGGACAAATTGTCACCTGTGCAAAGGAAATTAAGGAGAGTGTTCAGACATTCTTTAAGCTTGT
AAATAAATTTTTGGCTTTGTGTGCTGACTCTATCATTATTGGTGGAGCTAAACTTAAAGCCTTGAATTTAGGTGAAA
CATTTGTCACGCACTCAAAGGGATTGTACAGAAAGTGTGTTAAATCCAGAGAAGAAACTGGCCTACTCATGCCTCTA
AAAGCCCCAAAAGAAATTATCTTCTTAGAGGGAGAAACACTTCCCACAGAAGTGTTAACAGAGGAAGTTGTCTTGAA
AACTGGTGATTTACAACCATTAGAACAACCTACTAGTGAAGCTGTTGAAGCTCCATTGGTTGGTACACCAGTTTGTA
TTAACGGGCTTATGTTGCTCGAAATCAAAGACACAGAAAAGTACTGTGCCCTTGCACCTAATATGATGGTAACAAAC
AATACCTTCACACTCAAAGGCGGTGCACCAACAAAGGTTACTTTTGGTGATGACACTGTGATAGAAGTGCAAGGTTA
CAAGAGTGTGAATATCACTTTTGAACTTGATGAAAGGATTGATAAAGTACTTAATGAGAAGTGCTCTGCCTATACAG
TTGAACTCGGTACAGAAGTAAATGAGTTCGCCTGTGTTGTGGCAGATGCTGTCATAAAAACTTTGCAACCAGTATCT
GAATTACTTACACCACTGGGCATTGATTTAGATGAGTGGAGTATGGCTACATACTACTTATTTGATGAGTCTGGTGA
GTTTAAATTGGCTTCACATATGTATTGTTCTTTCTACCCTCCAGATGAGGATGAAGAAGAAGGTGATTGTGAAGAAG
AAGAGTTTGAGCCATCAACTCAATATGAGTATGGTACTGAAGATGATTACCAAGGTAAACCTTTGGAATTTGGTGCC
ACTTCTGCTGCTCTTCAACCTGAAGAAGAGCAAGAAGAAGATTGGTTAGATGATGATAGTCAACAAACTGTTGGTCA
ACAAGACGGCAGTGAGGACAATCAGACAACTACTATTCAAACAATTGTTGAGGTTCAACCTCAATTAGAGATGGAAC
TTACACCAGTTGTTCAGACTATTGAAGTGAATAGTTTTAGTGGTTATTTAAAACTTACTGACAATGTATACATTAAA
AATGCAGACATTGTGGAAGAAGCTAAAAAGGTAAAACCAACAGTGGTTGTTAATGCAGCCAATGTTTACCTTAAACA
TGGAGGAGGTGTTGCAGGAGCCTTAAATAAGGCTACTAACAATGCCATGCAAGTTGAATCTGATGATTACATAGCTA
CTAATGGACCACTTAAAGTGGGTGGTAGTTGTGTTTTAAGCGGACACAATCTTGCTAAACACTGTCTTCATGTTGTC
GGCCCAAATGTTAACAAAGGTGAAGACATTCAACTTCTTAAGAGTGCTTATGAAAATTTTAATCAGCACGAAGTTCT
ACTTGCACCATTATTATCAGCTGGTATTTTTGGTGCTGACCCTATACATTCTTTAAGAGTTTGTGTAGATACTGTTC
GCACAAATGTCTACTTAGCTGTCTTTGATAAAAATCTCTATGACAAACTTGTTTCAAGCTTTTTGGAAATGAAGAGT
GAAAAGCAAGTTGAACAAAAGATCGCTGAGATTCCTAAAGAGGAAGTTAAGCCATTTATAACTGAAAGTAAACCTTC
AGTTGAACAGAGAAAACAAGATGATAAGAAAATCAAAGCTTGTGTTGAAGAAGTTACAACAACTCTGGAAGAAACTA
AGTTCCTCACAGAAAACTTGTTACTTTATATTGACATTAATGGCAATCTTCATCCAGATTCTGCCACTCTTGTTAGT
GACATTGACATCACTTTCTTAAAGAAAGATGCTCCATATATAGTGGGTGATGTTGTTCAAGAGGGTGTTTTAACTGCT

GTGGTTATACCTACTAAAAAGGCTGGTGGCACTACTGAAATGCTAGCGAAAGCTTTGAGAAAAGTGCCAACAGACA
ATTATATAACCACTTACCCGGGTCAGGGTTTAAATGGTTACACTGTAGAGGAGGCAAAGACAGTGCTTAAAAAGTGT
AAAAGTGCCTTTTACATTCTACCATCTATTATCTCTAATGAGAAGCAAGAAATTCTTGGAACTGTTTCTTGGAATTT
GCGAGAAATGCTTGCACATGCAGAAGAAACACGCAAATTAATGCCTGTCTGTGTGGAAACTAAAGCCATAGTTTCAA
CTATACAGCGTAAATATAAGGGTATTAAAATACAAGAGGGTGTGGTTGATTATGGTGCTAGATTTTACTTTTACACC
AGTAAAACAACTGTAGCGTCACTTATCAACACACTTAACGATCTAAATGAAACTCTTGTTACAATGCCACTTGGCTA
TGTAACACATGGCTTAAATTTGGAAGAAGCTGCTCGGTATATGAGATCTCTCAAAGTGCCAGCTACAGTTTCTGTTT
CTTCACCTGATGCTGTTACAGCGTATAATGGTTATCTTACTTCTTCTTCTAAAACACCTGAAGAACATTTTATTGAA
ACCATCTCACTTGCTGGTTCCTATAAAGATTGGTCCTATTCTGGACAATCTACACAACTAGGTATAGAATTTCTTAA
GAGAGGTGATAAAAGTGTATATTACACTAGTAATCCTACCACATTCCACCTAGATGGTGAAGTTATCACCTTTGACA
ATCTTAAGACACTTCTTTCTTTGAGAGAAGTGAGGACTATTAAGGTGTTTACAACAGTAGACAACATTAACCTCCAC
ACGCAAGTTGTGGACATGTCAATGACATATGGACAACAGTTTGGTCCAACTTATTTGGATGGAGCTGATGTTACTAA
AATAAAACCTCATAATTCACATGAAGGTAAAACATTTTATGTTTTACCTAATGATGACACTCTACGTGTTGAGGCTT
TTGAGTACTACCACACAACTGATCCTAGTTTTCTGGGTAGGTACATGTCAGCATTAAATCACACTAAAAAGTGGAAA
TACCCACAAGTTAATGGTTTAACTTCTATTAAATGGGCAGATAACAACTGTTATCTTGCCACTGCATTGTTAACACT
CCAACAAATAGAGTTGAAGTTTAATCCACCTGCTCTACAAGATGCTTATTACAGAGCAAGGGCTGGTGAAGCTGCTA
ACTTTTGTGCACTTATCTTAGCCTACTGTAATAAGACAGTAGGTGAGTTAGGTGATGTTAGAGAAACAATGAGTTAC
TTGTTTCAACATGCCAATTTAGATTCTTGCAAAAGAGTCTTGAACGTGGTGTGTAAAACTTGTGGACAACAGCAGAC
AACCCTTAAGGGTGTAGAAGCTGTTATGTACATGGGCACACTTTCTTATGAACAATTTAAGAAAGGTGTTCAGATAC
CTTGTACGTGTGGTAAACAAGCTACAAAATATCTAGTACAACAGGAGTCACCTTTTGTTATGATGTCAGCACCACCT
GCTCAGTATGAACTTAAGCATGGTACATTTACTTGTGCTAGTGAGTACACTGGTAATTACCAGTGTGGTCACTATAA
ACATATAACTTCTAAAGAAACTTTGTATTGCATAGACGGTGCTTTACTTACAAAGTCCTCAGAATACAAAGGTCCTA
TTACGGATGTTTTCTACAAAGAAAACAGTTACACAACAACCATAAAACCAGTTACTTATAAATTGGATGGTGTTGTT
TGTACAGAAATTGACCCTAAGTTGGACAATTATTATAAGAAAGACAATTCTTATTTCACAGAGCAACCAATTGATCT
TGTACCAAACCAACCATATCCAAACGCAAGCTTCGATAATTTTAAGTTTGTATGTGATAATATCAAATTTGCTGATG
ATTTAAACCAGTTAACTGGTTATAAGAAACCTGCTTCAAGAGAGCTTAAAGTTACATTTTTCCCTGACTTAAATGGT
GATGTGGTGGCTATTGATTATAAACACTACACACCCTCTTTTAAGAAAGGAGCTAAATTGTTACATAAACCTATTGT
TTGGCATGTTAACAATGCAACTAATAAAGCCACGTATAAACCAAATACCTGGTGTATACGTTGTCTTTGGAGCACAA
AACCAGTTGAAACATCAAATTCGTTTGATGTACTGAAGTCAGAGGACGCGCAGGGAATGGATAATCTTGCCTGCGAA
GATCTAAAACCAGTCTCTGAAGAAGTAGTGGAAAATCCTACCATACAGAAAGACGTTCTTGAGTGTAATGTGAAAAC
TACCGAAGTTGTAGGAGACATTATACTTAAACCAGCAAATAATAGTTTAAAAATTACAGAAGAGGTTGGCCACACAG
ATCTAATGGCTGCTTATGTAGACAATTCTAGTCTTACTATTAAGAAACCTAATGAATTATCTAGAGTATTAGGTTTG
AAAACCCTTGCTACTCATGGTTTAGCTGCTGTTAATAGTGTCCCTTGGGATACTATAGCTAATTATGCTAAGCCTTT
TCTTAACAAAGTTGTTAGTACAACTACTAACATAGTTACACGGTGTTTAAACCGTGTTTGTACTAATTATATGCCTT
ATTTCTTTACTTTATTGCTACAATTGTGTACTTTTACTAGAAGTACAAATTCTAGAATTAAAGCATCTATGCCGACT
ACTATAGCAAAGAATACTGTTAAGAGTGTCGGTAAATTTTGTCTAGAGGCTTCATTTAATTATTTGAAGTCACCTAA
TTTTTCTAAACTGATAAATATTATAATTTGGTTTTTACTATTAAGTGTTTGCCTAGGTTCTTTAATCTACTCAACCG
CTGCTTTAGGTGTTTTAATGTCTAATTTAGGCATGCCTTCTTACTGTACTGGTTACAGAGAAGGCTATTTGAACTCT
ACTAATGTCACTATTGCAACCTACTGTACTGGTTCTATACCTTGTAGTGTTTGTCTTAGTGGTTTAGATTCTTTAGA
CACCTATCCTTCTTTAGAAACTATACAAATTACCATTTCATCTTTTAAATGGGATTTAACTGCTTTTGGCTTAGTTG
CAGAGTGGTTTTTGGCATATATTCTTTTCACTAGGTTTTTCTATGTACTTGGATTGGCTGCAATCATGCAATTGTTT
TTCAGCTATTTTGCAGTACATTTTATTAGTAATTCTTGGCTTATGTGGTTAATAATTAATCTTGTACAAATGGCCCC
GATTTCAGCTATGGTTAGAATGTACATCTTCTTTGCATCATTTTATTATGTATGGAAAAGTTATGTGCATGTTGTAG
ACGGTTGTAATTCATCAACTTGTATGATGTGTTACAAACGTAATAGAGCAACAAGAGTCGAATGTACAACTATTGTT
AATGGTGTTAGAAGGTCCTTTTATGTCTATGCTAATGGAGGTAAAGGCTTTTGCAAACTACACAATTGGAATTGTGT
TAATTGTGATACATTCTGTGCTGGTAGTACATTTATTAGTGATGAAGTTGCGAGAGACTTGTCACTACAGTTTAAAA
GACCAATAAATCCTACTGACCAGTCTTCTTACATCGTTGATAGTGTTACAGTGAAGAATGGTTCCATCCATCTTTAC
TTTGATAAAGCTGGTCAAAAGACTTATGAAAGACATTCTCTCTCTCATTTTGTTAACTTAGACAACCTGAGAGCTAA
TAACACTAAAGGTTCATTGCCTATTAATGTTATAGTTTTTGATGGTAAATCAAAATGTGAAGAATCATCTGCAAAAT
CAGCGTCTGTTTACTACAGTCAGCTTATGTGTCAACCTATACTGTTACTAGATCAGGCATTAGTGTCTGATGTTGGT
GATAGTGCGGAAGTTGCAGTTAAAATGTTTGATGCTTACGTTAATACGTTTTCATCAACTTTTAACGTACCAATGGA
AAAACTCAAAACACTAGTTGCAACTGCAGAAGCTGAACTTGCAAAGAATGTGTCCTTAGACAATGTCTTATCTACTT
TTATTTCAGCAGCTCGGCAAGGGTTTGTTGATTCAGATGTAGAAACTAAAGATGTTGTTGAATGTCTTAAATTGTCA
CATCAATCTGACATAGAAGTTACTGGCGATAGTTGTAATAACTATATGCTCACCTATAACAAAGTTGAAAACATGAC
ACCCCGTGACCTTGGTGCTTGTATTGACTGTAGTGCGCGTCATATTAATGCGCAGGTAGCAAAAAGTCACAACATTG
CTTTGATATGGAACGTTAAAGATTTCATGTCATTGTCTGAACAACTACGAAAACAAATACGTAGTGCTGCTAAAAAG
AATAACTTACCTTTTAAGTTGACATGTGCAACTACTAGACAAGTTGTTAATGTTGTAACAACAAAGATAGCACTTAA
GGGTGGTAAAATTGTTAATAATTGGTTGAAGCAGTTAATTAAAGTTACACTTGTGTTCCTTTTTGTTGCTGCTATTT
TCTATTTAATAACACCTGTTCATGTCATGTCTAAACATACTGACTTTTCAAGTGAAATCATAGGATACAAGGCTATT

GATGGTGGTGTCACTCGTGACATAGCATCTACAGATACTTGTTTTGCTAACAAACATGCTGATTTTGACACATGGTT
TAGCCAGCGTGGTGGTAGTTATACTAATGACAAAGCTTGCCCATTGATTGCTGCAGTCATAACAAGAGAAGTGGGTT
TTGTCGTGCCTGGTTTGCCTGGCACGATATTACGCACAACTAATGGTGACTTTTTGCATTTCTTACCTAGAGTTTTT
AGTGCAGTTGGTAACATCTGTTACACACCATCAAAACTTATAGAGTACACTGACTTTGCAACATCAGCTTGTGTTTT
GGCTGCTGAATGTACAATTTTTAAAGATGCTTCTGGTAAGCCAGTACCATATTGTTATGATACCAATGTACTAGAAG
GTTCTGTTGCTTATGAAAGTTTACGCCCTGACACACGTTATGTGCTCATGGATGGCTCTATTATTCAATTTCCTAAC
ACCTACCTTGAAGGTTCTGTTAGAGTGGTAACAACTTTTGATTCTGAGTACTGTAGGCACGGCACTTGTGAAAGATC
AGAAGCTGGTGTTTGTGTATCTACTAGTGGTAGATGGGTACTTAACAATGATTATTACAGATCTTTACCAGGAGTTT
TCTGTGGTGTAGATGCTGTAAATTTACTTACTAATATGTTTACACCACTAATTCAACCTATTGGTGCTTTGGACATA
TCAGCATCTATAGTAGCTGGTGGTATTGTAGCTATCGTAGTAACATGCCTTGCCTACTATTTTATGAGGTTTAGAAG
AGCTTTTGGTGAATACAGTCATGTAGTTGCCTTTAATACTTTACTATTCCTTATGTCATTCACTGTACTCTGTTTAA
CACCAGTTTACTCATTCTTACCTGGTGTTTATTCTGTTATTTACTTGTACTTGACATTTTATCTTACTAATGATGTT
TCTTTTTTAGCACATATTCAGTGGATGGTTATGTTCACACCTTTAGTACCTTTCTGGATAACAATTGCTTATATCAT
TTGTATTTCCACAAAGCATTTCTATTGGTTCTTTAGTAATTACCTAAAGAGACGTGTAGTCTTTAATGGTGTTTCCT
TTAGTACTTTTGAAGAAGCTGCGCTGTGCACCTTTTTGTTAAATAAAGAAATGTATCTAAAGTTGCGTAGTGATGTG
CTATTACCTCTTACGCAATATAATAGATACTTAGCTCTTTATAATAAGTACAAGTATTTTAGTGGAGCAATGGATAC
AACTAGCTACAGAGAAGCTGCTTGTTGTCATCTCGCAAAGGCTCTCAATGACTTCAGTAACTCAGGTTCTGATGTTC
TTTACCAACCACCACAAACCTCTATCACCTCAGCTGTTTTGCAGAGTGGTTTTAGAAAAATGGCATTCCCATCTGGT
AAAGTTGAGGGTTGTATGGTACAAGTAACTTGTGGTACAACTACACTTAACGGTCTTTGGCTTGATGACGTAGTTTA
CTGTCCAAGACATGTGATCTGCACCTCTGAAGACATGCTTAACCCTAATTATGAAGATTTACTCATTCGTAAGTCTA
ATCATAATTTCTTGGTACAGGCTGGTAATGTTCAACTCAGGGTTATTGGACATTCTATGCAAAATTGTGTACTTAAG
CTTAAGGTTGATACAGCCAATCCTAAGACACCTAAGTATAAGTTTGTTCGCATTCAACCAGGACAGACTTTTTCAGT
GTTAGCTTGTTACAATGGTTCACCATCTGGTGTTTACCAATGTGCTATGAGGCCCAATTTCACTATTAAGGGTTCAT
TCCTTAATGGTTCATGTGGTAGTGTTGGTTTTAACATAGATTATGACTGTGTCTCTTTTTGTTACATGCACCATATG
GAATTACCAACTGGAGTTCATGCTGGCACAGACTTAGAAGGTAACTTTTATGGACCTTTTGTTGACAGGCAAACAGC
ACAAGCAGCTGGTACGGACACAACTATTACAGTTAATGTTTTAGCTTGGTTGTACGCTGCTGTTATAAATGGAGACA
GGTGGTTTCTCAATCGATTTACCACAACTCTTAATGACTTTAACCTTGTGGCTATGAAGTACAATTATGAACCTCTA
ACACAAGACCATGTTGACATACTAGGACCTCTTTCTGCTCAAACTGGAATTGCCGTTTTAGATATGTGTGCTTCATT
AAAAGAATTACTGCAAAATGGTATGAATGGACGTACCATATTGGGTAGTGCTTTATTAGAAGATGAATTTACACCTT
TTGATGTTGTTAGACAATGCTCAGGTGTTACTTTCCAAAGTGCAGTGAAAAGAACAATCAAGGGTACACACCACTGG
TTGTTACTCACAATTTTGACTTCACTTTTAGTTTTAGTCCAGAGTACTCAATGGTCTTTGTTCTTTTTTTTGTATGA
AAATGCCTTTTTACCTTTTGCTATGGGTATTATTGCTATGTCTGCTTTTGCAATGATGTTTGTCAAACATAAGCATG
CATTTCTCTGTTTGTTTTTGTTACCTTCTCTTGCCACTGTAGCTTATTTTAATATGGTCTATATGCCTGCTAGTTGG
GTGATGCGTATTATGACATGGTTGGATATGGTTGATACTAGTTTGTCTGGTTTTAAGCTAAAAGACTGTGTTATGTA
TGCATCAGCTGTAGTGTTACTAATCCTTATGACAGCAAGAACTGTGTATGATGATGGTGCTAGGAGAGTGTGGACAC
TTATGAATGTCTTGACACTCGTTTATAAAGTTTATTATGGTAATGCTTTAGATCAAGCCATTTCCATGTGGGCTCTT
ATAATCTCTGTTACTTCTAACTACTCAGGTGTAGTTACAACTGTCATGTTTTTGGCCAGAGGTATTGTTTTTATGTG
TGTTGAGTATTGCCCTATTTTCTTCATAACTGGTAATACACTTCAGTGTATAATGCTAGTTTATTGTTTCTTAGGCT
ATTTTTGTACTTGTTACTTTGGCCTCTTTTGTTTACTCAACCGCTACTTTAGACTGACTCTTGGTGTTTATGATTAC
TTAGTTTCTACACAGGAGTTTAGATATATGAATTCACAGGGACTACTCCCACCCAAGAATAGCATAGATGCCTTCAA
ACTCAACATTAAATTGTTGGGTGTTGGTGGCAAACCTTGTATCAAAGTAGCCACTGTACAGTCTAAAATGTCAGATG
TAAAGTGCACATCAGTAGTCTTACTCTCAGTTTTGCAACAACTCAGAGTAGAATCATCATCTAAATTGTGGGCTCAA
TGTGTCCAGTTACACAATGACATTCTCTTAGCTAAAGATACTACTGAAGCCTTTGAAAAAATGGTTTCACTACTTTC
TGTTTTGCTTTCCATGCAGGGTGCTGTAGACATAAACAAGCTTTGTGAAGAAATGCTGGACAACAGGGCAACCTTAC
AAGCTATAGCCTCAGAGTTTAGTTCCCTTCCATCATATGCAGCTTTTGCTACTGCTCAAGAAGCTTATGAGCAGGCT
GTTGCTAATGGTGATTCTGAAGTTGTTCTTAAAAAGTTGAAGAAGTCTTTGAATGTGGCTAAATCTGAATTTGACCG
TGATGCAGCCATGCAACGTAAGTTGGAAAAGATGGCTGATCAAGCTATGACCCAAATGTATAAACAGGCTAGATCTG
AGGACAAGAGGGCAAAAGTTACTAGTGCTATGCAGACAATGCTTTTCACTATGCTTAGAAAGTTGGATAATGATGCA
CTCAACAACATTATCAACAATGCAAGAGATGGTTGTGTTCCCTTGAACATAATACCTCTTACAACAGCAGCCAAACT
AATGGTTGTCATACCAGACTATAACACATATAAAAATACGTGTGATGGTACAACATTTACTTATGCATCAGCATTGT
GGGAAATCCAACAGGTTGTAGATGCAGATAGTAAAATTGTTCAACTTAGTGAAATTAGTATGGACAATTCACCTAAT
TTAGCATGGCCTCTTATTGTAACAGCTTTAAGGGCCAATTCTGCTGTCAAATTACAGAATAATGAGCTTAGTCCTGT
TGCACTACGACAGATGTCTTGTGCTGCCGGTACTACACAAACTGCTTGCACTGATGACAATGCGTTAGCTTACTACA
ACACAACAAAGGGAGGTAGGTTTGTACTTGCACTGTTATCCGATTTACAGGATTTGAAATGGGCTAGATTCCCTAAG
AGTGATGGAACTGGTACTATCTATACAGAACTGGAACCACCTTGTAGGTTTGTTACAGACACACCTAAAGGTCCTAA
AGTGAAGTATTTATACTTTATTAAAGGATTAAACAACCTAAATAGAGGTATGGTACTTGGTAGTTTAGCTGCCACAG
TACGTCTACAAGCTGGTAATGCAACAGAAGTGCCTGCCAATTCAACTGTATTATCTTTCTGTGCTTTTGCTGTAGAT
GCTGCTAAAGCTTACAAAGATTATCTAGCTAGTGGGGGACAACCAATCACTAATTGTGTTAAGATGTTGTGTACACA
CACTGGTACTGGTCAGGCAATAACAGTTACACCGGAAGCCAATATGGATCAAGAATCCTTTGGTGGTGCATCGTGTT

GTCTGTACTGCCGTTGCCACATAGATCATCCAAATCCTAAAGGATTTTGTGACTTAAAAGGTAAGTATGTACAAATA
CCTACAACTTGTGCTAATGACCCTGTGGGTTTTACACTTAAAAACACAGTCTGTACCGTCTGCGGTATGTGGAAAGG
TTATGGCTGTAGTTGTGATCAACTCCGCGAACCCATGCTTCAGTCAGCTGATGCACAATCGTTTTTAAACGGGTTTG
CGGTGTAAGTGCAGCCCGTCTTACACCGTGCGGCACAGGCACTAGTACTGATGTCGTATACAGGGCTTTTGACATCT
ACAATGATAAAGTAGCTGGTTTTGCTAAATTCCTAAAAACTAATTGTTGTCGCTTCCAAGAAAAGGACGAAGATGAC
AATTTAATTGATTCTTACTTTGTAGTTAAGAGACACACTTTCTCTAACTACCAACATGAAGAAACAATTTATAATTT
ACTTAAGGATTGTCCAGCTGTTGCTAAACATGACTTCTTTAAGTTTAGAATAGACGGTGACATGGTACCACATATAT
CACGTCAACGTCTTACTAAATACACAATGGCAGACCTCGTCTATGCTTTAAGGCATTTTGATGAAGGTAATTGTGAC
ACATTAAAAGAAATACTTGTCACATACAATTGTTGTGATGATGATTATTTCAATAAAAAGGACTGGTATGATTTTGT
AGAAAACCCAGATATATTACGCGTATACGCCAACTTAGGTGAACGTGTACGCCAAGCTTTGTTAAAAACAGTACAAT
TCTGTGATGCCATGCGAAATGCTGGTATTGTTGGTGTACTGACATTAGATAATCAAGATCTCAATGGTAACTGGTAT
GATTTCGGTGATTTCATACAAACCACGCCAGGTAGTGGAGTTCCTGTTGTAGATTCTTATTATTCATTGTTAATGCC
TATATTAACCTTGACCAGGGCTTTAACTGCAGAGTCACATGTTGACACTGACTTAACAAAGCCTTACATTAAGTGGG
ATTTGTTAAAATATGACTTCACGGAAGAGAGGTTAAAACTCTTTGACCGTTATTTTAAATATTGGGATCAGACATAC
CACCCAAATTGTGTTAACTGTTTGGATGACAGATGCATTCTGCATTGTGCAAACTTTAATGTTTTATTCTCTACAGT
GTTCCCACCTACAAGTTTTGGACCACTAGTGAGAAAAATATTTGTTGATGGTGTTCCATTTGTAGTTTCAACTGGAT
ACCACTTCAGAGAGCTAGGTGTTGTACATAATCAGGATGTAAACTTACATAGCTCTAGACTTAGTTTTAAGGAATTA
CTTGTGTATGCTGCTGACCCTGCTATGCACGCTGCTTCTGGTAATCTATTACTAGATAAACGCACTACGTGCTTTTC
AGTAGCTGCACTTACTAACAATGTTGCTTTTCAAACTGTCAAACCCGGTAATTTTAACAAAGACTTCTATGACTTTG
CTGTGTCTAAGGGTTTCTTTAAGGAAGGAAGTTCTGTTGAATTAAAACACTTCTTCTTTGCTCAGGATGGTAATGCT
GCTATCAGCGATTATGACTACTATCGTTATAATCTACCAACAATGTGTGATATCAGACAACTACTATTTGTAGTTGA
AGTTGTTGATAAGTACTTTGATTGTTACGATGGTGGCTGTATTAATGCTAACCAAGTCATCGTCAACAACCTAGACA
AATCAGCTGGTTTTCCATTTAATAAATGGGGTAAGGCTAGACTTTATTATGATTCAATGAGTTATGAGGATCAAGAT
GCACTTTTCGCATATACAAAACGTAATGTCATCCCTACTATAACTCAAATGAATCTTAAGTATGCCATTAGTGCAAA
GAATAGAGCTCGCACCGTAGCTGGTGTCTCTATCTGTAGTACTATGACCAATAGACAGTTTCATCAAAAATTATTGA
AATCAATAGCCGCCACTAGAGGAGCTACTGTAGTAATTGGAACAAGCAAATTCTATGGTGGTTGGCACAACATGTTA
AAAACTGTTTATAGTGATGTAGAAAACCCTCACCTTATGGGTTGGGATTATCCTAAATGTGATAGAGCCATGCCTAA
CATGCTTAGAATTATGGCCTCACTTGTTCTTGCTCGCAAACATACAACGTGTTGTAGCTTGTCACACCGTTTCTATA
GATTAGCTAATGAGTGTGCTCAAGTATTGAGTGAAATGGTCATGTGTGGCGGTTCACTATATGTTAAACCAGGTGGA
ACCTCATCAGGAGATGCCACAACTGCTTATGCTAATAGTGTTTTTAACATTTGTCAAGCTGTCACGGCCAATGTTAA
TGCACTTTTATCTACTGATGGTAACAAAATTGCCGATAAGTATGTCCGCAATTTACAACACAGACTTTATGAGTGTC
TCTATAGAAATAGAGATGTTGACACAGACTTTGTGAATGAGTTTTACGCATATTTGCGTAAACATTTCTCAATGATG
ATACTCTCTGACGATGCTGTTGTGTGTTTCAATAGCACTTATGCATCTCAAGGTCTAGTGGCTAGCATAAAGAACTT
TAAGTCAGTTCTTTATTATCAAAACAATGTTTTTATGTCTGAAGCAAAATGTTGGACTGAGACTGACCTTACTAAAG
GACCTCATGAATTTTGCTCTCAACATACAATGCTAGTTAAACAGGGTGATGATTATGTGTACCTTCCTTACCCAGAT
CCATCAAGAATCCTAGGGGCCGGCTGTTTTGTAGATGATATCGTAAAAACAGATGGTACACTTATGATTGAACGGTT
CGTGTCTTTAGCTATAGATGCTTACCCACTTACTAAACATCCTAATCAGGAGTATGCTGATGTCTTTCATTTGTACT
TACAATACATAAGAAAGCTACATGATGAGTTAACAGGACACATGTTAGACATGTATTCTGTTATGCTTACTAATGAT
AACACTTCAAGGTATTGGGAACCTGAGTTTTATGAGGCTATGTACACACCGCATACAGTCTTACAGGCTGTTGGGGC
TTGTGTTCTTTGCAATTCACAGACTTCATTAAGATGTGGTGCTTGCATACGTAGACCATTCTTATGTTGTAAATGCT
GTTACGACCATGTCATATCAACATCACATAAATTAGTCTTGTCTGTTAATCCGTATGTTTGCAATGCTCCAGGTTGT
GATGTCACAGATGTGACTCAACTTTACTTAGGAGGTATGAGCTATTATTGTAAATCACATAAACCACCCATTAGTTT
TCCATTGTGTGCTAATGGACAAGTTTTTGGTTTATATAAAAATACATGTGTTGGTAGCGATAATGTTACTGACTTTA
ATGCAATTGCAACATGTGACTGGACAAATGCTGGTGATTACATTTTAGCTAACACCTGTACTGAAAGACTCAAGCTT
TTTGCAGCAGAAACGCTCAAAGCTACTGAGGAGACATTTAAACTGTCTTATGGTATTGCTACTGTACGTGAAGTGCT
GTCTGACAGAGAATTACATCTTTCATGGGAAGTTGGTAAACCTAGACCACCACTTAACCGAAATTATGTCTTTACTG
GTTATCGTGTAACTAAAAACAGTAAAGTACAAATAGGAGAGTACACCTTTGAAAAAGGTGACTATGGTGATGCTGTT
GTTTACCGAGGTACAACAACTTACAAATTAAATGTTGGTGATTATTTTGTGCTGACATCACATACAGTAATGCCATT
AAGTGCACCTACACTAGTGCCACAAGAGCACTATGTTAGAATTACTGGCTTATACCCAACACTCAATATCTCAGATG
AGTTTTCTAGCAATGTTGCAAATTATCAAAAGGTTGGTATGCAAAAGTATTCTACACTCCAGGGACCACCTGGTACT
GGTAAGAGTCATTTTGCTATTGGCCTAGCTCTCTACTACCCTTCTGCTCGCATAGTGTATACAGCTTGCTCTCATGC
CGCTGTTGATGCACTATGTGAGAAGGCATTAAAATATTTGCCTATAGATAAATGTAGTAGAATTATACCTGCACGTG
CTCGTGTAGAGTGTTTTGATAAATTCAAAGTGAATTCAACATTAGAACAGTATGTCTTTTGTACTGTAAATGCATTG
CCTGAGACGACAGCAGATATAGTTGTCTTTGATGAAATTTCAATGGCCACAAATTATGATTTGAGTGTTGTCAATGC
CAGATTACGTGCTAAGCACTATGTGTACATTGGCGACCCTGCTCAATTACCTGCACCACGCACATTGCTAACTAAGG
GCACACTAGAACCAGAATATTTCAATTCAGTGTGTAGACTTATGAAAACTATAGGTCCAGACATGTTCCTCGGAACT
TGTCGGCGTTGTCCTGCTGAAATTGTTGACACTGTGAGTGCTTTGGTTTATGATAATAAGCTTAAAGCACATAAAGA
CAAATCAGCTCAATGCTTTAAAATGTTTTATAAGGGTGTTATCACGCATGATGTTTCATCTGCAATTAACAGGCCAC
AAATAGGCGTGGTAAGAGAATTCCTTACACGTAACCCTGCTTGGAGAAAAGCTGTCTTTATTTCACCTTATAATTCA

CAGAATGCTGTAGCCTCAAAGATTTTGGGACTACCAACTCAAACTGTTGATTCATCACAGGGCTCAGAATATGACTA
TGTCATATTCACTCAAACCACTGAAACAGCTCACTCTTGTAATGTAAACAGATTTAATGTTGCTATTACCAGAGCAA
AAGTAGGCATACTTTGCATAATGTCTGATAGAGACCTTTATGACAAGTTGCAATTTACAAGTCTTGAAATTCCACGT
AGGAATGTGGCAACTTTACAAGCTGAAAATGTAACAGGACTCTTTAAAGATTGTAGTAAGGTAATCACTGGGTTACA
TCCTACACAGGCACCTACACACCTCAGTGTTGACACTAAATTCAAAACTGAAGGTTTATGTGTTGACATACCTGGCA
TACCTAAGGACATGACCTATAGAAGACTCATCTCTATGATGGGTTTTAAAATGAATTATCAAGTTAATGGTTACCCT
AACATGTTTATCACCCGCGAAGAAGCTATAAGACATGTACGTGCATGGATTGGCTTCGATGTCGAGGGGTGTCATGC
TACTAGAGAAGCTGTTGGTACCAATTTACCTTTACAGCTAGGTTTTTCTACAGGTGTTAACCTAGTTGCTGTACCTA
CAGGTTATGTTGATACACCTAATAATACAGATTTTTCCAGAGTTAGTGCTAAACCACCGCCTGGAGATCAATTTAAA
CACCTCATACCACTTATGTACAAAGGACTTCCTTGGAATGTAGTGCGTATAAAGATTGTACAAATGTTAAGTGACAC
ACTTAAAAATCTCTCTGACAGAGTCGTATTTGTCTTATGGGCACATGGCTTTGAGTTGACATCTATGAAGTATTTTG
TGAAAATAGGACCTGAGCGCACCTGTTGTCTATGTGATAGACGTGCCACATGCTTTTCCACTGCTTCAGACACTTAT
GCCTGTTGGCATCATTCTATTGGATTTGATTACGTCTATAATCCGTTTATGATTGATGTTCAACAATGGGGTTTTAC
AGGTAACCTACAAAGCAACCATGATCTGTATTGTCAAGTCCATGGTAATGCACATGTAGCTAGTTGTGATGCAATCA
TGACTAGGTGTCTAGCTGTCCACGAGTGCTTTGTTAAGCGTGTTGACTGGACTATTGAATATCCTATAATTGGTGAT
GAACTGAAGATTAATGCGGCTTGTAGAAAGGTTCAACACATGGTTGTTAAAGCTGCATTATTAGCAGACAAATTCCC
AGTTCTTCACGACATTGGTAACCCTAAAGCTATTAAGTGTGTACCTCAAGCTGATGTAGAATGGAAGTTCTATGATG
CACAGCCTTGTAGTGACAAAGCTTATAAAATAGAAGAATTATTCTATTCTTATGCCACACATTCTGACAAATTCACA
GATGGTGTATGCCTATTTTGGAATTGCAATGTCGATAGATATCCTGCTAATTCCATTGTTTGTAGATTTGACACTAG
AGTGCTATCTAACCTTAACTTGCCTGGTTGTGATGGTGGCAGTTTGTATGTAAATAAACATGCATTCCACACACCAG
CTTTTGATAAAAGTGCTTTTGTTAATTTAAAACAATTACCATTTTTCTATTACTCTGACAGTCCATGTGAGTCTCAT
GGAAAACAAGTAGTGTCAGATATAGATTATGTACCACTAAAGTCTGCTACGTGTATAACACGTTGCAATTTAGGTGG
TGCTGTCTGTAGACATCATGCTAATGAGTACAGATTGTATCTCGATGCTTATAACATGATGATCTCAGCTGGCTTTA
GCTTGTGGGTTTACAAACAATTTGATACTTATAACCTCTGGAACACTTTTACAAGACTTCAGAGTTTAGAAAATGTG
GCTTTTAATGTTGTAAATAAGGGACACTTTGATGGACAACAGGGTGAAGTACCAGTTTCTATCATTAATAACACTGT
TTACACAAAAGTTGATGGTGTTGATGTAGAATTGTTTGAAAATAAAACAACATTACCTGTTAATGTAGCATTTGAGC
TTTGGGCTAAGCGCAACATTAAACCAGTACCAGAGGTGAAAATACTCAATAATTTGGGTGTGGACATTGCTGCTAAT
ACTGTGATCTGGGACTACAAAAGAGATGCTCCAGCACATATATCTACTATTGGTGTTTGTTCTATGACTGACATAGC
CAAGAAACCAACTGAAACGATTTGTGCACCACTCACTGTCTTTTTTGATGGTAGAGTTGATGGTCAAGTAGACTTAT
TTAGAAATGCCCGTAATGGTGTTCTTATTACAGAAGGTAGTGTTAAAGGTTTACAACCATCTGTAGGTCCCAAACAA
GCTAGTCTTAATGGAGTCACATTAATTGGAGAAGCCGTAAAAACACAGTTCAATTATTATAAGAAAGTTGATGGTGT
TGTCCAACAATTACCTGAAACTTACTTTACTCAGAGTAGAAATTTACAAGAATTTAAACCCAGGAGTCAAATGGAAA
TTGATTTCTTAGAATTAGCTATGGATGAATTCATTGAACGGTATAAATTAGAAGGCTATGCCTTCGAACATATCGTT
TATGGAGATTTTAGTCATAGTCAGTTAGGTGGTTTACATCTACTGATTGGACTAGCTAAACGTTTTAAGGAATCACC
TTTTGAATTAGAAGATTTTATTCCTATGGACAGTACAGTTAAAAACTATTTCATAACAGATGCGCAAACAGGTTCAT
CTAAGTGTGTGTGTTCTGTTATTGATTTATTACTTGATGATTTTGTTGAAATAATAAAATCCCAAGATTTATCTGTA
GTTTCTAAGGTTGTCAAAGTGACTATTGACTATACAGAAATTTCATTTATGCTTTGGTGTAAAGATGGCCATGTAGA
AACATTTTACCCAAAATTACAATCTAGTCAAGCGTGGCAACCGGGTGTTGCTATGCCTAATCTTTACAAAATGCAAA
GAATGCTATTAGAAAAGTGTGACCTTCAAAATTATGGTGATAGTGCAACATTACCTAAAGGCATAATGATGAATGTC
GCAAAATATACTCAACTGTGTCAATATTTAAACACATTAACATTAGCTGTACCCTATAATATGAGAGTTATACATTT
TGGTGCTGGTTCTGATAAAGGAGTTGCACCAGGTACAGCTGTTTTAAGACAGTGGTTGCCTACGGGTACGCTGCTTG
TCGATTCAGATCTTAATGACTTTGTCTCTGATGCAGATTCAACTTTGATTGGTGATTGTGCAACTGTACATACAGCT
AATAAATGGGATCTCATTATTAGTGATATGTACGACCCTAAGACTAAAAATGTTACAAAAGAAAATGACTCTAAAGA
GGGTTTTTTCACTTACATTTGTGGGTTTATACAACAAAAGCTAGCTCTTGGAGGTTCCGTGGCTATAAAGATAACAG
AACATTCTTGGAATGCTGATCTTTATAAGCTCATGGGACACTTCGCATGGTGGACAGCCTTTGTTACTAATGTGAAT
GCGTCATCATCTGAAGCATTTTTAATTGGATGTAATTATCTTGGCAAACCACGCGAACAAATAGATGGTTATGTCAT
GCATGCAAATTACATATTTTGGAGGAATACAAATCCAATTCAGTTGTCTTCCTATTCTTTATTTGACATGAGTAAAT
TTCCCCTTAAATTAAGGGGTACTGCTGTTATGTCTTTAAAAGAAGGTCAAATCAATGATATGATTTTATCTCTTCTT
AGTAAAGGTAGACTTATAATTAGAGAAAACAACAGAGTTGTTATTTCTAGTGATGTTCTTGTTAACAACTAAACGAA
CAATGTTTGTTTTTCTTGTTTTATTGCCACTAGTCTCTAGTCAGTGTGTTAATCTTACAACCAGAACTCAATTACCC
CCTGCATACACTAATTCTTTCACACGTGGTGTTTATTACCCTGACAAAGTTTTCAGATCCTCAGTTTTACATTCAAC
TCAGGACTTGTTCTTACCTTTCTTTTCCAATGTTACTTGGTTCCATGCTATACATGTCTCTGGGACCAATGGTACTA
AGAGGTTTGATAACCCTGTCCTACCATTTAATGATGGTGTTTATTTTGCTTCCACTGAGAAGTCTAACATAATAAGA
GGCTGGATTTTTGGTACTACTTTAGATTCGAAGACCCAGTCCCTACTTATTGTTAATAACGCTACTAATGTTGTTAT
TAAAGTCTGTGAATTTCAATTTTGTAATGATCCATTTTTGGGTGTTTATTACCACAAAAACAACAAAAGTTGGATGG
AAAGTGAGTTCAGAGTTTATTCTAGTGCGAATAATTGCACTTTTGAATATGTCTCTCAGCCTTTTCTTATGGACCTT
GAAGGAAAACAGGGTAATTTCAAAAATCTTAGGGAATTTGTGTTTAAGAATATTGATGGTTATTTTAAAATATATTC
TAAGCACACGCCTATTAATTTAGTGCGTGATCTCCCTCAGGGTTTTTCGGCTTTAGAACCATTGGTAGATTTGCCAA
TAGGTATTAACATCACTAGGTTTCAAACTTTACTTGCTTTACATAGAAGTTATTTGACTCCTGGTGATTCTTCTTCA

GGTTGGACAGCTGGTGCTGCAGCTTATTATGTGGGTTATCTTCAACCTAGGACTTTTCTATTAAAATATAATGAAAA
TGGAACCATTACAGATGCTGTAGACTGTGCACTTGACCCTCTCTCAGAAACAAAGTGTACGTTGAAATCCTTCACTG
TAGAAAAAGGAATCTATCAAACTTCTAACTTTAGAGTCCAACCAACAGAATCTATTGTTAGATTTCCTAATATTACA
AACTTGTGCCCTTTTGGTGAAGTTTTTAACGCCACCAGATTTGCATCTGTTTATGCTTGGAACAGGAAGAGAATCAG
CAACTGTGTTGCTGATTATTCTGTCCTATATAATTCCGCATCATTTTCCACTTTTAAGTGTTATGGAGTGTCTCCTA
CTAAATTAAATGATCTCTGCTTTACTAATGTCTATGCAGATTCATTTGTAATTAGAGGTGATGAAGTCAGACAAATC
GCTCCAGGGCAAACTGGAAAGATTGCTGATTATAATTATAAATTACCAGATGATTTTACAGGCTGCGTTATAGCTTG
GAATTCTAACAATCTTGATTCTAAGGTTGGTGGTAATTATAATTACCTGTATAGATTGTTTAGGAAGTCTAATCTCA
AACCTTTTGAGAGAGATATTTCAACTGAAATCTATCAGGCCGGTAGCACACCTTGTAATGGTGTTGAAGGTTTTAAT
TGTTACTTTCCTTTACAATCATATGGTTTCCAACCCACTAATGGTGTTGGTTACCAACCATACAGAGTAGTAGTACT
TTCTTTTGAACTTCTACATGCACCAGCAACTGTTTGTGGACCTAAAAAGTCTACTAATTTGGTTAAAAACAAATGTG
TCAATTTCAACTTCAATGGTTTAACAGGCACAGGTGTTCTTACTGAGTCTAACAAAAAGTTTCTGCCTTTCCAACAA
TTTGGCAGAGACATTGCTGACACTACTGATGCTGTCCGTGATCCACAGACACTTGAGATTCTTGACATTACACCATG
TTCTTTTGGTGGTGTCAGTGTTATAACACCAGGAACAAATACTTCTAACCAGGTTGCTGTTCTTTATCAGGATGTTA
ACTGCACAGAAGTCCCTGTTGCTATTCATGCAGATCAACTTACTCCTACTTGGCGTGTTTATTCTACAGGTTCTAAT
GTTTTTCAAACACGTGCAGGCTGTTTAATAGGGGCTGAACATGTCAACAACTCATATGAGTGTGACATACCCATTGG
TGCAGGTATATGCGCTAGTTATCAGACTCAGACTAATTCTCCTCGGCGGGCACGTAGTGTAGCTAGTCAATCCATCA
TTGCCTACACTATGTCACTTGGTGCAGAAAATTCAGTTGCTTACTCTAATAACTCTATTGCCATACCCACAAATTTT
ACTATTAGTGTTACCACAGAAATTCTACCAGTGTCTATGACCAAGACATCAGTAGATTGTACAATGTACATTTGTGG
TGATTCAACTGAATGCAGCAATCTTTTGTTGCAATATGGCAGTTTTTGTACACAATTAAACCGTGCTTTAACTGGAA
TAGCTGTTGAACAAGACAAAAACACCCAAGAAGTTTTTGCACAAGTCAAACAAATTTACAAAACACCACCAATTAAA
GATTTTGGTGGTTTTAATTTTTCACAAATATTACCAGATCCATCAAAACCAAGCAAGAGGTCATTTATTGAAGATCT
ACTTTTCAACAAAGTGACACTTGCAGATGCTGGCTTCATCAAACAATATGGTGATTGCCTTGGTGATATTGCTGCTA
GAGACCTCATTTGTGCACAAAAGTTTAACGGCCTTACTGTTTTGCCACCTTTGCTCACAGATGAAATGATTGCTCAA
TACACTTCTGCACTGTTAGCGGGTACAATCACTTCTGGTTGGACCTTTGGTGCAGGTGCTGCATTACAAATACCATT
TGCTATGCAAATGGCTTATAGGTTTAATGGTATTGGAGTTACACAGAATGTTCTCTATGAGAACCAAAAATTGATTG
CCAACCAATTTAATAGTGCTATTGGCAAAATTCAAGACTCACTTTCTTCCACAGCAAGTGCACTTGGAAAACTTCAA
GATGTGGTCAACCAAAATGCACAAGCTTTAAACACGCTTGTTAAACAACTTAGCTCCAATTTTGGTGCAATTTCAAG
TGTTTTAAATGATATCCTTTCACGTCTTGACAAAGTTGAGGCTGAAGTGCAAATTGATAGGTTGATCACAGGCAGAC
TTCAAAGTTTGCAGACATATGTGACTCAACAATTAATTAGAGCTGCAGAAATCAGAGCTTCTGCTAATCTTGCTGCT
ACTAAAATGTCAGAGTGTGTACTTGGACAATCAAAAAGAGTTGATTTTTGTGGAAAGGGCTATCATCTTATGTCCTT
CCCTCAGTCAGCACCTCATGGTGTAGTCTTCTTGCATGTGACTTATGTCCCTGCACAAGAAAAGAACTTCACAACTG
CTCCTGCCATTTGTCATGATGGAAAAGCACACTTTCCTCGTGAAGGTGTCTTTGTTTCAAATGGCACACACTGGTTT
GTAACACAAAGGAATTTTTATGAACCACAAATCATTACTACAGACAACACATTTGTGTCTGGTAACTGTGATGTTGT
AATAGGAATTGTCAACAACACAGTTTATGATCCTTTGCAACCTGAATTAGACTCATTCAAGGAGGAGTTAGATAAAT
ATTTTAAGAATCATACATCACCAGATGTTGATTTAGGTGACATCTCTGGCATTAATGCTTCAGTTGTAAACATTCAA
AAAGAAATTGACCGCCTCAATGAGGTTGCCAAGAATTTAAATGAATCTCTCATCGATCTCCAAGAACTTGGAAAGTA
TGAGCAGTATATAAAATGGCCATGGTACATTTGGCTAGGTTTTATAGCTGGCTTGATTGCCATAGTAATGGTGACAA
TTATGCTTTGCTGTATGACCAGTTGCTGTAGTTGTCTCAAGGGCTGTTGTTCTTGTGGATCCTGCTGCAAATTTGAT
GAAGACGACTCTGAGCCAGTGCTCAAAGGAGTCAAATTACATTACACATAAACGAACTTATGGATTTGTTTATGAGA
ATCTTCACAATTGGAACTGTAACTTTGAAGCAAGGTGAAATCAAGGATGCTACTCCTTCAGATTTTGTTCGCGCTAC
TGCAACGATACCGATACAAGCCTCACTCCCTTTCGGATGGCTTATTGTTGGCGTTGCACTTCTTGCTGTTTTTCAGA
GCGCTTCCAAAATCATAACCCTCAAAAAGAGATGGCAACTAGCACTCTCCAAGGGTGTTCACTTTGTTTGCAACTTG
CTGTTGTTGTTTGTAACAGTTTACTCACACCTTTTGCTCGTTGCTGCTGGCCTTGAAGCCCCTTTTCTCTATCTTTA
TGCTTTAGTCTACTTCTTGCAGAGTATAAACTTTGTAAGAATAATAATGAGGCTTTGGCTTTGCTGGAAATGCCGTT
CCAAAAACCCATTACTTTATGATGCCAACTATTTTCTTTGCTGGCATACTAATTGTTACGACTATTGTATACCTTAC
AATAGTGTAACTTCTTCAATTGTCATTACTTCAGGTGATGGCACAACAAGTCCTATTTCTGAACATGACTACCAGAT
TGGTGGTTATACTGAAAAATGGGAATCTGGAGTAAAAGACTGTGTTGTATTACACAGTTACTTCACTTCAGACTATT
ACCAGCTGTACTCAACTCAATTGAGTACAGACACTGGTGTTGAACATGTTACCTTCTTCATCTACAATAAAATTGTT
GATGAGCCTGAAGAACATGTCCAAATTCACACAATCGACGGTTCATCCGGAGTTGTTAATCCAGTAATGGAACCAAT
TTATGATGAACCGACGACGACTACTAGCGTGCCTTTGTAAGCACAAGCTGATGAGTACGAACTTATGTACTCATTCG
TTTCGGAAGAGACAGGTACGTTAATAGTTAATAGCGTACTTCTTTTTCTTGCTTTCGTGGTATTCTTGCTAGTTACA
CTAGCCATCCTTACTGCGCTTCGATTGTGTGCGTACTGCTGCAATATTGTTAACGTGAGTCTTGTAAAACCTTCTTT
TTACGTTTACTCTCGTGTTAAAAATCTGAATTCTTCTAGAGTTCCTGATCTTCTGGTCTAAACGAACTAAATATTAT
ATTAGTTTTTCTGTTTGGAACTTTAATTTTAGCCATGGCAGATTCCAACGGTACTATTACCGTTGAAGAGCTTAAAA
AGCTCCTTGAACAATGGAACCTAGTAATAGGTTTCCTATTCCTTACATGGATTTGTCTTCTACAATTTGCCTATGCC
AACAGGAATAGGTTTTTGTATATAATTAAGTTAATTTTCCTCTGGCTGTTATGGCCAGTAACTTTAGCTTGTTTTGT
GCTTGCTGCTGTTTACAGAATAAATTGGATCACCGGTGGAATTGCTATCGCAATGGCTTGTCTTGTAGGCTTGATGT
GGCTCAGCTACTTCATTGCTTCTTTCAGACTGTTTGCGCGTACGCGTTCCATGTGGTCATTCAATCCAGAAACTAAC

ATTCTTCTCAACGTGCCACTCCATGGCACTATTCTGACCAGACCGCTTCTAGAAAGTGAACTCGTAATCGGAGCTGT
GATCCTTCGTGGACATCTTCGTATTGCTGGACACCATCTAGGACGCTGTGACATCAAGGACCTGCCTAAAGAAATCA
CTGTTGCTACATCACGAACGCTTTCTTATTACAAATTGGGAGCTTCGCAGCGTGTAGCAGGTGACTCAGGTTTTGCT
GCATACAGTCGCTACAGGATTGGCAACTATAAATTAAACACAGACCATTCCAGTAGCAGTGACAATATTGCTTTGCT
TGTACAGTAAGTGACAACAGATGTTTCATCTCGTTGACTTTCAGGTTACTATAGCAGAGATATTACTAATTATTATG
AGGACTTTTAAAGTTTCCATTTGGAATCTTGATTACATCATAAACCTCATAATTAAAAATTTATCTAAGTCACTAAC
TGAGAATAAATATTCTCAATTAGATGAAGAGCAACCAATGGAGATTGATTAAACGAACATGAAAATTATTCTTTTCT
TGGCACTGATAACACTCGCTACTTGTGAGCTTTATCACTACCAAGAGTGTGTTAGAGGTACAACAGTACTTTTAAAA
GAACCTTGCTCTTCTGGAACATACGAGGGCAATTCACCATTTCATCCTCTAGCTGATAACAAATTTGCACTGACTTG
CTTTAGCACTCAATTTGCTTTTGCTTGTCCTGACGGCGTAAAACACGTCTATCAGTTACGTGCCAGATCAGTTTCAC
CTAAACTGTTCATCAGACAAGAGGAAGTTCAAGAACTTTACTCTCCAATTTTTCTTATTGTTGCGGCAATAGTGTTT
ATAACACTTTGCTTCACACTCAAAAGAAAGACAGAATGATTGAACTTTCATTAATTGACTTCTATTTGTGCTTTTTA
GCCTTTCTGCTATTCCTTGTTTTAATTATGCTTATTATCTTTTGGTTCTCACTTGAACTGCAAGATCATAATGAAAC
TTGTCACGCCTAAACGAACATGAAATTTCTTGTTTTCTTAGGAATCATCACAACTGTAGCTGCATTTCACCAAGAAT
GTAGTTTACAGTCATGTACTCAACATCAACCATATGTAGTTGATGACCCGTGTCCTATTCACTTCTATTCTAAATGG
TATATTAGAGTAGGAGCTAGAAAATCAGCACCTTTAATTGAATTGTGCGTGGATGAGGCTGGTTCTAAATCACCCAT
TCAGTACATCGATATCGGTAATTATACAGTTTCCTGTTTACCTTTTACAATTAATTGCCAGGAACCTAAATTGGGTA
GTCTTGTAGTGCGTTGTTCGTTCTATGAAGACTTTTTAGAGTATCATGACGTTCGTGTTGTTTTAGATTTCATCTAA
ACGAACAAACTAAAATGTCTGATAATGGACCCCAAAATCAGCGAAATGCACCCCGCATTACGTTTGGTGGACCCTCA
GATTCAACTGGCAGTAACCAGAATGGAGAACGCAGTGGGGCGCGATCAAAACAACGTCGGCCCCAAGGTTTACCCAA
TAATACTGCGTCTTGGTTCACCGCTCTCACTCAACATGGCAAGGAAGACCTTAAATTCCCTCGAGGACAAGGCGTTC
CAATTAACACCAATAGCAGTCCAGATGACCAAATTGGCTACTACCGAAGAGCTACCAGACGAATTCGTGGTGGTGAC
GGTAAAATGAAAGATCTCAGTCCAAGATGGTATTTCTACTACCTAGGAACTGGGCCAGAAGCTGGACTTCCCTATGG
TGCTAACAAAGACGGCATCATATGGGTTGCAACTGAGGGAGCCTTGAATACACCAAAAGATCACATTGGCACCCGCA
ATCCTGCTAACAATGCTGCAATCGTGCTACAACTTCCTCAAGGAACAACATTGCCAAAAGGCTTCTACGCAGAAGGG
AGCAGAGGCGGCAGTCAAGCCTCTTCTCGTTCCTCATCACGTAGTCGCAACAGTTCAAGAAATTCAACTCCAGGCAG
CAGTAGGGGAACTTCTCCTGCTAGAATGGCTGGCAATGGCGGTGATGCTGCTCTTGCTTTGCTGCTGCTTGACAGAT
TGAACCAGCTTGAGAGCAAAATGTCTGGTAAAGGCCAACAACAACAAGGCCAAACTGTCACTAAGAAATCTGCTGCT
GAGGCTTCTAAGAAGCCTCGGCAAAAACGTACTGCCACTAAAGCATACAATGTAACACAAGCTTTCGGCAGACGTGG
TCCAGAACAAACCCAAGGAAATTTTGGGGACCAGGAACTAATCAGACAAGGAACTGATTACAAACATTGGCCGCAAA
TTGCACAATTTGCCCCCAGCGCTTCAGCGTTCTTCGGAATGTCGCGCATTGGCATGGAAGTCACACCTTCGGGAACG
TGGTTGACCTACACAGGTGCCATCAAATTGGATGACAAAGATCCAAATTTCAAAGATCAAGTCATTTTGCTGAATAA
GCATATTGACGCATACAAAACATTCCCACCAACAGAGCCTAAAAAGGACAAAAAGAAGAAGGCTGATGAAACTCAAG
CCTTACCGCAGAGACAGAAGAAACAGCAAACTGTGACTCTTCTTCCTGCTGCAGATTTGGATGATTTCTCCAAACAA
TTGCAACAATCCATGAGCAGTGCTGACTCAACTCAGGCCTAAACTCATGCAGACCACACAAGGCAGATGGGCTATAT
AAACGTTTTCGCTTTTCCGTTTACGATATATAGTCTACTCTTGTGCAGAATGAATTCTCGTAACTACATAGCACAAG
TAGATGTAGTTAACTTTAATCTCACATAGCAATCTTTAATCAGTGTGTAACATTAGGGAGGACTTGAAAGAGCCACC
ACATTTTCACCGAGGCCACGCGGAGTACGATCGAGTGTACAGTGAACAATGCTAGGGAGAGCTGCCTATATGGAAGA
GCCCTAATGTGTAAAATTAATTTTAGTAGTGCTATCCCCATGTGATTTTAATAGCTTCTTAGGAGAATGACAAAAAA
AAAAAAAAAAAAAAAAAAAAAAAAAAA

**>VT1**

ATCAAGGGCCTGTACCTGCCCCGCTAACAGACCAACCAGCTGAGCATCAGCTGCCGCAGCGTGCTGTAAACCAACT
TCAAGATCTGCGTGGCCGTGACCCGCCTGCACGCCTAATGCACCCACGCCGTGTAACTGATCACCAACTACTGCCG
CTAACAGGACACCAGCAACAGCAGCATCTTCTGCCGCCTGCTGACCGTGAGCAGCGTGCTGCAGCCCATCATCAGC
ACCAGCCGCTTCCGCCCCGGCGTGACCGAGCGCTAAGACGGCGAGCCCTGCCCCTGGTTCCAGCGCGAGAACACCC
GCCCCACCCAGTTCGCCTGCTTCACCGGCAGCCGCCGCGCCCGCACCTGGCTGTGGCGCCTGCGCGGCGGCGGCCT
GATCCGCGGCACCAGCACCAGCTAACGCTGGCACCTGTGGCTGAGCCGCAGCTAAAAGCGCCGCTTCGCCAGCACC
TAAACCGCCCTGTGCGTGCACCAGACCTTCGGCTGCAGCAACTGCACCAGCTGGAGCTGCTACGGCTAAGCCGGCA
GCCGCACCCGCCGCCACAGCGTGCGCAGCTAATGGTAAGACACCTGGTGCCCCTGCCCCAGCTGCGGCCGCAACAC
CAGCGGCCTGCCCCAGGGCAGCAGCAGCTAAGAGCGCTAATAACGCAGCTGGTGGCCCTAACTGCGCCGCCGCAGC
AAGGTGATCTAACTGCGCCGCCGCGCCTGGCACTAAAGCCTGTAACGCTTCAGCCGCAAGCTGGAGCACTAAACCT
AACAGTGGTGCTACCCCTAAACCCACGCCTAAGCCTAACGCCGCGGCATCCACAGCCTGTGCCGCTAACAGCTGCT
GTGGCCCTAATGGCTGCCCAGCTAAGTGCACTAACGCCCCAGCAGCACCTGCTGGTAAAGCTTCATGCACTTCGTG
CGCACCACCGGCCTGTACTAACACTAAGAGGGCTGCATCCTGCTGCCCTAAACCTAAGCCTAAAACTGCCTGGTGC
ACGGCACCTTCTAAAAGGAGCTGTAAATCGCCGACACCTTCTAAAACTAAATCGGCAAGGAGATCTAACACCTGCA
GTGGGGCATGAGCAAGTTCTGCATCAGCCTGAAGTTCCACAACCAGGACTACAGCACCAAGGGCTAAAAGGAGAAG
GCCTAATGGCTGTACGGCTAAAACAGCATCTGCCTGAGCAGCTGCGTGACCAAGTAAATGCAGCCCAACGTGCCCT
TCAACAGCCACGAGGTGTAAAGCCTGTGGTAAAACTTCATGGCCGACGGCCGCTTCTGCTAAAGCCACCTGCGCAT
CCTGTGGCACTAAGAGTTCGACTAACGCCGCTGCCACTACCTGTGGCTGCTGACCCCCAAGTGCTGCTGCTAAAAC
CTGCTGAGCAGCATGAGCCAGTTCCGCAGCCGCACCTAAGCCTAAAGCTGCCGCATCCCCTAATAAATCTGGCTGG
AGAACCACAGCAGCTAAGGCTGGAGCCACTACTGCCTGTGGCGCCTGTGCGTGCTGCTGTGCTGGCTGCCCTAACA
GGTGTGCCTGCTGGGCAGCACCTGCTAACGCTAACACCGCCTGTAACCCTACCGCTGCTGCTGGCGCCGCTTCCGC
CGCAGCTAATAACAGCCCAGCTAAAACACCCCCAAGCGCGAGAGCCAGCACCAGTACTGCTGGTAACTGTAAACCT
AATAACGCGACCGCCACTACTTCGGCATCTTCTTCTGCTTCCACAAGTGCTTCTGCGGCAACTGCGAGCGCTTCGG
CCTGTAAAGCATCCAGACCAACTGCTAAATCCTGTGGTAATTCTAAAGCTACAAGCGCAAGAGCTAAAAGCGCTGC
CTGGAGTACTGGTAAACCGAGATCAACACCGAGAGCAGCCTGTGCATCTGCATCCGCGGCTGCAGCTGCTGCACCA
TCAACTTCCTGCCCCACAGCTAAAACTGCAGCAAGTTCTGCGCCTGCTTCACCGAGGGCCGCTACAACAACACCCG
CTGGAACTTCACCGTGTTCACCGAGACCCACTAATGCTACGACGTGCACATCTAATTCGGCTACTAACAGAGCAGC
TGCAACGGCCTGCACTACCGCTGGTGCTGCAGCGTGGACTTCGCCGTGGCCAACTAACACCTGTGGCACTGCCTGT
AAAAGACCCAGACCCGCCCCTAACTGGCCTAACGCGAGGTGTAAGGCCGCTGCCGCGTGAGCTAACGCCGCCTGGG
CAACTGCTAAATCTACCTGAACCTGTGCCTGTAAAACTGCCGCTGGACCAACTGCCACCTGTGCAAGGGCAACTAA
GGCGAGTGCAGCGACATCCTGTAAGCCTGCAAGTAAATCTTCGGCTTCGTGTGCTAACTGTACCACTACTGGTGGA
GCTAAACCTAAAGCCTGGAGTTCCGCTAAAACATCTGCCACGCCCTGAAGGGCATCGTGCAGAAGGTGTGCTAAAT
CCAGCGCCGCAACTGGCCCACCCACGCCAGCAAGAGCCCCAAGCGCAACTACCTGCTGCGCGGCCGCAACACCAGC
CACCGCAGCGTGAACCGCGGCAGCTGCCTGGAGAACTGGTAATTCACCACCATCCGCACCACCTACTAATAAAGCT
GCTAAAGCAGCATCGGCTGGTACACCAGCCTGTACTAACGCGCCTACGTGGCCCGCAACCAGCGCCACCGCAAGGT
GCTGTGCCCCTGCACCTAATACGACGGCAACAAGCAGTACCTGCACACCCAGCGCCGCTGCACCAACAAGGGCTAC
TTCTGGTAATAACACTGCGACCGCAGCGCCCGCCTGCAGGAGTGCGAGTACCACTTCTAAACCTAATAAAGGACT
AATAAAGCACCTAATAAGAGGTGCTGTGCCTGTACAGCTAAACCCGCTACCGCAGCAAGTAAGTGCGCCTGTGCTG
CGGCCGCTGCTGCCACAAGAACTTCGCCACCAGCATCTAAATCACCTACACCACCGGCCACTAATTCCGCTAAGTG
GAGTACGGCTACATCCTGCTGATCTAATAAGTGTGGTAAGTGTAAATCGGCTTCACCTACGTGCTGTTCTTCCTGC
CCAGCCGCTAAGGCTAACGCCGCCGCTAACTGTAACGCCGCCGCGTGTAAGCCATCAACAGCATCTAAGTGTGGTA
CTAACGCTAACTGCCCCGCTAAACCTTCGGCATCTGGTGCCACTTCTGCTGCAGCAGCACCTAACGCCGCGCCCGC
CGCCGCCTGGTGCGCTAATAATAAAGCACCAACTGCTGGAGCACCCGCCGCCAGTAAGGCCAGAGCGACAACTACT
ACAGCAACAACTGCTAAGGCAGCACCAGCATCCGCGACGGCACCTACACCAGCTGCAGCGACTACTAAAGCGAGTA
ATTCTAATGGCTGTTCAAGACCTACTAACAGTGCATCCACTAAAAGTGCCGCCACTGCGGCCGCAGCTAAAAGGGC
AAGACCAACAGCGGCTGCTAATGCAGCCAGTGCCTGCCCTAAACCTGGCGCCGCTGCTGCCGCAGCCTGAAGTAAG
GCTACTAACAGTGCCACGCCAGCTAAATCTAATAACTGCACAGCTACTAATGGACCACCTAAAGCGGCTGGTAACT
GTGCTTCAAGCGCACCCAGAGCTGCTAAACCCTGAGCAGCTGCTGCCGCCCCAAGTGCTAACAGCGCTAACGCCAC
AGCACCAGCTAAGAGTGCCTGTAAAAGTTCTAAAGCGCCCGCAGCAGCACCTGCACCATCATCATCAGCTGGTACT
TCTGGTGCTAACCCTACACCTTCTTCAAGAGCCTGTGCCGCTACTGCAGCCACAAGTGCCTGCTGAGCTGCCTGTA
ATAAAAGAGCCTGTAACAGACCTGCTTCAAGCTGTTCGGCAACGAGGAGTAAAAGGCCAGCTAAACCAAGGACCGC
TAAGACAGCTAACGCGGCAGCTAAGCCATCTACAACTAAAAGTAAACCTTCAGCTAAACCGAGAAGACCCGCTAAT
AAGAGAACCAGAGCCTGTGCTAACGCAGCTACAACAACAGCGGCCGCAACTAAGTGCCCCACCGCAAGCTGGTGAC
CCTGTACTAACACTAATGGCAGAGCAGCAGCCGCTTCTGCCACAGCTGCTAATAACACTAACACCACTTCCTGAAG
GAGCGCTGCAGCATCTACAGCGGCTAATGCTGCAGCCGCGGCTGCTTCAACTGCTGCGGCTACACCTACTAAAAGG

GCTGGTGGCACTACTAAAACGCCAGCGAGAGCTTCGAGAAGAGCGCCAACCGCCAGCTGTACAACCACCTGCCCGG
CAGCGGCTTCAAGTGGCTGCACTGCCGCGGCGGCAAGGACAGCGCCTAAAAGGTGTAAAAGTGCCTGCTGCACAGC
ACCATCTACTACCTGTAATAAGAGGCCCGCAACAGCTGGAACTGCTTCCTGGAGTTCGCCCGCAACGCCTGCACCT
GCCGCCGCAACACCCAGATCAACGCCTGCCTGTGCGGCAACTAAAGCCACAGCTTCAACTACACCGCCTAAATCTA
AGGCTACTAAAACACCCGCGGCTGCGGCTAACTGTGGTGCTAAATCCTGCTGCTGCACCAGTAAAACAACTGCAGC
GTGACCTACCAGCACACCTAACGCAGCAAGTAAAACAGCTGCTACAACGCCACCTGGCTGTGCAACACCTGGCTGA
AGTTCGGCCGCAGCTGCAGCGTGTACGAGATCAGCCAGAGCGCCAGCTACAGCTTCTGCTTCTTCACCTAATGCTG
CTACAGCGTGTAATGGCTGAGCTACTTCTTCTTCTAAAACACCTAACGCACCTTCTACTAAAACCACCTGACCTGC
TGGTTCCTGTAACGCCTGGTGCTGTTCTGGACCATCTACACCACCCGCTACCGCATCAGCTAAGAGCGCTAATAAA
AGTGCATCCTGCACTAATAAAGCTACCACATCCCCCCCCGCTGGTAAAGCTACCACCTGTAACAGAGCTAAGACAC
CAGCTTCTTCGAGCGCAGCGAGGACTACTAAGGCGTGTACAACAGCCGCCAGCACTAACCCCCCCACGCCAGCTGC
GGCCACGTGAACGACATCTGGACCACCGTGTGGAGCAACCTGTTCGGCTGGAGCTAATGCTACTAAAACAAGACCA
GCTAATTCACCTAACGCTAAAACATCCTGTGCTTCACCTAATAATAACACAGCACCTGCTAAGGCTTCTAAGTGCT
GCCCCACAACTAAAGCTAATTCAGCGGCTAAGTGCACGTGAGCATCAAGAGCCACTAAAAGGTGGAGATCCCCACC
AGCTAATGGTTCAACTTCTACTAAATGGGCCGCTAACAGCTGCTGAGCTGCCACTGCATCGTGAACACCCCCACCA
ACCGCGTGGAGGTGTAAAGCACCTGCAGCACCCGCTGCCTGCTGCAGAGCAAGGGCTGGTAAAGCTGCTAACTGCT
GTGCACCTACCTGAGCCTGCTGTAATAAGACAGCCGCTAAGTGCGCTAATGCTAACGCAACAACGAGCTGCTGGTG
AGCACCTGCCAGTTCCGCTTCCTGCAGAAGAGCCTGGAGCGCGGCGTGTAAAACCTGTGGACCACCGCCGACAACC
CCTAAGGCTGCCGCAGCTGCTACGTGCACGGCCACACCTTCCTGTAAACCATCTAAGAGCGCTGCAGCGACACCCT
GTACGTGTGGTAAACCAGCTACAAGATCAGCAGCACCACCGGCGTGACCTTCTGCTACGACGTGAGCACCACCTGC
AGCGTGTAAACCTAAGCCTGGTACATCTACCTGTGCTAATAAGTGCACTGGTAACTGCCCGTGTGGAGCCTGTAAA
CCTACAACTTCTAACGCAACTTCGTGCTGCACCGCCGCTGCTTCACCTACAAGGTGCTGCGCATCCAGCGCAGCTA
CTACGGCTGCTTCCTGCAGCGCAAGCAGCTGCACAACAACCACAAGACCAGCTACCTGTAAATCGGCTGGTGCTGC
CTGTACCGCAACTAACCCTAAGTGGGCCAGCTGCTGTAAGAGCGCCAGTTCCTGTTCCACCGCGCCACCAACTAAA
GCTGCACCAAGCCCACCATCAGCAAGCGCAAGCTGCGCTAATTCTAAGTGTGCATGTAATAATACCAGATCTGCTA
ATAATTCAAGCCCGTGAACTGGCTGTAAGAGACCTGCTTCAAGCGCGCCTAAAGCTACATCTTCCCCTAACTGAAG
TGGTAATGCGGCGGCTACTAACTGTAAACCCTGCACACCCTGTTCTAAGAGCGCAGCTAAATCGTGACCTAAACCT
ACTGCCTGGCCTGCTAACAGTGCAACTAATAAAGCCACGTGTAAACCAAGTACCTGGTGTACACCCTGAGCCTGGA
GCACAAGACCAGCTAAAACATCAAGTTCGTGTAATGCACCGAGGTGCGCGGCCGCGCCGGCAACGGCTAAAGCTGC
CTGCGCCGCAGCAAGACCAGCCTGTAACGCAGCAGCGGCAAGAGCTACCACACCGAGCGCCGCAGCTAAGTGTAAT
GCGAGAACTACCGCAGCTGCCGCCGCCACTACACCTAAACCAGCAAGTAATAATTCAAGAACTACCGCCGCGGCTG
GCCCCACCGCAGCAACGGCTGCCTGTGCCGCCAGTTCTAAAGCTACTACTAAGAGACCTAATAAATCATCTAAAGC
ATCCGCTTCGAGAACCCCTGCTACAGCTGGTTCAGCTGCTGCTAATAATGCCCCCTGGGCTACTACAGCTAACTGT
GCTAAGCCTTCAGCTAACAGAGCTGCTAATACAACTACTAACACAGCTACACCGTGTTCAAGCCCTGCCTGTACTA
ACTGTACGCCCTGTTCCTGTACTTCATCGCCACCATCGTGTACTTCTACTAAAAGTACAAGTTCTAAAACTAAAGC
ATCTACGCCGACTACTACAGCAAGGAGTACTGCTAAGAGTGCCGCTAAATCCTGAGCCGCGGCTTCATCTAACTGT
TCGAGGTGACCTAATTCTTCTAAACCGACAAGTACTACAACCTGGTGTTCACCATCAAGTGCCTGCCCCGCTTCTT
CAACCTGCTGAACCGCTGCTTCCGCTGCTTCAACGTGTAATTCCGCCACGCCTTCCTGCTGTACTGGCTGCAGCGC
CGCCTGTTCGAGCTGTACTAATGCCACTACTGCAACCTGCTGTACTGGTTCTACACCCTGTAATGCCTGAGCTAAT
GGTTCCGCTTCTTCCGCCACCTGAGCTTCTTCCGCAACTACACCAACTACCACTTCATCTTCTAAATGGGCTTCAA
CTGCTTCTGGCTGAGCTGCCGCGTGGTGTTCGGCATCTACAGCTTCCACTAAGTGTTCCTGTGCACCTGGATCGGC
TGCAACCACGCCATCGTGTTCCAGCTGTTCTGCAGCACCTTCTACTAATAATTCCTGGCCTACGTGGTGAACAACT
AAAGCTGCACCAACGGCCCCGACTTCAGCTACGGCTAAAACGTGCACCTGCTGTGCATCATCCTGCTGTGCATGGA
GAAGCTGTGCGCCTGCTGCCGCCGCCTGTAATTCATCAACCTGTACGACGTGCTGCAGACCTAATAAAGCAACAAG
AGCCGCATGTACAACTACTGCTAATGGTGCTAAAAGGTGCTGCTGTGCCTGTGCTAATGGCGCTAACGCCTGCTGC
AGACCACCCAGCTGGAGCTGTGCTAACTGTAATACATCCTGTGCTGGTAATACATCTACTAATAATAAAGCTGCGA
GCGCCTGGTGACCACCGTGTAAAAGACCAACAAGAGCTACTAACCCGTGTTCCTGCACCGCTAATAATGCTACAGC
GAGGAGTGGTTCCACCCCAGCCTGCTGTAATAAAGCTGGAGCAAGGACCTGTAAAAGACCTTCAGCCTGAGCTTCT
GCTAACTGCGCCAGCCCGAGAGCTAATAACACTAACGCTTCATCGCCTACTAATGCTACAGCTTCTAATGGTAAAT
CAAGATGTAACGCATCATCTGCAAGATCAGCGTGTGCCTGCTGCAGAGCGCCTACGTGAGCACCTACACCGTGACC
CGCAGCGGCATCAGCGTGTAATGCTGGTAATAATGCGGCAGCTGCAGCTAAAACGTGTAATGCCTGCGCTAATACG
TGTTCATCAACTTCTAACGCACCAACGGCAAGACCCAGAACACCAGCTGCAACTGCCGCAGCTAAACCTGCAAGGA
GTGCGTGCTGCGCCAGTGCCTGATCTACTTCTACTTCAGCAGCAGCGCCCGCGTGTGCTAATTCCGCTGCCGCAAC
TAACGCTGCTGCTAAATGAGCTAAATCGTGACCAGCATCTAACACCGCAGCTACTGGCGCTAACTGTAATAACTGT
ACGCCCACCTGTAACAGAGCTAAAAGCACGACACCCCCTAACCCTGGTGCCTGTACTAACTGTAATGCGCCAGCTA
CTAATGCGCCGGCAGCAAGAAGAGCCAGCACTGCTTCGACATGGAGCGCTAACGCTTCCACGTGATCGTGTAAACC
ACCACCAAGACCAACACCTAATGCTGCTAAAAGGAGTAACTGACCTTCTAAGTGGACATGTGCAACTACTAAACCA
GCTGCTAATGCTGCAACAACAAGGACAGCACCTAAGGCTGGTAAAACTGCTAATAACTGGTGGAGGCCGTGAACTA

AAGCTACACCTGCGTGCCCTTCTGCTGCTGCTACTTCCTGTTCAACAACACCTGCAGCTGCCACGTGTAAACCTAC
TAACTGTTCAAGTAAAACCACCGCATCCAGGGCTACTAATGGTGGTGCCACAGCTAACACAGCATCTACCGCTACC
TGTTCTGCTAACAGACCTGCTAATTCTAACACATGGTGTAACCCGCCTGGTGGTAACTGTACTAATAACAGAGCCT
GCCCATCGACTGCTGCAGCCACAACAAGCGCAGCGGCTTCTGCCGCGCCTGGTTCGCCTGGCACGACATCACCCAC
AACTAATGGTAACTGTTCGCCTTCCTGACCTAAAGCTTCTAATGCAGCTGGTAACACCTGCTGCACACCATCAAGA
CCTACCGCGTGCACTAACTGTGCAACATCAGCCTGTGCTTCGGCTGCTAAATGTACAACTTCTAACGCTGCTTCTG
GTAAGCCAGCACCATCCTGCTGTAATACCAGTGCACCCGCCGCTTCTGCTGCCTGTAAAAGTTCACCCCCTAACAC
ACCCTGTGCGCCCACGGCTGGCTGTACTACAGCATCAGCTAACACCTGCCCTAACGCTTCTGCTAAAGCGGCAACA
ACTTCTAATTCTAAGTGCTGTAAGCCCGCCACCTGTAAAAGATCCGCAGCTGGTGCCTGTGCATCTACTAATGGTA
AATGGGCACCTAACAGTAACTGCTGCAGATCTTCACCCGCAGCTTCCTGTGGTGCCGCTGCTGCAAGTTCACCTAC
TAATACGTGTACACCACCAACAGCACCTACTGGTGCTTCGGCCACATCAGCATCTACAGCAGCTGGTGGTACTGCA
GCTACCGCAGCAACATGCCCTGCCTGCTGTTCTACGAGGTGTAAAAGAGCTTCTGGTAAATCCAGAGCTGCAGCTG
CCTGTAATACTTCACCATCCCCTACGTGATCCACTGCACCCTGTTCAACACCAGCCTGCTGATCCTGACCTGGTGC
CTGTTCTGCTACCTGCTGGTGCTGGACATCCTGAGCTACTAATAATGCTTCTTCTTCAGCACCTACAGCGTGGACG
GCTACGTGCACACCTTCAGCACCTTCCTGGACAACAACTGCCTGTACCACCTGTACTTCCACAAGGCCTTCCTGCT
GGTGCTGTAATAACTGCCCAAGGAGACCTGCAGCCTGTAATGGTGCTTCCTGTAATACTTCTAACGCAGCTGCGCC
GTGCACCTGTTCGTGAAGTAACGCAACGTGAGCAAGGTGGCCTAATAATGCGCCATCACCAGCTACGCCATCTAAT
AAATCCTGAGCAGCCTGTAATAAGTGCAGGTGTTCTAATGGAGCAACGGCTACAACTAACTGCAGCGCAGCTGCCT
GCTGAGCAGCCGCAAGGGCAGCCAGTAACTGCAGTAACTGCGCTTCTAATGCAGCCTGCCCACCACCACCAACCTG
TACCACCTGAGCTGCTTCGCCGAGTGGTTCTAAAAGAACGGCATCCCCATCTGGTAAAGCTAAGGCCTGTACGGCA
CCAGCAACCTGTGGTACAACTACACCTAACGCAGCCTGGCCTAATAACGCAGCCTGCTGAGCAAGACCTGCGACCT
GCACCTGTAACGCCACGCCTAACCCTAACTGTAACGCTTCACCCACAGCTAAGTGTAAAGCTAATTCCTGGGCACC
GGCTGGTAATGCAGCACCCAGGGCTACTGGACCTTCTACGCCAAGCTGTGCACCTAAGCCTAAGGCTAATACAGCC
AGAGCTAAGACACCTAAGTGTAAGTGTGCAGCCACAGCACCCGCACCGACTTCTTCAGCGTGAGCCTGCTGCAGTG
GTTCACCATCTGGTGCCTGCCCATGTGCTACGAGGCCCAGTTCCACTACTAAGGCTTCATCCCCTAATGGTTCATG
TGGTAATGCTGGTTCTAACACCGCCTGTAACTGTGCCTGTTCCTGCTGCACGCCCCCTACGGCATCACCAACTGGA
GCAGCTGCTGGCACCGCCTGCGCCGCTAACTGCTGTGGACCTTCTGCTAACAGGCCAACAGCACCAGCAGCTGGTA
CGGCCACAACTACTACAGCTAATGCTTCAGCCTGGTGGTGCGCTGCTGCTACAAGTGGCGCCAGGTGGTGAGCCAG
AGCATCTACCACAACAGCTAATAACTGTAACCCTGCGGCTACGAGGTGCAGCTGTAAACCAGCAACACCCGCCCCT
GCTAACACACCCGCACCAGCTTCTGCAGCAACTGGAACTGCCGCTTCCGCTACGTGTGCTTCATCAAGCGCATCAC
CGCCAAGTGGTACGAGTGGACCTACCACATCGGCTAATGCTTCATCCGCCGCTAAATCTACACCTTCTAATGCTGC
TAAACCATGCTGCGCTGCTACTTCCCCAAGTGCAGCGAGAAGAACAACCAGGGCTACACCCCCCTGGTGGTGACCC
ACAACTTCGACTTCACCTTCAGCTTCAGCCCCGAGTACAGCATGGTGTTCGTGCTGTTCTTCGTGTAAAAGTGCCT
GTTCACCTTCTGCTACGGCTACTACTGCTACGTGTGCTTCTGCAACGACGTGTGCCAGACCTAAGCCTGCATCAGC
CTGTTCGTGTTCGTGACCTTCAGCTGCCACTGCAGCCTGTTCTAATACGGCCTGTACGCCTGCTAACTGGGCGACG
CCTACTACGACATGGTGGGCTACGGCTAATACTAATTCGTGTGGTTCTAAGCCAAGCGCCTGTGCTACGTGTGCAT
CAGCTGCAGCGTGACCAACCCCTACGACAGCAAGAACTGCGTGTAATAATGGTGCTAAGAGAGCGTGGACACCTAC
GAGTGCCTGGACACCCGCCTGTAAAGCCTGCTGTGGTAATGCTTCCGCAGCAGCCACTTCCACGTGGGCAGCTACA
ACCTGTGCTACTTCTAACTGCTGCGCTGCAGCTACAACTGCCACGTGTTCGGCCAGCGCTACTGCTTCTACGTGTG
CTAAGTGCTGCCCTACTTCCTGCACAACTGGTAATACACCAGCGTGTACAACGCCAGCCTGCTGTTCCTGCGCCTG
TTCCTGTACCTGCTGCTGTGGCCCCTGCTGTTCACCCAGCCCCTGCTGTAAACCGACAGCTGGTGCCTGTAACTGC
TGAGCTTCTACACCGGCGTGTAAATCTACGAGTTCACCGGCACCACCCCCACCCAGGAGTAACACCGCTGCCTGCA
GACCCAGCACTAAATCGTGGGCTGCTGGTGGCAGACCCTGTACCAGAGCAGCCACTGCACCGTGTAAAACGTGCGC
TGCAAGGTGCACATCAGCAGCCTGACCCTGAGCTTCGCCACCACCCAGAGCCGCATCATCATCTAAATCGTGGGCA
GCATGTGCCCCGTGACCCAGTAACACAGCCTGAGCTAACGCTACTACTAAAGCCTGTAAAAGAACGGCTTCACCAC
CTTCTGCTTCGCCTTCCACGCCGGCTGCTGCCGCCACAAGCAGGCCCTGTAACGCAACGCCGGCCAGCAGGGCAAC
CTGACCAGCTACAGCCTGCGCGTGTAATTCCCCAGCATCATCTGCAGCTTCTGCTACTGCAGCCGCAGCCTGTAAG
CCGGCTGCTGCTAATGGTAATTCTAAAGCTGCAGCTAAAAGGTGGAGGAGGTGTTCGAGTGCGGCTAAATCTAAAT
CTAACCCTAATGCAGCCACGCCACCTAAGTGGGCAAGGACGGCTAAAGCAGCTACGACCCCAACGTGTAAACCGGC
TAAATCTAAGGCCAGGAGGGCAAGAGCTACTAATGCTACGCCGACAACGCCTTCCACTACGCCTAAAAGGTGGGCT
AATAATGCACCCAGCAGCACTACCAGCAGTGCAAGCGCTGGCTGTGCAGCCTGGAGCACAACACCAGCTACAACAG
CAGCCAGACCAACGGCTGCCACACCCGCCTGTAACACATCTAAAAGTACGTGTAATGGTACAACATCTACCTGTGC
ATCAGCATCGTGGGCAACCCCACCGGCTGCCGCTGCCGCTAATAAAACTGCAGCACCTAATAAAACTAATACGGCC
AGTTCACCTAATTCAGCATGGCCAGCTACTGCAACAGCTTCAAGGGCCAGTTCTGCTGCCAGATCACCGAGTAATA
AGCCTAAAGCTGCTGCACCACCACCGACGTGCTGTGCTGCCGCTACTACACCAACTGCCTGCACTAATAACAGTGC
GTGAGCCTGCTGCAGCACAACAAGGGCCGCTAAGTGTGCACCTGCACCGTGATCCGCTTCACCGGCTTCGAGATGG
GCTAAATCCCCTAAGAGTAATGGAACTGGTACTACCTGTACCGCACCGGCACCACCCTGTAAGTGTGCTACCGCCA
CACCTAACGCAGCTAAAGCGAGGTGTTCATCCTGTACTAACGCATCAAGCAGCCCAAGTAACGCTACGGCACCTGG

TAATTCAGCTGCCACAGCACCAGCACCAGCTGGTAATGCAACCGCAGCGCCTGCCAGTTCAACTGCATCATCTTCC
TGTGCTTCTGCTGCCGCTGCTGCTAAAGCCTGCAGCGCCTGAGCAGCTAATGGGGCACCACCAACCACTAACTGTG
CTAAGACGTGGTGTACACCCACTGGTACTGGAGCGGCAACAACAGCTACACCGGCAGCCAGTACGGCAGCCGCATC
CTGTGGTGGTGCATCGTGCTGAGCGTGCTGCCCCTGCCCCACCGCAGCAGCAAGAGCTAACGCATCCTGTAACTGA
AGCGCTAAGTGTGCACCAACACCTACAACCTGTGCTAATAACCCTGCGGCTTCTACACCTAAAGCACAGCCTGTA
CCGCCTGCGCTACGTGGAGCGCCTGTGGCTGTAACTGTAAAGCACCCCCCGCACCCACGCCAGCGTGAGCTAATGC
ACCATCGTGTTCAAGCGCGTGTGCGGCGTGAGCGCCGCCCGCCTGACCCCCTGCGGCACCGGCACCAGCACCGACG
TGGTGTACCGCGCCTTCGACATCTACAACGACAAGGTGGCCGGCTTCGCCAAGTTCCTGAAGACCAACTGCTGCCG
CTTCCAGGAGAAGGACGAGGACGACAACCTGATCGACAGCTACTTCGTGGTGAAGCGCCACACCTTCAGCAACTAC
CAGCACGAGGAGACCATCTACAACCTGCTGAAGGACTGCCCCGCCGTGGCCAAGCACGACTTCTTCAAGTTCCGCA
TCGACGGCGACATGGTGCCCCACATCAGCCGCCAGCGCCTGACCAAGTACACCATGGCCGACCTGGTGTACGCCCT
GCGCCACTTCGACGAGGGCAACTGCGACACCCTGAAGGAGATCCTGGTGACCTACAACTGCTGCGACGACGACTAC
TTCAACAAGAAGGACTGGTACGACTTCGTGGAGAACCCCGACATCCTGCGCGTGTACGCCAACCTGGGCGAGCGCG
TGCGCCAGGCCCTGCTGAAGACCGTGCAGTTCTGCGACGCCATGCGCAACGCCGGCATCGTGGGCGTGCTGACCCT
GGACAACCAGGACCTGAACGGCAACTGGTACGACTTCGGCGACTTCATCCAGACCACCCCCGGCAGCGGCGTGCCC
GTGGTGGACAGCTACTACAGCCTGCTGATGCCCATCCTGACCCTGACCCGCGCCCTGACCGCCGAGAGCCACGTGG
ACACCGACCTGACCAAGCCCTACATCAAGTGGGACCTGCTGAAGTACGACTTCACCGAGGAGCGCCTGAAGCTGTT
CGACCGCTACTTCAAGTACTGGGACCAGACCTACCACCCCAACTGCGTGAACTGCCTGGACGACCGCTGCATCCTG
CACTGCGCCAACTTCAACGTGCTGTTCAGCACCGTGTTCCCCCCCACCAGCTTCGGCCCCCTGGTGCGCAAGATCT
TCGTGGACGGCGTGCCCTTCGTGGTGAGCACCGGCTACCACTTCCGCGAGCTGGGCGTGGTGCACAACCAGGACGT
GAACCTGCACAGCAGCCGCCTGAGCTTCAAGGAGCTGCTGGTGTACGCCGCCGACCCCGCCATGCACGCCGCCAGC
GGCAACCTGCTGCTGGACAAGCGCACCACCTGCTTCAGCGTGGCCGCCCTGACCAACAACGTGGCCTTCCAGACCG
TGAAGCCCGGCAACTTCAACAAGGACTTCTACGACTTCGCCGTGAGCAAGGGCTTCTTCAAGGAGGGCAGCAGCGT
GGAGCTGAAGCACTTCTTCTTCGCCCAGGACGGCAACGCCGCCATCAGCGACTACGACTACTACCGCTACAACCTG
CCCACCATGTGCGACATCCGCCAGCTGCTGTTCGTGGTGGAGGTGGTGGACAAGTACTTCGACTGCTACGACGGCG
GCTGCATCAACGCCAACCAGGTGATCGTGAACAACCTGGACAAGAGCGCCGGCTTCCCCTTCAACAAGTGGGGCAA
GGCCCGCCTGTACTACGACAGCATGAGCTACGAGGACCAGGACGCCCTGTTCGCCTACACCAAGCGCAACGTGATC
CCCACCATCACCCAGATGAACCTGAAGTACGCCATCAGCGCCAAGAACCGCGCCCGCACCGTGGCCGGCGTGAGCA
TCTGCAGCACCATGACCAACCGCCAGTTCCACCAGAAGCTGCTGAAGAGCATCGCCGCCACCCGCGGCGCCACCGT
GGTGATCGGCACCAGCAAGTTCTACGGCGGCTGGCACAACATGCTGAAGACCGTGTACAGCGACGTGGAGAACCCC
CACCTGATGGGCTGGGACTACCCCAAGTGCGACCGCGCCATGCCCAACATGCTGCGCATCATGGCCAGCCTGGTGC
TGGCCCGCAAGCACACCACCTGCTGCAGCCTGAGCCACCGCTTCTACCGCCTGGCCAACGAGTGCGCCCAGGTGCT
GAGCGAGATGGTGATGTGCGGCGGCAGCCTGTACGTGAAGCCCGGCGGCACCAGCAGCGGCGACGCCACCACCGCC
TACGCCAACAGCGTGTTCAACATCTGCCAGGCCGTGACCGCCAACGTGAACGCCCTGCTGAGCACCGACGGCAACA
AGATCGCCGACAAGTACGTGCGCAACCTGCAGCACCGCCTGTACGAGTGCCTGTACCGCAACCGCGACGTGGACAC
CGACTTCGTGAACGAGTTCTACGCCTACCTGCGCAAGCACTTCAGCATGATGATCCTGAGCGACGACGCCGTGGTG
TGCTTCAACAGCACCTACGCCAGCCAGGGCCTGGTGGCCAGCATCAAGAACTTCAAGAGCGTGCTGTACTACCAGA
ACAACGTGTTCATGAGCGAGGCCAAGTGCTGGACCGAGACCGACCTGACCAAGGGCCCCCACGAGTTCTGCAGCCA
GCACACCATGCTGGTGAAGCAGGGCGACGACTACGTGTACCTGCCCTACCCCGACCCCAGCCGCATCCTGGGCGCC
GGCTGCTTCGTGGACGACATCGTGAAGACCGACGGCACCCTGATGATCGAGCGCTTCGTGAGCCTGGCCATCGACG
CCTACCCCCTGACCAAGCACCCCAACCAGGAGTACGCCGACGTGTTCCACCTGTACCTGCAGTACATCCGCAAGCT
GCACGACGAGCTGACCGGCCACATGCTGGACATGTACAGCGTGATGCTGACCAACGACAACACCAGCCGCTACTGG
GAGCCCGAGTTCTACGAGGCCATGTACACCCCCCACACCGTGCTGCAGGCCGTGGGCGCCTGCGTGCTGTGCAACA
GCCAGACCAGCCTGCGCTGCGGCGCCTGCATCCGCCGCCCCTTCCTGTGCTGCAAGTGCTGCTACGACCACGTGAT
CAGCACCAGCCACAAGCTGGTGCTGAGCGTGAACCCCTACGTGTGCAACGCCCCCGGCTGCGACGTGACCGACGTG
ACCCAGCTGTACCTGGGCGGCATGAGCTACTACTGCAAGAGCCACAAGCCCCCCATCAGCTTCCCCCTGTGCGCCA
ACGGCCAGGTGTTCGGCCTGTACAAGAACACCTGCGTGGGCAGCGACAACGTGACCGACTTCAACGCCATCGCCAC
CTGCGACTGGACCAACGCCGGCGACTACATCCTGGCCAACACCTGCACCGAGCGCCTGAAGCTGTTCGCCGCCGAG
ACCCTGAAGGCCACCGAGGAGACCTTCAAGCTGAGCTACGGCATCGCCACCGTGCGCGAGGTGCTGAGCGACCGCG
AGCTGCACCTGAGCTGGGAGGTGGGCAAGCCCCGCCCCCCCCTGAACCGCAACTACGTGTTCACCGGCTACCGCGT
GACCAAGAACAGCAAGGTGCAGATCGGCGAGTACACCTTCGAGAAGGGCGACTACGGCGACGCCGTGGTGTACCGC
GGCACCACCACCTACAAGCTGAACGTGGGCGACTACTTCGTGCTGACCAGCCACACCGTGATGCCCCTGAGCGCCC
CCACCCTGGTGCCCCAGGAGCACTACGTGCGCATCACCGGCCTGTACCCCACCCTGAACATCAGCGACGAGTTCAG
CAGCAACGTGGCCAACTACCAGAAGGTGGGCATGCAGAAGTACAGCACCCTGCAGGGCCCCCCCGGCACCGGCAAG
AGCCACTTCGCCATCGGCCTGGCCCTGTACTACCCCAGCGCCCGCATCGTGTACACCGCCTGCAGCCACGCCGCCG
TGGACGCCCTGTGCGAGAAGGCCCTGAAGTACCTGCCCATCGACAAGTGCAGCCGCATCATCCCCGCCCGCGCCCG
CGTGGAGTGCTTCGACAAGTTCAAGGTGAACAGCACCCTGGAGCAGTACGTGTTCTGCACCGTGAACGCCCTGCCC
GAGACCACCGCCGACATCGTGGTGTTCGACGAGATCAGCATGGCCACCAACTACGACCTGAGCGTGGTGAACGCCC

GCCTGCGCGCCAAGCACTACGTGTACATCGGCGACCCCGCCCAGCTGCCCGCCCCCCGCACCCTGCTGACCAAGGG
CACCCTGGAGCCCGAGTACTTCAACAGCGTGTGCCGCCTGATGAAGACCATCGGCCCCGACATGTTCCTGGGCACC
TGCCGCCGCTGCCCCGCCGAGATCGTGGACACCGTGAGCGCCCTGGTGTACGACAACAAGCTGAAGGCCCACAAGG
ACAAGAGCGCCCAGTGCTTCAAGATGTTCTACAAGGGCGTGATCACCCACGACGTGAGCAGCGCCATCAACCGCCC
CCAGATCGGCGTGGTGCGCGAGTTCCTGACCCGCAACCCCGCCTGGCGCAAGGCCGTGTTCATCAGCCCCTACAAC
AGCCAGAACGCCGTGGCCAGCAAGATCCTGGGCCTGCCCACCCAGACCGTGGACAGCAGCCAGGGCAGCGAGTACG
ACTACGTGATCTTCACCCAGACCACCGAGACCGCCCACAGCTGCAACGTGAACCGCTTCAACGTGGCCATCACCCG
CGCCAAGGTGGGCATCCTGTGCATCATGAGCGACCGCGACCTGTACGACAAGCTGCAGTTCACCAGCCTGGAGATC
CCCCGCCGCAACGTGGCCACCCTGCAGGCCGAGAACGTGACCGGCCTGTTCAAGGACTGCAGCAAGGTGATCACCG
GCCTGCACCCCACCCAGGCCCCCACCCACCTGAGCGTGGACACCAAGTTCAAGACCGAGGGCCTGTGCGTGGACAT
CCCCGGCATCCCCAAGGACATGACCTACCGCCGCCTGATCAGCATGATGGGCTTCAAGATGAACTACCAGGTGAAC
GGCTACCCCAACATGTTCATCACCCGCGAGGAGGCCATCCGCCACGTGCGCGCCTGGATCGGCTTCGACGTGGAGG
GCTGCCACGCCACCCGCGAGGCCGTGGGCACCAACCTGCCCCTGCAGCTGGGCTTCAGCACCGGCGTGAACCTGGT
GGCCGTGCCCACCGGCTACGTGGACACCCCCAACAACACCGACTTCAGCCGCGTGAGCGCCAAGCCCCCCCCCGGC
GACCAGTTCAAGCACCTGATCCCCCTGATGTACAAGGGCCTGCCCTGGAACGTGGTGCGCATCAAGATCGTGCAGA
TGCTGAGCGACACCCTGAAGAACCTGAGCGACCGCGTGGTGTTCGTGCTGTGGGCCCACGGCTTCGAGCTGACCAG
CATGAAGTACTTCGTGAAGATCGGCCCCGAGCGCACCTGCTGCCTGTGCGACCGCCGCGCCACCTGCTTCAGCACC
GCCAGCGACACCTACGCCTGCTGGCACCACAGCATCGGCTTCGACTACGTGTACAACCCCTTCATGATCGACGTGC
AGCAGTGGGGCTTCACCGGCAACCTGCAGAGCAACCACGACCTGTACTGCCAGGTGCACGGCAACGCCCACGTGGC
CAGCTGCGACGCCATCATGACCCGCTGCCTGGCCGTGCACGAGTGCTTCGTGAAGCGCGTGGACTGGACCATCGAG
TACCCCATCATCGGCGACGAGCTGAAGATCAACGCCGCCTGCCGCAAGGTGCAGCACATGGTGGTGAAGGCCGCCC
TGCTGGCCGACAAGTTCCCCGTGCTGCACGACATCGGCAACCCCAAGGCCATCAAGTGCGTGCCCCAGGCCGACGT
GGAGTGGAAGTTCTACGACGCCCAGCCCTGCAGCGACAAGGCCTACAAGATCGAGGAGCTGTTCTACAGCTACGCC
ACCCACAGCGACAAGTTCACCGACGGCGTGTGCCTGTTCTGGAACTGCAACGTGGACCGCTACCCCGCCAACAGCA
TCGTGTGCCGCTTCGACACCCGCGTGCTGAGCAACCTGAACCTGCCCGGCTGCGACGGCGGCAGCCTGTACGTGAA
CAAGCACGCCTTCCACACCCCCGCCTTCGACAAGAGCGCCTTCGTGAACCTGAAGCAGCTGCCCTTCTTCTACTAC
AGCGACAGCCCCTGCGAGAGCCACGGCAAGCAGGTGGTGAGCGACATCGACTACGTGCCCCTGAAGAGCGCCACCT
GCATCACCCGCTGCAACCTGGGCGGCGCCGTGTGCCGCCACCACGCCAACGAGTACCGCCTGTACCTGGACGCCTA
CAACATGATGATCAGCGCCGGCTTCAGCCTGTGGGTGTACAAGCAGTTCGACACCTACAACCTGTGGAACACCTTC
ACCCGCCTGCAGAGCCTGGAGAACGTGGCCTTCAACGTGGTGAACAAGGGCCACTTCGACGGCCAGCAGGGCGAGG
TGCCCGTGAGCATCATCAACAACACCGTGTACACCAAGGTGGACGGCGTGGACGTGGAGCTGTTCGAGAACAAGAC
CACCCTGCCCGTGAACGTGGCCTTCGAGCTGTGGGCCAAGCGCAACATCAAGCCCGTGCCCGAGGTGAAGATCCTG
AACAACCTGGGCGTGGACATCGCCGCCAACACCGTGATCTGGGACTACAAGCGCGACGCCCCCGCCCACATCAGCA
CCATCGGCGTGTGCAGCATGACCGACATCGCCAAGAAGCCCACCGAGACCATCTGCGCCCCCCTGACCGTGTTCTT
CGACGGCCGCGTGGACGGCCAGGTGGACCTGTTCCGCAACGCCCGCAACGGCGTGCTGATCACCGAGGGCAGCGTG
AAGGGCCTGCAGCCCAGCGTGGGCCCCAAGCAGGCCAGCCTGAACGGCGTGACCCTGATCGGCGAGGCCGTGAAGA
CCCAGTTCAACTACTACAAGAAGGTGGACGGCGTGGTGCAGCAGCTGCCCGAGACCTACTTCACCCAGAGCCGCAA
CCTGCAGGAGTTCAAGCCCCGCAGCCAGATGGAGATCGACTTCCTGGAGCTGGCCATGGACGAGTTCATCGAGCGC
TACAAGCTGGAGGGCTACGCCTTCGAGCACATCGTGTACGGCGACTTCAGCCACAGCCAGCTGGGCGGCCTGCACC
TGCTGATCGGCCTGGCCAAGCGCTTCAAGGAGAGCCCCTTCGAGCTGGAGGACTTCATCCCCATGGACAGCACCGT
GAAGAACTACTTCATCACCGACGCCCAGACCGGCAGCAGCAAGTGCGTGTGCAGCGTGATCGACCTGCTGCTGGAC
GACTTCGTGGAGATCATCAAGAGCCAGGACCTGAGCGTGGTGAGCAAGGTGGTGAAGGTGACCATCGACTACACCG
AGATCAGCTTCATGCTGTGGTGCAAGGACGGCCACGTGGAGACCTTCTACCCCAAGCTGCAGAGCAGCCAGGCCTG
GCAGCCCGGCGTGGCCATGCCCAACCTGTACAAGATGCAGCGCATGCTGCTGGAGAAGTGCGACCTGCAGAACTAC
GGCGACAGCGCCACCCTGCCCAAGGGCATCATGATGAACGTGGCCAAGTACACCCAGCTGTGCCAGTACCTGAACA
CCCTGACCCTGGCCGTGCCCTACAACATGCGCGTGATCCACTTCGGCGCCGGCAGCGACAAGGGCGTGGCCCCCGG
CACCGCCGTGCTGCGCCAGTGGCTGCCCACCGGCACCCTGCTGGTGGACAGCGACCTGAACGACTTCGTGAGCGAC
GCCGACAGCACCCTGATCGGCGACTGCGCCACCGTGCACACCGCCAACAAGTGGGACCTGATCATCAGCGACATGT
ACGACCCCAAGACCAAGAACGTGACCAAGGAGAACGACAGCAAGGAGGGCTTCTTCACCTACATCTGCGGCTTCAT
CCAGCAGAAGCTGGCCCTGGGCGGCAGCGTGGCCATCAAGATCACCGAGCACAGCTGGAACGCCGACCTGTACAAG
CTGATGGGCCACTTCGCCTGGTGGACCGCCTTCGTGACCAACGTGAACGCCAGCAGCAGCGAGGCCTTCCTGATCG
GCTGCAACTACCTGGGCAAGCCCCGCGAGCAGATCGACGGCTACGTGATGCACGCCAACTACATCTTCTGGCGCAA
CACCAACCCCATCCAGCTGAGCAGCTACAGCCTGTTCGACATGAGCAAGTTCCCCCTGAAGCTGCGCGGCACCGCC
GTGATGAGCCTGAAGGAGGGCCAGATCAACGACATGATCCTGAGCCTGCTGAGCAAGGGCCGCCTGATCATCCGCG
AGAACAACCGCGTGGTGATCAGCAGCGACGTGCTGGTGAACAACTAAACCAACAACGTGTGCTTCAGCTGCTTCAT
CGCCACCAGCCTGTAAAGCGTGTGCTAAAGCTACAACCAGAACAGCATCACCCCCTGCATCCACTAATTCTTCCAC
ACCTGGTGCCTGCTGCCCTAACAGAGCTTCCAGATCCTGAGCTTCACCTTCAACAGCGGCCTGGTGCTGACCTTCC
TGTTCCAGTGCTACCTGGTGCCCTGCTACACCTGCCTGTGGGACCAGTGGTACTAAGAGGTGTAATAACCCTGCCC

CACCATCTAATAATGGTGCCTGTTCTGCTTCCACTAAGAGGTGTAACACAACAAGCGCCTGGACTTCTGGTACTAC
TTCCGCTTCGAGGACCCCGTGCCCACCTACTGCTAATAACGCTACTAATGCTGCTACTAAAGCCTGTAAATCAGCA
TCCTGTAATAAAGCATCTTCGGCTGCCTGCTGCCCCAGAAGCAGCAGAAGCTGGACGGCAAGTAAGTGCAGAGCCT
GTTCTAATGCGAGTAACTGCACTTCTAAATCTGCCTGAGCGCCTTCAGCTACGGCCCCTAACGCAAGACCGGCTAA
TTCCAGAAGAGCTAAGGCATCTGCGTGTAAGAGTACTAATGGCTGTTCTAAAACATCTTCTAAGCCCACGCCTACT
AATTCAGCGCCTAAAGCCCCAGCGGCTTCTTCGGCTTCCGCACCATCGGCCGCTTCGCCAACCGCTACTAACACCA
CTAAGTGAGCAACTTCACCTGCTTCACCTAAAAGCTGTTCGACAGCTGGTAATTCTTCTTCCGCCTGGACAGCTGG
TGCTGCAGCCTGCTGTGCGGCCTGAGCAGCACCTAAGACTTCAGCATCAAGATCTAATAAAAGTGGAACCACTACC
GCTGCTGCCGCCTGTGCACCTAACCCAGCCTGCGCAACAAGGTGTACGTGGAGATCCTGCACTGCCGCAAGCGCAA
CCTGAGCAACTTCTAACTGTAAAGCCCCACCAACCGCATCTACTGCTAAATCAGCTAATACTACAAGCTGGTGCCC
TTCTGGTAAAGCTTCTAACGCCACCAGATCTGCATCTGCCTGTGCCTGGAGCAGGAGGAGAACCAGCAGCTGTGCT
GCTAACTGTTCTGCCCCATCTAATTCCGCATCATCTTCCACTTCTAAGTGCTGTGGAGCGTGAGCTACTAAATCAA
GTAAAGCCTGCTGTACTAATGCCTGTGCCGCTTCATCTGCAACTAACGCTAATAAAGCCAGACCAACCGCAGCCGC
GCCAACTGGAAGGACTGCTAACTGTAACTGTAAATCACCCGCTAATTCTACCGCCTGCGCTACAGCCTGGAGTTCT
AACAGAGCTAATTCTAAGGCTGGTGGTAACTGTAACTGCCCGTGTAAATCGTGTAAGAGGTGTAAAGCCAGACCTT
CTAAGAGCGCTACTTCAACTAAAACCTGAGCGGCCGCTAACACACCCTGTAATGGTGCTAACGCTTCTAACTGCTG
CTGAGCTTCACCATCATCTGGTTCCCCACCCACTAATGGTGCTGGCTGCCCACCATCCAGAGCAGCAGCACCTTCT
TCTAAACCAGCACCTGCACCAGCAACTGCCTGTGGACCTAAAAGGTGTACTAATTCGGCTAAAAGCAGATGTGCCA
GTTCCAGCTGCAGTGGTTCAACCGCCACCGCTGCAGCTACTAAGTGTAACAGAAGGTGAGCGCCTTCCCCACCATC
TGGCAGCGCCACTGCTAACACTACTAATGCTGCCCCTAAAGCACCGACACCTAAGACAGCTAACACTACACCATGT
TCTTCTGGTGGTGCCAGTGCTACAACACCCGCAACAAGTACTTCTAACCCGGCTGCTGCAGCCTGAGCGGCTGCTA
ACTGCACCGCAGCCCCTGCTGCTACAGCTGCCGCAGCACCTACAGCTACCTGGCCTGCCTGTTCTACCGCTTCTAA
TGCTTCAGCAACACCTGCCGCCTGTTCAACCGCGGCTAAACCTGCCAGCAGCTGATCTAAGTGTAACACACCCACT
GGTGCCGCTACATGCGCTAACTGAGCGACAGCGACTAATTCAGCAGCGCCGGCACCTAATGCAGCTAAAGCATCCA
CCACTGCCTGCACTACGTGACCTGGTGCCGCAAGTTCAGCTGCCTGCTGTAATAACTGTACTGCCACACCCACAAG
TTCTACTACTAATGCTACCACCGCAACAGCACCAGCGTGTACGACCAGGACATCAGCCGCCTGTACAACGTGCACC
TGTGGTAATTCAACTAAATGCAGCAGAGCTTCGTGGCCATCTGGCAGTTCCTGTACACCATCAAGCCCTGCTTCAA
CTGGAACAGCTGCTAAACCCGCCAGAAGCACCCCCGCAGCTTCTGCACCAGCCAGACCAACCTGCAGAACACCACC
AACTAACGCTTCTGGTGGTTCTAATTCTTCACCAACATCACCCGCAGCATCAAGACCAAGCAGGAGGTGATCTACT
AACGCAGCACCTTCCAGCAGAGCGACACCTGCCGCTGCTGGCTGCACCAGACCATCTGGTAACTGCCCTGGTAATA
CTGCTGCTAACGCCCCCACCTGTGCACCAAGGTGTAACGCCCCTACTGCTTCGCCACCTTCGCCCACCGCTAAAAC
GACTGCAGCATCCACTTCTGCACCGTGAGCGGCTACAACCACTTCTGGCTGGACCTGTGGTGCCGCTGCTGCATCA
CCAACACCATCTGCTACGCCAACGGCCTGTAAGTGTAATGGTACTGGAGCTACACCGAGTGCAGCCTGTAAGAGCC
CAAGATCGACTGCCAGCCCATCTAATAATGCTACTGGCAGAACAGCCGCCTGACCTTCTTCCACAGCAAGTGCACC
TGGAAGACCAGCCGCTGCGGCCAGCCCAAGTGCACCAGCTTCAAGCACGCCTGCTAAACCACCTAACTGCAGTTCT
GGTGCAACTTCAAGTGCTTCAAGTAATACCCCTTCACCAGCTAACAGAGCTAAGGCTAAAGCGCCAACTAATAAGT
GGACCACCGCCAGACCAGCAAGTTCGCCGACATCTGCGACAGCACCATCAACTAAAGCTGCCGCAACCAGAGCTTC
TGCTAAAGCTGCTGCTACTAAAACGTGCGCGTGTGCACCTGGACCATCAAGAAGAGCTAATTCCTGTGGAAGGGCC
TGAGCAGCTACGTGCTGCCCAGCGTGAGCACCAGCTGGTGCAGCCTGCTGGCCTGCGACCTGTGCCCCTGCACCCG
CAAGGAGCTGCACAACTGCAGCTGCCACCTGAGCTAATGGAAGAGCACCCTGAGCAGCTAACGCTGCCTGTGCTTC
AAGTGGCACACCCTGGTGTGCAACACCAAGGAGTTCCTGTAAACCACCAACCACTACTACCGCCAGCACATCTGCG
TGTGGTAACTGTAATGCTGCAACCGCAACTGCCAGCAGCACAGCCTGTAAAGCTTCGCCACCTAAATCCGCCTGAT
CCAGGGCGGCGTGCGCTAAATCTTCTAAGAGAGCTACATCACCCGCTGCTAATTCCGCTAACACCTGTGGCACTAA
TGCTTCAGCTGCAAGCACAGCAAGCGCAACTAACCCCCCCAGTAAGGCTGCCAGGAGTTCAAGTAAATCAGCCACC
GCAGCCCCCGCACCTGGAAGGTGTAAGCCGTGTACAAGATGGCCATGGTGCACCTGGCCCGCTTCTACAGCTGGCT
GGACTGCCACAGCAACGGCGACAACTACGCCCTGCTGTACGACCAGCTGCTGTAACTGAGCCAGGGCCTGCTGTTC
CTGTGGATCCTGCTGCAGATCTAATAACGCCGCCTGTAAGCCAGCGCCCAGCGCAGCCAGATCACCCTGCACATCA
ACGAGCTGATGGACCTGTTCATGCGCATCTTCACCATCGGCACCGTGACCCTGAAGCAGGGCGAGATCAAGGACGC
CACCCCCAGCGACTTCGTGCGCGCCACCGCCACCATCCCCATCCAGGCCAGCCTGCCCTTCGGCTGGCTGATCGTG
GGCGTGGCCCTGCTGGCCGTGTTCCAGAGCGCCAGCAAGATCATCACCCTGAAGAAGCGCTGGCAGCTGGCCCTGA
GCAAGGGCGTGCACTTCGTGTGCAACCTGCTGCTGCTGTTCGTGACCGTGTACAGCCACCTGCTGCTGGTGGCCGC
CGGCCTGGAGGCCCCCTTCCTGTACCTGTACGCCCTGGTGTACTTCCTGCAGAGCATCAACTTCGTGCGCATCATC
ATGCGCCTGTGGCTGTGCTGGAAGTGCCGCAGCAAGAACCCCCTGCTGTACGACGCCAACTACTTCCTGTGCTGGC
ACACCAACTGCTACGACTACTGCATCCCCTACAACAGCGTGACCAGCAGCATCGTGATCACCAGCGGCGACGGCAC
CACCAGCCCCATCAGCGAGCACGACTACCAGATCGGCGGCTACACCGAGAAGTGGGAGAGCGGCGTGAAGGACTGC
GTGGTGCTGCACAGCTACTTCACCAGCGACTACTACCAGCTGTACAGCACCCAGCTGAGCACCGACACCGGCGTGG
AGCACGTGACCTTCTTCATCTACAACAAGATCGTGGACGAGCCCGAGGAGCACGTGCAGATCCACACCATCGACGG
CAGCAGCGGCGTGGTGAACCCCGTGATGGAGCCCATCTACGACGAGCCCACCACCACCACCAGCGTGCCCCTGTAA

GCCCAGGCCGACGAGTACGAGCTGATGTACAGCTTCGTGAGCGAGGAGACCGGCACCCTGATCGTGAACAGCGTGC
TGCTGTTCCTGGCCTTCGTGGTGTTCCTGCTGGTGACCCTGGCCATCCTGACCGCCCTGCGCCTGTGCGCCTACTG
CTGCAACATCGTGAACGTGAGCCTGGTGAAGCCCAGCTTCTACGTGTACAGCCGCGTGAAGAACCTGAACAGCAGC
CGCGTGCCCGACCTGCTGGTGTAAACCAACTAAATCCTGTACTAATTCTTCTGCCTGGAGCTGTAATTCTAACCCT
GGCAGATCCCCACCGTGCTGCTGCCCCTGAAGAGCCTGAAGAGCAGCCTGAACAACGGCACCTAATAATAAGTGAG
CTACAGCCTGCACGGCTTCGTGTTCTACAACCTGCCCATGCCCACCGGCATCGGCTTCTGCATCTAACTGAGCTAA
TTCAGCAGCGGCTGCTACGGCCAGTAACTGTAACTGGTGCTGTGCCTGCTGCTGTTCACCGAGTAAATCGGCAGCC
CCGTGGAGCTGCTGAGCCAGTGGCTGGTGCTGTAAGCCTAATGCGGCAGCGCCACCAGCCTGCTGCTGAGCGACTG
CCTGCGCGTGCGCGTGCCCTGCGGCCACAGCATCCAGAAGCTGACCTTCTTCAGCACCTGCCACAGCATGGCCCTG
TTCTAACCCGACCGCTTCTAAAAGGTGAACAGCTAAAGCGAGCTGTAAAGCTTCGTGGACATCTTCGTGCTGCTGG
ACACCATCTAAGACGCCGTG[redacted]

**>VT2**

ATTAAAGGACTGTACCTGCCCCGGTGACAGACCAACCAGCTGTCAATCAGCTGTCGGTCAGTGCTGTGAACAAACT
TTAAAATCTGCGTCGCCGTGACCAGGCTGCATGCCTGATGCACCCATGCCGTCTGACTCATCACCAACTACTGCAG
ATGACAGGATACCAGCAACAGCTCCATCTTTTGCCGGCTGCTGACCGTGAGCTCCGTGCTGCAGCCTATCATCTCA
ACCAGCCGGTTCAGACCTGGCGTGACTGAGAGATGAGACGGCGAGCCATGCCCATGGTTCCAGCGGGAAAACACCA

GACCAACCCAGTTCGCCTGCTTCACCGGCAGCCGCAGGGCCAGGACATGGCTGTGGAGACTGCGAGGAGGCGGACT
GATCCGCGGGACTAGTACCAGTTGACGCTGGCACCTGTGGCTGTCCCGGAGCTAAAAGAGGCGCTTCGCCTCTACA
TAAACAGCCCTGTGTGTGCACCAGACCTTCGGGTGCTCCAATTGCACTTCCTGGAGTTGCTACGGCTGAGCCGGCT
CTAGAACCCGGCGGCATAGCGTGAGGAGCTGATGGTGAGATACCTGGTGTCCCTGCCCCAGTTGCGGCAGGAATAC
CTCCGGCCTGCCCCAGGGCTCCAGCAGCTGAGAACGCTAATAACGGAGCTGGTGGCCCTGACTGCGGAGACGGTCC
AAGGTGATCTGACTGAGAAGGCGAGCCTGGCACTGAAGTCTCTGAAGATTTTCACGGAAGCTGGAGCACTGAACCT
GACAGTGGTGTTATCCCTGAACCCACGCCTAAGCCTGAAGGAGAGGAATCCATTCTCTGTGCAGATGACAGCTGCT
GTGGCCATAATGGCTGCCATCATGAGTGCACTGACGGCCCTCAAGCACCTGCTGGTAAAGTTTCATGCACTTCGTG
CGGACCACAGGCCTGTACTAGCACTAAGAGGGGTGTATTCTGCTGCCCTGAACCTGAGCCTGAAACTGTCTGGTGC
ATGGCACATTCTGAAAAGAGCTCTGAATCGCCGACACCTTTTGAAACTGAATCGGAAAGGAGATCTGACACCTGCA
GTGGGGTATGAGCAAGTTCTGTATCAGCCTGAAATTTCACAACCAGGATTACAGCACAAAGGGCTGAAAGGAGAAG
GCATGATGGCTGTACGGCTGAAACTCCATCTGTCTCAGCTCTTGCGTGACTAAATGAATGCAGCCTAACGTGCCCT
TTAACTCACATGAGGTTTGATCCCTGTGGTGAAACTTCATGGCCGACGGCCGGTTTTGCTGAAGCCACCTGCGGAT
CCTTTGGCACTGAGAGTTCGATTAGCGGCGCTGTCATTACCTGTGGCTGCTCACACCAAAGTGCTGTTGCTGAAAC
CTGCTGAGCAGCATGAGCCAGTTCCGGTCTAGGACCTGAGCCTGATCCTGCAGGATTCCCTGATGAATCTGGCTGG
AGAACCACTCTTCTTGAGGCTGGTCCCACTATTGCCTGTGGAGACTGTGCGTGCTGCTGTGTTGGCTGCCTTGACA
GGTGTGCCTGCTGGGGAGTACATGCTGAAGGTAACACAGGCTGTGACCCTATAGATGCTGTTGGCGGAGATTCAGG
CGGAGCTAGTGACAGCCTAGCTGAAACACCCCCAAGCGGGAGAGCCAGCACCAGTACTGCTGGTGACTGTGAACAT
AGTGACGGGATCGCCATTACTTCGGCATTTTTTTCTGCTTCCACAAGTGCTTCTGCGGAAACTGCGAGAGGTTTGG
CCTGTGATCCATTCAGACAAACTGCTGAATCCTGTGGTGATTTTGATCCTACAAGAGAAAGTCCTGAAAGCGGTGT
CTCGAGTATTGGTGAACAGAGATTAACACCGAGAGCTCCCTGTGCATCTGCATTCGGGGCTGCTCCTGCTGTACCA
TTAACTTTCTGCCTCACAGCTGAAACTGCAGTAAGTTTTGCGCCTGCTTTACAGAGGGCAGATATAACAATACAAG
ATGGAATTTCACAGTTTTCACCGAGACCCACTAGTGTTACGACGTGCACATCTGATTTGGATACTAGCAGTCCTCC
TGCAATGGGCTCCACTACAGGTGGTGCTGCTCAGTGGATTTCGCAGTGGCCAATTGACACCTCTGGCACTGTCTGT
GAAAGACCCAGACCAGACCCTGACTGGCCTGAAGGGAGGTGTGAGGAAGGTGTAGGGTGTCTTAAAGGCGGCTGGG
AAACTGTTAAATCTATTTGAACCTGTGTCTGTAGAATTGTCGGTGGACCAACTGCCACCTGTGTAAAGGTAATTGA
GGCGAGTGTTCCGACATCCTGTGAGCCTGCAAGTGAATCTTTGGGTTTGTGTGCTAGCTGTACCACTATTGGTGGA
GCTGAACCTGATCCCTGGAGTTCAGGTGAAATATTTGCCACGCGCTCAAGGGAATAGTGCAGAAGGTGTGCTGAAT
TCAGAGAAGAAATTGGCCTACTCACGCCAGCAAAAGCCCCAAGAGAAATTACCTGCTGCGCGGCAGAAATACCAGC
CACCGCAGCGTGAATAGAGGCTCTTGTCTGGAAAACTGGTGATTCACAACAATCAGAACCACATATTGATGATCTT
GCTAAAGCTCTATTGGCTGGTACACAAGCCTGTACTAACGCGCCTACGTGGCCAGAAACCAGAGACACCGGAAGGT
GCTGTGCCCATGCACTTGATACGACGGCAATAAACAGTACCTGCACACCCAGAGAAGATGCACCAACAAGGGGTAT
TTCTGGTGATGACATTGCGACAGATCTGCCCGCCTGCAGGAGTGTGAGTATCACTTCTGAACATGATGAAAAGACT
GATGATCTACCTAATAAGAGGTGCTGTGTCTGTATTCATGAACCAGATATAGGAGCAAATGAGTGAGACTGTGCTG
CGGAAGATGTTGCCACAAAAACTTTGCAACATCCATCTAAATTACTTATACCACCGGACACTGATTTAGATGAGTG
GAGTACGGTTACATCTTACTGATCTGATGAGTGTGGTGAGTCTGAATCGGGTTTACCTATGTGCTTTTTTTTCTGC
CTAGCAGATAAGGCTAACGGAGGAGGTGACTGTGAAGAAGACGTGTGTAAGCCATCAATAGTATCTGAGTGTGGTA
CTGACGGTGACTGCCCCGGTGAACATTTGGCATTTGGTGTCACTTCTGCTGCAGTAGCACATGACGGCGGGCTCGT
AGACGGCTGGTCAGGTAGTGATGATCTACTAACTGTTGGTCCACCCGAAGGCAGTGAGGACAGTCCGACAATTACT
ACTCTAACAACTGTTGAGGAAGCACCTCCATCCGGGACGGCACATACACAAGCTGCTCAGACTACTAAAGTGAGTG
ATTCTGATGGCTGTTCAAAACCTACTAGCAGTGCATCCATTGAAAGTGCAGGCACTGTGGCAGGTCCTAGAAGGGA
AAGACCAACAGTGGGTGTTGATGTTCCCAGTGTCTGCCCTGAACTTGGAGAAGATGCTGCCGCTCCTTAAAGTGAG
GCTACTAACAGTGTCACGCCAGCTAAATCTAGTGACTGCACAGCTACTGATGGACCACTTAAAGCGGGTGGTGACT
GTGCTTCAAAAGGACTCAGTCCTGCTGAACCCTCAGCAGCTGTTGTAGACCTAAATGCTAACAGAGATGAAGGCAC
AGCACAAGCTAAGAATGCCTGTGAAAATTCTGATCCGCAAGATCCTCCACATGCACCATCATCATCAGCTGGTACT
TCTGGTGCTAGCCTTACACTTTCTTCAAGTCGCTGTGCAGGTACTGCAGCCATAAGTGTCTGCTCAGTTGTCTGTG
ATAAAAAAGCCTGTGACAGACCTGCTTTAAGCTGTTTGGAAACGAGGAATGAAAGGCCAGCTGAACTAAGGACAGG
TGAGATAGCTGAAGGGGCTCTTGAGCCATCTACAATTGAAAATGAACCTTCAGCTAAACCGAGAAGACCCGGTGAT
GAGAGAACCAGAGCCTGTGCTGAAGGAGTTATAACAACTCTGGCCGCAACTGAGTGCCACACCGCAAACTGGTGAC
CCTGTACTGACACTGATGGCAGTCTTCCTCCCGGTTTTGCCACTCCTGCTGATGACACTAGCACCACTTCCTGAAG
GAGCGCTGTAGCATCTACTCTGGCTAGTGCTGCTCCAGGGGCTGCTTCAACTGCTGCGGGTACACTTACTGAAAGG
GGTGGTGGCACTACTGAAACGCCTCTGAGAGCTTCGAAAAAAGCGCTAACCGGCAGCTGTATAATCATCTGCCCGG
AAGCGGCTTCAAATGGCTGCACTGCAGAGGAGGCAAGGACAGCGCCTGAAAGGTGTGAAAATGCCTGCTGCATTCC
ACAATTTACTACCTGTGATGAGAGGCCCGGAACAGTTGGAACTGCTTCCTGGAATTCGCCAGGAATGCCTGTACCT
GTCGGCGAAATACACAGATCAACGCCTGCCTGTGCGGGAATTGATCCCATTCCTTCAACTACACTGCCTAAATCTG
AGGATACTGAAATACACGGGGATGTGGATAACTGTGGTGTTGAATCCTGCTGCTGCACCAGTAAAACAATTGCAGC
GTGACCTACCAGCACACCTAGAGGAGCAAGTAGAACAGCTGTTACAACGCAACCTGGCTGTGTAATACCTGGCTGA
AGTTTGGCAGGTCTTGTTCTGTGTACGAGATCTCCCAGTCCGCTAGTTACAGCTTCTGCTTTTTTACCTGATGCTG

TTACAGCGTGTGATGGCTGTCCTACTTTTTTTTTGAAACACCTGAAGGACATTTTACTAAAACCACCTGACCTGC
TGGTTTCTGTGAAGACTGGTGCTGTTTTGGACCATCTACACTACCCGTTACCGGATTTCATGAGAACGGTGATGAA
AGTGCATCCTGCACTAGTAAAGCTACCACATCCCTCCTCGCTGGTGATCCTACCACCTGTGACAGAGCTGAGACAC
TAGCTTCTTTGAGAGGAGCGAGGATTATTAAGGGGTGTACAATAGCAGGCAGCACTGACCACCCCACGCCAGCTGC
GGCCATGTGAACGACATCTGGACCACCGTGTGGAGTAACCTGTTCGGCTGGTCCTGATGCTACTAGAACAAGACCA
GCTGATTCACATGAAGGTGAAACATCCTGTGCTTTACCTGATGATGACACAGCACCTGCTGAGGCTTCTGAGTGCT
GCCTCACAACTGAAGCTGATTCAGCGGCTAAGTGCATGTGAGCATTAAGTCCCACTGAAAGGTGGAGATCCCCACT
TCTTGATGGTTTAACTTCTATTGAATGGGCAGATGACAGCTGCTGTCTTGTCACTGCATCGTGAACACACCTACCA
ACCGGGTGGAGGTGTGAAGCACCTGTTCTACCCGCTGTCTGCTGCAGAGCAAGGGGTGGTAAAGTTGCTAACTGCT
GTGTACCTATCTGAGCCTGCTCTGATGAGATTCCAGATGAGTGAGATGATGCTGACGGAATAATGAGCTCCTGGTG
TCCACATGCCAGTTTAGATTCCTGCAGAAGAGCCTGGAGAGGGGCGTGTGAAATCTGTGGACTACTGCCGATAATC
CATAGGGCTGCCGCTCCTGCTACGTGCATGGGCACACCTTCCTGTAAACTATCTGAGAGCGCTGTTCAGATACCCT
GTACGTGTGGTAAACCAGCTATAAGATATCATCCACTACCGGCGTGACCTTCTGCTATGATGTGAGCACCACATGC
AGCGTTTGAACCTGAGCCTGGTATATCTACCTGTGCTGATGAGTGCACTGGTGACTGCCTGTGTGGTCCCTGTGAA
CATACAACTTCTGAAGAAATTTCGTGCTGCACCGGCGGTGTTTCACCTACAAAGTGCTGCGGATCCAGCGGTCTTA
CTACGGCTGTTTCCTGCAGCGGAAGCAGCTGCACAACAACCACAAAACCTCCTATCTGTGAATCGGCTGGTGTTGC
CTGTATCGCAACTAGCCATGAGTGGGCCAGCTGCTGTGAGAGAGACAATTTCTGTTTCACAGGGCCACCAATTGAT
CCTGCACAAAGCCTACAATCAGTAAGCGGAAGCTGCGGTAGTTTTAAGTCTGTATGTGATGATATCAGATTTGCTG
ATGATTCAAGCCAGTGAACTGGCTGTGAGAGACATGCTTCAAGCGTGCCTGAAGCTACATCTTTCCATGACTGAAA
TGGTAGTGCGGGGGCTACTGACTGTAAACCCTGCACACCCTGTTTTGAGAGAGAAGCTGAATTGTGACTTGAACAT
ATTGCCTGGCCTGCTAACAGTGCAATTAATGATCCCACGTGTGAACCAAGTACCTGGTGTATACACTGAGCCTCGA
ACACAAGACTAGCTGAAATATTAAGTTCGTGTGATGTACCGAGGTCCGGGGCAGGGCCGGCAACGGCTGATCCTGC
CTGCGGCGGTCCAAGACCTCTCTGTGAAGAAGCTCCGGCAAAAGCTACCACACAGAGCGCAGGTCATGAGTGTAAT
GTGAAAACTACAGATCCTGTCGCAGGCATTACACATGAACCAGCAAGTGATGATTCAAGAACTACAGAAGGGGCTG
GCCCCACAGAAGCAACGGCTGCCTGTGCCGGCAGTTCTAGAGCTATTACTGAGAGACCTGATGAATCATCTAATCC
ATCAGGTTCGAAAATCCCTGCTACTCTTGGTTTAGCTGCTGCTGATGATGTCCACTGGGCTACTACAGCTGACTCT
GCTGAGCCTTCAGCTGACAGTCCTGCTAATACAACTACTGACACAGCTACACCGTGTTCAAGCCTTGCCTGTACTG
ACTGTACGCCCTGTTTCTGTATTTCATCGCCACCATTGTGTATTTCTATTGAAAGTACAAATTCTGAAATTGAAGC
ATCTACGCCGATTATTACAGCAAGGAGTACTGCTGAGAGTGTAGGTAAATTCTGTCCCGGGGATTCATCTGACTGT
TTGAGGTGACTTAGTTCTTTTAAACTGATAAGTACTACAACCTGGTGTTCACCATCAAGTGCCTGCCAAGGTTCTT
CAATCTGCTGAACCGGTGCTTCAGGTGCTTCAATGTGTAATTCAGGCATGCCTTTCTGCTGTACTGGCTGCAGAGA
AGGTTGTTCGAGCTGTACTAATGCCACTATTGCAACCTGCTGTACTGGTTTTACACCCTGTGATGCCTCAGCTGAT
GGTTTCGGTTCTTTCGGCACCTGAGCTTCTTCCGCAATTATACAAATTACCATTTTATTTTCTGAATGGGCTTTAA
CTGTTTCTGGCTGAGCTGCAGAGTGGTGTTCGGGATCTACAGTTTTCATTGAGTGTTTCTGTGCACATGGATCGGC
TGTAACCATGCCATCGTGTTCCAGCTGTTTTGTAGCACCTTCTATTGATGATTCCTGGCTTATGTGGTGAACAACT
GATCCTGCACAAACGGGCCAGACTTCAGCTATGGATGAAACGTGCACCTGCTGTGTATCATTCTGCTGTGCATGGA
GAAGCTCTGTGCCTGTTGTCGGAGGCTGTAATTCATCAACCTGTATGACGTGCTGCAGACCTGATGAAGTAACAAG
AGTAGAATGTACAATTATTGCTGATGGTGCTGAAAGGTGCTGCTGTGCTTGTGCTGATGGCGGTGACGGCTGTTGC
AGACCACCCAGCTGGAGCTGTGCTGACTGTGATATATCTTGTGCTGGTAATACATCTACTGATGATGAAGCTGCGA
GCGGCTGGTGACAACCGTGTGAAAGACCAATAAGTCCTATTGACCTGTGTTTCTGCACAGATGATAGTGTTACTCT
GAAGAGTGGTTCCACCCATCTCTGCTGTAGTAATCCTGGAGCAAAGATCTGTGAAAGACCTTTTCCCTGTCTTTCT
GCTGACTGAGGCAGCCCGAGTCCTGATAACATTGACGCTTCATTGCATATTAGTGTTATTCCTTCTGATGGTGAAT
CAAGATGTGACGCATTATTTGCAAGATCTCCGTGTGCCTGCTGCAGTCCGCCTACGTGAGCACCTATACAGTGACT
AGGTCCGGCATTAGCGTGTGATGTTGGTGATGATGTGGATCCTGTAGCTGAAACGTGTGATGTCTGCGCTAATATG
TGTTCATTAATTTCTGAAGGACAAACGGCAAGACACAGAACACCTCCTGTAACTGCCGCTCCTAAACATGCAAAGA
ATGTGTGCTGAGGCAGTGCCTGATTTACTTCTATTTTAGCAGCTCTGCCCGGGTGTGTTGATTCAGGTGTAGAAAC
TAACGCTGTTGCTGAATGTCCTGAATCGTGACAAGCATCTAACATCGCTCTTACTGGAGGTGACTGTGATGACTGT
ACGCCCACCTTTGACAGAGCTGAAAACACGATACCCCATGACCCTGGTGCCTCTACTAGCTGTAATGCGCCAGCTA
TTAGTGCGCCGGAAGCAAAAAGTCTCAGCACTGTTTCGACATGGAGAGATGAAGGTTCCACGTGATTGTGTGAACA
ACCACTAAGACAAACACCTGATGCTGTTAAAAAGAATGACTGACCTTCTGAGTGGATATGTGCAACTACTGAACTT
CTTGCTGATGTTGCAATAACAAGGATAGCACATAGGGGTGGTAAAATTGCTGATGACTGGTGGAGGCCGTGAACTG
ATCCTACACATGTGTGCCATTCTGCTGCTGCTACTTCCTGTTCAACAACACTTGCTCCTGTCACGTCTGAACTTAT
TGACTGTTTAAGTGAAATCATAGAATCCAGGGCTATTGATGGTGGTGCCACTCCTGACACAGCATTTACAGGTATC
TGTTTTGTTGACAGACATGTTAATTCTGACACATGGTGTGACCTGCTTGGTGGTGACTGTACTAATGACAGAGCCT
GCCCATCGACTGCTGTAGCCACAACAAGCGCAGCGGCTTCTGCCGGGCTTGGTTTGCCTGGCACGACATTACACAC
AATTGATGGTGACTGTTCGCCTTCCTGACTTGATCTTTCTGATGCTCCTGGTAGCACCTGCTGCACACTATCAAGA
CTTACCGGGTGCACTGACTGTGCAATATCAGCCTGTGCTTTGGCTGCTGAATGTACAACTTTTGACGGTGCTTTTG
GTAGGCCAGCACAATCCTGCTGTGATACCAGTGCACCCGGCGGTTTTGTTGTCTGTGAAAGTTTACCCCCTGACAT

ACCCTGTGTGCCCACGGCTGGCTCTATTATAGCATCAGCTAACACCTGCCATGACGCTTTTGCTGAAGCGGAACA
ACTTCTAATTTTGAGTTCTGTGAGCACGGCACCTGTGAAAGATCAGATCCTGGTGCCTGTGCATTTACTGATGGTG
AATGGGAACCTGACAGTAACTGTTGCAGATCTTTACCAGAAGCTTCCTGTGGTGCAGATGCTGCAAGTTCACATAC
TGATACGTGTACACTACCAATAGCACCTACTGGTGCTTCGGCCACATCAGCATTTACTCCAGCTGGTGGTACTGCA
GCTACAGGAGTAACATGCCATGCCTGCTGTTCTACGAGGTGTAGAAGTCTTTTTGGTGAATACAGAGCTGTTCATG
TCTGTGATATTTCACAATCCCCTACGTGATCCACTGCACACTGTTCAATACAAGCCTGCTGATTCTCACTTGGTGT
CTCTTCTGCTACCTGCTTGTGCTGGACATCCTGAGCTATTAATAATGCTTTTTCTTCAGCACATACAGCGTGGACG
GGTATGTGCACACCTTCAGCACCTTCCTGGACAACAATTGTCTGTACCACCTGTACTTTCATAAAGCATTCCTGCT
GGTGCTGTGATGACTGCCTAAGGAGACCTGCTCCCTTTGATGGTGTTTCCTGTGATATTTCTGAAGATCCTGCGCC
GTGCACCTGTTCGTGAAGTGAAGGAATGTGAGTAAAGTGGCCTAATGATGCGCCATTACAAGCTACGCAATCTGAT
GAATCCTGAGCTCTCTGTGATGAGTGCAGGTGTTTTAATGGAGCAACGGGTATAATTGACTGCAGAGGTCATGCCT
GCTGTCTAGCCGGAAAGGGTCCCAGTAACTGCAGTGACTGCGCTTCTGATGCTCACTGCCTACCACCACCAACCTG
TACCATCTGTCCTGCTTTGCCGAATGGTTCTGAAAGAACGGGATTCCTATCTGGTAATCCTGAGGCCTGTACGGTA
CCTCTAATCTGTGGTATAATTACACCTAACGAAGCCTGGCCTGATGACGGAGCCTGCTGTCTAAAACCTGCGACCT
GCACCTGTAAAGGCATGCCTGACCCTGACTGTGACGCTTTACACACAGCTGAGTGTGATCCTGATTTCTGGGAACC
GGCTGGTGATGCAGTACTCAGGGCTACTGGACATTCTACGCCAAGCTCTGTACATGAGCTTAAGGATGATACTCCC
AGAGCTAAGACACCTGAGTCTGAGTGTGTAGCCACTCTACCCGGACCGATTTCTTTAGCGTGAGCCTGCTGCAGTG
GTTCACTATCTGGTGCCTGCCCATGTGTTATGAAGCTCAGTTCCATTACTGAGGGTTCATCCCTTAATGGTTCATG
TGGTGATGCTGGTTCTAACACAGACTGTGACTGTGCCTGTTTCTGCTGCACGCCCCCTACGGGATTACCAACTGGA
GCAGCTGTTGGCACCGGCTGCGCCGCTGACTGCTGTGGACTTTCTGTTGACAGGCCAACTCCACATCCTCCTGGTA
CGGCCACAACTACTACAGCTGATGCTTCTCCCTGGTGGTGCGGTGCTGCTATAAGTGGAGACAGGTGGTGAGCCAG
TCCATCTATCATAACTCCTGATGACTGTAGCCTTGCGGGTATGAGGTCCAGCTGTGAACAAGTAATACAAGACCCT
GCTGACACACCCGAACCTCCTTCTGCTCCAACTGGAACTGCAGATTCAGGTACGTTTGCTTCATCAAGAGGATCAC
CGCTAAGTGGTACGAGTGGACATACCACATTGGCTGATGCTTCATTAGAAGATGAATCTACACCTTTTGATGCTGC
TGAACCATGCTGAGGTGCTATTTCCCTAAGTGCTCCGAGAAAAACAATCAGGGCTATACCCCTCTGGTGGTGACCC
ACAACTTCGACTTCACATTCAGTTTCTCCCCTGAGTATTCAATGGTGTTCGTGCTGTTTTTCGTGTGAAAATGTCT
GTTCACCTTTTGTTACGGATACTACTGCTATGTGTGCTTCTGCAACGACGTGTGCCAGACTTGAGCCTGCATTTCA
CTTTTCGTGTTTGTGACCTTTTCTTGTCACTGCAGTCTGTTTTGATACGGCCTGTACGCCTGCTAACTCGGTGATG
CCTATTACGACATGGTGGGGTATGGATGATACTGATTTGTGTGGTTCTGAGCCAAAAGACTGTGCTACGTGTGTAT
CTCCTGCAGCGTCACTAACCCCTACGACTCCAAAAATTGCGTGTGATGATGGTGCTGAGAAAGCGTGGACACCTAC
GAGTGCCTGGACACCCGGCTGTGATCTCTCCTGTGGTGATGTTTCCGGTCTTCCCATTTTCACGTGGGCTCTTACA
ATCTGTGTTACTTCTGACTGCTGAGATGCTCCTATAATTGCCATGTGTTCGGGCAGAGGTATTGTTTTTACGTTTG
TTAAGTGCTGCCCTACTTCCTGCATAACTGGTAGTACACCAGCGTGTACAACGCCTCCCTGCTGTTCCTGCGCCTC
TTTCTGTACCTGCTGCTGTGGCCTCTGCTGTTCACCCAGCCTCTGCTGTGAACTGACAGCTGGTGCCTGTGACTGC
TCAGTTTTTACACAGGCGTGTGAATCTACGAGTTCACAGGCACTACCCCAACCCAGGAGTGACACAGATGTCTGCA
GACCCAGCATTGAATCGTGGGGTGCTGGTGGCAGACACTGTATCAGTCAAGCCATTGTACAGTGTGAAACGTGAGA
TGCAAAGTGCACATCAGCTCCCTGACACTGTCCTTTGCCACTACTCAGAGCCGGATCATCATCTGAATTGTGGGCA
GCATGTGTCCCGTGACACAGTGACACAGCCTGAGCTGACGATACTACTAATCCCTGTAGAAAAACGGGTTCACCAC
ATTCTGCTTTGCATTTCACGCTGGCTGCTGCAGACACAAGCAGGCTCTGTGAAGGAATGCCGGCCAGCAAGGAAAC
CTGACCTCCTACAGTTTGAGGGTCTGATTTCCCTCTATCATTTGTTCCTTCTGCTATTGCTCAAGGTCGCTGTAAG
CCGGCTGTTGCTAATGGTGATTTTGAAGCTGTTCCTGAAAGGTGGAGGAAGTCTTCGAGTGCGGGTAAATTTGAAT
TTGACCCTGATGCAGCCACGCCACCTGAGTGGGCAAGGATGGGTGAAGCTCCTATGATCCTAACGTGTGAACCGGC
TGAATCTGAGGACAGGAGGGCAAGAGCTATTGATGCTACGCCGATAACGCCTTCCACTACGCTTGAAAGGTGGGCT
AATAGTGCACACAGCAGCACTATCAGCAGTGCAAGAGATGGCTGTGCTCCCTGGAGCACAACACTTCATACAATAG
CTCTCAGACTAATGGTTGTCATACCCGGCTGTGACACATTTGAAAATACGTGTGATGGTACAACATCTACCTGTGC
ATCAGCATCGTGGGCAACCCAACCGGATGCCGGTGCAGGTAATGAAACTGTAGCACATAATGAAATTGATATGGCC
AGTTTACTTGATTCAGCATGGCCTCTTATTGCAATAGCTTTAAGGGGCAGTTCTGCTGCCAGATTACCGAGTGATG
AGCCTGATCATGCTGCACCACAACCGACGTGCTGTGCTGTCGGTACTACACCAACTGCTTGCATTGATAACAGTGT
GTGTCCCTGCTGCAGCATAACAAAGGCCGCTAAGTGTGCACCTGTACTGTGATCAGGTTCACCGGCTTTGAGATGG
GATGAATCCCCTGAGAGTGATGGAACTGGTACTATCTGTACAGGACCGGCACCACCCTGTGAGTGTGCTACAGGCA
CACTTGAAGGAGCTGAAGTGAGGTGTTCATCCTGTATTGAAGGATCAAGCAGCCTAAGTGAAGGTACGGACCTGG
TGATTCTCCTGCCACAGTACCAGCACCTCCTGGTGATGTAATCGGTCTGCATGTCAGTTTAACTGCATCATCTTTC
TGTGCTTTTGTTGCAGGTGCTGCTGATCTCTGCAGAGACTGAGCTCTTGATGGGGAACCACAAATCACTGACTGTG
CTGAGACGTCGTGTATACCCACTGGTACTGGTCCGGCAATAACAGCTACACCGGCTCACAGTACGGCAGTCGGATT
CTGTGGTGGTGCATCGTGCTGAGCGTGCTGCCCCTGCCACACCGGTCCAGCAAGTCCTAAAGGATCCTGTGACTGA
AAAGATGAGTGTGCACCAATACCTATAACCTGTGTTGATGACCATGTGGATTCTACACCTGAAAGCACTCTCTGTA
CAGACTGAGGTACGTGGAGCGCCTGTGGCTGTGACTGTGAAGCACTCCACGCACACACGCTAGTGTGAGCTGATGT
ACAATCGTGTTCAAAAGGGTGTGTGGCGTGAGCGCTGCCCGGCTGACACCTTGCGGAACTGGAACCAGCACTGACG

TGGTGTACCGGGCCTTCGACATCTACAACGATAAGGTGGCTGGCTTCGCAAAGTTTCTGAAGACAAACTGTTGCAG
GTTCCAGGAGAAAGACGAGGACGACAATCTGATCGACAGTTATTTCGTGGTCAAACGGCACACCTTCAGCAACTAT
CAGCACGAAGAAACCATTTACAACCTGCTGAAAGATTGCCCCGCAGTGGCCAAGCACGACTTTTTCAAGTTCCGGA
TAGACGGCGACATGGTGCCTCACATCTCCCGGCAGAGACTTACAAAGTATACAATGGCTGACCTGGTTTACGCACT
GAGGCACTTTGACGAGGGCAACTGTGATACCCTGAAGGAGATCCTGGTGACTTACAATTGCTGTGACGACGACTAC
TTTAACAAGAAAGACTGGTACGACTTTGTGGAGAACCCCGACATCCTGCGGGTCTACGCCAATCTGGGCGAAAGGG
TGCGGCAGGCCCTGCTGAAGACCGTGCAGTTCTGCGACGCCATGCGGAACGCTGGCATCGTTGGCGTGCTGACACT
GGACAACCAGGACCTGAACGGGAACTGGTACGATTTTGGCGACTTTATCCAGACCACCCCCGGCAGCGGCGTGCCC
GTGGTGGATAGCTATTACTCCCTGCTGATGCCTATCCTGACCCTGACTAGGGCTCTGACCGCTGAATCTCACGTGG
ACACCGATCTGACAAAGCCTTACATCAAATGGGATCTGCTCAAGTACGACTTCACAGAGGAAAGACTGAAACTGTT
TGACAGATATTTCAAGTATTGGGATCAGACATATCACCCAAACTGCGTGAACTGCCTGGATGATAGGTGCATTCTG
CATTGTGCCAATTTTAATGTGCTGTTCTCTACCGTGTTCCCTCCCACCAGTTTTGGCCCGCTGGTGCGGAAAATCT
TCGTGGATGGGGTGCCTTTCGTGGTGTCTACAGGGTACCACTTCAGAGAGCTGGGGGTGGTTCACAATCAGGATGT
GAATCTGCATAGCTCTAGACTGTCCTTCAAGGAGCTGCTGGTTTACGCCGCCGACCCCGCCATGCACGCTGCCTCC
GGGAACCTGCTGCTGGACAAGAGGACCACCTGCTTTAGCGTGGCCGCCCTGACCAACAACGTGGCCTTCCAGACAG
TGAAGCCTGGCAACTTTAACAAAGACTTCTACGACTTCGCTGTGAGCAAGGGCTTCTTTAAGGAAGGCTCCAGCGT
GGAGCTGAAGCACTTCTTTTTTGCTCAGGACGGCAACGCAGCCATCAGCGACTACGACTACTACAGGTACAACCTG
CCAACCATGTGCGATATCCGCCAGCTGCTGTTTGTGGTGGAAGTGGTGGACAAGTACTTCGACTGTTACGACGGGG
GCTGTATCAACGCAAACCAGGTAATCGTGAATAATCTCGATAAGTCCGCTGGCTTCCCCTTCAACAAGTGGGGAAA
GGCCCGGCTGTACTACGACTCCATGTCTTATGAGGACCAGGATGCCCTGTTCGCTTACACTAAGCGGAACGTGATT
CCCACCATTACTCAGATGAATTTGAAGTACGCCATCTCTGCCAAGAATCGGGCAAGGACAGTGGCTGGCGTGTCCA
TCTGCAGCACAATGACAAACCGGCAGTTTCACCAGAAACTGCTCAAGTCCATCGCCGCCACTAGAGGAGCGACAGT
GGTCATCGGCACCAGTAAGTTTTACGGGGGCTGGCATAACATGCTTAAGACTGTGTACTCCGACGTGGAGAACCCT
CACCTGATGGGCTGGGATTATCCCAAGTGCGATCGCGCCATGCCCAACATGCTGCGGATTATGGCCAGCCTGGTGC
TGGCCAGGAAGCACACCACATGTTGTAGCCTGTCCCACCGGTTTTACCGGCTGGCCAACGAGTGCGCCCAGGTGCT
GTCTGAAATGGTTATGTGTGGCGGAAGCCTGTATGTCAAGCCAGGCGGAACCTCCTCAGGAGATGCCACTACAGCC
TATGCCAACTCCGTGTTTAACATCTGCCAGGCCGTGACCGCCAACGTGAATGCCCTCCTGTCTACCGATGGCAACA
AGATCGCCGATAAGTACGTGAGGAACCTGCAGCACAGACTGTATGAATGTCTGTACAGAAATAGAGACGTGGATAC
CGACTTTGTGAACGAGTTTTACGCCTACCTGCGGAAACATTTCAGCATGATGATCCTGTCCGACGACGCCGTGGTG
TGCTTCAATTCCACATATGCCAGCCAGGGACTGGTGGCAAGCATCAAGAATTTCAAAAGTGTGCTGTATTACCAGA
ATAACGTGTTCATGTCCGAGGCCAAGTGCTGGACCGAGACCGATCTGACCAAGGGCCCCCACGAGTTTTGCAGCCA
GCACACAATGCTGGTGAAGCAGGGCGATGACTACGTGTACCTGCCATACCCAGATCCATCCCGGATCCTGGGCGCC
GGCTGCTTCGTGGATGATATCGTGAAGACTGATGGGACCCTGATGATTGAGAGATTTGTGTCCCTGGCCATCGACG
CCTACCCCCTGACAAAGCACCCAAACCAGGAGTATGCCGATGTGTTCCACCTGTATCTGCAGTATATCAGGAAACT
GCACGACGAGCTGACCGGCCACATGTTGGACATGTATAGCGTGATGCTGACCAACGACAATACATCAAGGTACTGG
GAGCCCGAGTTCTACGAGGCCATGTACACTCCACACACCGTGCTGCAGGCTGTGGGGGCCTGCGTGCTTTGCAATA
GCCAGACAAGCCTGAGGTGCGGAGCCTGCATCCGGCGCCCCTTTCTGTGTTGCAAATGCTGTTATGACCACGTGAT
CTCTACTAGCCACAAGCTGGTGCTGTCAGTCAACCCATATGTGTGTAACGCCCCTGGCTGTGATGTGACTGACGTG
ACTCAGCTGTATCTGGGCGGGATGAGCTACTACTGTAAGTCTCACAAACCCCCTATCAGCTTCCCCCTGTGCGCCA
ATGGCCAGGTGTTCGGCCTGTACAAGAATACCTGTGTGGGCTCAGACAATGTCACCGATTTCAATGCCATTGCCAC
CTGCGATTGGACCAACGCCGGCGACTACATCCTGGCAAACACCTGTACTGAGCGGCTGAAACTGTTCGCCGCCGAG
ACACTGAAGGCCACCGAGGAGACCTTCAAACTGTCCTACGGCATCGCCACAGTGCGGGAGGTGCTGTCCGATAGGG
AGCTGCATCTGAGCTGGGAGGTGGGAAAACCTAGACCACCCCTGAATAGAAACTACGTGTTTACAGGCTATAGGGT
GACTAAGAATAGTAAAGTGCAGATTGGTGAGTACACATTTGAGAAGGGGGACTACGGCGACGCCGTGGTGTACAGG
GGAACCACAACCTATAAGCTGAACGTTGGGGACTACTTTGTGCTGACCTCCCATACCGTGATGCCACTGTCTGCCC
CCACACTGGTGCCTCAGGAGCACTACGTGCGCATCACCGGCCTGTACCCTACCCTGAATATCAGCGACGAGTTTTC
CTCTAATGTGGCCAACTACCAGAAAGTGGGGATGCAGAAGTATTCAACACTGCAGGGCCCTCCCGGCACCGGCAAG
TCACACTTCGCTATCGGCCTGGCCCTGTATTATCCCAGCGCCAGAATCGTGTACACCGCTTGTAGCCACGCCGCAG
TGGATGCCCTCTGTGAGAAGGCCCTGAAATACCTGCCTATCGATAAGTGTTCTCGGATCATCCCAGCCAGAGCCAG
GGTGGAGTGCTTTGACAAATTTAAGGTGAACAGCACACTGGAACAGTATGTGTTTTGCACTGTGAACGCCCTGCCT
GAAACCACCGCCGACATCGTGGTGTTCGACGAGATTTCCATGGCCACCAATTACGATCTGAGCGTCGTGAATGCCA
GGCTGCGCGCTAAGCACTATGTGTACATTGGGGATCCCGCCCAGCTGCCCGCTCCTAGGACACTGCTGACCAAAGG
CACCCTGGAGCCTGAGTACTTCAACTCAGTCTGCAGGCTGATGAAGACTATTGGCCCAGATATGTTTCTGGGCACC
TGTAGACGCTGTCCAGCCGAAATCGTGGATACCGTGAGTGCCCTGGTGTATGACAATAAGCTGAAGGCCCACAAGG
ATAAGTCCGCCCAGTGTTTTAAGATGTTCTATAAGGGCGTGATTACCCACGATGTCAGCTCTGCTATCAATAGGCC
ACAGATCGGCGTGGTGAGAGAGTTCCTCACAAGGAACCCAGCCTGGAGAAAGGCCGTGTTCATTTCCCCCTATAAC
TCTCAGAATGCTGTGGCATCTAAAATCTTGGGTTTGCCCACTCAGACTGTGGACAGCTCTCAGGGGTCCGAGTACG
ACTACGTGATCTTCACCCAGACCACAGAGACTGCCCACAGTTGCAACGTGAATAGGTTTAACGTGGCCATCACACG

GGCCAAGGTGGGCATCCTGTGCATAATGAGCGACAGGGACCTCTACGACAAGCTTCAGTTTACT[redacted]TGCGCCACCGTGCACACCGCTAACAAGTGGGACCTGATCATCTCCGACATGT
ACGACCCAAAGACAAAGAACGTGACTAAGGAGAATGACTCTAAGGAGGGGTTCTTCACCTACATTTGTGGATTCAT
TCAGCAGAAGCTCGCCCTGGGCGGGAGCGTGGCTATCAAGATCACCGAGCACAGCTGGAATGCTGATCTGTATAAG
CTGATGGGCCACTTTGCCTGGTGGACCGCCTTCGTGACAAACGTGAATGCCTCTTCTTCCGAAGCTTTTCTGATCG
GCTGTAACTACCTGGGAAAACCACGAGAGCAGATCGACGGCTACGTGATGCACGCAAACTACATCTTCTGGAGGAA
TACCAACCCCATCCAGCTGTCCAGCTACAGCCTGTTTGATATGTCCAAATTCCCCCTGAAGCTGAGGGGGACAGCC
GTGATGTCTCTGAAGGAGGGGCAGATTAACGACATGATCCTGAGCCTGCTGTCCAAAGGAAGACTGATCATCAGGG
AGAACAACAGAGTTGTGATCTCCAGCGACGTGCTGGTGAACAATTGAACCAACAATGTGTGCTTCAGCTGCTTCAT
CGCCACTTCTCTGTGATCCGTGTGCTAATCCTATAATCAGAATTCTATTACTCCATGCATCCACTGATTTTTCCAC
ACCTGGTGCCTGCTGCCATAGCAGAGCTTCCAGATCCTGTCTTTCACATTCAACTCCGGCCTGGTGCTGACCTTCT
TGTTCCAGTGCTACCTGGTGCCATGCTATACTTGTCTGTGGGATCAGTGGTATTGAGAGGTTTGATGACCCTGCCC
TACCATCTGATAATGGTGTCTGTTCTGCTTCCACTAAGAGGTGTGACACAACAAGAGACTGGACTTCTGGTATTAC
TTCAGATTTGAAGATCCTGTGCCCACCTACTGCTAGTAGCGGTACTGATGTTGTTATTGATCCCTGTGAATCAGCA
TTCTGTGATGAAGTATCTTCGGCTGCCTGCTGCCTCAGAAGCAGCAGAAGCTGGACGGCAAATGAGTGCAGTCCCT
GTTTTGATGCGAATAACTGCACTTTTGAATCTGCCTGTCCGCCTTCTCTTACGGCCCCTGAAGAAAGACTGGGTGA
TTTCAGAAGTCCTGAGGCATCTGCGTGTAAGAGTATTGATGGCTGTTTTAAAATATCTTCTAAGCCCACGCCTACT
AATTTTCCGCCTGATCTCCCTCCGGGTTCTTCGGCTTTAGAACAATCGGACGCTTTGCCAACAGATACTAGCACCA
TTAGGTGAGTAACTTCACATGCTTCACCTGAAAGCTGTTTGATAGCTGGTGATTCTTTTTCCGGCTGGACTCCTGG

TGTTGCAGCCTCCTGTGCGGCCTGTCTTCCACCTGAGACTTTAGCATCAAAATCTGATGAAAATGGAACCACTATA
GATGCTGCCGCCTGTGCACCTGACCATCCCTGCGCAATAAGGTGTACGTGGAGATACTGCATTGCAGAAAAAGAAA
CCTGTCCAACTTCTGACTGTAATCCCCCACTAACCGAATCTACTGCTGAATCAGTTGATATTACAAGCTGGTGCCC
TTCTGGTGATCCTTCTAAAGACATCAGATCTGCATCTGCCTGTGCTTGGAGCAGGAGGAGAATCAGCAGCTGTGTT
GCTGACTGTTTTGTCCGATCTGATTTAGAATTATCTTTCACTTTTGAGTGCTGTGGTCTGTGAGTTACTAGATCAA
ATGATCCCTGCTGTACTGATGCCTCTGCAGATTCATCTGTAATTGAAGATGATGATCCCAGACCAACCGCTCAAGG
GCTAACTGGAAGGATTGCTAACTGTGACTGTGAATTACACGCTAGTTCTACAGGCTGAGATATTCACTGGAGTTCT
AACAGTCTTGATTCTGAGGCTGGTGGTGACTGTAGCTGCCAGTCTAAATCGTGTGAGAGGTATGAAGCCAGACTTT
TTGAGAGCGCTACTTCAATTGAAATCTGAGTGGAAGATGACACACTCTCTAATGGTGCTGACGCTTCTGACTCCTG
CTGTCCTTCACTATCATCTGGTTTCCTACCCACTGATGGTGTTGGCTGCCCACCATCCAGAGTTCCAGCACCTTTT
TTTAAACCAGCACCTGTACTAGCAACTGTCTGTGGACCTGAAAGGTGTATTGATTCGGCTGAAAACAGATGTGTCA
GTTCCAGCTGCAGTGGTTCAACAGGCACCGCTGTAGCTACTGAGTGTGACAGAAAGTGTCCGCCTTTCCCACAATC
TGGCAGCGGCACTGCTGACACTACTGATGCTGCCCTTGATCAACCGACACATGAGACAGCTAGCACTACACCATGT
TTTTCTGGTGGTGCCAGTGCTACAACACAAGGAACAAATATTTCTAACCTGGCTGCTGCAGCCTGAGCGGCTGCTG
ACTGCACAGGAGCCCTTGCTGTTACAGCTGCAGGTCCACCTACAGCTACCTGGCATGCCTGTTTTACCGGTTCTGA
TGCTTCTCCAACACCTGCAGACTGTTCAACAGGGGCTGAACCTGCCAGCAGCTGATCTGAGTGTGACACACCCACT
GGTGTCGGTACATGAGATGACTGTCTGACAGCGATTGATTTTCCAGTGCCGGCACATGATGCTCCTGATCTATCCA
CCACTGCCTGCACTACGTGACCTGGTGTCGCAAGTTCAGTTGCCTCCTGTGATAACTGTATTGTCACACTCATAAG
TTCTATTACTGATGTTATCATCGGAATAGCACCAGCGTGTACGATCAGGACATTAGCCGGCTGTATAATGTGCACC
TGTGGTAATTTAACTGAATGCAGCAGAGCTTTGTGGCCATCTGGCAGTTTCTGTACACCATTAAGCCTTGTTTCAA
CTGGAACTCATGTTAAACACGGCAGAAGCACCCCCGGTCCTTCTGCACATCTCAGACAAACCTGCAGAACACCACC
AATTGACGGTTCTGGTGGTTTTGATTTTTCACTAACATTACACGGAGCATTAAGACCAAGCAGGAGGTGATTTATT
GAAGAAGCACTTTTCAGCAGTCTGATACCTGCAGGTGCTGGCTGCATCAGACAATTTGGTAGCTGCCCTGGTGATA
CTGCTGTTGACGGCCCCACCTGTGTACTAAGGTGTGAAGGCCATACTGCTTCGCCACCTTCGCTCATAGGTGAAAT
GATTGCAGTATTCATTTCTGCACCGTGTCCGGATACAATCACTTCTGGCTGGATCTGTGGTGCAGATGTTGCATCA
CTAACACAATTTGCTACGCCAACGGGCTGTGAGTGTGATGGTACTGGAGCTACACAGAGTGCAGCCTGTGAGAGCC
AAAGATCGATTGCCAGCCTATCTGATGATGCTACTGGCAGAATTCTAGGCTGACCTTTTTTCACAGCAAGTGTACA
TGGAAGACTAGCAGGTGCGGACAGCCTAAGTGCACTAGCTTTAAGCATGCTTGCTGAACCACTTAGCTGCAGTTTT
GGTGCAACTTCAAGTGTTTCAAGTGATATCCCTTCACCAGCTGACAGAGCTGAGGGTAATCTGCCAACTGATGAGT
GGACCACCGGCAGACCTCCAAATTCGCCGATATTTGTGACAGCACCATTAACTGAAGCTGCAGGAATCAGAGCTTT
TGCTGATCCTGCTGCTACTAGAATGTGAGGGTGTGTACCTGGACTATCAAAAAAAGCTGATTCCTGTGGAAGGGCC
TGTCTAGCTATGTGCTGCCTAGCGTGAGTACAAGCTGGTGCTCCCTGCTGGCCTGTGACCTGTGCCCATGCACAAG
GAAGGAGCTGCACAACTGCTCCTGCCATCTGAGCTGATGGAAGTCCACACTGTCCTCATGACGCTGCCTGTGTTTC
AAGTGGCACACCCTCGTCTGTAATACCAAAGAATTCCTGTAAACGACAAACCACTACTACAGACAGCACATCTGCG
TGTGGTGACTGTGATGCTGCAATCGGAACTGCCAGCAGCATAGTCTGTGAAGTTTCGCCACATGAATCAGGCTGAT
CCAGGGGGGCGTCAGGTGAATCTTCTGAGAGAGCTACATAACCCGGTGTTGATTCAGATAGCACCTGTGGCACTGA
TGCTTCTCTTGTAAGCACTCTAAGCGGAACTGACCCCCCCAGTAGGGGTGTCAGGAGTTCAAATGAATCTCCCACA
GATCTCCCAGGACCTGGAAAGTGTAGGCCGTGTATAAGATGGCCATGGTGCATCTGGCCAGGTTTTACTCCTGGCT
GGACTGCCACTCCAATGGAGACAATTACGCCCTGCTGTACGACCAGCTTCTGTGACTGAGCCAGGGGCTGCTGTTC
CTGTGGATTCTGCTGCAGATCTGATGACGGAGGCTGTGAGCCAGCGCCCAGCGGTCCCAGATCACCCTCCACATCA
ACGAGCTGATGGACCTTTTCATGCGCATCTTTACTATCGGCACCGTGACTCTGAAGCAGGGCGAGATTAAGGACGC
CACTCCAAGCGACTTTGTGAGAGCAACCGCCACCATCCCCATTCAGGCTTCCCTCCCATTTGGATGGCTGATCGTG
GGGGTGGCCCTGCTGGCCGTGTTTCAGTCCGCCTCTAAGATCATTACCCTGAAGAAGAGATGGCAGCTGGCTCTGA
GCAAGGGCGTGCACTTCGTCTGCAACCTGCTGCTGCTGTTCGTGACAGTGTACTCTCACCTGCTGCTGGTGGCCGC
CGGCCTGGAGGCCCCTTTCCTGTATCTGTACGCCCTCGTGTACTTCTTACAGAGCATCAATTTCGTGCGCATCATC
ATGCGGCTGTGGCTGTGTTGGAAATGCAGGTCAAAAAACCCACTGCTGTACGACGCCAACTATTTCCTGTGTTGGC
ACACCAACTGTTACGACTACTGTATCCCCTACAACTCCGTGACCTCTAGCATCGTGATCACATCCGGGGATGGCAC
CACCTCTCCCATCAGCGAACACGATTATCAGATCGGTGGCTACACCGAGAAGTGGGAGAGCGGAGTGAAGGATTGC
GTGGTGCTGCACAGCTATTTTACATCTGACTATTACCAGCTGTATTCTACCCAGCTGTCTACAGACACCGGCGTGG
AGCACGTGACCTTCTTTATCTATAACAAGATCGTGGACGAGCCTGAAGAGCACGTGCAGATTCACACCATAGACGG
CAGCTCTGGAGTGGTGAATCCAGTGATGGAGCCTATCTATGACGAGCCAACTACCACCACAAGCGTGCCCCTGTGA
GCCCAGGCCGACGAGTACGAACTGATGTATTCTTTCGTGAGCGAGGAGACCGGAACCCTGATTGTGAACAGTGTGC
TCCTGTTCCTGGCCTTCGTTGTGTTTCTGCTGGTGACCCTCGCTATCCTCACCGCCCTGCGGCTCTGCGCTTACTG
CTGCAACATCGTGAATGTGTCACTGGTGAAGCCTTCCTTCTACGTCTATAGCAGGGTCAAGAACCTGAATTCCTCC
CGGGTGCCCGACCTGCTGGTGTGAACAAACTGAATCCTGTATTGATTTTTTTGTTTGGAGCTGTGATTCTGACCTT
GGCAGATCCCAACAGTGCTGCTGCCCCTGAAGAGTCTGAAGTCCTCCCTGAATAATGGCACCTGATGATGAGTGAG
TTACTCTCTGCACGGCTTCGTGTTTTACAACCTGCCTATGCCCACTGGTATTGGCTTCTGCATTTGACTGAGCTAA
TTTAGTAGCGGGTGCTACGGGCAGTGACTGTGACTGGTGCTGTGTCTGCTCCTGTTTACAGAGTGAATCGGCTCTC

CTGTGGAGCTGCTGAGCCAGTGGCTGGTGTTGTGAGCTTGATGTGGATCCGCCACCAGCCTGCTGCTGAGCGACTG
CCTGCGGGTGCGCGTGCCTTGCGGCCACAGCATTCAGAAGCTGACCTTCTTTAGCACCTGCCACAGCATGGCCCTG
TTCTGACCTGACAGATTTTAAAAAGTGAACTCCTAGAGCGAACTGTGATCCTTCGTGGACATTTTCGTGCTGCTGG
ACACCATCTGAGATGCCGTTACCAGTCGGACATGTCTGAAGAAGTCCCTGCTGCTGCATCACGAGCGTTTCCTGAT
CACTAATTGGGAACTCCGGAGCGTGTGACAGGTGACCCAGGTGCTGCTGCACACCGTGGCCACAGGACTCGCCACC
ATCAATTAGACTCAGACCATCCCAGTGGCCGTGACAATCCTGTTGTGTCTGTATTCCAAGTGACAACAGATGTTTC
ACCTGGTGGATTTCCAGGTTACCATTGCTGAGATCCTCCTGATCATCATGAGGACCTTTAAGGTGTCTATCTGGAA
CCTGGATTACATTATCAACCTGATTATCAAGAACCTGAGCAAGAGCCTCACTGAGAACAAGTACAGCCAGCTGGAT
GAGGAGCAGCCCATGGAAATCGATTGAACCAATATGAAGATCATTCTGTTCCTGGCCCTGATCACCCTGGCCACAT
GCGAACTTTACCACTATCAGGAGTGCGTGCGGGGCACCACCGTGCTGCTGAAGGAACCCTGTAGCAGCGGCACTTA
CGAAGGCAACTCCCCATTCCACCCTCTGGCAGATAACAAGTTCGCACTCACATGTTTCAGCACACAGTTTGCTTTC
GCCTGCCCAGATGGAGTCAAGCACGTGTACCAGCTGAGAGCTCGCTCCGTGAGCCCTAAGCTCTTCATCCGGCAGG
AAGAGGTGCAGGAGCTGTACAGCCCAATCTTCCTGATTGTCGCCGCCATCGTGTTCATCACCCTGTGCTTTACACT
GAAGCGGAAAACCGAGTGACTGAACTTCCACTGACTCACTAGCATCTGCGCTTTCTAGCCATTCTGTTACAGTCTG
TTTTGACTGTGTCTGCTGTCCTTCGGATCCCATCTGAACTGTAAAATTATTATGAAACTGGTGACCCCCAAGCGGA
CTTGAAATTTCCTGTTTAGCTGAGAGTCCAGCCAGCTGTGACTGCATTTCACAAAGAATGTGGTGTACTCACACGT
GCTGAACATCAATCATATGTGATTGATGACCAGAGTGCTGTTCACCAGTATCCTGAATGGCATCCTTGAGTAAGAG
CTGGAGAACCAGCACCTGTGACTGAACTGCGCCTGGATGAGGCTGGTGCTGAACCACCCATTTAGCACCAGCATCT
CCGTGATAATTCAGTTCCCTGTGTATCTCCTGCAGCTGATTGCCCGGAATCTGAACTGGGTTGTGCTGTGATGCGT
GGTGCGGAGCATGAAAACCTTCTGAAGCATCATGACCTTCGTGCTGTTTTGAATCTCCAGCAAACGGACCAACTGA
AACGTGTAATGATGGACCCCCAAGTCAGCCAAGTGTACACCCCACTACGTTTGGTGGACTCTCCGGTTCAACTGGC
AGTGACCAGAGTGGCGGACCCAGTGGGGCGCCATCAAAACTACCAGCGCCCCCAGGTTTACCCAGTGATACTGCGT
GCTCGTGCACAGGAGCCACTCCACATGGCAGGGTAGACCTTAAATTCCATCACGCACCAGAAGGTCCAATTAGCAC
CAGTGACAGTCCAGATGACCCAACTGGCTGCTGCCTAAGAGCTATCAGACCAATTCCTGGTGGTAGAGATGAAATG
AGAGGTCCCAGTCCAAGATGGTGTTCCTGCTGCCCAGGAACTGGGCCAGATCTTGGACCTCCCTGTGGTGTTGACA
GAGAAGGCACCACATGGGATGCAATTGAGGCAGCCTGGAGTACACCAAACGCAGCCACTGGCACCCACAGTCTTGC
TGACAGTGCTGTAACAGGGCCACCACTTCTTCCCGCAACAATATCGCCAAGAGACTCCTGAGGCGGCGGGAGCAGA
GAAGGCAGTCCAGCCTGTTCTCTTTCCTGATCACCTGAAGCCAGCAGTTTAAGAAATTTAACTCCAGACAGCAGTG
AGGAAATTTCTCTTGTTGAAACGGCTGGCAGTGGAGGTGATGCTGTTCATGCTTCGCCGCCGCCTGACAGATTGAG
CCCGCTTGAGAGCAGAACGTGTGGTGAAGACCCACTACCACCAGGCCCAATTGCCACTGAGAAATCTGCTGCTGAG
GGTTTTGAGAGGCCAGTGCCAAGACCTACTGCCACTGATCTATCCAGTGTAATACCAGCTTCCGGCAGACCTGGTC
CCGCACAAACCCCAGGAAGTTCTGGGGCCCTGGGACAAACCAGACCAGGAATTGACTGCAGACCCTGGCCGCCAAC
TGTACCATCTGCCCCCAAAGATTTTCCGTCCTGCGCAACGTGGCCCACTGGCACGGCAGTCACACTTTCGGCAACG
TGGTCGACCTCCACCGGTGTCACCAGATTGGCTGACAGCGGTCTAAATTCCAGAGGAGCAGTCACTTCGCCGAATG
AGCCTACTGAAGGATCCAGAACATTCCCACAAATAGAGCCTGAAAGGGCCAGAAGGAGGAGGGATGATGAAACTCC
TCTCTGACCGCCGAGACCGAGGAGACCGCCAACTGCGACAGCTCTAGCTGTTGCCGGTTCGGGTGATTCCTGCAGA
CAATCGCTACCATCCACGAGCAGTGTTGACTGAATAGCGGCCTGAACTCCTGTCGACCTCACAAGGCTGACGGCCT
GTATAAGCGGTTCAGATTCAGCGTGTACGACATTTGAAGCACCCTGGTGCAGAACGAGTTTAGCTGACTGCACTCC
ACCTCTCGCTGTAGCTGACTCTGATCCCATATTGCCATCTTTAATCAGTGCGTCACCCTGGGCCGGACATGAAAGT
CTCATCACATCTTTACTGAGGCTACTAGATCTACCATCGAATGTACCGTGAACAATGCCAGGGAGAGTTGCCTGTA
CGGCCGGGCTCTGATGTGCAAGATCAACTTTAGCAGCGCCATCCCCATGTGATTCTGATAGCTGCTGCGGAGGATG
ACCAAGAAAAAGAAAAAGAAAAAGAAGAAAAAGAA

**>VT3**

ATTAAAGGACTCTACCTGCCAAGGTGACAGACCAACCAACTTTCAATTAGTTGTCGATCCGTTTTGTGAA
CCAACTTTAAAATCTGCGTCGCTGTCACGCGGCTGCATGCTTGATGCACCCACGCTGTATGACTCATTAC
AAACTACTGCCGATAACAGGACACGTCCAATTCCTCCATTTTCTGTAGACTTTTGACGGTCTCCAGTGTG
CTGCAACCAATTATCAGCACGAGCCGGTTTAGGCCGGGCGTAACTGAACGCTAAGATGGGGAACCCTGTC
CATGGTTTCAGCGAGAGAATACCAGACCCACGCAGTTCGCGTGTTTCACGGGGTCACGCCGGGCCAGAAC
ATGGCTGTGGAGACTTCGAGGCGGAGGCTTGATCCGGGGGACTTCAACTAGCTAAAGGTGGCACTTGTGG
CTTAGTCGGTCATGAAAGAGGAGGTTCGCGAGCACATAGACAGCTCTGTGCGTGCATCAGACATTTGGAT
GTAGCAATTGTACTAGTTGGTCTTGCTATGGTTGAGCTGGAAGCCGAACCCGCAGGCATTCTGTCAGGAG
CTGATGGTAAGACACTTGGTGCCCCTGCCCTAGTTGTGGAAGGAATACATCAGGGCTTCCACAGGGAAGC

TCAAGCTGAGAGCGGTGATGACGAAGCTGGTGGCCGTAGTTGCGCCGCCGCTCTAAAGTAATATAATTGC
GGCGAAGAGCGTGGCATTAATCACTTTAGCGCTTTAGTCGGAAGTTGGAACACTAAACGTAGCAATGGTG
TTATCCGTAGACCCACGCTTAAGCCTGACGCCGCGGGATTCACAGCCTTTGTAGGTAGCAACTCCTGTGG
CCTTAATGGTTGCCATCTTAAGTACACTAACGGCCCAGCTCCACATGCTGGTAATCTTTCATGCATTTTG
TCCGAACCACAGGACTCTATTAACACTAGGAGGGGTGTATTTTGCTTCCATAAACTTAAGCCTAGAATTG
TCTTGTGCATGGCACATTTTAGAAAGAGTTGTAAATAGCTGACACATTCTGAAATTGAATTGGTAAAGAG
ATATGACATCTTCAATGGGGAATGTCTAAATTCTGTATCTCATTGAAGTTTCATAATCAGGACTACTCAA
CAAAGGGATGAAAGGAGAAAGCTTGATGGCTTTACGGTTAGAATTCCATATGCCTGAGTAGCTGCGTCAC
AAAATAAATGCAACCAAACGTTCCTTTTAACAGCCATGAGGTGTAGTCCCTTTGGTGAAATTTCATGGCA
GACGGTCGCTTTTGTTAGTCCCATCTGAGAATATTGTGGCATTAGGAATTTGACTAAAGGCGGTGTCACT
ATTTGTGGCTTCTTACTCCCAAATGCTGCTGTTAAAATCTGTTGTCTTCCATGAGTCAATTTCGGTCAAG
AACGTGAGCATAAAGCTGTAGGATCCCTTAATAGATATGGCTGGAGAACCACAGCTCATAAGGTTGGAGC
CATTACTGTCTGTGGCGCCTTTGTGTCCTGCTGTGCTGGCTGCCATAGCAGGTATGTCTCCTGGGATCAA
CGTGCTAGCGCTAGCACCGCCTGTGACCGTATCGATGCTGCTGGAGGAGGTTCCGAAGGAGTTAATGACA
ACCCTCTTAAAACACTCCCAAACGAGAGTCCCAGCATCAGTATTGCTGGTAGCTTTGAACATAGTGAAGG
GATCGCCATTACTTCGGAATATTTTTCTGCTTCCATAAGTGTTTCTGTGGTAACTGCGAGCGATTCGGGC
TGTAAAGCATTCAAACGAACTGTTAGATACTCTGGTAGTTCTAGTCATACAAGCGGAAAAGTTAGAAGCG
ATGCCTCGAGTACTGGTAGACTGAGATTAACACAGAAAGTAGTCTCTGTATATGCATTCGAGGTTGCTCT
TGTTGCACAATTAACTTCCTCCCACACTCCTGAAACTGCTCAAAGTTTTGTGCCTGTTTCACTGAAGGTC
GCTATAACAATACGCGCTGGAACTTCACCGTATTTACCGAAACACATTAGTGCTACGATGTACATATATG
ATTCGGGTACTAGCAAAGTAGTTGTAATGGATTGCACTATCGATGGTGTTGCAGTGTGGATTTCGCCGTG
GCGAACTAACACCTGTGGCATTGCCTTTAGAAGACACAGACAAGGCCGTGACTCGCCTAACGCGAAGTCT
AGGGTCGGTGCCGCGTATCATAGCGCAGATTGGGAAACTGCTAGATTTACCTGAACCTTTGCCTTTAAAA
TTGTAGATGGACCAATTGCCATCTCTGCAAGGGTAACTAGGGCGAGTGCTCCGATATCCTCTAAGCGTGT
AAATGAATTTTTGGATTCGTGTGTTGACTCTATCATTATTGGTGGTCCTAAACATGATCTCTGGAGTTTC
GATGAAACATCTGTCACGCTCTGAAGGGTATCGTCCAAAAAGTCTGCTGAATACAGCGCAGAAATTGGCC
TACACATGCATCAAAATCTCCTAAACGGAATTATCTCCTCAGGGGGCGGAATACCAGTCACCGGTCTGTT
AACAGAGGAAGCTGTCTTGAGAATTGGTGATTTACTACGATACGCACAACTTACTAATAGTCCTGCTAGA
GTTCAATCGGGTGGTATACGTCCCTTTATTAGAGAGCATACGTCGCACGAAACCAGCGGCACCGGAAAGT
TCTCTGTCCATGCACTTGATATGACGGTAACAAACAGTACCTGCACACTCAGCGACGCTGTACCAATAAG
GGTTACTTTTGGTAGTGACATTGTGACAGATCTGCCAGGCTCCAGGAATGTGAGTACCACTTCTAGACCT
GATAAAAAGATTAGTAAAGTACGTAATAGGAAGTTCTGTGTTTGTATTCCTGAACTAGGTACCGAAGTAA
ATGAGTGAGGCTTTGTTGTGGGCGGTGCTGCCACAAAAATTTTGCGACGTCCATCTAGATTACATATACA
ACTGGACACTGATTCCGATAAGTAGAGTATGGCTACATACTCCTGATCTGATGAGTCTGG[redacted]

[redacted]TGATCTTGCACAAAACCCAC
GATCTCCAAAAGGAAGCTTAGGTAGTTCTAGGTTTGCATGTAATGATATCAAATATGCTAGTGATTCAAA
CCCGTCAACTGGCTCTAGGAAACATGTTTTAAGAGAGCCTAGTCCTATATTTTTCCCTAGTTGAAGTGGT
AGTGCGGCGGGTACTAACTCTAAACTCTTCATACGCTTTTCTAGGAAAGATCTTAAATAGTTACATAAAC
CTACTGCCTGGCGTGCTGACAATGCAATTAATGATCACACGTGTAAACGAAATACTTGGTGTATACCCTC
TCTCTTGAACACAAGACCTCTTGAAATATAAAGTTCGTGTAATGTACAGAAGTGAGGGGCAGGGCGGGCA
ATGGTTGAAGTTGCTTGCGGAGGTCCAAGACCTCACTTTAACGATCTAGCGGAAAATCTTATCACACTGA
GAGGAGATCATGAGTGTAATGTGAGAATTATCGATCCTGCAGGAGGCACTACACTTAGACAAGTAAATAG
TAGTTTAAAAACTATCGCAGGGGATGGCCCCATCGGAGTAATGGGTGCCTGTGCAGACAATTCTAATCAT
ACTATTGAGAGACTTGATGAATTATTTAATCTATACGGTTTGAGAACCCTTGCTACAGCTGGTTCTCATG
CTGCTAATAATGTCCCCTTGGTTATTACAGTTGACTCTGTTAAGCATTCTCTTAGCAGAGTTGCTGATAC
AACTATTAGCATTCATATACAGTTTTCAAGCCATGTCTGTATTAACTCTACGCGCTTTTCCTCTATTTTA
TAGCAACTATAGTTTACTTTTATTGAAAGTACAAGTTTTAAAACTGAAGTATATACGCCGACTATTATTC
AAAAGAATACTGTTGAGAATGTCGCTAGATCCTTAGCAGAGGTTTTATCTGACTGTTTGAAGTTACTTAG
TTTTTCTAAACCGACAAGTACTATAATCTGGTATTCACGATAAAGTGTCTCCCACGATTCTTTAACCTGC
TGAACAGATGCTTCAGATGCTTCAACGTCTGATTTCGCCATGCTTTCTTGCTCTACTGGCTGCAGCGCAG
ACTGTTTGAGCTGTATTAGTGCCATTACTGCAACCTGCTCTATTGGTTCTATACGCTTTAGTGTCTCTCT
TAATGGTTCAGGTTTTTCAGGCACCTGTCCTTCTTCAGAAATTACACCAACTATCACTTCATTTTCTAAA
TGGGTTTCAATTGCTTCTGGCTGTCCTGTCGGGTAGTGTTTGGGATATATAGCTTCCACTAGGTGTTTCT
CTGTACATGGATAGGTTGCAACCACGCTATTGTCTTTCAACTCTTTTGTAGTACCTTTTACTAATGATTT
TTGGCATACGTTGTGAACAATTAGTCATGCACTAACGGTCCAGACTTCAGTTACGGATAAAATGTGCATC
TGCTGTGCATTATCTTGCTCTGCATGGAGAAACTGTGTGCATGTTGCCGACGGCTTTAGTTCATAAATCT
TTACGACGTACTTCAAACCTAATAATCCAATAAATCTCGCATGTATAACTACTGTTAGTGGTGCTAAAAG
GTTCTTCTGTGCCTGTGTTAATGGAGGTGAAGGCTGCTTCAAACAACACAACTTGAGCTTTGCTAGCTCT
AATACATACTTTGCTGGTGATATATTTATTGATGATGATCATGTGAGCGCCTTGTGACTACCGTCTAGAA
AACAAATAAGTCCTACTAACCAGTGTTTCTCCATAGATGATAGTGCTATTCCGAGGAGTGGTTTCACCCT
TCACTTTTGTGATAGAGCTGGTCCAAGGATCTGTGAAAGACATTTAGTCTTTCCTTTTGTTAACTTCGGC
AGCCGGAATCCTGATAGCACTAACGATTCATTGCATACTAATGCTATAGCTTCTGATGGTAAATCAAAAT
GTAAAGGATTATCTGTAAAATAAGTGTATGCCTCCTCCAGAGTGCCTACGTTAGCACCTACACGGTAACT
CGGAGTGGGATATCCGTCTGATGTTGGTGATAGTGTGGTTCTTGCTCATGAAACGTTTAGTGTCTGCGCT

AATACGTCTTTATAAATTTCTAAAGAACTAATGGTAAGACACAGAACACATCTTGCAATTGCAGGTCCTG
AACGTGCAAGGAGTGTGTGCTGCGCCAGTGCCTGATATACTTTTACTTCAGTAGTAGTGCGAGAGTTTGC
TAATTTAGATGCAGAAATTAACGCTGCTGTTGAATGTCTTGAATAGTTACCTCAATCTAGCACAGATCCT
ACTGGAGATAGTTGTGATAACTCTATGCTCACCTTTGACAATCATAGAAGCATGACACCCCGTAGCCGTG
GTGCCTTTACTAGCTCTGATGTGCGTCCTATTAGTGCGCCGGGAGTAAAAAGTCACAGCACTGCTTCGAT
ATGGAGCGCTAGCGATTTCACGTTATTGTGTAAACGACGACTAAAACGAATACTTAATGTTGTTGAAAGG
AATAATTGACTTTTTGAGTTGACATGTGTAACTATTAAACTAGCTGTTAGTGCTGTAACAACAAAGATAG
CACGTAGGGGTGGTAGAATTGTTGATAACTTGTAGAGGCCGTCAATTAAAGTTATACGTGCGTTCCCTTC
TGCTGTTGTTACTTCCTTTTTAACAACACGTGTTCATGTCATGTGTAAACATATTGACTTTTTAAATAGA
ATCATCGCATTCAGGGGTATTAGTGGTGGTGTCACTCATGACATTCTATCTATCGCTATCTCTTTTGTTA
GCAAACGTGCTAATTCTAGCACATGGTATAGCCCGCATGGTGGTAGCTCTATTGATGACAAAGCCTCCCT
ATCGATTGTTGTTCCCACAATAAGAGATCCGGTTTTTGCCGCGCCTGGTTTGCATGGCACGATATCACGC
ATAACTGATGGTGACTCTTCGCATTTTTGACTTAGTCTTTTTAGTGTAGCTGGTAACATCTCCTTCATAC
AATAAAGACATATAGGGTACATTAACTGTGTAATATAAGCTTGTGTTTCGGATGCTAGATGTACAATTTT
TAGCGCTGCTTCTGGTGAGCCTCAACCATTCTGCTGTAGTATCAATGCACGAGACGGTTTTGTTGTTTGT
AAAAGTTCACGCCATAACATACCCTTTGTGCACACGGATGGCTTTATTATAGTATATCATGACACCTTCC
CTGACGGTTTTGCTAATCAGGCAACAACTTTTAATTCTGAGTCTTGTGAGCACGGCACCTCTGAAAAATT
CGCAGTTGGTGTCTTTGCATCTATTGATGGTAAATGGGCACTTAACAATGACTCTTGCAAATTTTTACGC
GAAGCTTCTTGTGGTGTAGATGTTGTAAATTCACATACTAATATGTCTATACCACAAACAGTACTTATTG
GTGTTTCGGTCACATATCTATTTACTCAAGCTGGTGGTATTGCAGCTATCGATCTAACATGCCCTGCCTG
CTGTTCTACGAAGTATAGAAAAGTTTCTGGTAGATCCAATCATGTTCATGCCTTTAGTATTTCACCATAC
CGTATGTAATTCATTGCACATTGTTCAACACCTCACTTTTGATACTTACCTGGTGCCTGTTCTGTTACTT
GCTGGTACTTGATATCCTGTCTTATTGATAATGCTTCTTTTTTAGTACTTATAGTGTCGATGGATACGTG
CACACTTTCAGTACATTTCTGGATAACAACTGCCTCTATCATCTCTACTTCCATAAGGCATTCCTCCTTG
TATTGTGATAACTCCCAAAGGAGACATGCAGTTTGTGATGGTGCTTCCTGTGATATTTCTAGCGGTCCTG
CGCCGTCCACCTTTTCGTGAAGTAGCGCAATGTGTCCAAGGTCGCATAGTAGTGTGCCATCACTAGTTAC
GCCATCTAATGAATTCTCTCCAGCCTCTGATAGGTCCAAGTGTTCTGATGGTCAAACGGCTACAACTAGC
TTCAAAGGAGTTGCCTTCTCAGCTCAAGGAAGGGATCACAATAGCTTCAGTAACTTCGATTTTAATGTTC
ACTTCCTACGACGACAAATCTTTATCACCTCTCATGTTTCGCTGAATGGTTTTGAAAGAACGGTATACCG
ATTTGGTAGAGTTAAGGACTGTATGGAACGTCTAACCTGTGGTATAATTACACATGAAGGTCCCTGGCCT
AATAGCGGTCACTTCTTTCCAAAACGTGCGATCTCCATTTGTAAAGACATGCGTAACCCTAGTTGTGACG
ATTCACACATAGTTAGGTATAATCTTAATTCCTCGGTACCGGGTGGTGATGCTCCACGCAGGGATATTGG
ACCTTTTACGCTAAGTTGTGTACTTAGGCATAGGGGTAATACTCCCAAAGTTAAGACACGTAAGTGTAAG
TGTGTTCTCACAGTACACGAACCGACTTCTTCTCTGTCTCTCTCCTGCAATGGTTTACTATTTGGTGCCT
TCCGATGTGCTATGAAGCTCAATTTCACTATTGAGGGTTTATCCCGTGATGGTTTATGTGGTGATGTTGG
TTCTAACACAGGCTCTAGCTCTGCTTGTTTTTGTTGCACGCCCCGTACGGCATTACGAACTGGTCCTCAT
GCTGGCATCGGCTTCGAAGATAGCTTCTGTGGACGTTCTGCTGACAAGCGAATTCAACAAGTTCCTGGTA
TGGTCATAACTACTATTCCTGATGCTTCTCACTGGTAGTTAGGTGTTGCTATAAGTGGAGGCAGGTCGTT
TCCCAATCAATATATCATAATAGTTAGTAGCTTTAACCGTGTGGTTATGAAGTACAGTTGTAGACCTCCA
ATACTCGACCCTGTTAGCATACTCGCACTTCTTTTTGCTCCAACTGGAATTGCCGCTTCAGATATGTTTG
TTTTATAAAGAGAATCACGGCGAAATGGTATGAATGGACGTATCATATCGGATGATGCTTCATTCGACGG
TAAATTTACACTTTCTAGTGTTGCTGAACGATGTTGAGATGTTACTTCCCGAAATGTTCAGAAAAGAATA
ATCAGGGGTATACGCCCTTGGTTGTCACTCACAATTTTGATTTCACTTTTTCTTTTTCACCTGAATACTC
AATGGTATTTGTCCTCTTTTTCGTATGAAAATGTTTGTTCACCTTTTGCTACGGATATTATTGTTATGTT
TGCTTTTGCAATGACGTGTGTCAGACCTGAGCGTGCATAAGCCTGTTCGTTTTCGTCACTTTCAGTTGCC
ACTGTTCTTTGTTCTGATATGGACTTTACGCCTGCTAACTCGGCGACGCGTATTACGATATGGTTGGTTA
TGGTTGATACTGATTCGTCTGGTTTTAAGCTAAAAGATTGTGCTACGTGTGCATAAGTTGTTCAGTTACT
AATCCATACGATAGTAAAAACTGTGTTTAGTAATGGTGCTGAGAGAGTGTTGATACTTATGAGTGTCTTG
ATACAAGGCTGTGAAGTTTGCTTTGGTGATGTTTCCGCTCCTCACACTTTCACGTTGGCTCCTACAATCT
GTGCTATTTTTAACTTCTGCGCTGCTCTTATAATTGCCATGTCTTTGGTCAGCGATACTGTTTTTATGTT
TGCTAGGTCTTGCCGTATTTTCTCCATAATTGGTGATACACGTCCGTGTACAACGCTTCACTGTTGTTCC
TTCGGCTGTTTCTTTATTTGCTGTTGTGGCCTCTCTTGTTCACTCAACCACTTCTGTGAACTGACTCCTG
GTGCCTTTAGCTCCTGAGCTTCTACACGGGAGTTTGAATTTACGAATTTACGGGGACTACGCCGACACAA

GAATAACACCGCTGTCTTCAAACGCAGCACTAAATTGTGGGTTGCTGGTGGCAAACGTTGTACCAATCTT
CCCACTGCACAGTGTGAAATGTCCGCTGTAAGGTGCACATATCTTCATTGACACTCTCCTTTGCCACTAC
ACAGAGTCGGATCATCATCTAGATTGTAGGTTCCATGTGCCCAGTCACGCAGTAGCATAGCCTGTCTTGA
AGGTATTACTGATCCCTTTAAAAAAACGGTTTCACCACTTTCTGTTTTGCTTTTCACGCGGGTTGTTGTC
GGCATAAGCAAGCTCTTTAACGGAACGCGGGACAACAGGGCAATCTCACGTCTTACTCTCTTCGAGTCTG
ATTTCCCAGTATAATTTGTAGTTTTTGTTATTGCAGCCGCTCCCTGTGAGCGGGGTGTTGCTGATGGTAA
TTTTAATCTTGCTCTTAGAAGGTAGAAGAAGTCTTCGAGTGCGGGTAGATTTAGATTTAGCCTTGATGTT
CACATGCTACGTAGGTGGGAAAGGACGGCTAAAGCTCCTACGACCCTAACGTATAGACTGGGTAAATCTA
GGGCCAGGAAGGCAAGTCTTATTGATGTTATGCGGATAACGCGTTCCATTACGCTTGAAAGGTAGGTTAA
TAGTGCACTCAGCAGCATTATCAGCAGTGTAAACGCTGGCTCTGCAGTTTGGAGCATAATACTTCCTATA
ACTCCTCCCAAACAAACGGCTGTCATACACGCCTTTAGCATATCTAAAAGTACGTTTAATGGTACAACAT
ATACCTTTGTATATCAATCGTCGGTAATCCAACCGGCTGTCGATGCCGCTGATAGAATTGCTCAACATAA
TAGAACTAGTACGGACAATTTACTTGATTTTCCATGGCATCATACTGCAATAGTTTTAAGGGCCAATTCT
GTTGCCAAATAACAGAGTAATAGGCCTAGTCCTGTTGCACGACGACGGATGTACTGTGTTGCAGGTACTA
TACCAATTGTCTGCACTAGTGACAGTGCGTCAGTCTTTTGCAGCACAACAAAGGTAGATAAGTCTGCACC
TGCACTGTAATACGGTTTACGGGTTTTGAGATGGGTTAAATTCCTTAAGAGTAGTGGAACTGGTACTACC
TCTATCGGACGGGAACAACCCTTTGAGTATGTTATAGGCACACGTAACGGTCTTAATCCGAAGTCTTCAT
ATTGTACTAACGAATTAAACAACCGAAGTAACGCTATGGCACCTGGTAATTCAGCTGCCATTCAACCAGC
ACCTCCTGGTAATGTAACCGCTCCGCGTGTCAATTCAACTGTATTATTTTTCTGTGTTTTTGTTGTAGAT
GTTGCTGAAGCCTGCAAAGGCTCTCCTCATGATGGGGAACAACCAATCACTGACTGTGTTGAGACGTGGT
CTATACGCATTGGTATTGGTCTGGAAACAACTCATACACGGGATCACAATATGGATCCCGGATACTCTGG
TGGTGCATCGTCCTCTCTGTGTTGCCCCTCCCACACAGAAGTTCAAAGTCTTGACGAATCCTGTAACTGA
AGAGATAAGTCTGCACGAACACATACAACCTGTGTTGATAGCCTTGCGGGTTCTATACTTAGAAGCACTC
TCTTTACCGATTGCGGTATGTGGAACGGCTGTGGCTCTAGCTGTAGAGTACGCCGAGGACCCACGCCTCT
GTTAGCTAGTGCACCATTGTGTTTAAGCGAGTATGCGGTGTATCTGCGGCTAGGCTCACACCCTGTGGCA
CTGGGACCTCAACTGATGTAGTGTACCGAGCATTCGATATATATAACGACAAGGTTGCGGGATTTGCAAA
ATTTCTCAAAACGAATTGTTGTAGGTTTCAGGAGAAAGATGAAGACGATAACCTTATTGATTCTTATTTT
GTGGTTAAGAGACATACTTTCTCTAATTATCAGCATGAGGAAACGATCTATAACCTGTTGAAAGATTGCC
CTGCTGTAGCGAAACATGACTTCTTTAAGTTCCGCATCGATGGGGATATGGTTCCACACATATCTAGACA
GCGCCTCACTAAATACACAATGGCTGATTTGGTATATGCGTTGCGGCACTTCGATGAAGGAAATTGCGAT
ACTCTTAAGGAGATCCTGGTGACATATAACTGTTGCGACGATGACTATTTCAACAAGAAGGATTGGTATG
ATTTCGTGGAGAACCCCGATATTCTGCGGGTTTACGCCAACCTTGGTGAGCGGGTCAGACAAGCCCTCCT
CAAGACCGTACAGTTCTGTGACGCCATGAGAAACGCCGGCATTGTGGGCGTGTTGACTTTGGATAATCAG
GATCTCAACGGTAATTGGTATGACTTTGGTGATTTCATACAGACAACTCCAGGATCTGGAGTGCCGGTAG
TAGATTCTTACTACAGCCTCCTTATGCCAATACTCACCTTGACGCGGGCGCTGACGGCTGAGTCCCACGT
GGACACTGATCTGACCAAACCTTACATAAAGTGGGACCTGCTGAAATACGATTTTACGGAAGAGCGCCTG
AAGTTGTTTGATAGGTACTTTAAATATTGGGACCAGACATACCACCCAAATTGCGTTAATTGTCTTGATG
ACCGATGCATTCTTCACTGTGCGAATTTCAATGTACTCTTCTCCACCGTGTTCCCGCCCACGTCATTTGG
ACCCCTCGTCCGAAAGATTTTCGTTGACGGCGTTCCGTTTGTTGTATCCACGGGATATCATTTTAGAGAA
CTCGGGGTGGTTCATAATCAAGATGTTAATCTGCACTCCAGTCGCTTGTCTTTCAAGGAACTCTTGGTCT
ACGCTGCTGACCCCGCAATGCATGCTGCCTCAGGCAACCTGCTTCTCGATAAGCGGACCACGTGTTTCTC
TGTCGCTGCGCTGACTAATAACGTCGCATTCCAAACCGTCAAGCCAGGCAACTTCAATAAGGATTTTTAC
GATTTCGCGGTCTCCAAGGGATTCTTTAAGGAAGGCAGTAGTGTGGAGTTGAAACACTTCTTCTTCGCAC
AGGACGGGAACGCGGCGATAAGTGATTATGACTACTATCGATACAACCTGCCTACTATGTGCGACATTCG
CCAGCTGCTTTTTGTTGTGGAGGTAGTCGATAAATACTTTGATTGTTATGATGGCGGCTGCATAAATGCA
AACCAGGTCATCGTGAATAACTTGGATAAAAGTGCCGGTTTTCCGTTCAATAAGTGGGTAAAGCCAGGT
TGTATTATGATTCCATGAGTTATGAAGATCAGGATGCTTTGTTCGCGTACACAAAGAGAAACGTGATCCC
GACAATTACACAAATGAACTTGAAGTATGCCATCTCTGCGAAGAATAGAGCTCGCACCGTTGCTGGGGTC
AGTATATGCTCTACCATGACAAACCGACAGTTTCATCAAAAATTGCTGAAAAGCATTGCGGCTACTAGAG
GCGCGACGGTAGTTATAGGCACTTCCAAATTCTACGGCGGCTGGCACAACATGCTTAAGACCGTGTATAG
TGATGTAGAAAACCCACATTTGATGGGATGGGACTATCCCAAATGTGACCGAGCAATGCCGAACATGTTG
CGAATCATGGCCTCCCTCGTTCTTGCGAGGAAACACACGACCTGTTGCTCCCTCAGCCACCGGTTTTACA
GGCTCGCGAATGAGTGCGCCCAAGTCCTCAGCGAAATGGTAATGTGCGGAGGGTCACTCTACGTCAAACC

GGGCGGAACATCATCAGGAGATGCCACCACCGCTTATGCTAATTCTGTCTTTAATATTTGTCAAGCTGTT
ACCGCAAATGTTAACGCACTCCTTAGTACGGACGGAAATAAGATCGCCGACAAATATGTTAGAAATCTTC
AACATAGGCTGTATGAGTGTTTGTACCGAAATAGGGACGTTGATACAGATTTTGTGAACGAGTTCTACGC
ATACTTGCGGAAGCACTTTTCAATGATGATACTGAGTGACGATGCGGTTGTATGCTTTAATAGCACGTAC
GCTTCACAGGGACTGGTGGCCAGTATTAAGAATTTTAAGAGTGTTCTCTACTACCAGAATAATGTCTTTA
TGAGTGAAGCAAAGTGTTGGACTGAAACTGACCTGACTAAGGGGCCACACGAGTTTTGTTCTCAGCATAC
GATGCTGGTAAAACAGGGGGACGATTATGTGTATCTCCCTTACCCAGATCCGTCAAGAATCTTGGGGGCC
GGTTGTTTCGTGGACGATATAGTGAAAACCGATGGTACCCTGATGATCGAGCGGTTTGTGTCTTTGGCCA
TTGATGCGTATCCGCTTACAAAACACCCGAACCAAGAGTACGCTGACGTATTCCATCTGTACCTCCAATA
CATAAGGAAGCTGCATGATGAATTGACAGGACATATGCTCGATATGTACTCCGTTATGCTGACCAACGAC
AACACTTCCCGATATTGGGAGCCCGAGTTCTACGAGGCTATGTACACGCCCCACACAGTCCTTCAGGCCG
TAGGCGCCTGTGTTTTGTGCAATTCCCAAACCAGCCTCAGGTGTGGGGCCTGTATCCGACGACCGTTTTT
GTGTTGTAAATGTTGTTATGATCACGTCATCTCTACTTCACATAAACTCGTTCTTAGCGTCAACCCTTAT
GTTTGCAATGCCCCTGGGTGCGACGTAACTGATGTGACGCAGCTTTATCTCGGTGGGATGAGCTATTACT
GTAAGTCCCATAAGCCCCCGATCTCCTTCCCGCTTTGCGCGAATGGACAAGTGTTCGGGCTGTACAAAAA
TACATGTGTAGGGTCTGATAATGTAACGGACTTTAATGCCATAGCGACCTGCGACTGGACGAATGCCGGA
GATTATATACTGGCAAACACCTGCACGGAAAGGTTGAAGCTCTTCGCGGCAGAAACCCTGAAGGCGACGG
AGGAAACCTTCAAACTTTCTTATGGCATCGCAACGGTACGCGAAGTATTGTCAGACAGAGAGCTTCACTT
GTCATGGGAGGTAGGAAAACCCAGACCCCCCTTGAACCGGAATTATGTGTTCACCGGATACCGCGTCACC
AAGAACTCAAAGGTCCAGATAGGTGAGTATACATTCGAGAAGGGTGACTACGGTGACGCGGTAGTGTACC
GGGGGACCACGACGTATAAACTTAACGTCGGTGACTATTTTGTGCTTACGAGCCACACAGTTATGCCACT
GTCAGCTCCAACGCTCGTCCCTCAGGAACATTACGTTAGGATCACCGGTTTGTATCCTACCTTGAATATC
TCAGACGAGTTTAGTTCAAATGTAGCCAACTATCAGAAGGTCGGGATGCAAAAATATTCAACGCTCCAGG
GGCCACCCGGAACGGGTAAATCTCATTTTGCTATAGGGCTGGCATTGTATTACCCGAGCGCAAGGATAGT
TTACACGGCCTGTTCCCATGCTGCTGTCGATGCACTGTGTGAGAAAGCCCTCAAATACTTGCCCATTGAT
AAGTGCAGCCGAATTATACCTGCGAGGGCAAGGGTGGAATGTTTCGACAAATTCAAAGTCAACTCAACTC
TTGAACAGTATGTTTTTTGCACAGTTAATGCGCTTCCAGAGACCACGGCTGATATAGTGGTTTTCGACGA
GATCTCCATGGCAACAAACTACGATTTGTCTGTCGTCAACGCGCGACTGAGGGCCAAACACTATGTATAC
ATCGGGGACCCTGCACAGCTCCCCGCTCCAAGAACTCTCCTCACAAAGGGAACTCTGGAGCCCGAATATT
TTAACAGCGTTTGTAGGCTTATGAAGACAATAGGACCGGATATGTTTTTGGGCACATGTCGAAGGTGCCC
AGCGGAAATCGTGGATACGGTATCCGCTCTTGTATATGACAATAAGTTGAAAGCACATAAGGACAAATCT
GCCCAATGCTTCAAAATGTTTTATAAGGGCGTTATAACCCACGATGTCTCCTCTGCAATCAATCGACCAC
AGATAGGGGTTGTGAGAGAATTCCTGACGAGGAATCCTGCTTGGCGAAAGGCTGTGTTTATAAGTCCATA
CAACAGCCAGAACGCGGTAGCGTCAAAGATCCTTGGACTCCCCACTCAAACGGTGGACAGCAGTCAGGGC
AGTGAATATGACTATGTTATCTTTACACAGACAACCGAGACTGCCCACTCCTGCAATGTTAACCGGTTCA
ATGTGGCAATTACGCGCGCTAAAGTCGGCATATTGTGCATCATGAGCGATCGAGACTTGTACGATAAATT
GCAATTTACAAGTTTGGAAATACCCCGGCGGAATGTAGCAACATTGCAAGCGGAAAATGTCACTGGACTT
TTCAAAGACTGTAGTAAAGTCATTACCGGACTGCATCCTACGCAAGCTCCCACTCATCTGAGTGTCGACA
CCAAGTTCAAAACAGAAGGTCTCTGCGTGGACATCCCCGGCATACCCAAGGATATGACTTACCGGCGCCT
CATAAGTATGATGGGTTTTAAGATGAATTACCAGGTCAATGGCTACCCAAACATGTTTATTACCCGAGAA
GAGGCCATTAGGCACGTGCGAGCATGGATAGGGTTTGACGTAGAGGGGTGCCACGCCACACGCGAAGCAG
TCGGAACGAACCTCCCTCTTCAGCTGGGTTTCAGTACAGGGGTCAATCTTGTTGCAGTACCAACGGGCTA
TGTAGACACTCCCAATAACACAGATTTCTCACGAGTGTCCGCAAAACCTCCACCAGGGGACCAATTTAAA
CATCTGATCCCCTTGATGTACAAAGGACTGCCCTGGAACGTAGTTCGGATCAAGATTGTTCAAATGCTCT
CTGACACCCTCAAGAACCTCAGCGATAGAGTCGTTTTCGTCCTTTGGGCACACGGTTTTGAGCTCACATC
TATGAAGTACTTTGTTAAGATCGGTCCTGAACGCACGTGTTGTCTTTGTGACCGAAGGGCTACTTGCTTC
AGCACCGCAAGTGACACGTATGCTTGCTGGCACCACAGTATCGGTTTTGACTATGTTTACAATCCGTTCA
TGATCGACGTGCAACAGTGGGGTTTTACCGGGAATCTCCAATCTAACCATGACCTCTACTGTCAGGTTCA
CGGGAACGCTCATGTTGCCAGCTGTGACGCCATTATGACGCGATGCCTGGCCGTGCATGAGTGCTTCGTC
AAACGAGTTGATTGGACCATCGAGTATCCAATTATTGGAGATGAGCTGAAAATAAATGCAGCTTGCCGAA
AGGTGCAACACATGGTAGTTAAGGCGGCATTGTTGGCGGACAAGTTCCCAGTGCTTCATGACATTGGAAA
TCCCAAAGCCATAAAGTGCGTTCCCCAGGCGGACGTTGAATGGAAATTCTACGACGCACAACCGTGCTCC
GATAAGGCATACAAAATTGAAGAACTGTTTTACTCCTATGCAACGCACTCCGATAAATTCACCGACGGGG

TCTGCCTGTTTTGGAACTGTAACGTGGATAGGTACCCTGCAAACAGCATAGTCTGCAGATTCGATACAAG
AGTTTTGTCAAACCTGAACCTTCCTGGTTGTGACGGCGGAAGTTTGTATGTCAACAAACATGCATTTCAC
ACTCCCGCGTTTGACAAAAGCGCATTTGTTAACCTTAAACAGTTGCCTTTTTTTTATTACAGTGATAGCC
CATGTGAATCACATGGTAAACAAGTTGTATCCGACATCGACTATGTACCATTGAAATCCGCAACCTGTAT
CACCCGCTGTAACTTGGGGGGAGCTGTGTGCAGGCATCACGCGAACGAATATCGATTGTATCTCGACGCT
TACAACATGATGATTAGTGCCGGGTTTAGCTTGTGGGTATACAAACAGTTTGATACGTATAATCTTTGGA
ACACTTTCACCCGGTTGCAGAGTTTGGAAAACGTTGCGTTTAACGTAGTTAATAAAGGGCATTTTGACGG
ACAACAGGGTGAAGTCCCAGTCTCTATTATTAATAATACCGTTTATACGAAGGTGGATGGTGTTGACGTG
GAGTTGTTTGAGAATAAGACCACACTCCCTGTGAACGTCGCGTTTGAACTGTGGGCGAAAAGGAATATTA
AACCGGTCCCAGAAGTCAAAATTTTGAACAACCTCGGCGTTGACATTGCCGCTAACACAGTCATATGGGA
CTATAAGCGGGATGCTCCGGCACATATAAGCACGATTGGGGTTTGCAGTATGACGGACATCGCTAAAAAG
CCGACGGAGACGATTTGCGCTCCCCTTACTGTGTTTTTCGATGGCAGAGTAGATGGACAAGTGGATCTTT
TTCGGAATGCCAGAAATGGAGTCCTTATCACCGAAGGCTCTGTGAAGGGACTCCAACCCTCAGTTGGTCC
TAAGCAGGCGTCACTCAATGGTGTCACCCTTATCGGAGAAGCCGTCAAAACACAGTTCAACTACTACAAG
AAAGTTGATGGCGTGGTCCAACAGCTGCCAGAAACCTATTTTACGCAAAGTAGGAATTTGCAGGAGTTTA
AACCTAGAAGCCAAATGGAGATTGACTTTCTCGAGCTTGCAATGGATGAGTTTATAGAGCGATATAAATT
GGAAGGATACGCCTTCGAGCATATAGTCTATGGCGATTTCTCCCATTCACAACTGGGCGGACTCCATCTT
CTCATCGGGTTGGCCAAGCGATTTAAAGAGTCTCCCTTCGAGCTGGAGGATTTCATTCCAATGGACAGCA
CGGTAAAAAACTACTTTATAACTGACGCTCAAACGGGCTCTAGTAAATGTGTTTGCTCTGTGATTGACTT
GTTGCTCGATGACTTTGTAGAAATAATCAAATCACAAGACCTGTCTGTGGTAAGTAAGGTTGTTAAGGTG
ACTATCGATTATACCGAAATAAGTTTCATGCTGTGGTGTAAAGACGGGCACGTTGAAACGTTCTATCCGA
AGCTCCAAAGTTCACAGGCTTGGCAACCTGGAGTGGCCATGCCTAATCTGTACAAAATGCAGAGAATGTT
GCTCGAGAAGTGTGATCTGCAGAATTATGGCGACTCCGCCACGCTGCCAAAGGGCATTATGATGAATGTC
GCTAAGTACACTCAACTCTGCCAATACTTGAACACTCTCACCCTCGCAGTTCCTTATAACATGCGCGTGA
TCCACTTCGGAGCTGGCTCAGATAAGGGAGTGGCGCCGGGGACTGCTGTTCTCCGGCAGTGGCTTCCTAC
GGGGACTCTCTTGGTTGACTCTGACTTGAACGATTTCGTTAGTGACGCTGACAGCACACTGATCGGCGAT
TGTGCGACTGTGCACACTGCCAATAAGTGGGATCTCATCATTTCCGACATGTACGACCCCAAAACCAAAA
ATGTAACTAAAGAGAATGACAGTAAAGAGGGCTTTTTTACATACATCTGCGGGTTTATCCAGCAAAAGTT
GGCCCTGGGCGGTTCCGTTGCTATTAAGATCACTGAGCACAGTTGGAATGCGGACCTGTATAAACTTATG
GGGCATTTCGCCTGGTGGACGGCCTTCGTTACGAATGTGAATGCTTCTTCCAGCGAGGCCTTTCTTATAG
GGTGCAATTACCTGGGCAAACCTCGCGAGCAAATTGACGGCTACGTAATGCACGCCAATTATATTTTCTG
GCGAAATACGAATCCAATTCAATTGTCATCATATAGTTTGTTTGATATGTCCAAGTTCCCTCTCAAGCTG
CGGGGTACAGCAGTTATGTCCCTCAAAGAAGGACAGATAAACGACATGATCTTGAGCCTTCTCTCTAAAG
GCAGATTGATCATCAGAGAAAACAACCGCGTCGTCATATCCAGTGACGTGCTCGTCAATAATTGAACAAA
CAATGTCTGTTTTTCATGCTTCATAGCGACCAGCTTGTAGTCTGTGTGCTAATCATACAATCAGAATAGC
ATCACACCATGTATTCATTAGTTTTTCCATACTTGGTGTCTGTTGCCTTAGCAGTCTTTCCAAATCCTCA
GCTTTACCTTCAACTCCGGCCTTGTTTTGACATTTCTCTTCCAGTGTTATCTGGTTCCCTGTTACACGTG
TCTTTGGGACCAGTGGTATTAAGAGGTATGATAACCTTGTCCAACTATTTGATAGTGGTGTCTCTTCTGC
TTCCATTAAGAAGTCTGACACAACAAACGCTTGGACTTTTGGTATTACTTTAGGTTTGAAGACCCAGTGC
CCACATACTGCTGATGAAGGTACTGATGCTGCTACTAAAGTCTGTGAATCAGTATTTTGTAATAGAGCAT
CTTCGGCTGCCTCTTGCCACAAAAACAACAGAAGCTCGATGGCAAGTAAGTTCAATCTCTGTTCTAGTGC
GAATAACTGCATTTCTAGATATGTCTCAGCGCTTTTAGTTATGGTCCATAACGAAAAACTGGCTAATTTC
AGAAGTCATGAGGCATCTGTGTGTGAGAATACTAGTGGTTGTTCTAAAATATTTTTTGAGCCCATGCATA
CTAGTTTAGTGCCTGATCTCCTAGCGGCTTTTTTGGATTCCGCACTATTGGTAGATTCGCGAATAGATAC
TGACACCACTAAGTATCCAATTTTACTTGTTTTACATAAAAGCTCTTTGATTCTTGGTAATTTTTTTTA
GGCTCGATTCTTGGTGTTGCTCACTCTTGTGCGGTTTGAGTAGTACATAGGATTTCAGCATAAAAATCTG
ATAAAAGTGGAATCACTATAGGTGCTGCCGGCTGTGCACGTAGCCTTCACTCAGAAACAAAGTCTACGTT
GAAATTCTGCACTGCCGCAAAAGAAATCTGAGTAACTTTTAATTGTGATCCCCTACGAATAGGATATATT
GCTGAATCTCTTAATATTACAAGTTGGTGCCATTTTGGTAATCATTCTAACGGCACCAGATCTGCATTTG
CCTGTGTCTTGAGCAAGAGGAGAACCAACAGTTGTGCTGTTGACTGTTTTGCCCAATCTAGTTCCGAATT
ATATTTCACTTCTGAGTACTCTGGTCCGTTAGTTACTAAATTAAGTAGTCCCTTCTGTATTGATGTTTGT
GCAGGTTCATATGCAATTGACGGTAGTGATCTCAGACAAATAGATCCCGCGCAAATTGGAAGGATTGTTA
ACTTTGACTTTAGATCACAAGGTAATTTTATCGGCTGCGCTATTCCTTGGAGTTCTAGCAGAGCTAATTT

TGAGGCTGGTGGTGACTGTAACTCCCAGTCTAGATCGTGTAGGAGGTCTAGTCACAGACATTCTAGGAGC
GGTATTTCAACTGAAACCTTTCAGGACGGTGACATACGTTGTAGTGGTGCTAACGGTTTTAGCTTCTTCT
GAGTTTCACGATCATTTGGTTTCCTACACATTGATGGTGTTGGCTGCCCACTATACAGTCTAGTAGCACA
TTTTTTTAAACCAGTACGTGCACATCAAATTGCCTGTGGACATAGAAAGTTTATTAGTTCGGTTAAAAGC
AAATGTGCCAATTCCAATTGCAGTGGTTTAATCGACACAGGTGTAGCTATTGAGTCTAACAAAAAGTAAG
TGCATTTCCAACTATATGGCAACGGCACTGTTAGCATTACTGATGTTGTCCCTAATCAACAGATACTTAA
GATAGTTAACACTATACTATGTTCTTTTGGTGGTGTCAATGCTATAACACACGAAACAAATATTTCTAGC
CGGGATGTTGTTCCTTGAGCGGGTGCTGACTCCACCGATCTCCGTGTTGTTACTCCTGTCGCTCAACATA
TTCCTACCTTGCCTGCCTCTTTTATAGGTTTTAGTGTTTCTCAAATACGTGTCGCCTGTTTAACAGGGGT
TAGACTTGTCAGCAGCTCATTTGAGTATAGCATACTCACTGGTGTAGATACATGCGATAGCTGAGTGATT
CAGACTAATTTAGCAGCGCAGGAACGTGATGTAGTTAATCTATACACCACTGTCTTCATTACGTTACCTG
GTGCCGAAAATTCTCATGCCTCCTTTGATAGCTCTACTGTCATACACATAAGTTCTATTACTAATGTTAC
CATAGGAATTCAACCTCCGTGTATGACCAGGATATTAGCCGGCTCTATAATGTTCATCTTTGGTGATTTA
ATTAAATGCAACAGTCATTTGTTGCCATCTGGCAGTTTCTTTATACAATTAAACCGTGCTTCAATTGGAA
CTCTTGTTAAACCCGCCAAAAACACCCTCGGAGCTTCTGTACTAGTCAGACAAACCTTCAGAACACAACA
AATTGAAGGTTCTGGTGGTTTTAATTTTTCACGAACATTACCCGGAGTATCAAGACAAAGCAAGAAGTTA
TTTATTAGAGGTCTACATTCCAGCAGAGCGACACCTGTCGGTGTTGGCTTCATCAAACCATATGGTAACT
GCCTTGGTAATACTGCTGCTGAAGGCCCCATTTGTGCACGAAGGTCTGACGGCCTTATTGTTTCGCTACA
TTTGCACATCGATGAAACGATTGTAGTATCCATTTCTGTACGGTCAGTGGCTATAATCACTTTTGGCTTG
ATTTGTGGTGTCGGTGTTGTATTACCAATACAATATGTTATGCAAATGGTCTCTGAGTTTAGTGGTACTG
GTCTTACACAGAATGTAGCCTGTAAGAACCGAAGATCGATTGTCAACCCATATAATAGTGCTATTGGCAA
AACTCTAGACTTACTTTCTTCCATTCCAAGTGCACCTGGAAGACAAGTCGATGTGGGCAACCAAAATGCA
CCAGCTTTAAACATGCTTGTTGAACGACATAACTGCAGTTTTGGTGTAACTTTAAATGTTTTAAATAATA
TCCCTTTACTTCATGACAGAGCTGAGGGTGAAGCGCGAACTAATAAGTTGATCACAGGCAAACGAGTAAG
TTCGCGGACATTTGCGATAGTACTATTAATTAGAGTTGCCGCAATCAATCATTCTGCTAAAGTTGTTGCT
ACTAAAATGTGCGAGTCTGCACGTGGACCATTAAGAAAAGCTAATTTCTGTGGAAAGGGCTCAGCTCATA
TGTGCTTCCTTCTGTTTCTACATCCTGGTGTTCACTTCTCGCGTGTGACCTGTGTCCTTGTACAAGGAAG
GAACTGCACAACTGTTCTTGTCATCTCAGTTAATGGAAGAGCACGCTCTCCTCATGACGGTGCCTCTGTT
TCAAGTGGCACACATTGGTATGCAACACTAAAGAGTTCCTGTGAACTACGAATCATTATTATAGACAACA
TATCTGCGTCTGGTAGCTGTAGTGTTGTAACAGAAATTGTCAGCAACACTCCCTTTGATCCTTCGCAACA
TGAATCAGACTCATTCAGGGGGGTGTGCGCTAGATCTTCTAAGAATCCTATATAACCCGATGTTAGTTCC
GATAGCACCTCTGGCATTAGTGCTTTTCTTGTAAGCACTCAAAAAGAAACTAGCCACCGCAGTAGGGTTG
CCAAGAGTTTAAGTAAATATCACATAGGTCACCCCGCACATGGAAAGTATAAGCAGTGTATAAGATGGCG
ATGGTGCATTTGGCCAGGTTCTACAGTTGGCTTGACTGTCACTCTAATGGTGATAACTATGCTTTGCTTT
ATGACCAGTTGTTGTAATTGAGCCAGGGGTTGCTGTTCCTTTGGATACTCCTGCAGATCTAGTAACGGCG
ATTGTAGGCATCCGCGCAGCGCTCTCAAATTACGCTCCATATAAATGAACTCATGGATCTTTTCATGAGG
ATATTTACCATAGGAACGGTCACGCTCAAACAGGGTGAAATAAAGGACGCTACGCCCTCCGATTTTGTGC
GAGCAACTGCGACGATTCCGATTCAAGCATCACTTCCGTTTGGCTGGTTGATTGTCGGGGTCGCTCTTCT
TGCGGTGTTCCAGAGTGCGTCAAAGATCATTACCCTGAAAAAGAGGTGGCAGCTGGCTCTGAGTAAGGGA
GTGCACTTTGTCTGTAACCTCTTGCTCCTTTTTGTGACGGTATATTCACATCTTCTGTTGGTCGCGGCCG
GACTCGAGGCGCCCTTCCTGTATCTGTATGCACTGGTGTACTTCCTGCAGTCAATAAATTTCGTAAGAAT
TATAATGCGGCTCTGGTTGTGCTGGAAGTGTCGGAGTAAGAATCCTCTCCTCTACGATGCTAACTATTTC
TTGTGTTGGCACACCAACTGCTATGACTACTGTATCCCCTATAATTCTGTCACATCTAGTATCGTTATTA
CTTCTGGAGATGGAACCACCTCACCCATCTCTGAACATGACTATCAGATAGGAGGCTACACTGAAAAGTG
GGAATCTGGTGTCAAGGATTGTGTGGTGCTGCACTCTTATTTTACTTCAGACTATTATCAGCTTTATAGT
ACGCAACTTTCAACAGACACAGGTGTGGAGCACGTGACGTTCTTTATATACAACAAGATAGTAGACGAAC
CAGAGGAACACGTACAGATACACACAATTGATGGTTCTAGCGGAGTCGTAAATCCTGTGATGGAGCCAAT
TTACGACGAACCGACGACAACCACCTCAGTACCCTTGTAAGCCCAAGCAGACGAATATGAGCTGATGTAC
TCTTTTGTATCAGAGGAAACCGGGACACTCATCGTAAACTCCGTACTCCTTTTCTTGGCGTTTGTGGTTT
TTCTTCTTGTTACGTTGGCAATTCTCACGGCCCTGAGACTGTGTGCCTATTGTTGCAATATCGTGAACGT
GTCTCTCGTCAAACCGTCTTTTTATGTTTATAGTCGAGTGAAAAACCTGAATAGTAGTCGAGTCCCCGAT
CTCCTGGTCTGAACAAATTGAATTTTGTATTAGTTCTTCTGCTTGGAGCTTTAATTTTAACCCTGGCAAA
TACCAACAGTGCTGCTCCCTCTCAAATCTCTTAAGTCATCTCTCAACAACGGAACTTGATGATAAGTAAG

TTACTCCTTGCACGGGTTTGTCTTTTACAACTTGCCTATGCCTACCGGCATCGGCTTTTGTATTTAGCTT
AGTTAATTCTCTTCCGGGTGTTATGGACAATAACTGTAGCTTGTCCTTTGCCTGCTCCTGTTCACCGAAT
AAATAGGATCTCCGGTGGAGCTCCTCAGCCAATGGTTGGTCCTCTGAGCGTGATGTGGCAGTGCTACATC
CCTGCTTCTCAGTGATTGCCTTAGGGTAAGGGTTCCGTGTGGCCACTCTATACAGAAACTTACATTCTTT
TCCACGTGTCACAGTATGGCGCTGTTCTAACCGGACCGATTTTAAAAAGTGAACTCCTAGTCCGAACTCT
GAAGTTTCGTCGACATTTTCGTCCTGCTGGACACCATCTAGGATGCCGTGACAAGCCGCACCTGCTTGAA
GAAGTCCCTTCTGCTGCACCACGAGAGATTCCTCATAACCAACTGGGAACTCCGATCTGTTTAACAAGTA
ACGCAAGTCTTGCTTCACACTGTTGCGACGGGGCTCGCTACTATTAATTGAACTCAGACGATTCCAGTTG
CAGTTACGATCCTCCTCTGTCTGTATTCAAAGTGACAGCAAATGTTCCATTTGGTTGATTTTCAAGTGAC
CATTGCGGAGATCCTTTTGATAATCATGAGAACTTTTAAAGTATCCATTTGGAATCTTGATTATATTATC
AATTTGATTATCAAGAATCTGTCAAAATCTTTGACGGAAAACAAGTACAGCCAACTGGATGAGGAGCAAC
CTATGGAGATCGACTGAACGAATATGAAAATTATTTTGTTTTTGGCACTCATAACTCTTGCTACCTGCGA
ACTCTATCATTATCAAGAATGTGTGCGGGGTACCACGGTATTGCTTAAAGAGCCGTGCAGTAGCGGGACT
TACGAGGGCAATAGCCCATTCCACCCCCTTGCAGACAATAAGTTTGCGTTGACATGCTTTAGTACTCAAT
TTGCGTTCGCGTGTCCTGATGGCGTTAAACACGTCTACCAATTGCGGGCTAGATCTGTAAGCCCTAAACT
GTTCATCAGGCAAGAAGAAGTTCAAGAACTTTACAGCCCGATATTTCTTATTGTAGCCGCTATTGTTTTC
ATCACGCTTTGTTTTACCCTTAAGCGGAAGACCGAATAACTGAACTTCCATTAGCTCACTTCTATCTGTG
CATTTTAACCATTCTGCTACAGCTTGTTTTAGCTGTGTCTTCTTAGTTTTGGTTCTCACCTTAATTGCAA
AATAATTATGAAGTTGGTCACCCCGAAAAGGACATGAAACTTTTTGTTCTCCTGAGAGAGTTCACAGCTC
TAGCTGCATTTTACGAAAAACGTAGTTTATTCACATGTACTTAACATAAACCATATGTGACTCATGACTC
GAGTGCTCTTCACGAGTATTCTCAACGGGATCTTGGAGTAAGAACTGGAAAATCAACATTTGTAATTGAA
TTGCGCCTGGATGAGGCTTGTGCTTAACCACCCGTTTTCCACTTCCATAAGCGTGATTATTCAATTCCCG
GTGTATCTCCTTCAGCTTATTGCCCGGAACCTTAATTGGGTCGTGCTGTAATGCGTGGTGCGATCAATGA
AAACCTTTTAATCAATTATGACCTTTGTGCTCTTCTAAATAAGTAGTAAGCGCACAAACTGAAATGTATA
GTAATGGACCCCCAAATCTGCCAAGTGCACTCCACATTATGTCTGGTGGACTCTGCGATTTAACTGGCAA
TGACCTGAATGGCGGACTCAGTGGGGCGCTATAAAGACTACATCAGCTCCTCGATTCACGCAATGATACT
GCGTACTTGTTCATAGATCACATTCCACATGGCAAGGGAGGCCTTAAATTCCTAGTCGGACACGAAGATC
CAACTGACATCAGTAGCAGTCCCGATGACCTAATTGGTTGCTTCCTAAATCTTACCAAACGAATTCTTGG
TGGTGAAGGTAGAATGAACGCTCACAGTCAAAAATGGTTTTCCTGCTTCCCCGAAACTGGGCTCGCAGTT
GGACTAGTCTGTGGTGTTAACAGCGCCGACACCATATGGGGTGCAATTAGGGATCACTCGAATACACAAA
ACGGTCCCACTGGCATCCTCAATCTTGCTGACAGTGCTGTAATCGAGCGACGACCAGTTCTAGGAACAAT
ATCGCGAAGAGGCTCCTTAGGAGGCGAGAGCAGCGGCGCCAGAGCTCTCTCTTCTCATTCCTCATTACCT
GATCCCAACAATTTAAAAAGTTCAACTCCCGGCAGCAGTAGGGTAACTTTTCCTGTTAAAATGGATGGCA
GTGGCGCTAGTGCTGCTCATGCTTCGCGGCAGCGTGACAAATAGAACCAGCATGAGAACAGAACGTCTGG
TAGCGGCCCACGACCACACGACCTAATTGCCACTAAGAAATATGTTGTTAGGGCTTTTAAGAAGCCAGTG
CTAAAACTTACTGTCATTAGTCAATACAGTGTAATACGAGCTTTCGACAGACGTGGAGTAGAACAAATCC
GCGAAAATTCTGGGGCCCTGGTACCAATCAAACTAGGAATTGACTGCAAACCCTCGCAGCTAATTGTACC
ATATGCCCCCAACGGTTCAGCGTACTTAGGAATGTGGCTCACTGGCATGGGAGCCATACATTCGGAAACG
TCGTTGATTTGCATCGCTGTCACCAAATCGGGTAACAACGATCCAAGTTTCAACGATCAAGCCATTTCGC
AGAGTAAGCATACTGAAGAATTCAAAATATTCCGACAAATCGCGCCTGAAAGGGCCAAAAGGAAGAAGGT
TAATGAAATAGTTCACTCACTGCTGAAACTGAGGAGACTGCCAATTGTGATTCTTCAAGTTGCTGCAGAT
TTGGATAATTTCTCCAAACGATAGCGACTATTCACGAACAATGCTAGCTCAATAGCGGATTGAACTCTTG
TCGACCACACAAGGCAGACGGTCTCTATAAGCGCTTTAGGTTTAGCGTTTATGATATCTAAAGTACCTTG
GTGCAGAATGAGTTCTCCTAGCTCCACTCAACGAGTCGATGTTCTTAACTTTAGAGCCACATAGCGATAT
TTAATCAATGTGTTACCCTGGGCCGCACGTAGAAAAGTCATCATATATTTACGGAAGCTACGCGGTCAAC
CATTGAGTGCACTGTTAATAATGCGAGGGAAAGCTGCCTCTATGGTAGGGCCTTGATGTGTAAGATAAAT
TTTAGTAGTGCCATTCCCATGTAGTTCTGATAACTTTTGCGACGCATGACGAAGAAGAAAAAGAAAAAAA
AGAAGAAAAAAAA

**>VT4**

ATCAAGGGCCTGTACCTGCCTAGATGACAGACCAACCAGCTGAGCATCTCCTGCAGATCCGTGCTGTGAA

CAAACTTCAAGATCTGCGTGGCCGTGACCAGACTGCACGCCTGATGTACACATGCCGTGTGACTGATCAC
CAACTACTGCAGATGACAGGACACCAGCAACAGCAGCATCTTCTGCCGGCTGCTGACCGTGTCATCTGTG
CTGCAGCCTATCATCAGCACCAGCAGATTCAGACCCGGCGTGACCGAGAGATGAGATGGCGAACCCTGTC
CATGGTTCCAGAGAGAGAACACCCGGCCTACACAGTTCGCCTGCTTTACCGGAAGCAGACGGGCCAGAAC
CTGGCTGTGGCGACTTAGAGGCGGAGGCCTGATCAGAGGCACCTCCACATCTTGAAGATGGCACCTGTGG
CTGAGCAGAAGCTGAAAGCGGAGATTCGCCAGCACCTGAACAGCCCTGTGTGTGCACCAGACCTTCGGCT
GTAGCAACTGCACCTCCTGGTCCTGTTATGGCTGAGCCGGCAGCAGAACAAGACGGCACAGCGTCAGATC
CTAGTGGTAGGACACATGGTGCCCCTGTCCTAGCTGCGGCAGAAATACCTCTGGACTGCCTCAGGGCAGC
AGCAGCTGAGAGCGCTGATGAAGATCCTGGTGGCCCTGACTGCGGCGGAGAAGCAAAGTGATCTAACTGA
GGCGCAGAGCCTGGCACTGAAGCCTGTGACGGTTCAGCAGAAAGCTGGAACACTGAACCTGACAGTGGTG
CTACCCCTGAACACACGCCTGAGCCTGAAGAAGAGGCATCCACAGCCTGTGCAGATAACAGCTGCTGTGG
CCTTGATGGCTGCCCAGCTGAGTGCACTGAAGGCCTAGCAGCACCTGTTGGTAGAGCTTCATGCACTTCG
TGCGGACCACCGGCCTGTACTGACACTGAGAGGGCTGTATCCTGCTGCCTTGAACCTGAGCTTGAAACTG
CCTGGTGCACGGCACCTTTTGAAAAGAGCTGTGAATCGCCGACACCTTCTGAAACTGAATCGGCAAAGAG
ATCTGACACCTCCAGTGGGGCATGAGCAAGTTCTGCATCAGCCTGAAGTTCCACAACCAGGACTACAGCA
CCAAGGGCTGAAAAGAAAAAGCCTGATGGCTGTACGGCTGAAACAGCATCTGCCTGAGCAGCTGCGTGAC
CAAGTGAATGCAGCCCAACGTGCCCTTCAACAGCCACGAAGTGTGATCCCTGTGGTAGAATTTCATGGCC
GACGGCCGGTTCTGCTGATCCCACCTGAGAATCCTGTGGCACTGAGAGTTCGACTGACGGCGGTGTCACT
ATCTGTGGCTGCTCACCCCTAAGTGTTGCTGCTGAAACCTGCTGAGCAGCATGTCCCAGTTCAGAAGCCG
GACTTGAGCCTGATCCTGCAGAATCCCCTGATGAATCTGGCTGGAAAACCACAGCTCCTGAGGCTGGTCC
CACTACTGCCTTTGGAGACTGTGCGTGCTGCTGTGTTGGCTGCCATGACAAGTGTGCCTGCTGGGCTCCA
CTTGCTGAAGATGACACCGGCTGTGACCCTACCGGTGCTGCTGGCGGAGGTTTCGGAGAAGCTGATGACA
GCCCTCCTGAAACACCCCAAAGAGAGAGAGCCAGCACCAGTACTGCTGGTAGCTGTGAACTTGATGACGG
GACAGACACTACTTCGGGATATTCTTCTGCTTCCACAAGTGCTTTTGCGGCAACTGCGAGAGATTCGGCC
TGTGATCCATCCAGACCAACTGCTGAATTCTGTGGTAGTTCTGAAGCTACAAGCGGAAGTCTTGAAAGCG
GTGCCTGGAATACTGGTAGACCGAGATCAACACCGAGAGCAGCCTGTGTATCTGTATCCGGGGCTGCTCC
TGCTGCACCATCAACTTTCTGCCCCACAGCTGAAACTGCAGCAAGTTTTGCGCCTGTTTCACCGAGGGCA
GATACAACAACACCCGGTGGAATTTCACCGTGTTCACCGAGACACACTGATGCTACGACGTGCACATCTG
ATTCGGCTACTGACAGTCCAGCTGCAACGGCCTGCACTACAGATGGTGCTGCTCCGTGGATTTCGCCGTG
GCCAATTGACATCTGTGGCATTGCCTGTGAAAGACCCAGACCAGGCCTTGACTGGCCTGAAGAGAAGTCT
GAGGCCGGTGCAGAGTGTCCTAAAGAAGGCTGGGCAATTGCTGAATCTACCTGAACCTGTGCCTGTAAAA
CTGCCGGTGGACCAATTGCCACCTGTGCAAGGGCAATTGAGGCGAGTGCAGCGACATCCTCTGAGCCTGC
AAGTGAATCTTCGGCTTCGTGTGCTGACTGTACCACTACTGGTGGTCCTGAACATGAAGCCTGGAATTTC
GCTGAAACATCTGTCACGCCCTGAAGGGCATCGTGCAGAAAGTGTGCTGAATCCAGCGGCGGAACTGGCC
TACTCACGCCAGCAAGAGCCCCAAGAGAAACTATCTGCTGAGAGGCCGGAACACAAGCCACAGAAGCGTG
AACAGAGGCTCCTGCCTCGAGAACTGGTAGTTCACCACCATCCGGACCACCTACTGATGAAGCTGCTGAA
GCTCCATCGGCTGGTACACCAGCCTGTATTGAAGGGCCTACGTGGCCCGGAACCAGCGGCATAGAAAAGT
GCTGTGCCCCTGCACCTGATACGACGGCAACAAGCAGTACCTGCACACCCAGCGGAGATGTACCAACAAG
GGCTACTTCTGGTAGTGACACTGCGACAGAAGCGCCAGGCTGCAAGAGTGCGAGTACCACTTCTGAACCT
AGTGAAAGGACTGATGATCCACCTGATGAGAGGTGCTGTGTCTGTACAGCTGAACCCGGTACAGATCCAA
ATGAGTGCGGCTGTGCTGCGGCCGGTGCTGTCACAAGAATTTCGCCACCAGCATCTGAATCACCTACACC
ACAGGCCACTAGTTCCGCTGAGTGGAATACGGCTACATCCTGCTGATCTGATGAGTGTGGTGAGTGTGAA
TCGGCTTTACCTACGTGCTGTTCTTCCTGCCTAGCAGATGAGGCTGACGGCGGAGATGACTGTGAAGGCG
GAGAGTCTGAGCCATCAACAGCATCTGAGTGTGGTACTGAAGATGACTGCCCAGATGAACCTTCGGAATC
TGGTGCCACTTCTGCTGCAGCTCCACATGAAGAAGGGCCAGACGCCGGCTCGTCAGATGATGATGAAGCA
CCAACTGCTGGTCCACCAGACGGCAATGAGGCCAGAGCGACAACTACTACTCCAACAACTGCTGAGGCAG
CACCAGCATCAGAGATGGCACCTACACCAGCTGCAGCGACTACTGAAGCGAGTGATTCTGATGGCTGTTC
AAGACCTACTGACAGTGCATCCACTGAAAGTGCAGACACTGCGGCAGAAGCTGAAAGGGCAAGACCAACA
GCGGCTGCTGATGCTCTCAGTGCCTGCCTTGAACATGGCGGAGGTGCTGCAGATCCCTGAAATGAGGCTA
TTGACAGTGCCACGCCAGCTGAATCTGATGACTGCACAGCTACTGATGGACCACCTGAAGCGGCTGGTAG
CTGTGTTTCAAGCGGACCCAGAGCTGCTGAACCCTGTCCAGCTGCTGCAGACCCAAGTGCTGACAGAGAT
GACGGCACTCCACCAGCTGAGAGTGCCTGTGAAAGTTCTGAAGCGCCAGAAGCAGCACCTGTACCATTAT
CATCAGCTGGTACTTTTGGTGCTGACCCTACACCTTCTTCAAGAGCCTGTGCCGGTACTGCAGCCACAAG

TGCCTGCTGAGCTGCCTGTGATAGAAGTCCCTGTGACAGACCTGCTTCAAGCTGTTCGGCAACGAAGAGT
GAAAGGCCTCCTGAACAAAGGACCGCTGAGACAGCTGAAGAGGCAGCTGAGCTATCTACAACTGAAAGTA
GACCTTCAGCTAGACCGAGAAAACCCGCTGATGAGAGAACCAGTCTCTGTGCTGACGGTCCTACAACAAC
TCCGGCAGAAACTGAGTGCCCCACCGGAAGCTGGTCACCCTGTACTGACACTGATGGCAGAGCAGCAGCC
GGTTCTGCCACTCCTGTTGATGACACTGACACCACTTTCTGAAAGAGCGGTGCAGCATCTACTCCGGCTG
ATGCTGTAGCCGGGGCTGCTTCAATTGCTGCGGCTACACCTACTGAAAAGGCTGGTGGCACTACTGAAAC
GCCAGCGAGAGCTTCGAGAAGTCCGCCAACAGACAGCTGTACAACCATCTGCCTGGCAGCGGCTTCAAGT
GGCTGCATTGCAGAGGCGGCAAGGACAGCGCCTGAAAAGTGTGAAAATGCCTGCTGCACAGCACCATCTA
CTACCTGTGATGAGAGGCCCGGAACAGCTGGAACTGCTTCCTGGAATTCGCCCGGAACGCCTGCACCTGT
AGACGGAACACACAGATCAACGCCTGCCTGTGCGGCAACTGAAGCCACAGCTTCAACTACACCGCCTGAA
TTTGAGGGTACTAGAACACCAGAGGCTGCGGCTGACTGTGGTGCTGAATTCTGCTGCTGCACCAGTGAAA
CAACTGCAGCGTGACCTACCAGCACACCTGACGCAGCAAGTGAAACTCCTGCTACAACGCCACCTGGCTG
TGCAATACCTGGCTGAAGTTCGGCAGATCCTGCTCCGTGTACGAGATCAGCCAGAGCGCCAGCTACAGCT
TTTGCTTCTTCACCTGATGCTGCTACAGCGTGTGATGGCTGAGCTACTTCTTCTTCTGAAACACCTGAAG
GACCTTCTACTAGAACCACCTGACCTGCTGGTTTCTGTGACGCCTGGTGCTGTTTTGGACCATCTACACC
ACCAGATACCGGATCTCTTGAGAGCGGTAGTGAAAGTGTATCCTGCACTGATGAAGCTATCACATCCCTC
CACGGTGGTAGAGCTACCATCTGTGACAGTCTTGAGATACCAGCTTCTTCGAGCGGAGCGAGGACTACTG
AGGCGTGTACAACAGCCGGCAGCACTGACCTCCACACGCCTCTTGTGGACACGTGAACGACATCTGGACC
ACCGTGTGGTCTAACCTGTTCGGCTGGTCCTGATGTTACTGAAACAAGACCTCCTGATTCACCTGACGGT
AGAACATCCTGTGCTTCACCTAGTGATGACATAGCACCTGTTGAGGGTTCTGAGTGCTGCCCCACAACTG
ATCCTGATTTTCCGGCTGAGTGCATGTGTCCATCAAGTCTCACTGAAAGGTGGAAATCCCCACCAGCTAG
TGGTTCAACTTCTACTGAATGGGCCGCTGACAGCTGCTGTCCTGCCACTGCATTGTGAACACCCCTACCA
ACAGAGTGGAAGTGTGATCCACCTGTAGCACCCGGTGTCTGCTCCAGAGCAAAGGATGGTAGTCCTGTTG
ACTGCTGTGCACCTACCTGTCTCTGCTCTGATGAGACTCCCGCTGAGTTAGATGATGTTGACGGAACAAC
GAGCTGCTGGTGTCTACCTGCCAGTTCCGGTTCCTGCAGAAGTCTCTGGAACGGGGCGTGTGAAACCTGT
GGACCACAGCCGATAATCCCTGAGGCTGCAGGTCCTGTTATGTGCACGGCCACACCTTTCTGTGAACCAT
CTGAGAGAGATGCAGCGACACACTGTACGTGTGGTAGACCAGCTACAAGATCAGCTCTACAACCGGCGTG
ACCTTCTGCTACGATGTGTCCACCACCTGTTCCGTGTGAACCTGAGCCTGGTACATCTACCTGTGCTGAT
GAGTGCATTGGTAGCTGCCAGTGTGGTCCCTGTAAACCTACAACTTCTGACGGAACTTCGTGCTGCATCG
GCGGTGTTTCACCTACAAGGTGCTGAGAATCCAGCGGAGCTACTACGGCTGCTTTCTGCAGCGGAAGCAG
CTGCACAACAACCACAAGACAAGCTACCTGTAAATCGGCTGGTGTTGCCTGTACCGGAATTGACCCTGAG
TGGGCCAGCTGCTCTGAGAGCGGCAGTTCCTGTTTCACAGAGCCACCAACTGAAGCTGCACCAAGCCTAC
CATCAGCAAGCGGAAGCTGAGATGATTCTGAGTGTGCATGTGATGATACCAGATCTGCTGATGATTCAAG
CCCGTGAACTGGCTCTGAGAGACATGCTTCAAGCGGGCCTGAAGCTACATCTTCCCATGACTGAAGTGGT
AGTGCGGCGGCTACTGACTGTGAACCCTGCACACCCTGTTCTGAGAGCGGAGCTGAATCGTGACCTGAAC
CTACTGCCTGGCCTGCTGACAGTGCAACTGATGAAGCCACGTGTGAACAAAGTACCTGGTGTACACACTG
AGCCTGGAACACAAGACCAGCTGAAACATCAAGTTCGTGTGATGCACCGAAGTGCGGGGCAGAGCCGGCA
ATGGATGAAGCTGTCTGCGGCGGAGCAAGACAAGCCTGTGAAGAAGCAGCGGCAAGAGCTACCACACCGA
GCGGAGAAGCTGAGTGTGATGCGAGAACTACAGAAGCTGCAGACGGCACTACACCTGAACCAGCAAGTGA
TAGTTCAAGAATTATCGGAGAGGCTGGCCCCACCGGTCCAATGGCTGTCTGTGCAGACAGTTCTGATCCT
ACTACTGAGAAACCTGATGAATCATCTGATCCATCCGCTTCGAGAACCCCTGCTACAGCTGGTTCTCCTG
CTGTTGATGATGCCCTCTGGGCTACTACAGCTGACTGTGCTGAGCCTTCAGCTGACAGAGCTGCTGATAC
AACTACTGACACAGCTACACCGTGTTCAAGCCCTGCCTGTATTGACTGTACGCCCTGTTCCTGTACTTTA
TCGCCACCATCGTGTACTTCTACTGAAAGTACAAGTTCTGAAACTAGAGCATCTACGCCGACTACTACTC
CAAAGAATACTGCTGAGAGTGCCGCTGAATCCTGAGCCGGGGCTTCATCTGACTGTTCGAAGTGACCTGA
TTCTTCTGAACCGACAAGTACTACAACCTGGTGTTCACCATCAAGTGCCTGCCTCGGTTCTTCAACCTGC
TGAACCGGTGCTTCAGATGCTTCAACGTGTGATTCCGGCACGCCTTTCTGCTGTACTGGCTGCAGAGAAG
GCTGTTCGAGCTGTACTGATGCCACTACTGTAACCTGCTCTACTGGTTCTACACCCTGTGATGCCTGAGC
TGATGGTTCCGGTTCTTCCGGCACCTGAGCTTTTTCCGGAACTACACCAACTACCATTTCATCTTCTAGA
TGGGCTTCAACTGCTTCTGGCTGAGCTGCAGAGTGGTGTTCGGCATCTACAGCTTTCACTGAGTGTTTCT
GTGCACCTGGATCGGCTGCAACCACGCCATCGTGTTCCAGCTGTTCTGCTCTACCTTCTACTAATGATTC
CTGGCCTACGTGGTCAACAACTGATCCTGCACAAACGGCCCCGACTTCAGCTACGGCTGAAATGTGCATC
TGCTGTGCATCATCCTGCTCTGCATGGAAAGCTGTGCGCCTGCTGTCGGCGGCTGTGATTCATTAACCT

GTACGACGTGCTGCAGACCTGATGATCCAACAAGAGCCGGATGTACAATTACTGCTAGTGGTGCTGAAAG
GTGCTGCTGTGCCTGTGTTGATGGCGCTAACGGCTGCTCCAGACAACCCAGTTGGAGCTGTGTTGACTGT
GATATATCCTGTGCTGGTAGTACATCTACTGATGATGATCCTGCGAGCGGCTGGTCACCACCGTGTGAAA
GACCAACAAGTCCTACTGACCCGTGTTCCTGCACCGCTGATGATGCTACTCCGAGGAATGGTTTCACCCC
AGTCTGCTCTGATGATCTTGGAGCAAGGACCTCTGAAAGACCTTCAGCCTGAGCTTCTGCTGACTGAGAC
AGCCCGAGAGCTGATGACACTGACGGTTTATCGCCTACTAGTGCTACTCCTTCTAGTGGTAGATCAAGAT
GTGACGGATCATCTGCAAGATCAGCGTGTGCCTGCTGCAGAGCGCCTACGTGTCCACCTACACAGTGACC
AGATCCGGCATCTCCGTGTGATGTTGGTAGTGATGCGGCAGCTGCTCCTGAAACGTGTAATGCCTGCGGT
AGTACGTGTTCATCAATTTCTGACGGACCAACGGCAAGACCCAGAACACCAGCTGCAACTGCAGATCCTG
AACCTGCAAAGAATGCGTGCTGCGGCAGTGCCTGATCTACTTTTACTTCAGCAGCAGCGCCAGAGTGTGC
TGATTCCGGTGCAGAAACTGACGGTGCTGTTGAATGTCCTGAATCGTCACCAGCATCTGACACCGCAGCT
ACTGGCGCTGACTCTGATGACTGTATGCCCACCTGTGACAGTCCTGAAAGCACGACACCCCTTGACCTTG
GTGTCTGTACTAACTGTGATGCGCCTCCTACTGATGCGCCGGCAGCAAGAAGTCCCAGCACTGCTTCGAC
ATGGAAAGATGACGGTTCCACGTGATCGTGTGAACCACCACCAAGACCAATACCTGATGCTGCTAGAAAG
AGTGACTGACCTTCTGAGTCGACATGTGCAACTATTGAACCTCCTGCTAGTGCTGCAACAACAAGGACAG
CACCTGAGGCTGGTAGAACTGCTGATAACTGGTGGAAGCCGTCAACTGAAGCTATACCTGCGTGCCCTTC
TGCTGCTGCTACTTCCTGTTCAACAATACCTGCAGCTGCCACGTGTAGACCTACTGACTCTTCAAGTGAA
ACCACCGGATCCAGGGCTACTGATGGTGGTGCCACAGCTGACACTCCATCTACAGATACCTGTTCTGTTG
ACAGACCTGCTAGTTCTGACACATGGTGTAACCCGCCTGGTGGTAGCTGTATTGATGACAGAGCCTGCCT
ATCGACTGCTGCAGCCACAACAAGAGAAGCGGCTTCTGCAGAGCTTGGTTTGCCTGGCACGACATCACCC
ACAATTGATGGTAGCTTTTCGCCTTCCTGACCTGAAGCTTTTGATGCTCCTGGTAGCACCTCCTGCACAC
AATCAAGACCTACCGGGTGCACTGACTCTGCAACATCTCCCTGTGCTTCGGCTGCTGAATGTACAACTTC
TGACGGTGCTTTTGGTAGGCCAGCACCATCCTGCTGTGATACCAGTGCACCAGACGGTTCTGCTGCCTGT
GAAAGTTCACCCCTTGACACACCCTGTGTGCCCACGGCTGGCTGTACTACAGCATCAGCTGACATCTGCC
CTGACGGTTTTGCTGAAGCGGCAACAATTTCTGATTCTGAGTGCTCTGAGCCCGGCACCTGTGAAAAATC
AGATCCTGGTGCCTGTGCATCTACTGATGGTAGATGGGCACCTGACAGTGACTGCTGCAGATCTTCACCC
GGTCCTTTCTGTGGTGCCGGTGCTGCAAGTTCACCTACTGATACGTGTACACCACCAACAGCACCTACTG
GTGCTTTGGCCACATCAGCATCTACAGCAGCTGGTGGTACTGCAGCTACCGGTCCAACATGCCCTGCCTG
CTGTTCTACGAAGTGTGAAAGAGCTTCTGGTAGATCCAGAGCTGCTCTTGCCTGTGATACTTCACAATCC
CCTACGTGATCCACTGCACCCTGTTCAACACCAGCCTGCTGATCCTGACCTGGTGTCTGTTCTGCTACCT
GCTGGTGCTGGACATCCTGAGCTACTGATGATGCTTCTTCTTCTCCACCTACTCCGTGGACGGCTACGTG
CACACCTTCAGCACCTTTCTGGACAACAACTGCCTGTACCACCTGTACTTCCACAAGGCTTTTCTGCTGG
TCCTGTGATGACTGCCCAAAGAAACCTGCAGCCTGTGATGGTGCTTCCTCTGATACTTCTGAAGATCCTG
CGCCGTGCACCTGTTCGTGAAGTGACGGAACGTGTCCAAGGTGGCCTGATGATGTGCCATCACCAGCTAC
GCCATCTGATGAATCCTGTCCTCTCTGTGATGAGTCCAGGTGTTCTGATGGTCCAACGGCTACAACTGAC
TGCAGCGGAGCTGTCTGCTGAGCAGCAGAAAGGGCAGCCAGTGACTTCAGTGACTGAGGTTCTGATGCAG
CCTGCCTACCACCACCAATCTGTATCACCTGAGCTGCTTCGCCGAGTGGTTCTGAAAGAACGGCATCCCC
ATCTGGTAGAGCTGAGGCCTGTACGGCACAAGCAACCTGTGGTACAACTACACCTGACGGTCCCTGGCTT
GATGAAGAAGCCTGCTGTCCAAGACCTGCGACCTGCATCTGTGAAGGCACGCCTGACCTTGACTGTGACG
GTTCACCCACAGCTGAGTGTGAAGCTGATTCCTCGGCACCGGCTGGTAGTGTAGCACCCAAGGCTACTGG
ACCTTCTACGCCAAGCTGTGCACATGAGCCTGAGGCTGATACAGCCAGAGCTGAGACACCTGAGTCTGAG
TGTGCAGCCACAGCACCCGGACCGATTTCTTTAGCGTGTCCCTGCTGCAGTGGTTCACCATTTGGTGCCT
GCCTATGTGCTACGAGGCCCAGTTCCACTACTGAGGCTTCATCCCTTGATGGTTCATGTGGTAGTGCTGG
TTCTGACACCGGCTGTGACTGTGCCTGTTCCTGCTGCATGCCCCTTACGGCATCACCAATTGGAGCAGCT
GTTGGCACAGACTGCGGAGATGACTGCTGTGGACCTTTTGCTGACAAGCCAACTCCACCAGCTCTTGGTA
CGGCCACAACTACTACAGCTGATGTTTCAGCCTGGTCGTGCGGTGCTGCTACAAATGGCGACAGGTGGTG
TCCCAGAGCATCTACCACAATAGCTGATAACTGTGACCCTGCGGCTACGAGGTGCAGCTGTGAACCTCCA
ACACACGGCCTTGCTGACACACCAGAACAAGCTTTTGCTCCAACTGGAACTGCCGGTTCAGATACGTGTG
CTTTATCAAGCGGATCACCGCCAAGTGGTACGAGTGGACCTACCACATCGGCTGATGCTTCATCAGGCGC
TGAATCTACACCTTTTGATGCTGCTGAACCATGCTGCGGTGCTACTTCCCCAAGTGCAGCGAGAAGAACA
ACCAGGGCTACACCCCTCTGGTGGTCACCCACAACTTCGACTTCACCTTTAGCTTCAGCCCCGAGTACAG
CATGGTGTTCGTGCTGTTCTTCGTGTGAAAGTGCCTGTTTACCTTCTGCTACGGCTACTACTGCTATGTG
TGCTTCTGCAACGACGTGTGCCAGACCTGAGCCTGTATCTCCCTGTTTGTGTTCGTGACCTTCAGCTGCC

ACTGTTCCCTGTTCTGATACGGCCTGTATGCCTGCTGACTGGGCGACGCCTACTACGACATGGTCGGATA
CGGCTGATACTGATTCGTGTGGTTTTGAGCCAAGCGGCTGTGCTACGTGTGCATCTCCTGCAGCGTGACA
AACCCCTACGACAGCAAGAACTGCGTGTGATGATGGTGTTGAGAGAGCGTGGACACCTACGAGTGCCTGG
ACACCAGACTGTGATCTCTGCTTTGGTAGTGTTTCCGGTCCAGCCACTTCCACGTGGGCAGCTACAACCT
GTGCTACTTTTGACTGCTGAGGTGCTCCTACAACTGCCACGTGTTCGGCCAGCGGTACTGCTTCTACGTG
TGTTGAGTGCTGCCCTACTTCCTGCACAACTGGTAGTACACCTCCGTGTACAACGCCTCTCTGCTGTTCC
TGAGACTGTTCCTGTACTTGCTGCTGTGGCCCCTGCTGTTTACCCAGCCTCTGCTGTGAACTGACAGCTG
GTGCCTCTAACTGCTGAGCTTCTACACAGGCGTGTGAATCTACGAGTTCACCGGCACAACCCCTACACAA
GAGTGACACAGATGCCTGCAGACCCAGCACTGAATCGTTGGCTGCTGGTGGCAGACCCTGTACCAGAGCA
GTCATTGCACCGTGTGAAACGTCCGGTGCAAGGTGCACATCAGCTCCCTGACACTGAGCTTCGCCACCAC
ACAGAGCCGGATCATCATCTGAATCGTGGGGAGCATGTGCCCCGTGACACAGTGACACTCCCTGAGCTGA
CGGTACTACTGATCTCTGTGAAAGAATGGCTTCACCACCTTTTGCTTCGCCTTCCACGCCGGCTGCTGCA
GACACAAACAGGCCCTGTGAAGAAACGCCGGCCAGCAGGGCAATCTGACCAGCTACTCTCTGAGAGTGTG
ATTCCCGAGCATCATCTGCAGCTTTTGCTACTGCAGCAGAAGCCTCTGAGCCGGCTGTTGCTGATGGTAG
TTCTGAAGCTGCTCCTGAAAGGTGGAAGAGGTGTTCGAGTGCGGCTGAATCTGAATTTGACCCTGATGCA
GCCACGCCACCTGAGTGGGCAAAGATGGCTGAAGCAGCTACGACCCCAACGTGTGAACCGGATGAATCTG
AGGCCAAGAGGGCAAGAGCTACTGATGCTACGCCGACAATGCCTTCCACTACGCCTGAAAAGTGGGCTGA
TGATGCACCCAGCAGCACTACCAGCAGTGCAAGCGGTGGCTGTGTAGCCTGGAACACAACACCAGCTACA
ACAGCAGCCAGACCAACGGCTGCCACACCAGACTGTGACACATCTAGAAATACGTGTGATGGTACAACAT
CTACCTGTGCATCAGCATCGTGGGCAACCCCACCGGCTGTCGGTGCAGATAGTGAAACTGCAGCACCTGA
TAGAACTGATACGGCCAGTTCACCTGATTCAGCATGGCCAGCTACTGCAACAGCTTCAAGGGCCAGTTCT
GCTGCCAGATCACCGAGTGATGAGCCTGATCCTGCTGCACCACCACCGATGTGCTGTGCTGTCGGTACTA
CACCAACTGCCTGCACTGATGACAGTGCGTGTCCCTGCTCCAGCACAACAAGGGCAGATGAGTGTGCACC
TGTACCGTGATCAGGTTCACCGGCTTCGAGATGGGATGAATCCCCTGAGAGTGATGGAACTGGTACTACC
TGTACAGAACCGGCACCACACTTTGAGTGTGCTACCGGCACACCTGACGGTCCTGAAGCGAGGTGTTCAT
CCTGTACTGACGGATCAAGCAGCCCAAGTGAAGATACGGCACCTGGTAGTTCAGCTGCCACAGCACCAGC
ACCTCTTGGTAGTGCAACAGAAGCGCCTGCCAGTTCAACTGTATCATCTTCCTGTGCTTTTGCTGCCGGT
GCTGCTGAAGCCTGCAGAGACTGTCTAGCTGATGGGGCACCACCAACCACTGACTGTGCTGAGATGTGGT
GTACACACACTGGTATTGGAGCGGCAACAACAGCTATACCGGCAGCCAGTACGGCAGCCGGATTCTGTGG
TGGTGTATCGTGCTGAGCGTGCTGCCTCTGCCTCACAGAAGCAGCAAGAGCTGAAGAATCCTGTGACTGA
AGCGCTGAGTGTGTACCAACACCTACAACCTGTGCTGATGACCCTGCGGCTTCTACACCTGAAAGCACAG
CCTGTACCGGCTGAGATACGTGGAACGGCTGTGGCTGTGACTGTAAAGCACCCCTAGAACACACGCCAGC
GTGTCCTGATGCACCATCGTGTTCAAGAGAGTGTGCGGCGTTAGCGCCGCCAGACTGACACCTTGTGGAA
CAGGCACCTCTACCGACGTGGTGTACCGGGCCTTCGACATCTACAACGATAAGGTGGCCGGCTTCGCCAA
GTTCCTGAAAACAAACTGCTGCCGCTTCCAAGAGAAGGACGAGGACGACAACCTGATCGACAGCTACTTC
GTGGTCAAGAGGCACACCTTCAGCAATTACCAGCACGAGGAAACCATCTACAATCTGCTGAAGGACTGCC
CTGCCGTGGCCAAGCACGACTTCTTCAAGTTCAGAATCGACGGCGACATGGTGCCTCACATCTCCAGACA
GCGGCTGACCAAGTACACCATGGCCGATCTGGTGTACGCCCTGAGACACTTCGACGAGGGCAACTGCGAC
ACCCTGAAAGAGATCCTGGTCACCTACAATTGCTGCGACGACGACTACTTCAACAAGAAGGATTGGTACG
ACTTCGTCGAGAACCCCGACATCCTGCGCGTGTACGCCAATCTGGGCGAGAGAGTTAGACAGGCACTGCT
GAAAACCGTGCAGTTCTGCGACGCCATGAGGAACGCCGGAATTGTGGGAGTGCTGACCCTGGACAACCAG
GACCTGAACGGCAATTGGTATGATTTCGGCGACTTCATCCAGACCACACCTGGCAGCGGAGTGCCTGTGG
TGGATAGCTACTACAGCCTGCTGATGCCCATCCTGACACTGACAAGAGCCCTGACAGCCGAGAGCCACGT
GGACACCGATCTGACCAAGCCTTACATCAAGTGGGACCTGCTGAAGTACGATTTCACCGAGGAAAGGCTG
AAACTGTTCGACCGGTACTTCAAGTACTGGGACCAGACCTACCATCCTAACTGCGTGAACTGCCTGGACG
ACCGGTGCATTCTGCACTGCGCCAACTTCAACGTGCTGTTCAGCACCGTGTTTCCACCTACCAGCTTCGG
ACCTCTCGTGCGCAAGATCTTCGTGGATGGCGTGCCCTTTGTGGTGTCCACCGGCTACCACTTCAGAGAA
CTGGGCGTCGTGCACAATCAGGACGTGAACCTGCACAGCAGCAGACTGAGCTTCAAAGAACTGCTGGTGT
ATGCCGCCGATCCTGCCATGCATGCCGCCTCTGGAAATCTGCTGCTCGACAAGCGGACCACCTGTTTCTC
TGTGGCCGCTCTGACCAACAACGTGGCCTTCCAGACAGTGAAGCCCGGCAATTTCAACAAGGACTTCTAC
GATTTCGCCGTGTCCAAGGGCTTCTTCAAAGAGGGCAGCAGCGTCGAGCTGAAGCACTTCTTCTTCGCCC
AAGACGGCAACGCCGCCATCTCCGACTACGATTACTACCGGTACAACCTGCCTACCATGTGCGACATCCG
GCAGCTGCTGTTCGTGGTGGAAGTGGTGGACAAGTACTTCGACTGCTACGACGGCGGCTGTATCAACGCC

AATCAAGTGATCGTGAACAACCTGGACAAGAGCGCCGGCTTTCCCTTCAACAAATGGGGCAAAGCCCGGC
TGTACTACGACAGCATGTCTTACGAGGATCAGGATGCCCTGTTCGCCTACACCAAGCGGAACGTGATCCC
CACCATCACACAGATGAACCTGAAGTACGCCATCAGCGCCAAGAACCGGGCCAGAACAGTTGCTGGCGTG
TCCATCTGTAGCACCATGACCAACCGGCAGTTCCACCAGAAGCTGCTGAAGTCTATCGCCGCCACAAGAG
GCGCCACAGTGGTCATCGGCACCAGCAAGTTTTATGGCGGCTGGCACAACATGCTGAAAACCGTGTACAG
CGACGTGGAAAACCCTCACCTGATGGGCTGGGACTACCCCAAGTGCGATAGAGCCATGCCTAACATGCTG
CGGATCATGGCCAGTCTGGTGCTGGCCAGAAAGCACACCACCTGTTGTAGCCTGAGCCACCGGTTTTACC
GGCTGGCCAATGAATGTGCCCAGGTGCTGAGCGAGATGGTCATGTGTGGCGGCAGCCTGTATGTGAAGCC
TGGCGGAACATCTAGCGGCGACGCCACAACAGCCTACGCCAACAGCGTGTTCAACATCTGCCAGGCCGTG
ACCGCCAATGTGAATGCCCTGCTGAGCACCGACGGCAACAAGATCGCCGACAAATACGTGCGGAACCTGC
AGCACAGACTGTACGAGTGCCTGTACCGGAACAGAGATGTGGACACCGACTTCGTGAACGAGTTCTACGC
CTACCTGCGGAAGCACTTCAGCATGATGATCCTGAGCGACGACGCCGTCGTGTGCTTCAATAGCACCTAC
GCCTCTCAAGGCCTGGTGGCCAGCATCAAGAACTTCAAGAGCGTGCTGTACTACCAGAACAACGTGTTCA
TGAGCGAGGCCAAGTGCTGGACCGAGACAGACCTGACAAAGGGCCCTCACGAGTTCTGCAGCCAGCACAC
CATGCTGGTTAAGCAGGGCGACGACTACGTGTACCTGCCTTATCCAGATCCTAGCCGGATCCTCGGAGCC
GGCTGCTTCGTGGATGACATCGTGAAAACCGATGGCACCCTGATGATCGAGAGATTCGTGTCCCTGGCTA
TCGACGCCTATCCTCTGACAAAGCACCCCAATCAAGAGTACGCCGACGTGTTCCACCTGTACCTGCAGTA
CATCCGGAAGCTGCACGATGAGCTGACCGGCCACATGCTGGACATGTACAGCGTGATGCTGACCAACGAC
AACACCAGCAGATACTGGGAGCCCGAGTTTTACGAGGCCATGTACACCCCTCACACCGTCCTGCAAGCTG
TGGGAGCTTGCGTGCTGTGCAATAGCCAGACCAGCCTGAGATGCGGGGCCTGTATCAGACGGCCTTTCCT
GTGTTGCAAGTGCTGCTACGACCACGTGATCAGCACCAGCCACAAACTGGTGCTGTCCGTGAATCCCTAC
GTGTGCAATGCCCCTGGCTGCGACGTGACCGATGTGACACAGCTGTATCTCGGCGGCATGAGCTACTACT
GCAAGAGCCACAAGCCTCCAATCAGCTTCCCTCTGTGCGCCAACGGACAGGTGTTCGGCCTGTACAAGAA
TACCTGCGTGGGCAGCGACAATGTGACCGACTTCAACGCCATTGCCACCTGTGACTGGACCAACGCCGGC
GATTACATCCTGGCCAATACCTGCACCGAGCGGCTGAAGCTGTTTGCCGCCGAAACACTGAAGGCCACCG
AGGAAACCTTCAAGCTGAGCTACGGAATCGCCACCGTGCGGGAAGTGCTGTCTGATAGAGAGCTGCACCT
GAGCTGGGAAGTGGGCAAGCCTAGACCTCCTCTGAACCGGAATTACGTGTTCACCGGCTACAGAGTGACC
AAGAACAGCAAGGTGCAGATCGGCGAGTACACCTTCGAGAAGGGCGATTATGGCGACGCTGTGGTGTACA
GAGGCACCACCACCTACAAGCTGAACGTGGGCGACTACTTCGTGCTGACAAGCCACACCGTGATGCCTCT
GTCTGCCCCTACACTGGTGCCCCAAGAACACTATGTGCGGATCACCGGACTGTACCCCACACTGAACATC
AGCGACGAGTTCAGCTCCAACGTGGCCAATTACCAGAAAGTGGGCATGCAGAAGTATAGCACACTGCAGG
GCCCACCTGGCACCGGCAAATCTCACTTTGCCATTGGACTGGCCCTGTACTATCCCAGCGCCAGAATCGT
GTACACCGCCTGTTCTCACGCCGCCGTTGATGCTCTGTGTGAAAAGGCCCTGAAGTATCTGCCCATCGAC
AAGTGCAGCCGGATCATCCCTGCCAGAGCCAGAGTGGAATGCTTCGACAAGTTCAAAGTGAACAGCACCC
TGGAACAGTACGTGTTCTGCACCGTGAACGCCCTGCCTGAAACCACCGCCGATATCGTGGTGTTCGACGA
GATCAGCATGGCCACCAACTACGACCTGAGCGTGGTCAACGCCAGACTGAGAGCCAAGCACTATGTGTAC
ATCGGCGACCCCGCTCAGCTGCCCGCTCCTAGAACACTGCTGACTAAGGGCACACTGGAACCCGAGTACT
TCAACTCCGTGTGCCGGCTGATGAAGACAATCGGCCCCGACATGTTCCTGGGCACCTGTAGAAGATGCCC
TGCCGAGATCGTGGATACCGTGTCTGCCCTGGTGTACGACAACAAACTGAAAGCCCACAAGGACAAGTCT
GCCCAGTGCTTCAAGATGTTCTACAAGGGCGTGATCACCCACGACGTGTCCAGCGCCATCAACAGACCTC
AGATCGGAGTTGTGCGCGAGTTCCTGACCAGAAATCCCGCTTGGCGGAAGGCCGTGTTTATCAGCCCTTA
CAACAGCCAGAACGCCGTGGCCTCCAAGATCCTGGGACTGCCTACACAGACCGTGGATAGCTCTCAGGGC
AGCGAGTACGATTACGTGATCTTCACCCAGACCACCGAGACTGCCCACAGCTGCAACGTGAACAGATTCA
ATGTGGCCATCACCAGGGCCAAAGTGGGAATCCTGTGCATCATGTCCGACCGGGACCTGTACGATAAGCT
GCAGTTCACCAGCCTGGAAATCCCCAGACGGAATGTGGCTACACTGCAGGCCGAGAATGTGACCGGCCTG
TTCAAGGACTGCAGCAAAGTGATCACCGGACTGCACCCTACACAGGCCCCTACACACCTGTCCGTGGACA
CCAAGTTTAAGACCGAGGGCCTGTGCGTCGACATCCCTGGCATCCCTAAGGACATGACCTACCGGCGGCT
GATCTCCATGATGGGCTTCAAGATGAACTACCAAGTGAACGGCTACCCCAACATGTTCATCACCAGAGAA
GAGGCCATCCGGCATGTGCGCGCCTGGATCGGATTTGATGTGGAAGGCTGTCACGCCACCAGGGAAGCCG
TGGGAACAAATCTGCCTCTGCAGCTGGGCTTTAGCACCGGCGTTAACCTGGTGGCTGTGCCTACCGGCTA
TGTGGACACCCCTAACAACACCGACTTCAGCCGGGTGTCCGCCAAACCTCCTCCAGGCGACCAGTTCAAG
CACCTGATTCCTCTGATGTACAAGGGCCTGCCTTGGAACGTCGTGCGGATCAAGATCGTGCAGATGCTGA
GCGACACCCTGAAGAACCTGTCCGACAGAGTGGTGTTCGTGCTGTGGGCCCATGGCTTTGAGCTGACCAG

CATGAAGTACTTTGTGAAGATCGGCCCCGAGCGGACCTGCTGCCTTTGTGATAGAAGGGCCACCTGTTTC
AGCACCGCCAGCGATACCTATGCCTGCTGGCACCACAGCATCGGCTTCGACTACGTGTACAACCCCTTCA
TGATCGACGTGCAGCAGTGGGGCTTCACCGGCAACCTGCAGAGCAACCACGATCTGTACTGTCAGGTGCA
CGGCAATGCCCACGTGGCCTCTTGTGATGCCATCATGACCAGATGCCTGGCCGTGCACGAGTGCTTCGTG
AAGAGAGTGGACTGGACCATCGAGTACCCCATCATCGGCGACGAGCTGAAGATCAACGCCGCCTGCAGAA
AGGTGCAGCACATGGTGGTTAAGGCCGCTCTGCTGGCCGACAAGTTTCCCGTGCTGCACGACATCGGCAA
CCCCAAGGCCATTAAGTGTGTGCCCCAGGCCGACGTGGAATGGAAGTTCTATGATGCCCAGCCTTGCAGC
GACAAGGCCTACAAGATCGAGGAACTGTTCTACAGCTACGCCACACACAGCGATAAGTTCACCGACGGCG
TGTGCCTGTTCTGGAACTGCAACGTGGACAGATACCCCGCCAACAGCATCGTGTGCAGATTCGACACCAG
AGTGCTGAGCAACCTGAACCTGCCTGGCTGTGATGGCGGCAGCCTGTACGTGAACAAGCACGCCTTTCAC
ACCCCTGCCTTCGACAAGAGCGCCTTCGTGAACCTGAAGCAGCTGCCATTCTTCTACTACAGCGACAGCC
CCTGCGAGAGCCACGGAAAGCAGGTCGTGTCCGACATCGATTACGTGCCCCTGAAGTCCGCCACCTGTAT
CACCAGATGCAATCTCGGCGGAGCCGTGTGTAGACACCACGCCAATGAGTACCGGCTGTACCTGGACGCC
TACAACATGATGATCAGCGCCGGCTTCAGCCTGTGGGTGTACAAGCAGTTCGATACCTACAACCTGTGGA
ACACCTTCACCAGACTGCAGAGCCTCGAGAACGTGGCCTTCAACGTGGTCAACAAGGGCCACTTCGATGG
CCAGCAAGGCGAGGTGCCAGTGTCCATCATCAACAATACCGTGTACACAAAGGTGGACGGCGTGGACGTG
GAACTGTTTGAGAACAAGACCACACTGCCCGTGAATGTGGCCTTTGAACTGTGGGCCAAGCGGAACATCA
AGCCCGTGCCTGAAGTGAAGATCCTGAACAACCTGGGAGTCGATATCGCCGCCAACACCGTGATCTGGGA
CTACAAGAGAGATGCCCCTGCTCACATCAGCACCATCGGCGTGTGTAGCATGACCGACATTGCCAAGAAG
CCCACCGAGACAATCTGCGCCCCTCTGACCGTGTTCTTCGACGGCAGAGTTGATGGCCAGGTGGACCTGT
TCAGAAACGCCAGAAACGGCGTGCTGATCACCGAGGGATCTGTGAAGGGACTCCAGCCTAGCGTGGGACC
TAAACAGGCCTCTCTGAATGGCGTGACCCTGATCGGAGAGGCCGTGAAAACCCAGTTCAACTACTACAAG
AAAGTGGACGGGGTTGTGCAGCAGCTCCCCGAGACATACTTCACCCAGAGCCGGAACCTGCAAGAGTTCA
AGCCTCGGAGCCAGATGGAAATCGACTTCCTGGAACTGGCCATGGACGAGTTCATCGAGCGGTACAAGCT
GGAAGGCTACGCCTTCGAGCACATCGTGTACGGCGATTTCAGCCACTCTCAGCTCGGCGGACTGCATCTG
CTGATTGGCCTGGCCAAGAGATTCAAAGAGAGCCCCTTCGAGCTGGAAGATTTCATCCCCATGGACAGCA
CCGTGAAGAACTACTTCATCACAGACGCCCAGACCGGCAGCTCTAAGTGCGTGTGTTCCGTGATCGATCT
GCTGCTGGACGACTTCGTGGAAATCATCAAGAGCCAGGACCTGTCTGTGGTGTCCAAGGTGGTCAAAGTG
ACCATCGACTACACCGAGATCAGCTTCATGCTCTGGTGCAAGGACGGACACGTGGAAACATTCTACCCCA
AGCTCCAGTCCTCTCAGGCTTGGCAACCTGGCGTGGCCATGCCTAACCTGTACAAGATGCAGCGGATGCT
GCTCGAGAAGTGCGACCTGCAGAACTACGGCGATAGCGCCACTCTGCCCAAGGGAATCATGATGAACGTG
GCAAAGTACACCCAGCTGTGCCAGTACCTGAACACCCTGACTCTGGCAGTGCCCTACAATATGAGAGTGA
TCCACTTCGGAGCCGGAAGCGACAAAGGCGTGGCACCTGGAACAGCTGTGCTGAGACAATGGCTGCCTAC
AGGCACCCTGCTGGTGGACAGCGACCTGAACGACTTTGTGTCCGACGCCGACAGCACACTGATCGGCGAT
TGTGCCACAGTGCACACCGCCAACAAGTGGGACCTGATCATCAGCGATATGTACGACCCCAAGACCAAGA
ACGTGACCAAAGAGAACGACAGCAAAGAGGGCTTCTTCACCTACATCTGCGGCTTCATCCAGCAGAAGCT
GGCCCTCGGAGGAAGCGTGGCCATCAAGATCACAGAGCACAGCTGGAACGCCGACCTGTACAAGCTGATG
GGCCACTTTGCCTGGTGGACCGCCTTCGTGACCAATGTGAATGCCAGCAGCAGCGAGGCCTTCCTGATCG
GCTGTAACTACCTGGGCAAGCCCAGAGAGCAGATCGACGGCTACGTGATGCACGCCAACTACATCTTTTG
GCGGAACACAAACCCCATCCAGCTGAGCAGCTACAGCCTGTTCGATATGAGCAAGTTCCCTCTGAAGCTG
CGGGGCACAGCCGTGATGTCTCTGAAAGAGGGCCAGATCAACGACATGATCCTGAGCCTGCTGAGCAAGG
GCAGACTGATCATCCGGGAAAACAACCGGGTCGTGATCAGCAGCGACGTGCTGGTCAACAACTGAACAAA
CAACGTGTGCTTCAGCTGCTTTATCGCCACCAGCCTGTGAAGCGTGTGCTGAAGCTACAACCAGAACAGC
ATCACCCCTTGCATCCACTGATTCTTCCACACCTGGTGCCTGCTGCCTTGACAGAGCTTCCAGATCCTGT
CCTTCACCTTCAACAGCGGCCTGGTGCTGACATTCCTGTTTCAGTGCTATCTGGTGCCCTGCTACACCTG
TCTGTGGGACCAGTGGTACTGAGAAGTGTGATGACCCTGTCCTACCATCTGATGATGGTGCCTGTTCTGC
TTCCACTGAGAGGTGTAACACAACAAGCGGCTGGACTTCTGGTACTACTTCAGATTCGAGGACCCCGTGC
CTACCTACTGCTGATGACGGTACTGATGCTGCTACTGATCCCTGTGAATCAGCATCCTGTGATGATCCAT
CTTCGGCTGTCTGCTGCCCCAGAAGCAGCAGAAACTGGACGGCAAATGAGTGCAGTCCCTGTTCTGATGC
GAGTGACTGCACTTCTGAATCTGCCTGAGCGCCTTCAGCTACGGCCCCTGAAGAAAGACCGGCTGATTTC
AGAAGTCCTGAGGCATCTGCGTCTGAGAGTACTGATGGCTGTTTTGAAACATCTTCTGAGCCCACGCCTA
CTGATTCAGCGCCTGATCTCCTAGCGGCTTCTTCGGCTTCAGAACCATCGGCAGATTCGCCAACCGGTAC
TGACACCACTGAGTGTCCAACTTCACCTGTTTCACCTGAAAGCTGTTCGACAGCTGGTAGTTCTTCTTCA

GACTGGACTCCTGGTGTTGCAGCCTGCTGTGTGGCCTGAGCAGCACCTGAGACTTCTCTATCAAGATCTA
GTGAAAGTGGAACCACTACCGGTGCTGCCGGCTGTGTACATGACCCAGCCTGCGGAACAAGGTGTACGTG
GAAATCCTGCACTGCCGGAAGCGGAACCTGAGCAACTTCTGACTGTGAAGCCCCACCAACCGGATCTACT
GTTGAATCAGCTAGTACTACAAGCTCGTGCCCTTCTGGTAGAGCTTCTGACGCCACCAGATCTGCATCTG
TCTGTGCCTGGAACAAGAGGAAAACCAGCAGCTCTGCTGCTGACTGTTCTGCCCCATCTGATTCCGGATC
ATCTTCCACTTTTGAGTGCTTTGGAGCGTGTCCTACTGAATCAAGTGATCCCTGCTGTATTGATGCCTGT
GCCGGTTCATCTGCAACTGAAGATGATGAAGCCAGACCAACAGAAGCCGGGCCAACTGGAAGGACTGTTG
ACTGTGACTGTAGATCACCCGCTGATTCTACCGGCTGCGGTACAGCCTGGAATTCTGACAGTCCTGATTC
TGAGGATGGTGGTAGCTGTGACTGCCCGTGTGAATCGTGTAGGAAGTGTAATCCCAGACCTTCTGAGAGC
GGTACTTCAACTGAAACCTGAGCGGCAGATGACACACCCTGTGATGGTGCTGAAGATTCTGACTGCTGCT
GTCCTTTACCATCATCTGGTTCCCCACACACTGATGGTGTTGGCTGCCCACCATCCAGAGCAGCAGCACA
TTCTTTTGAACCAGCACCTGTACCAGCAACTGCCTGTGGACCTGAAAGGTGTACTGATTCGGCTGAAAGC
AGATGTGCCAGTTCCAGCTCCAGTGGTTCAACAGACACCGGTGCAGCTACTGAGTGTGACAGAAGGTGTC
CGCCTTTCCAACCATCTGGCAGCGGCACTGCTGACACTACTGATGTTGCCCCTGATCCACCGACACCTGA
GATAGCTGACATTACACCATGTTCTTTTGGTGGTGCCAGTGCTACAATACCCGGAACAAGTACTTCTGAC
CCGGCTGCTGTAGCCTGAGCGGATGCTGACTGCACAGAAGCCCTTGCTGCTACAGCTGCAGAAGCACCTA
CAGCTACCTGGCCTGCCTGTTTTACCGCTTCTGATGCTTCAGCAACACCTGTCGGCTGTTCAACCGGGGC
TGAACCTGTCAGCAGCTGATCTAAGTGTGACACACTCACTGGTGCCGGTACATGAGATGACTGAGCGACA
GCGACTGATTCTCCAGCGCCGGCACCTGATGTAGCTGATCCATCCACCACTGCCTGCACTACGTGACCTG
GTGCAGAAAGTTCTCTTGTCTGCTGTGATGACTCTACTGTCACACCCACAAGTTCTACTACTAATGCTAC
CACCGGAACTCCACCAGCGTGTACGACCAGGACATCAGCAGACTGTACAACGTGCACCTGTGGTAGTTCA
ACTAGATGCAGCAGTCCTTCGTGGCTATCTGGCAGTTCCTGTACACCATCAAGCCCTGCTTCAACTGGAA
TAGCTGCTGAACCCGGCAGAAACACCCCAGATCATTCTGCACAAGCCAGACAAACCTGCAGAACACCACC
AACTGACGGTTCTGGTGGTTCTAGTTCTTCACGAACATCACCCGGTCCATCAAGACAAAGCAAGAAGTGA
TCTACTGACGGTCCACCTTCCAGCAGAGCGATACCTGCAGATGCTGGCTGCACCAGACCATCTGGTAGCT
GCCCTGGTAGTACTGCTGCTGAAGGCCTCACCTGTGCACCAAAGTGTGACGGCCTTACTGCTTCGCCACA
TTCGCCCACCGCTGAAACGACTGCTCTATCCACTTCTGCACCGTGTCCGGCTACAACCACTTCTGGCTGG
ACCTGTGGTGCCGGTGCTGCATCACCAACACCATCTGCTACGCCAACGGCCTCTGAGTGTGATGGTACTG
GTCCTACACCGAGTGCAGCCTCTGAGAGCCCAAGATCGACTGCCAGCCTATCTGATGATGCTACTGGCAG
AACAGCCGCCTGACATTCTTCCACAGCAAGTGCACCTGGAAAACCAGCAGATGCGGCCAGCCTAAGTGCA
CCAGCTTCAAGCACGCCTGCTGAACCACCTGACTGCAGTTCTGGTGCAATTTCAAGTGCTTCAAGTGATA
CCCCTTCACCAGCTGACAGAGCTGAGGCTGAAGCGCCAACTGATGAGTGGACCACAGACAGACCAGCAAG
TTCGCCGACATCTGCGACAGCACCATCAACTGAAGCTGCCGGAACCAGAGCTTCTGCTGATCCTGCTGCT
ACTGAAACGTCCGCGTGTGTACCTGGACAATCAAGAAAAGCTGATTCCTGTGGAAGGGCCTGAGCAGCTA
CGTGCTGCCTTCTGTGTCCACCTCCTGGTGTTCTCTGCTGGCCTGCGATCTGTGCCCCTGCACAAGAAAA
GAGCTGCACAATTGCAGCTGCCACCTGAGCTGATGGAAGTCCACACTGTCCAGCTGACGGTGCCTGTGCT
TTAAGTGGCACACCCTCGTGTGCAATACCAAAGAATTTCTGTGAACCACCAACCACTACTACCGGCAGCA
CATCTGCGTGTGGTAGCTGTGATGCTGCAACCGGAACTGCCAGCAGCACAGCCTGTGATCCTTCGCCACC
TGAATCCGGCTGATCCAAGGCGGCGTCAGATGAATCTTCTGAGAGTCCTACATCACCCGGTGCTGATTCA
GATGACACCTGTGGCACTGATGCTTTAGCTGCAAGCACAGCAAGCGGAACTGACCTCCTCAGTAGGGCTG
CCAAGAGTTCAAGTGAATCAGCCACAGAAGCCCCAGAACCTGGAAAGTCTGAGCCGTGTACAAGATGGCC
ATGGTGCATCTGGCCCGGTTCTACAGCTGGCTGGATTGCCACAGCAACGGCGACAATTACGCCCTGCTGT
ACGACCAGCTGCTGTGACTGTCTCAGGGCCTGCTGTTTCTGTGGATTCTGCTGCAGATCTGATGACGGCG
GCTCTGAGCCAGCGCTCAGAGATCTCAGATCACCCTGCACATCAACGAACTGATGGACCTGTTCATGCGG
ATCTTCACCATCGGCACCGTGACACTGAAGCAGGGCGAGATCAAGGACGCCACACCTAGCGATTTTGTGC
GGGCCACCGCCACAATTCCAATCCAGGCCTCTCTGCCTTTCGGCTGGCTGATTGTGGGAGTTGCACTGCT
GGCCGTGTTTCAGAGCGCCAGCAAGATCATCACCCTGAAGAAGAGATGGCAGCTGGCCCTGTCTAAGGGC
GTGCACTTCGTGTGTAACCTGCTGCTGCTGTTCGTGACCGTGTACAGCCATCTGCTGCTGGTGGCTGCTG
GACTGGAAGCCCCTTTCCTGTACCTGTACGCTCTGGTGTACTTCCTGCAGAGCATCAACTTCGTGCGGAT
CATCATGCGGCTGTGGCTGTGTTGGAAGTGCAGAAGCAAGAACCCTCTGCTGTATGACGCCAACTACTTC
CTGTGCTGGCACACCAATTGCTACGACTACTGCATCCCCTACAACAGCGTGACCAGCTCCATCGTGATCA
CAAGCGGCGACGGCACCACCTCTCCAATCAGCGAGCACGATTATCAGATCGGCGGCTACACAGAGAAGTG
GGAGAGCGGCGTGAAGGACTGTGTGGTGCTGCACAGCTACTTCACCTCCGACTACTACCAGCTGTACTCT

ACCCAGCTGAGCACCGACACAGGCGTGGAACACGTGACCTTTTTCATCTACAACAAGATCGTGGACGAGC
CCGAAGAACACGTGCAGATCCACACAATCGATGGCAGCAGCGGCGTGGTCAACCCCGTGATGGAACCCAT
CTACGACGAGCCTACCACCACCACAAGCGTGCCACTCTAAGCCCAGGCCGATGAGTACGAGCTGATGTAC
AGCTTCGTGTCCGAGGAAACCGGCACACTGATCGTGAACAGCGTCCTGCTGTTCCTGGCCTTCGTCGTGT
TTCTGCTCGTGACCCTGGCCATCCTGACCGCTCTGAGACTGTGCGCCTACTGCTGCAACATCGTGAACGT
GTCCCTGGTCAAGCCCAGCTTCTACGTGTACTCCAGAGTGAAGAACCTGAACAGCTCCAGGGTGCCCGAC
CTGCTGGTGTAAACAAACTGAATCCTGTACTGATTCTTCTGCCTCGAGCTGTGATTCTGACCTTGGCAGA
TCCCCACCGTGCTGCTGCCTCTGAAGTCCCTGAAGTCTAGCCTGAACAACGGCACCTGATGATGAGTGTC
CTACAGCCTGCACGGCTTCGTGTTCTACAACCTGCCTATGCCTACCGGCATCGGATTCTGCATCTGACTG
TCCTGATTCAGCTCCGGCTGCTACGGCCAGTGACTGTAACTGGTGCTGTGCCTGCTCCTGTTCACCGAGT
GAATCGGCAGCCCTGTGGAACTGCTGAGCCAGTGGCTGGTGCTTTGAGCCTGATGTGGCTCTGCCACAAG
CCTGCTGCTCAGCGATTGCCTGAGAGTGCGAGTGCCTTGTGGCCACTCTATCCAGAAGCTGACCTTCTTC
AGCACCTGTCACTCTATGGCCCTGTTCTGACCCGACCGGTTCTGAAAAGTGAACAGCTGATCTGAGCTGT
GAAGCTTCGTGGACATCTTCGTGCTGCTCGACACAATCTGAGATGCCGTGACCAGCAGAACCTGCCTGAA
GAAGTCCCTGCTGCTGCACCACGAGCGGTTTCTGATCACCAACTGGGAGCTGCGGAGCGTGTGACAAGTG
ACACAGGTCCTGCTGCACACAGTGGCCACAGGACTGGCCACCATCAACTGAACCCAGACAATCCCTGTGG
CCGTGACCATCCTGCTGTGCCTGTACAGCAAGTGACAGCAGATGTTCCACCTGGTGGACTTCCAAGTGAC
AATCGCCGAGATCCTGCTGATCATCATGCGGACCTTCAAGGTGTCCATCTGGAACCTGGACTACATCATC
AACCTCATCATCAAGAACCTGAGCAAGAGCCTGACCGAGAACAAGTACAGCCAGCTGGACGAGGAACAGC
CCATGGAAATCGACTGAACGAATATGAAGATCATCCTGTTTCTGGCCCTGATCACCCTGGCCACATGCGA
GCTGTACCACTACCAAGAGTGTGTGCGGGGCACCACCGTGCTGCTGAAAGAGCCTTGTAGCAGCGGCACC
TACGAGGGCAACAGCCCTTTTCATCCCCTGGCCGACAACAAGTTCGCCCTGACCTGTTTCAGCACCCAGT
TCGCCTTCGCCTGTCCTGATGGCGTGAAGCACGTGTACCAGCTGAGAGCCAGAAGCGTGTCCCCTAAGCT
GTTCATCCGGCAAGAGGAAGTGCAAGAGCTGTACTCCCCTATCTTCCTGATCGTGGCCGCCATCGTGTTC
ATCACCCTGTGCTTCACCCTGAAGAGAAAGACCGAGTGACTGAACTTCCACTGACTGACCAGCATCTGCG
CCTTTTGACCCTTCTGCTACAGCCTGTTCTGACTGTGCCTGCTGAGCTTCGGCAGCCACCTGAACTGCAA
GATCATTATGAAGCTGGTCACCCCTAAGCGGACCTGAAACTTCCTGTTCTCCTGAGAGTCTAGCCAGCTG
TGACTGCACTTCACCAAGAACGTGGTGTACTCCCACGTGCTGAACATCAACCACATGTGACTGATGACCC
GGGTGCTGTTCACCAGCATCCTGAACGGCATCCTGGAATGAGAGCTGGAAAACCAGCACCTGTGACTGAA
TTGCGCCTGGATGCGGCTGGTGCTGAATCACCCTTTTAGCACCTCCATCAGCGTGATCATTCAGTTCCCC
GTGTACCTGCTGCAGCTGATCGCCCGGAATCTGAACTGGGTCGTGCTGTGATGTGTCGTGCGGAGCATGA
AGACCTTTTGATCCATCATGACCTTCGTGCTGTTCTGAATCAGCAGCAAGCGGACCAACTGAAACGTGTG
ATGATGGACCCCTAAGAGCGCCAAGTGCACCCCTCACTACGTGTGGTGGACCCTGAGATTCAACTGGCAG
TGACCTGAGTGGCGGACACAGTGGGGAGCCATCAAGACAACAAGCGCCCCTCGGTTCACCCAGTGATACT
GCGTGCTGGTGCACAGAAGCCACAGCACTTGGCAGGGCAGACCTTGAATCCCCAGCCGGACCAGAAGATC
CAACTGACACCAGTGACAGAGCAGATGACCCAACTGGCTGCTGCCCAAGAGCTACCAGACCAACAGCTGG
TGGTAGCGCTGAAACGAGCGGAGCCAGAGCAAGATGGTGTTCCTGCTGCCTCGGAACTGGGCCAGATCTT
GGACAAGCCTGTGGTGCTGACAGCGGAGACACCACATGGGCTGCAATTGAGGCAGCCTCGAGTACACAAA
GAGAAGCCACTGGCACCCTCAGAGCTGCTGACAGTGCTGCAATAGAGCCACCACCAGCAGCCGGAACAAT
ATCGCCAAGCGGCTGCTGCGGAGAAGAGAGCAGAGAAGGCAGTCCAGCCTGTTTAGCTTCCTGATTACCT
GAAGCCAGCAGTTCAAGAAGTTCAACAGCCGGCAGCAGTAGGGCAACTTCTCCTGCTGAAATGGCTGGCA
GTGGCGCTGATGCTGCTCCTGTTTTGCTGCCGCCTGACAGATCGAGCCTGCCTGAGAGCAGAATGTGTGG
TAGAGGCCCACCACCACAAGGCCCAATTGCCACTGAGAGATCTGCTGCTGAGGCTTTTGAGAGGCCAGCG
CCAAGACCTACTGCCACTGATCCATCCAGTGCAACACCAGCTTCCGGCAGACCTGGTCCAGAACAAACCC
CAGAAAGTTTTGGGGCCCTGGCACCAACCAGACCAGAAACTGACTGCAGACCCTGGCCGCCAATTGCACC
ATCTGTCCCCAGAGATTCAGCGTGCTGAGAAACGTGGCCCACTGGCATGGCAGCCACACCTTTGGCAATG
TGGTGGATCTGCACAGGTGCCACCAGATCGGCTGACAGAGAAGCAAGTTCCAGCGGAGCAGCCACTTCGC
CGAATGAGCCTACTGACGGATCCAGAACATCCCCACCAACCGGGCCTGAAAAGGCCAGAAAGAAGAGGGC
TGATAGAACAGCTCCCTGACCGCCGAGACAGAGGAAACCGCCAACTGCGATAGCTCCAGCTGCTGCAGAT
TCGGCTGATTCCTGCAGACCATTGCCACCATCCACGAGCAGTGCTGACTGAACAGCGGCCTGAACAGCTG
CAGACCTCACAAGGCCGATGGCCTGTACAAGCGGTTCCGGTTCAGCGTGTACGACATCTGAAGCACCCTG
GTGCAGAACGAGTTCTCTTGACTGCACAGCACAAGCCGGTGCAGCTGACTGTGAAGCCATATCGCCATCT
TCAACCAGTGCGTGACACTGGGGAGAACCTGAAAGTCCCACCACATCTTCACCGAGGCCACCAGGTCCAC

CATCGAGTGCACAGTGAACAACGCCAGAGAGAGCTGCCTGTACGGCAGAGCCCTGATGTGCAAGATTAAC
TTCAGCAGCGCTATCCCCATGTGATTCTGATGACTGCTGCGGCGGATGACCAAGAAGAAGAAAAAGAAAA
AGAAGAAGAAGAA

**5′ UTR**

WT

ATTAAAGGTTTATACCTTCCCAGGTAACAAACCAACCAACTTTCGATCTCTTGTAGATCTGTTCTCTAAACGAACTT
TAAAATCTGTGTGGCTGTCACTCGGCTGCATGCTTAGTGCACTCACGCAGTATAATTAATAACTAATTACTGTCGTT
GACAGGACACGAGTAACTCGTCTATCTTCTGCAGGCTGCTTACGGTTTCGTCCGTGTTGCAGCCGATCATCAGCACA
TCTAGGTTTCGTCCGGGTGTGACCGAAAGGTAAG

VT1

ATCAAGGGCCTGTACCTGCCCCGCTAACAGACCAACCAGCTGAGCATCAGCTGCCGCAGCGTGCTGTAAACCAACTT
CAAGATCTGCGTGGCCGTGACCCGCCTGCACGCCTAATGCACCCACGCCGTGTAACTGATCACCAACTACTGCCGCT
AACAGGACACCAGCAACAGCAGCATCTTCTGCCGCCTGCTGACCGTGAGCAGCGTGCTGCAGCCCATCATCAGCACC
AGCCGCTTCCGCCCCGGCGTGACCGAGCGCTAAG

VT2

ATTAAAGGACTGTACCTGCCCCGGTGACAGACCAACCAGCTGTCAATCAGCTGTCGGTCAGTGCTGTGAACAAACTT
TAAAATCTGCGTCGCCGTGACCAGGCTGCATGCCTGATGCACCCATGCCGTCTGACTCATCACCAACTACTGCAGAT
GACAGGATACCAGCAACAGCTCCATCTTTTGCCGGCTGCTGACCGTGAGCTCCGTGCTGCAGCCTATCATCTCAACC
AGCCGGTTCAGACCTGGCGTGACTGAGAGATGAG

VT3

ATTAAAGGCCTCTATTTGCCACGATAGCAAACGAACCAACTCTCAATTTCTTGTAGGTCCGTACTTTAAACCAACTT
CAAGATTTGCGTCGCTGTGACCAGACTCCATGCGTAATGCACCCATGCTGTGTGACTTATTACTAACTATTGCCGAT
GACAAGACACCTCTAACAGTAGCATCTTCTGTCGACTCCTGACGGTGTCTAGTGTTTTGCAACCCATCATTTCTACT
AGCAGATTTCGACCCGGTGTAACTGAGAGGTAGG

VT4

ATCAAGGGCCTGTACCTGCCTAGATGACAGACCAACCAGCTGAGCATCTCCTGCAGATCCGTGCTGTGAACAAACTT
CAAGATCTGCGTGGCCGTGACCAGACTGCACGCCTGATGTACACATGCCGTGTGACTGATCACCAACTACTGCAGAT
GACAGGACACCAGCAACAGCAGCATCTTCTGCCGGCTGCTGACCGTGTCATCTGTGCTGCAGCCTATCATCAGCACC
AGCAGATTCAGACCCGGCGTGACCGAGAGATGAG

**3′ UTR**

WT

TAAACGTTTTCGCTTTTCCGTTTACGATATATAGTCTACTCTTGTGCAGAATGAATTCTCGTAACTACATAGCACAA
GTAGATGTAGTTAACTTTAATCTCACATAGCAATCTTTAATCAGTGTGTAACATTAGGGAGGACTTGAAAGAGCCAC
CACATTTTCACCGAGGCCACGCGGAGTACGATCGAGTGTACAGTGAACAATGCTAGGGAGAGCTGCCTATATGGAAG
AGCCCTAATGTGTAAAATTAATTTTAGTAGTGCTATCCCCATGTGATTTTAATAGCTTCTTAGGAGAATGACAAAAA
AAAAAAAAAAAAAAAAAAAAAAAAAAAAAA

VT1

ACAAGCGCTTCCGCTTCAGCGTGTACGACATCTAAAGCACCCTGGTGCAGAACGAGTTCAGCTAACTGCACAGCACC
AGCCGCTGCAGCTAACTGTAAAGCCACATCGCCATCTTCAACCAGTGCGTGACCCTGGGCCGCACCTAAAAGAGCCA
CCACATCTTCACCGAGGCCACCCGCAGCACCATCGAGTGCACCGTGAACAACGCCCGCGAGAGCTGCCTGTACGGCC
GCGCCCTGATGTGCAAGATCAACTTCAGCAGCGCCATCCCCATGTAATTCTAATAACTGCTGCGCCGCATGACCAAG

AAGAAGAAGAAGAAGAAGAAGAAGAAGAAA

VT2

ATAAGCGGTTCAGATTCAGCGTGTACGACATTTGAAGCACCCTGGTGCAGAACGAGTTTAGCTGACTGCACTCCACC
TCTCGCTGTAGCTGACTCTGATCCCATATTGCCATCTTTAATCAGTGCGTCACCCTGGGCCGGACATGAAAGTCTCA
TCACATCTTTACTGAGGCTACTAGATCTACCATCGAATGTACCGTGAACAATGCCAGGGAGAGTTGCCTGTACGGCC
GGGCTCTGATGTGCAAGATCAACTTTAGCAGCGCCATCCCCATGTGATTCTGATAGCTGCTGCGGAGGATGACCAAG
AAAAAGAAAAAGAAAAAGAAGAAAAAGAA

VT3

ATAAGAGATTTAGGTTCTCAGTTTACGACATATGATCCACCCTCGTACAGAACGAGTTCTCCTAACTGCACAGTACG
AGCCGCTGTTCCTAGCTGTGATCCCATATTGCCATCTTTAACCAATGTGTGACGCTCGGCCGGACGTAGAAGAGTCA
TCATATATTCACCGAGGCGACGCGGTCCACGATCGAATGCACTGTGAACAATGCCCGCGAGAGTTGCCTCTATGGAA
GAGCGTTGATGTGCAAGATTAACTTTTCATCCGCCATTCCGATGTGATTCTAATGATTGCTCCGCCGAATGACAAAG
AAAAAGAAAAAGAAGAAGAAAAAAAAAGAAA

VT4

ACAAGCGGTTCCGGTTCAGCGTGTACGACATCTGAAGCACCCTGGTGCAGAACGAGTTCTCTTGACTGCACAGCACA
AGCCGGTGCAGCTGACTGTGAAGCCATATCGCCATCTTCAACCAGTGCGTGACACTGGGGAGAACCTGAAAGTCCCA
CCACATCTTCACCGAGGCCACCAGGTCCACCATCGAGTGCACAGTGAACAACGCCAGAGAGAGCTGCCTGTACGGCA
GAGCCCTGATGTGCAAGATTAACTTCAGCAGCGCTATCCCCATGTGATTCTGATGACTGCTGCGGCGGATGACCAAG
AAGAAGAAAAAGAAAAAGAAGAAGAAAAA

**Number of RBP motif binding positions in the reference sequence (WT) and the variants**

| | | | WHOLE GENOME | | | |
|---|---|---|---|---|---|---|
| S/N | RBP | WT | VT1 | VT2 | VT3 | VT4 |
| 1 | A1CF | 705 | 141 | 74 | 309 | 180 |
| 2 | ANKHD1 | 46 | 23 | 28 | 65 | 37 |
| 3 | BOLL | 790 | 97 | 314 | 589 | 181 |
| 4 | BRUNOL4 | 213 | 10 | 215 | 214 | 135 |
| 5 | BRUNOL5 | 252 | 23 | 256 | 243 | 180 |
| 6 | BRUNOL6 | 76 | 32 | 138 | 82 | 101 |
| 7 | CELF1 | 549 | 21 | 199 | 345 | 88 |
| 8 | CNOT4 | 238 | 25 | 208 | 157 | 254 |
| 9 | CPEB1 | 695 | 20 | 159 | 531 | 32 |
| 10 | CPEB2 | 275 | 0 | 101 | 259 | 39 |
| 11 | CPEB4 | 495 | 0 | 161 | 435 | 46 |
| 12 | DAZ3 | 807 | 10 | 209 | 697 | 104 |
| 13 | DAZAP | 322 | 2 | 28 | 261 | 5 |
| 14 | EIF4G2 | 115 | 760 | 225 | 204 | 271 |
| 15 | ELAVL4 | 856 | 120 | 146 | 32 | 26 |
| 16 | ENOX1 | 29 | 22 | 146 | 32 | 26 |
| 17 | ESRP1 | 2 | 12 | 44 | 7 | 1 |
| 18 | ESRP2 | 2 | 0 | 13 | 22 | 1 |
| 19 | EWSR1 | 32 | 58 | 109 | 35 | 109 |
| 20 | FMR1 | 25 | 27 | 40 | 13 | 40 |
| 21 | FUBP1 | 554 | 26 | 40 | 40 | 40 |
| 22 | FUBP3 | 1081 | 59 | 59 | 723 | 35 |
| 23 | FUS | 10 | 59 | 34 | 48 | 21 |
| 24 | FXR1 | 85 | 86 | 95 | 79 | 108 |
| 25 | FXR2 | 4 | 17 | 25 | 12 | 25 |
| 26 | G3BP2 | 34 | 9 | 42 | 30 | 24 |
| 27 | HNRNPA0 | 334 | 0 | 32 | 249 | 6 |
| 28 | HNRNPA1 | 179 | 37 | 68 | 157 | 62 |
| 29 | HNRNPA1L2 | 30 | 0 | 7 | 21 | 3 |
| 30 | HNRNPA2B1 | 138 | 18 | 105 | 166 | 56 |
| 31 | HNRNPC | 1020 | 0 | 234 | 791 | 61 |
| 32 | HNRNPCL1 | 1068 | 0 | 254 | 834 | 61 |
| 33 | HNRNPD | 370 | 12 | 42 | 193 | 2 |
| 34 | HNRNPDL | 1249 | 352 | 191 | 726 | 54 |
| 35 | HNRNPF | 136 | 23 | 138 | 142 | 70 |
| 36 | HNRNPH1 | 33 | 32 | 40 | 32 | 36 |
| 37 | HNRNPH2 | 5 | 0 | 42 | 23 | 3 |
| 38 | HNRNPK | 62 | 599 | 243 | 120 | 232 |

| | | | | | | |
|---|---|---|---|---|---|---|
| 39 | HNRNPL | 708 | 184 | 243 | 399 | 444 |
| 40 | HNRNPM | 89 | 1 | 26 | 69 | 15 |
| 41 | HNRNPU | 126 | 26 | 95 | 120 | 70 |
| 42 | HNRPLL | 198 | 50 | 130 | 118 | 140 |
| 43 | HuR | 372 | 69 | 69 | 263 | 9 |
| 44 | IGF2BP1 | 236 | 13 | 78 | 136 | 63 |
| 45 | IGF2BP2 | 276 | 51 | 115 | 128 | 100 |
| 46 | IGF2BP3 | 163 | 18 | 29 | 87 | 38 |
| 47 | ILF2 | 10 | 9 | 29 | 43 | 15 |
| 48 | KHDRBS1 | 200 | 41 | 22 | 107 | 4 |
| 49 | KHDRBS2 | 341 | 159 | 33 | 170 | 3 |
| 50 | KHDRBS3 | 498 | 88 | 54 | 239 | 1 |
| 51 | KHSRP | 499 | 0 | 149 | 369 | 59 |
| 52 | LIN28A | 23 | 22 | 42 | 15 | 15 |
| 53 | MATR3 | 150 | 43 | 42 | 85 | 40 |
| 54 | MBNL1 | 453 | 1397 | 445 | 356 | 503 |
| 55 | MSI1 | 200 | 2 | 21 | 216 | 20 |
| 56 | NOVA1 | 165 | 146 | 151 | 155 | 122 |
| 57 | NUPL2 | 378 | 97 | 99 | 220 | 71 |
| 58 | PABPC1 | 206 | 14 | 53 | 122 | 30 |
| 59 | PABPC3 | 211 | 99 | 84 | 135 | 88 |
| 60 | PABPC4 | 359 | 36 | 77 | 214 | 48 |
| 61 | PABPC5 | 154 | 2 | 43 | 69 | 18 |
| 62 | PABPN1 | 178 | 26 | 61 | 68 | 58 |
| 63 | PABPN1L | 231 | 33 | 56 | 130 | 31 |
| 64 | PCBP1 | 22 | 259 | 85 | 51 | 45 |
| 65 | PCBP2 | 79 | 484 | 186 | 131 | 173 |
| 66 | PCBP3 | 67 | 2 | 37 | 81 | 24 |
| 67 | PCBP4 | 7 | 65 | 72 | 49 | 61 |
| 68 | PRR3 | 889 | 99 | 197 | 488 | 78 |
| 69 | PTB3 | 111 | 3 | 33 | 148 | 12 |
| 70 | PTBP3 | 115 | 3 | 36 | 150 | 12 |
| 71 | PUF60 | 271 | 0 | 104 | 356 | 32 |
| 72 | PUM1 | 150 | 43 | 42 | 85 | 69 |
| 73 | PUM2 | 259 | 13 | 57 | 164 | 23 |
| 74 | QKI | 178 | 197 | 51 | 109 | 21 |
| 75 | RALY | 473 | 0 | 117 | 371 | 34 |
| 76 | RBFOX1 | 29 | 95 | 75 | 38 | 53 |
| 77 | RBFOX2 | 17 | 105 | 61 | 40 | 64 |
| 78 | RBFOX3 | 17 | 105 | 61 | 40 | 64 |
| 79 | RBM15B | 791 | 25 | 144 | 482 | 8 |

| | | | | | | |
|---|---|---|---|---|---|---|
| 80 | RBM22 | 13 | 223 | 147 | 36 | 196 |
| 81 | RBM23 | 3 | 214 | 59 | 32 | 22 |
| 82 | RBM24 | 439 | 118 | 481 | 329 | 320 |
| 83 | RBM25 | 2 | 0 | 21 | 14 | 0 |
| 84 | RBM28 | 32 | 12 | 38 | 39 | 28 |
| 85 | RBM3 | 33 | 18 | 25 | 46 | 16 |
| 86 | RBM38 | 121 | 54 | 202 | 123 | 171 |
| 87 | RBM4 | 16 | 264 | 40 | 42 | 42 |
| 88 | RBM41 | 707 | 105 | 141 | 408 | 101 |
| 89 | RBM42 | 209 | 323 | 80 | 110 | 63 |
| 90 | RBM45 | 367 | 1365 | 634 | 473 | 951 |
| 91 | RBM46 | 172 | 18 | 46 | 73 | 19 |
| 92 | RBM47 | 420 | 32 | 144 | 243 | 50 |
| 93 | RBM4B | 4 | 67 | 28 | 39 | 20 |
| 94 | RBM5 | 28 | 9 | 39 | 26 | 8 |
| 95 | RBM6 | 126 | 1067 | 277 | 159 | 505 |
| 96 | RBM8A | 6 | 164 | 15 | 26 | 13 |
| 97 | RBMS1 | 224 | 6 | 43 | 207 | 17 |
| 98 | RBMS2 | 244 | 1 | 24 | 193 | 6 |
| 99 | RBMS3 | 601 | 11 | 103 | 522 | 17 |
| 100 | RC3H1 | 361 | 19 | 107 | 248 | 62 |
| 101 | SAMD4A | 3 | 56 | 43 | 8 | 41 |
| 102 | SART3 | 212 | 14 | 54 | 128 | 30 |
| 103 | SF1 | 468 | 384 | 77 | 361 | 21 |
| 104 | SFPQ | 515 | 225 | 275 | 368 | 165 |
| 105 | SNRNP70 | 89 | 72 | 51 | 44 | 83 |
| 106 | SNRPA | 140 | 284 | 231 | 98 | 288 |
| 107 | SRSF1 | 13 | 37 | 53 | 17 | 12 |
| 108 | SRSF10 | 93 | 368 | 161 | 94 | 221 |
| 109 | SRSF11 | 8 | 10 | 23 | 10 | 7 |
| 110 | SRSF2 | 56 | 520 | 211 | 61 | 234 |
| 111 | SRSF4 | 48 | 454 | 131 | 39 | 214 |
| 112 | SRSF5 | 22 | 1293 | 226 | 45 | 333 |
| 113 | SRSF7 | 82 | 484 | 438 | 190 | 372 |
| 114 | SRSF8 | 18 | 588 | 141 | 32 | 176 |
| 115 | SRSF9 | 41 | 63 | 95 | 81 | 59 |
| 116 | TAF15 | 46 | 10 | 59 | 58 | 17 |
| 117 | TARDBP | 80 | 36 | 139 | 86 | 122 |
| 118 | TIA1 | 740 | 0 | 216 | 620 | 58 |
| 119 | TRA2A | 82 | 18 | 48 | 35 | 84 |
| 120 | TRNAU1AP | 694 | 0 | 127 | 433 | 11 |

| | | | | | | |
|---|---|---|---|---|---|---|
| 121 | TUT1 | 112 | 26 | 41 | 114 | 33 |
| 122 | U2AF2 | 293 | 5 | 122 | 291 | 18 |
| 123 | UNK | 284 | 2 | 57 | 233 | 37 |
| 124 | YBX1 | 152 | 534 | 144 | 90 | 297 |
| 125 | YBX2 | 183 | 570 | 154 | 90 | 299 |
| 126 | ZC3H10 | 5 | 303 | 35 | 8 | 68 |
| 127 | ZC3H14 | 240 | 0 | 81 | 276 | 14 |
| 128 | ZCRB1 | 767 | 85 | 80 | 391 | 7 |
| 129 | ZFP36 | 684 | 225 | 108 | 474 | 12 |
| 130 | ZNF326 | 308 | 8 | 98 | 272 | 44 |
| 131 | ZNF638 | 216 | 35 | 67 | 178 | 70 |

| | | | 5' UNTRANSLATED REGION | | | |
|---|---|---|---|---|---|---|
| S/N | RBPs | WT | JCat | VB | IDT | GA |
| 1 | A1CF | 8 | 0 | 0 | 2 | 0 |
| 2 | ANKHD1 | 0 | 0 | 0 | 1 | 0 |
| 3 | BOLL | 0 | 0 | 0 | 1 | 0 |
| 4 | BRUNOL4 | 2 | 0 | 0 | 5 | 0 |
| 5 | BRUNOL5 | 3 | 0 | 0 | 5 | 3 |
| 6 | BRUNOL6 | 1 | 0 | 0 | 0 | 1 |
| 7 | CELF1 | 3 | 0 | 0 | 3 | 0 |
| 8 | CNOT4 | 0 | 0 | 0 | 0 | 3 |
| 9 | CPEB1 | 0 | 0 | 0 | 1 | 0 |
| 10 | CPEB2 | 0 | 0 | 3 | 0 | 0 |
| 11 | CPEB4 | 0 | 0 | 2 | 2 | 0 |
| 12 | DAZ3 | 4 | 0 | 0 | 2 | 0 |
| 13 | DAZAP | 0 | 0 | 0 | 0 | 0 |
| 14 | EIF4G2 | 3 | 10 | 6 | 0 | 4 |
| 15 | ELAVL4 | 4 | 0 | 0 | 2 | 0 |
| 16 | ENOX1 | 0 | 0 | 0 | 2 | 0 |
| 17 | ESRP1 | 0 | 0 | 0 | 0 | 0 |
| 18 | ESRP2 | 0 | 0 | 0 | 0 | 0 |
| 19 | EWSR1 | 2 | 0 | 0 | 0 | 0 |
| 20 | FMR1 | 2 | 0 | 0 | 0 | 1 |
| 21 | FUBP1 | 1 | 0 | 0 | 3 | 0 |
| 22 | FUBP3 | 8 | 0 | 0 | 1 | 0 |
| 23 | FUS | 0 | 6 | 0 | 0 | 0 |
| 24 | FXR1 | 3 | 0 | 2 | 1 | 3 |
| 25 | FXR2 | 0 | 0 | 0 | 0 | 0 |
| 26 | G3BP2 | 0 | 0 | 0 | 0 | 0 |
| 27 | HNRNPA0 | 1 | 0 | 0 | 0 | 0 |
| 28 | HNRNPA1 | 0 | 0 | 0 | 0 | 0 |
| 29 | HNRNPA1L2 | 0 | 0 | 0 | 0 | 0 |
| 30 | HNRNPA2B1 | 0 | 0 | 0 | 0 | 0 |
| 31 | HRNPC | 0 | 0 | 1 | 6 | 0 |
| 32 | HNRNPCL1 | 0 | 0 | 1 | 4 | 0 |
| 33 | HNRNPD | 4 | 0 | 0 | 3 | 0 |
| 34 | HNRNPDL | 7 | 1 | 1 | 0 | 0 |
| 35 | HNRNPF | 0 | 0 | 0 | 0 | 0 |
| 36 | HNRNPH1 | 0 | 0 | 0 | 0 | 0 |
| 37 | HNRNPH2 | 0 | 0 | 0 | 0 | 0 |
| 38 | HNRNPK | 6 | 6 | 5 | 3 | 1 |

| | | | | | | |
|---|---|---|---|---|---|---|
| 39 | HNRNPL | 1 | 2 | 5 | 2 | 3 |
| 40 | HNRNPM | 0 | 0 | 0 | 0 | 0 |
| 41 | HNRNPU | 0 | 0 | 0 | 3 | 0 |
| 42 | HNRPLL | 0 | 1 | 1 | 0 | 0 |
| 43 | HUR | 0 | 0 | 0 | 0 | 0 |
| 44 | IGF2BP1 | 0 | 0 | 0 | 0 | 0 |
| 45 | IGF2BP2 | 3 | 0 | 0 | 0 | 0 |
| 46 | IGF2BP3 | 0 | 0 | 0 | 0 | 0 |
| 47 | ILF2 | 0 | 0 | 0 | 0 | 0 |
| 48 | KHDRBS1 | 1 | 0 | 0 | 0 | 0 |
| 49 | KHDRBS2 | 5 | 0 | 0 | 0 | 0 |
| 50 | KHDRBS3 | 4 | 0 | 0 | 0 | 0 |
| 51 | KHSRP | 0 | 0 | 0 | 1 | 0 |
| 52 | LIN28A | 0 | 0 | 0 | 0 | 0 |
| 53 | MATR3 | 2 | 0 | 0 | 0 | 0 |
| 54 | MBNL1 | 1 | 16 | 7 | 3 | 5 |
| 55 | MSI1 | 0 | 0 | 0 | 0 | 0 |
| 56 | NOVA1 | 6 | 3 | 1 | 0 | 4 |
| 57 | NUPL2 | 0 | 0 | 0 | 0 | 0 |
| 58 | PABPC1 | 0 | 0 | 0 | 0 | 0 |
| 59 | PABPC3 | 0 | 0 | 1 | 0 | 0 |
| 60 | PABPC4 | 0 | 0 | 0 | 0 | 0 |
| 61 | PABPC5 | 0 | 0 | 0 | 0 | 0 |
| 62 | PABPN1 | 0 | 0 | 0 | 0 | 0 |
| 63 | PABPN1L | 0 | 0 | 1 | 0 | 0 |
| 64 | PCBP1 | 0 | 0 | 0 | 0 | 0 |
| 65 | PCBP2 | 3 | 0 | 0 | 0 | 0 |
| 66 | PCBP3 | 0 | 0 | 0 | 0 | 0 |
| 67 | PCBP4 | 0 | 0 | 1 | 0 | 0 |
| 68 | PRR3 | 8 | 0 | 0 | 4 | 0 |
| 69 | PTB3 | 5 | 0 | 0 | 1 | 0 |
| 70 | PTBP3 | 5 | 0 | 0 | 1 | 0 |
| 71 | PUF60 | 4 | 0 | 0 | 1 | 0 |
| 72 | PUM1 | 10 | 0 | 0 | 0 | 0 |
| 73 | PUM2 | 0 | 0 | 1 | 0 | 1 |
| 74 | QKI | 7 | 2 | 1 | 2 | 1 |
| 75 | RALY | 0 | 0 | 0 | 0 | 0 |
| 76 | RBFOX1 | 0 | 4 | 3 | 0 | 3 |
| 77 | RBFOX2 | 0 | 3 | 3 | 0 | 4 |
| 78 | RBFOX3 | 0 | 3 | 3 | 0 | 4 |
| 79 | RBM15B | 3 | 0 | 1 | 3 | 0 |

| | | | | | | |
|---|---|---|---|---|---|---|
| 80 | RBM22 | 2 | 3 | 6 | 0 | 6 |
| 81 | RBM23 | 0 | 2 | 0 | 0 | 0 |
| 82 | RBM24 | 3 | 0 | 1 | 1 | 0 |
| 83 | RBM25 | 0 | 0 | 0 | 0 | 0 |
| 84 | RBM28 | 0 | 0 | 0 | 0 | 0 |
| 85 | RBM3 | 0 | 0 | 1 | 1 | 0 |
| 86 | RBM38 | 3 | 0 | 0 | 0 | 0 |
| 87 | RBM4 | 0 | 2 | 0 | 0 | 0 |
| 88 | RBM41 | 2 | 0 | 0 | 3 | 0 |
| 89 | RBM42 | 5 | 1 | 0 | 3 | 0 |
| 90 | RBM45 | 5 | 9 | 7 | 4 | 8 |
| 91 | RBM46 | 3 | 0 | 0 | 0 | 0 |
| 92 | RBM47 | 3 | 0 | 0 | 0 | 0 |
| 93 | RBM4B | 0 | 0 | 0 | 0 | 0 |
| 94 | RBM5 | 0 | 0 | 0 | 0 | 0 |
| 95 | RBM6 | 3 | 15 | 7 | 3 | 7 |
| 96 | RBM8A | 0 | 2 | 0 | 0 | 0 |
| 97 | RBMS1 | 0 | 0 | 0 | 0 | 0 |
| 98 | RBMS2 | 0 | 0 | 0 | 0 | 0 |
| 99 | RBMS3 | 0 | 0 | 0 | 0 | 0 |
| 100 | RC3H1 | 0 | 0 | 0 | 4 | 0 |
| 101 | SAMD4A | 0 | 0 | 0 | 0 | 0 |
| 102 | SART3 | 0 | 0 | 0 | 0 | 0 |
| 103 | SF1 | 8 | 1 | 0 | 1 | 0 |
| 104 | SFPQ | 1 | 0 | 0 | 1 | 0 |
| 105 | SNRNP70 | 0 | 0 | 0 | 0 | 0 |
| 106 | SNRPA | 1 | 4 | 1 | 0 | 6 |
| 107 | SRSF1 | 0 | 0 | 0 | 0 | 0 |
| 108 | SRSF10 | 3 | 8 | 4 | 0 | 9 |
| 109 | SRSF11 | 0 | 0 | 0 | 0 | 0 |
| 110 | SRSF2 | 1 | 16 | 5 | 0 | 14 |
| 111 | SRSF4 | 4 | 19 | 8 | 0 | 15 |
| 112 | SRSF5 | 2 | 20 | 8 | 0 | 13 |
| 113 | SRSF7 | 2 | 4 | 1 | 2 | 3 |
| 114 | SRSF8 | 2 | 10 | 4 | 0 | 7 |
| 115 | SRSF9 | 0 | 1 | 0 | 0 | 1 |
| 116 | TAF15 | 0 | 0 | 0 | 0 | 0 |
| 117 | TARDBP | 2 | 0 | 0 | 0 | 0 |
| 118 | TIA1 | 0 | 0 | 1 | 3 | 0 |
| 119 | TRA2A | 0 | 0 | 0 | 0 | 1 |
| 120 | TRNAU1AP | 3 | 0 | 0 | 1 | 0 |

| | | | | | | |
|---|---|---|---|---|---|---|
| 121 | TUT1 | 0 | 0 | 0 | 1 | 0 |
| 122 | U2AF2 | 0 | 0 | 2 | 0 | 0 |
| 123 | UNK | 0 | 0 | 0 | 0 | 0 |
| 124 | YBX1 | 3 | 17 | 7 | 1 | 15 |
| 125 | YBX2 | 6 | 17 | 7 | 1 | 17 |
| 126 | ZC3H10 | 0 | 2 | 0 | 0 | 0 |
| 127 | ZC3H14 | 0 | 0 | 0 | 0 | 0 |
| 128 | ZCRB1 | 8 | 0 | 1 | 2 | 0 |
| 129 | ZFP36 | 3 | 0 | 0 | 1 | 0 |
| 130 | ZNF326 | 0 | 0 | 0 | 1 | 0 |
| 131 | ZNF638 | 2 | 0 | 0 | 0 | 0 |

| | | | 3' UNTRANSLATED REGION | | | |
|---|---|---|---|---|---|---|
| S/N | RBPs | WT | JCat | VB | IDT | GA |
| 1 | A1CF | 15 | 4 | 0 | 2 | 0 |
| 2 | ANKHD1 | 0 | 2 | 0 | 2 | 2 |
| 3 | BOLL | 0 | 0 | 0 | 2 | 0 |
| 4 | BRUNOL4 | 0 | 0 | 0 | 0 | 0 |
| 5 | BRUNOL5 | 0 | 0 | 0 | 0 | 0 |
| 6 | BRUNOL6 | 0 | 0 | 0 | 0 | 0 |
| 7 | CELF1 | 5 | 0 | 1 | 4 | 0 |
| 8 | CNOT4 | 6 | 0 | 0 | 4 | 6 |
| 9 | CPEB1 | 7 | 0 | 0 | 0 | 0 |
| 10 | CPEB2 | 2 | 0 | 0 | 0 | 0 |
| 11 | CPEB4 | 9 | 0 | 0 | 0 | 0 |
| 12 | DAZ3 | 4 | 3 | 3 | 0 | 0 |
| 13 | DAZAP | 6 | 0 | 0 | 2 | 0 |
| 14 | EIF4G2 | 0 | 6 | 3 | 6 | 4 |
| 15 | ELAVL4 | 8 | 3 | 0 | 1 | 0 |
| 16 | ENOX1 | 0 | 0 | 0 | 1 | 0 |
| 17 | ESRP1 | 0 | 0 | 0 | 0 | 0 |
| 18 | ESRP2 | 0 | 0 | 0 | 0 | 0 |
| 19 | EWSR1 | 0 | 0 | 0 | 0 | 0 |
| 20 | FMR1 | 0 | 0 | 0 | 0 | 0 |
| 21 | FUBP1 | 4 | 0 | 0 | 0 | 0 |
| 22 | FUBP3 | 8 | 0 | 0 | 5 | 0 |
| 23 | FUS | 0 | 0 | 0 | 0 | 0 |
| 24 | FXR1 | 0 | 1 | 1 | 2 | 2 |
| 25 | FXR2 | 0 | 0 | 0 | 0 | 0 |
| 26 | G3BP2 | 0 | 0 | 1 | 0 | 0 |
| 27 | HNRNPA0 | 8 | 0 | 0 | 1 | 0 |
| 28 | HNRNPA1 | 7 | 0 | 0 | 0 | 0 |
| 29 | HNRNPA1L2 | 1 | 0 | 0 | 0 | 0 |
| 30 | HNRNPA2B1 | 4 | 3 | 0 | 0 | 0 |
| 31 | HRNPC | 8 | 0 | 0 | 0 | 0 |
| 32 | HNRNPCL1 | 9 | 0 | 0 | 0 | 0 |
| 33 | HNRNPD | 5 | 0 | 0 | 0 | 0 |
| 34 | HNRNPDL | 24 | 3 | 2 | 5 | 0 |
| 35 | HNRNPF | 1 | 0 | 0 | 0 | 0 |
| 36 | HNRNPH1 | 2 | 2 | 0 | 1 | 2 |
| 37 | HNRNPH2 | 0 | 0 | 1 | 0 | 0 |
| 38 | HNRNPK | 0 | 1 | 0 | 0 | 3 |

| | | | | | | |
|---|---|---|---|---|---|---|
| 39 | HNRNPL | 1 | 0 | 0 | 0 | 0 |
| 40 | HNRNPM | 0 | 0 | 0 | 0 | 0 |
| 41 | HNRNPU | 0 | 0 | 0 | 0 | 0 |
| 42 | HNRPLL | 1 | 0 | 0 | 0 | 0 |
| 43 | HUR | 0 | 0 | 0 | 0 | 0 |
| 44 | IGF2BP1 | 1 | 0 | 0 | 0 | 0 |
| 45 | IGF2BP2 | 10 | 0 | 1 | 2 | 0 |
| 46 | IGF2BP3 | 10 | 0 | 1 | 2 | 0 |
| 47 | ILF2 | 0 | 0 | 0 | 0 | 0 |
| 48 | KHDRBS1 | 11 | 0 | 2 | 2 | 1 |
| 49 | KHDRBS2 | 15 | 0 | 3 | 4 | 1 |
| 50 | KHDRBS3 | 18 | 0 | 3 | 4 | 1 |
| 51 | KHSRP | 3 | 0 | 0 | 0 | 0 |
| 52 | LIN28A | 1 | 3 | 0 | 0 | 2 |
| 53 | MATR3 | 3 | 0 | 1 | 1 | 0 |
| 54 | MBNL1 | 1 | 12 | 1 | 0 | 3 |
| 55 | MSI1 | 9 | 0 | 0 | 0 | 0 |
| 56 | NOVA1 | 3 | 3 | 2 | 1 | 1 |
| 57 | NUPL2 | 11 | 3 | 4 | 5 | 3 |
| 58 | PABPC1 | 11 | 3 | 7 | 9 | 4 |
| 59 | PABPC3 | 10 | 0 | 2 | 2 | 0 |
| 60 | PABPC4 | 11 | 0 | 8 | 10 | 2 |
| 61 | PABPC5 | 10 | 0 | 5 | 2 | 1 |
| 62 | PABPN1 | 11 | 3 | 3 | 3 | 3 |
| 63 | PABPN1L | 9 | 1 | 2 | 2 | 0 |
| 64 | PCBP1 | 0 | 0 | 0 | 0 | 0 |
| 65 | PCBP2 | 0 | 1 | 0 | 0 | 2 |
| 66 | PCBP3 | 3 | 0 | 0 | 0 | 0 |
| 67 | PCBP4 | 0 | 0 | 0 | 0 | 0 |
| 68 | PRR3 | 0 | 5 | 4 | 4 | 5 |
| 69 | PTB3 | 0 | 0 | 0 | 0 | 0 |
| 70 | PTBP3 | 0 | 0 | 0 | 0 | 0 |
| 71 | PUF60 | 0 | 0 | 4 | 0 | 0 |
| 72 | PUM1 | 17 | 1 | 2 | 1 | 1 |
| 73 | PUM2 | 6 | 0 | 0 | 0 | 0 |
| 74 | QKI | 0 | 0 | 0 | 0 | 0 |
| 75 | RALY | 1 | 0 | 0 | 0 | 0 |
| 76 | RBFOX1 | 0 | 0 | 0 | 0 | 0 |
| 77 | RBFOX2 | 0 | 0 | 0 | 0 | 0 |
| 78 | RBFOX3 | 0 | 0 | 0 | 0 | 0 |
| 79 | RBM15B | 7 | 0 | 1 | 4 | 0 |

| | | | | | | |
|---|---|---|---|---|---|---|
| 80 | RBM22 | 0 | 1 | 0 | 0 | 5 |
| 81 | RBM23 | 0 | 2 | 0 | 0 | 1 |
| 82 | RBM24 | 7 | 0 | 0 | 2 | 3 |
| 83 | RBM25 | 0 | 0 | 0 | 0 | 0 |
| 84 | RBM28 | 4 | 0 | 1 | 0 | 0 |
| 85 | RBM3 | 0 | 0 | 0 | 2 | 0 |
| 86 | RBM38 | 0 | 0 | 0 | 0 | 0 |
| 87 | RBM4 | 0 | 6 | 0 | 4 | 0 |
| 88 | RBM41 | 5 | 0 | 2 | 1 | 0 |
| 89 | RBM42 | 0 | 1 | 1 | 0 | 0 |
| 90 | RBM45 | 1 | 11 | 1 | 4 | 0 |
| 91 | RBM46 | 0 | 0 | 1 | 2 | 0 |
| 92 | RBM47 | 8 | 0 | 0 | 2 | 3 |
| 93 | RBM4B | 1 | 2 | 0 | 2 | 0 |
| 94 | RBM5 | 0 | 2 | 0 | 0 | 2 |
| 95 | RBM6 | 4 | 12 | 0 | 2 | 6 |
| 96 | RBM8A | 0 | 5 | 0 | 2 | 0 |
| 97 | RBMS1 | 5 | 0 | 0 | 3 | 0 |
| 98 | RBMS2 | 4 | 0 | 0 | 3 | 0 |
| 99 | RBMS3 | 9 | 0 | 1 | 6 | 0 |
| 100 | RC3H1 | 10 | 0 | 0 | 2 | 0 |
| 101 | SAMD4A | 0 | 0 | 0 | 0 | 0 |
| 102 | SART3 | 11 | 3 | 7 | 9 | 4 |
| 103 | SF1 | 10 | 3 | 1 | 0 | 0 |
| 104 | SFPQ | 15 | 2 | 2 | 1 | 0 |
| 105 | SNRNP70 | 0 | 1 | 1 | 0 | 0 |
| 106 | SNRPA | 0 | 14 | 6 | 6 | 12 |
| 107 | SRSF1 | 1 | 0 | 2 | 0 | 0 |
| 108 | SRSF10 | 2 | 5 | 4 | 4 | 8 |
| 109 | SRSF11 | 0 | 0 | 0 | 0 | 0 |
| 110 | SRSF2 | 0 | 7 | 3 | 2 | 3 |
| 111 | SRSF4 | 0 | 7 | 0 | 0 | 6 |
| 112 | SRSF5 | 0 | 15 | 2 | 0 | 5 |
| 113 | SRSF7 | 1 | 4 | 1 | 2 | 5 |
| 114 | SRSF8 | 0 | 3 | 0 | 0 | 0 |
| 115 | SRSF9 | 0 | 2 | 3 | 0 | 1 |
| 116 | TAF15 | 0 | 0 | 0 | 0 | 0 |
| 117 | TARDBP | 4 | 0 | 0 | 1 | 4 |
| 118 | TIA1 | 6 | 0 | 0 | 0 | 0 |
| 119 | TRA2A | 0 | 3 | 2 | 2 | 3 |
| 120 | TRNAU1AP | 13 | 0 | 1 | 1 | 0 |

| | | | | | | |
|---|---|---|---|---|---|---|
| 121 | TUT1 | 1 | 0 | 0 | 0 | 0 |
| 122 | U2AF2 | 3 | 0 | 0 | 0 | 0 |
| 123 | UNK | 8 | 0 | 0 | 4 | 2 |
| 124 | YBX1 | 0 | 9 | 3 | 0 | 5 |
| 125 | YBX2 | 0 | 11 | 3 | 0 | 4 |
| 126 | ZC3H10 | 0 | 3 | 1 | 1 | 1 |
| 127 | ZC3H14 | 1 | 0 | 0 | 0 | 0 |
| 128 | ZCRB1 | 11 | 0 | 0 | 2 | 0 |
| 129 | ZFP36 | 6 | 1 | 0 | 0 | 0 |
| 130 | ZNF326 | 0 | 0 | 0 | 6 | 0 |
| 131 | ZNF638 | 0 | 0 | 0 | 1 | 0 |
